# Recurrent but short-lived duplications of centromeric proteins in holocentric *Caenorhabditis* species

**DOI:** 10.1101/2022.03.31.486469

**Authors:** Lews Caro, Pravrutha Raman, Florian A. Steiner, Michael Ailion, Harmit S. Malik

**Author notes:** These authors contributed equally (Lews Caro, Pravrutha Raman). Correspondence to be addressed to: Harmit Singh Malik, 1100 Fairview Avenue N. A2-205, Seattle, WA 98109 USA.

## Abstract

Centromeric histones (CenH3s) are essential for chromosome inheritance during cell division in most eukaryotes. *CenH3* genes have rapidly evolved and undergone repeated gene duplications and diversification in many plant and animal species. In *Caenorhabditis,* two independent duplications of *CenH3* (named *hcp-3* for *<u>H</u>olo<u>C</u>entric chromosome-binding <u>P</u>rotein 3*) have been previously identified: in *C. elegans* and *C. remanei*. Here, using phylogenomic analyses in *Caenorhabditis,* we find strict retention of the ancestral *hcp-3* gene and eight additional independent *hcp-3* duplications, most of which are only found in one or two species. *hcp-3L* (*hcp-3-like*) paralogs are expressed in both sexes (males and females/ hermaphrodites) and have a conserved histone fold domain. We identified novel N-terminal protein motifs, including putative kinetochore protein-interacting motifs and a potential separase cleavage site, which are well-conserved across *Caenorhabditis* HCP-3 proteins. Other N-terminal motifs vary in their retention across paralogs or species, revealing potential sub-functionalization or functional loss following duplication. *C. afra* encodes an unprecedented protein fusion, where the *hcp-3* paralog fused to duplicated segments from *hcp-4* (nematode CENP-C). Extending our analyses beyond CenH3, we found gene duplications of six inner and outer kinetochore genes in *Caenorhabditis*, including co-retention of different kinetochore protein paralogs in a few species. Our findings suggest that centromeric protein duplications occur frequently in *Caenorhabditis* nematodes, are selectively retained under purifying selection but only for short evolutionary periods, then degenerate or are lost entirely. We hypothesize that unique challenges associated with holocentricity in *Caenorhabditis* may lead to this rapid ‘revolving door’ of kinetochore protein paralogs.

## Introduction

The faithful inheritance of genetic material is indispensable for all life. In most eukaryotes, faithful inheritance of chromosomes relies on the centromeric histone H3 variant (*CenH3*) to attach chromosomes to microtubules. CenH3 acts both as a structural component of the multi-subunit complex that links chromosomes to microtubules for segregation, and as the epigenetic mark that defines and maintains the centromeric location(s) on chromosomes (Allshire and Karpen 2008; De Wulf and Earnshaw 2008; Fukagawa and Earnshaw 2014; McKinley and Cheeseman 2016; Ali-Ahmad and Sekulić 2020; Mellone and Fachinetti 2021). CenH3 is critical for chromosome segregation during mitosis and meiosis. Mutations or misregulation of CenH3 have severe consequences for fertility and viability in many species (Stoler, et al. 1995; Buchwitz, et al. 1999; Howman, et al. 2000; Blower and Karpen 2001). CenH3 is therefore expected to be conserved across eukaryotes and expected to evolve under strong evolutionary constraints to maintain functionality.

Despite this expectation for strong conservation, *CenH3* genes have rapidly evolved in animal and plant species (Malik and Henikoff 2001; Talbert, et al. 2004; Schueler, et al. 2010). This rapid evolution is hypothesized to result from a unique genetic conflict that stems from asymmetric female meiosis in animals and plants, in which only one of four meiotic products gets selected to be included in the oocyte nucleus. As a result of this bottleneck, chromosomes compete with each other for inclusion into the egg in a process termed ‘centromere drive’ (Henikoff, et al. 2001; Malik 2009; Schueler, et al. 2010; Lampson and Black 2017). This competition favors changes in centromeric DNA that result in over-recruitment of centromeric proteins (Chmátal, et al. 2014; Akera, et al. 2017; Iwata-Otsubo, et al. 2017). Conversely, genes encoding centromeric proteins evolve rapidly to blunt the ‘selfish advantage’ of cheating centromeres to restore parity and ameliorate the deleterious effects of centromere-drive (Kumon, et al. 2021). Thus, in many animal and plant species, CenH3 proteins evolve rapidly despite being essential for mitotic fidelity.

CenH3 proteins can also function differently during meiotic and mitotic segregations. Some plant *CenH3* mutants only show defects during meiosis, but not mitosis (Lermontova, et al. 2011; Ravi, et al. 2011; Schubert, et al. 2014). Conflicting evolutionary selective pressures on *CenH3* between these functions (*e.g*., mitotic versus meiotic, conserved versus rapidly evolving) could be resolved by gene duplication, which allows the duplicate (paralog) and ancestral genes to specialize in different functions (Gallach and Betrán 2011). Indeed, *CenH3* genes have also undergone repeated gene duplications not just in plant but also in several animal species including cows, fruit flies, mosquitoes, and nematodes (Li and Huang 2008; Zedek and Bureš 2016; Kursel and Malik 2017; Ishii, et al. 2020; Kursel, et al. 2020; Despot-Slade, et al. 2021; Elisafenko, et al. 2021; Kursel, et al. 2021). Cytological evidence in *Drosophila virilis* suggests that divergent *CenH3* paralogs can acquire separate, tissue-specific functions (Kursel, et al. 2021).

Although *CenH3* has undergone duplication and diversification in *Drosophila* and mosquito species, four genera of insects have completely lost *CenH3* (Drinnenberg, et al. 2014). *CenH3* loss appears to correlate with transitions from monocentricity, in which centromeric determinants are concentrated in one genomic region, to holocentricity, in which centromeres are dispersed along the length of their chromosomes. Thus, holocentricity may impose unique selective pressures that shape the path of *CenH3* and kinetochore evolution (Marques and Pedrosa-Harand 2016; Cortes-Silva, et al. 2020; Senaratne, et al. 2022; Wang, et al. 2022).

In contrast to holocentric insects, *CenH3* homologs are present in other holocentric animal and plant species (Drinnenberg, et al. 2014). Moreover, several nematode clades encode duplications and diversification of *CenH3* genes (Despot-Slade, et al. 2021). Holocentric chromosome segregation in nematodes has been best studied in *C. elegans*, which encodes two *CenH3* paralogs. The first of these to be characterized was *hcp-3*, which encodes a protein required for recruiting all other kinetochore proteins and is essential for embryonic mitotic divisions in *C. elegans* (Buchwitz, et al. 1999; Oegema, et al. 2001). However, HCP-3 appears to be dispensable for oocyte meiotic segregation (Monen, et al. 2005). A second *CenH3* paralog in *C. elegans*, CPAR-1, shares high sequence similarity to HCP-3 in the histone fold domain but is diverged in the N-terminal domain (Monen, et al. 2015). Although CPAR-1 is enriched in meiotic chromosomes, it does not appear to play an essential role in meiosis or mitosis. Indeed, it does not appear to localize to centromeres at all, and its precise function is not well understood (Gassmann, et al. 2012; Monen, et al. 2015). An independent *hcp-3* duplication occurred in a related species, *C. remanei* (Monen, et al. 2015), but its function is also unknown.

The growing collection of *Caenorhabditis* species and their genome sequences (Stevens, et al. 2019) (unpublished genomes at http://caenorhabditis.org/) provides a rich dataset for identifying the evolutionary trajectory of their *CenH3* genes. Taking advantage of this resource, we performed detailed phylogenomic analyses to understand the evolution of *CenH3* genes in *Caenorhabditis*. Our studies reveal that thirteen out of thirty-two *Caenorhabditis* species encode two or more *CenH3* paralogs, which were the result of at least ten independent duplication events. We confirm these paralogs are expressed in both sexes in representative species. We identify novel, conserved protein motifs within the N-terminal domains of *Caenorhabditis* CenH3 proteins that are likely important for interactions with other kinetochore proteins and for centromere biology. Although some motifs are strictly retained, others display variable instances of loss and retention between ancestral and duplicate genes, revealing clues to their sub-functionalization. In a possible case of neofunctionalization, we find an unusual *CenH3* paralog in *C. afra* that encodes a CENP-C-CenH3 fusion protein. Extending our analyses beyond *CenH3*, we find independent duplications of other inner and outer kinetochore proteins, revealing a remarkable pace of diversification of the kinetochore within *Caenorhabditis* nematodes. Our analyses thus reveal an unusual ‘revolving door’ of CenH3 protein duplications, with retention only over short evolutionary periods, in contrast to the strict, long-lived retention of paralogs seen in *Drosophila*, mosquito, and plant species. We hypothesize that this pattern may result from the unusual mechanisms of centromere establishment and inheritance in holocentric organisms.

## Results

### *hcp-3* has duplicated at least ten independent times in *Caenorhabditis*

Global efforts to isolate and sequence *Caenorhabditis* species have recently resulted in several well-assembled genomes from highly diverged species (Stevens, et al. 2019) (unpublished genomes at http://caenorhabditis.org/). We used this resource for phylogenomic analyses of *CenH3* evolution. We used *C. elegans* HCP-3 as a query for tBLASTn searches against genome sequences from 32 *Caenorhabditis* species (Altschul, et al. 1990; Altschul, et al. 1997; Stevens, et al. 2019) (http://caenorhabditis.org/) to identify all *hcp-3* homologs (*hcp-3*-like) genes. All putative *hcp-3* homologs (Supplementary Data S1) and their syntenic location (surrounding genes) were recorded (Figure 1). Core histone H3 and H3 variant genes were also obtained in these analyses but were easily distinguished because of their high homology to each other. Since our focus was on putative *hcp-3* orthologs and paralogs, we ignored both highly conserved core histone H3 and H3 variant proteins, as well as species-specific instances of highly diverged H3-like genes such as *F20D6.9* (also referred to as *D6H3*) from *C. elegans* (Henikoff, et al. 2000; Delaney, et al. 2018).

**Figure 1.**
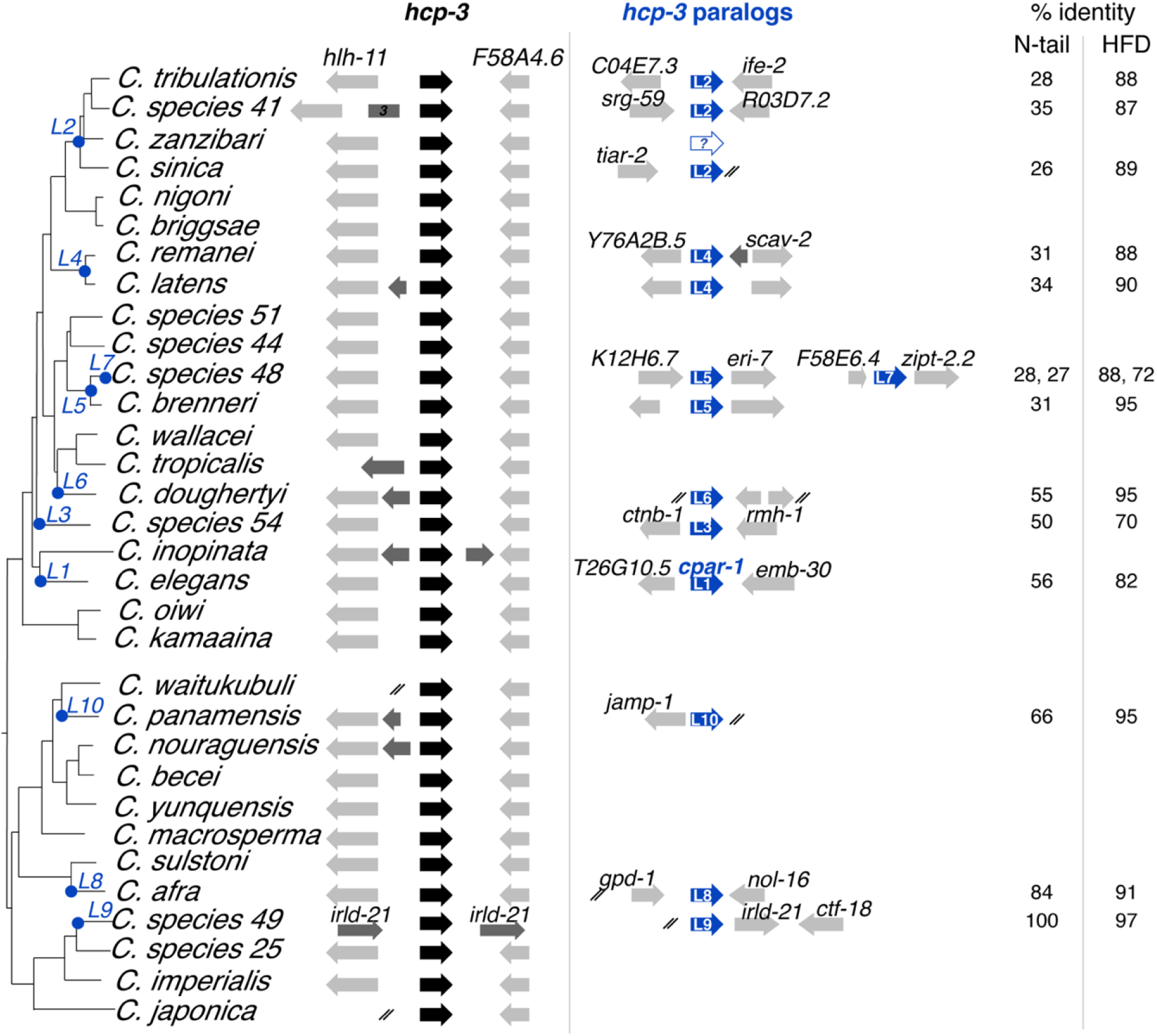
Identification and phylogenomic analysis of ancestral and duplicate *hcp-3* genes in *Caenorhabditis* species. A schematic representation of centromeric histone genes (*hcp-3*, black) and their duplicates (*hcp-3L*, blue) are shown alongside a *Caenorhabditis* species tree. *hcp-3* duplication events are represented on the species tree with a blue dot and numbered *L1* through *L10*, with paralogs arising from independent duplications assigned different numbers. Genes in the syntenic neighborhood near *hcp-3* and *hcp-3L* are represented in grey and labelled with their orthologous gene names in *C. elegans*. In some cases, 1-3 genes were inserted between *hlh-11* and *F58A4.6* within the syntenic neighborhood of *hcp-3*. White arrow with question mark represents a potential loss of *hcp-3L2* in *C. zanzibari*. Ends of genomic scaffolds are denoted with two slashes. Percent amino-acid identity between the paralog and ancestral *hcp-3* of each species (in the N-terminal tail and histone fold domain (HFD)) are shown.

Unlike holocentric insects (Drinnenberg, et al. 2014), we found that *hcp-3* orthologs are strictly retained in all *Caenorhabditis* species. They are found in shared syntenic locations, between genes homologous to *C. elegans hlh-11* and *F58A4.6*, in 28 of 32 species (Figure 1). For three of the four remaining species, at least partial synteny is maintained downstream of *hcp-3* (genes *F58A4.6, pri-1*, and *bbs-4*) but upstream synteny is either not maintained (in *C. tropicalis*) or cannot be discerned due to short genomic scaffolds (*C. waitukubuli* and *C. japonica*) (Figure 1). Only in *C. species 49 (C. sp49*) was *hcp-3* not found in this shared syntenic locus. Based on its presence in the ancestral locus in its sister species *C. sp25* and all other species, we infer that this movement of *hcp-3* is specific to *C. sp49. C. sp49* encodes two *CenH3* paralogs, both found in new syntenic loci that are not shared with sister species. We arbitrarily assign one homolog as *hcp-3* and the other as *hcp-3L9* (further explained below).

In addition to *hcp-3* orthologs, we found that thirteen out of thirty-two examined species encode at least one additional *hcp-3*-like sequence. We refer to these paralogs as *“hcp-3L”* genes (for *<u>hcp-3 L</u>ike*) (Figure 1). These *hcp-3L* genes include previously reported *hcp-3* duplications in *C. remanei* and *C. elegans* (Monen, et al. 2005; Monen, et al. 2015), which we refer to as *hcp-3L4* and *hcp-3L1*, respectively. We also identified one additional *hcp-3L* paralog in *C. tribulationis, C. sp41, C. sinica, C. latens, C. brenneri, C. doughertyi, C. sp54, C. panamensis, C. afra*, and *C. sp49*, and two, independent *hcp-3L* paralogs in *C. sp48*. All *hcp-3L* genes encode proteins with conserved Histone Fold Domains (HFD) (See supplementary Data), which are between 70-97% identical to the HFD of HCP-3 from the same species (Figure 1). In contrast, their N-terminal domains show high divergence from HCP-3 orthologs (26-100% identical, Figure 1). This pattern is consistent with overall trends of *CenH3* evolution, where the HFDs are more evolutionarily constrained due to interactions with other histones, whereas the N-terminal domains can be so divergent that they cannot even be reliably aligned across different lineages (Malik and Henikoff 2001).

We next used a combination of syntenic and phylogenetic analyses to determine whether *hcp-3L* paralogs were shared between different species, which would indicate their functional co-retention with *hcp-3* orthologs for long evolutionary periods. The highly divergent N-terminal tail sequences of *hcp-3* and their paralogs cannot be reliably aligned and can distort our interpretations. We first performed a maximum likelihood phylogenetic analysis based on an amino acid sequence alignment of the HFD (Supplementary Figure S1). We found that the protein-based phylogeny suffered from poor resolution, was unable to resolve most of the important branches and groupings of interest, and was even incongruous with the well-accepted *Caenorhadbitis* phylogeny.

Therefore, we built a maximum likelihood phylogenetic tree using a codon-based alignment of the conserved HFD cDNA sequence (Figure 2). This phylogeny is much better resolved and largely supports our inferences from the shared synteny analyses. For example, both synteny and phylogenetic analyses suggest that the duplication that gave rise to *hcp-3L4* occurred prior to the common ancestor of *C. latens* and *C. remanei* (Figure 2). Similarly, we can infer that *hcp-3L5* duplicated in the common ancestor of *C. sp48* and *C. brenneri*. In contrast, the *hcp-3L* paralogs in *C. doughertyi, C. sp54, C. elegans, C. panamensis, C. afra, C. sp49*, and the additional *hcp-3L* paralog in *C. sp48* each arose via seven independent duplications (Figure 2). In each of these seven species, the *hcp-3L* paralogs are present in unique genomic locations (Figure 1) and typically group most closely with *hcp-3* orthologs from the same species (Figure 2).

**Figure 2.**
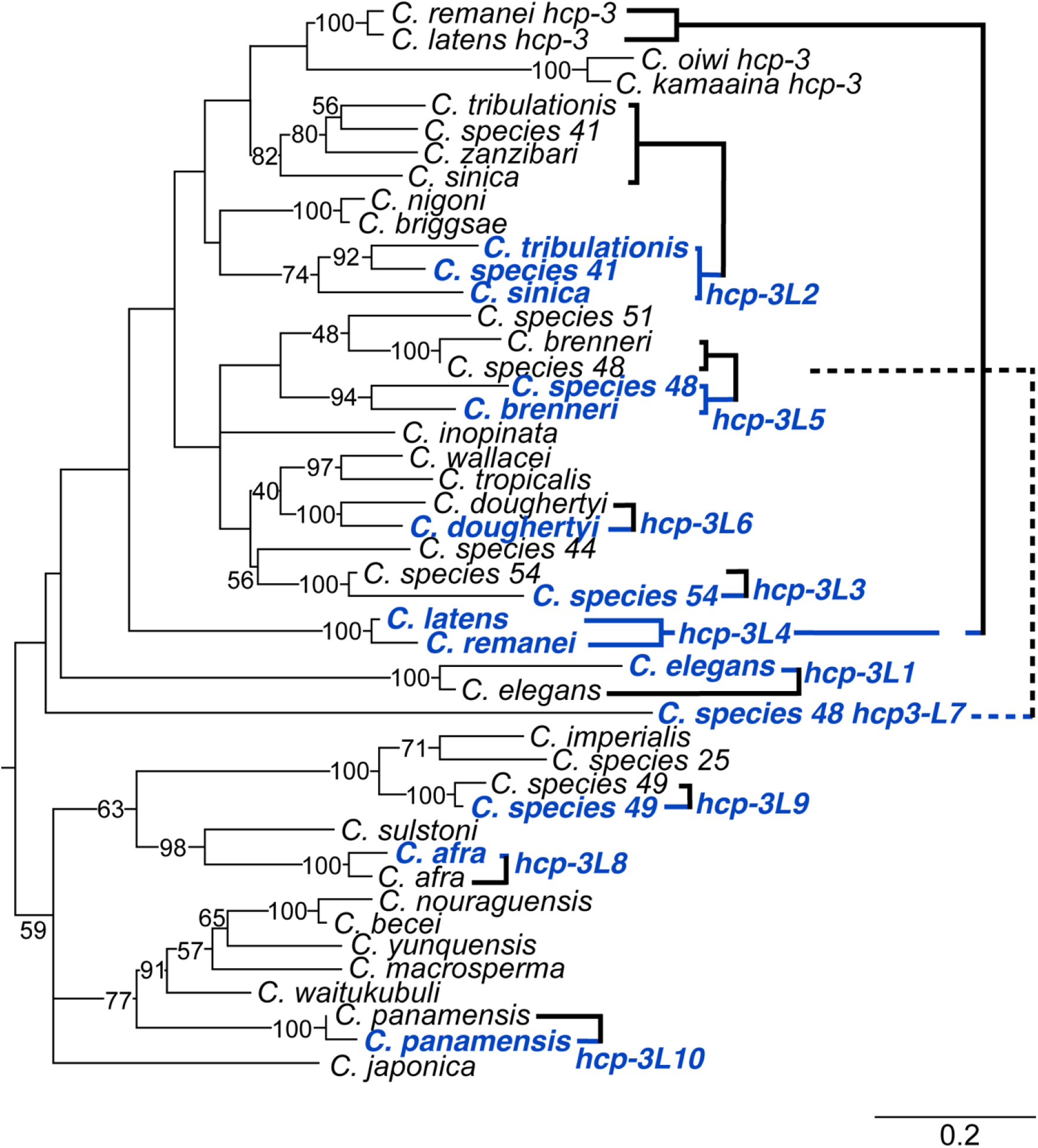
Maximum-likelihood phylogenetic tree showing relationship between *hcp-3* paralogs in *Caenorhabditis* species. A maximum likelihood tree of a DNA, codon-based alignment of the HFD of ancestral *hcp-3* (black) and *hcp-3* paralogs (blue) is shown. Bootstrap values of 40 and above are indicated. Branch lengths are scaled of number of substitutions per site (bottom-right shows scale bar).

The only discrepancy between the synteny and phylogenetic analyses was for *hcp-3L* genes found in *C. tribulationis, C. sp41* and *C. sinica*. These species are part of a group, with *C. sinica* believed to be an outgroup to *C. tribulationis, C. sp41*, and *C. zanzibari*. Different genomic locations of *hcp-3L* duplicates between *C. tribulationis, C. sp41 and C. sinica* (Figure 1) would suggest that the duplications are the result of independent duplication events. However, the small size of *C. sinica* genomic scaffolds weaken any claim of lack of synteny. Moreover, our phylogenetic analyses group *hcp-3L* genes from these species together with a high degree of confidence (Figure 2), suggesting that *hcp-3L2* is the result of a single duplication event, followed by transposition of this gene to a new locus in *C. sinica*.

Based on this inference, we infer that the absence of *hcp-3L2* in *C. zanzibari* could be the result of gene loss although there is no evidence of *hcp-3L* loss in any other species. An alternative possibility is that *C. zanzibari* may be ancestral to *C. tribulationis* and *C. sinica* for the *hcp-3L* syntenic location, in contrast to the accepted species phylogeny, and may have never acquired a *hcp-3L* paralog. Recent studies have revealed a widespread role for introgression or incomplete lineage sorting, leading to different genomic locations having vastly different evolutionary histories (Hobolth, et al. 2011; Mailund, et al. 2014; Ginsberg, et al. 2019; Suvorov, et al. 2022). Thus, it is formally possible that this *C. zanzibari* never acquired *hcp-3L2*. However, based on the well resolved species phylogeny of this quartet of species, we favor the first possibility that *C. zanzibari* acquired, then lost *hcp-3L2*.

We examined the expression of *hcp-3* and *hcp-3L* genes across representative *Caenorhabditis* species. We used RT-PCR analyses using specific primers on template RNA collected from a mixed population of males and females or hermaphrodites at various larval stages (see Methods). All analyzed species expressed both ancestral and duplicate *hcp-3* genes (Figure 3, Supplementary Figure S2), adding support to their functional retention. Since some *Drosophila CenH3* paralogs are specifically enriched in testes (Kursel and Malik 2017), we also investigated whether *Caenorhabditis hcp-3L* genes have sex-restricted expression. We performed RT-PCR on RNA collected from L4/young adult males or from L4/young adult hermaphrodites or females. Unlike *Drosophila CenH3* paralogs, we did not find sex-restricted expression of any *hcp-3L* genes (Figure 3, Supplementary Figure S2); instead, they appear to be expressed in both sexes.

**Figure 3.**
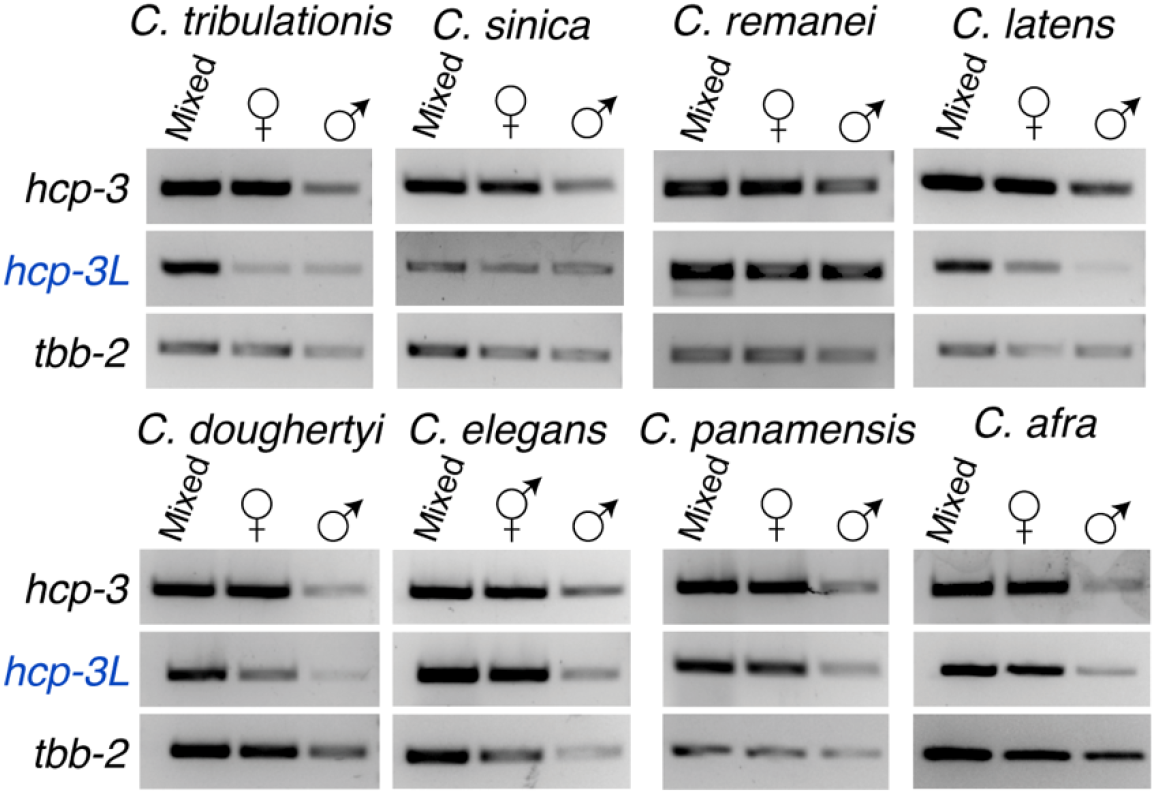
Expression analysis of ancestral and duplicate *hcp-3* genes. RT-PCR of ancestral *hcp-3* (top), *hcp-3L* (middle), or *tbb-2* (bottom; loading control) in species with *hcp-3* duplicates. RNA from a mixed worm population of various larval stages, L4 or young adult females/hermaphrodites or L4 or young adult males were used.

Overall, our analyses reveal that *hcp-3* has duplicated at least ten independent times within *Caenorhabditis* species. In contrast to plants, *Drosophila*, and mosquito species (Kursel and Malik 2017; Kursel, et al. 2020; Kursel, et al. 2021), we observed only a few cases of *hcp-3L* paralogs that are shared across two or three *Caenorhabditis* sister species, although this may partly reflect density of species sampling in these different taxonomic groups. Our findings suggest that most of the *hcp-3L* paralogs we have found are relatively young, if the relative ages of *Caenorhabditis* and *Drosophila* species analyzed are comparable (Cutter 2008).

### Motif retention and loss in the N-terminal region of HCP-3 and HCP-3L proteins

Although CenH3 proteins all have a relatively conserved HFD, their N-terminal tails are often so divergent that they cannot be aligned nor even be considered homologous across different lineages (Malik and Henikoff 2001). We took advantage of our comprehensive identification of *hcp-3* and *hcp-3L* paralogs to identify conserved motifs in the N-terminal tails of CenH3 proteins in *Caenorhabditis* species. Such a methodology represents a powerful, alignment-independent means to identify important conservation of motifs even in highly divergent protein domains. Similar analyses have identified N-terminal motifs in CenH3 proteins from many other lineages including *Drosophila*, mosquitos, and plants (Maheshwari, et al. 2015; Kursel and Malik 2017; Kursel, et al. 2020). Although the previously identified N-terminal tail motifs are often highly conserved within a lineage, they are not conserved across different lineages, suggesting that these motifs might represent lineage-specific interaction motifs between CenH3 and other kinetochore proteins.

Recent studies show that the N-terminal tail of *C. elegans* HCP-3 interacts with the inner kinetochore protein KNL-2 via a predicted structured region (de Groot, et al. 2021; Prosée, et al. 2021). This interaction between KNL-2 and HCP-3 is necessary for the establishment of centromeres in the hermaphrodite germ line, prior to the first embryonic mitosis (Prosée, et al. 2021). However, previous studies have been unable to delineate protein motifs in the HCP-3 N-terminal tail owing to its rapid divergence across species. To define putative motifs in CenH3 proteins and determine their retention across species and across paralogs, we performed motif analysis on HCP-3 and HCP-3L sequences from *Caenorhabditis* species as previously described (Kursel and Malik 2017). We identified motifs *de novo* by using MEME suite software (Bailey, et al. 2015) for all *Caenorhabditis* species containing only a single *hcp-3* gene, *i.e*., lacking a *CenH3* duplicate. We reasoned that *CenH3* genes present in a single copy are more likely to have retained all motifs essential for their functions. Using this analysis, we identified 13 motifs within HCP-3 (Figure 4A), numbered sequentially from the N-terminus. The C-terminal motifs 12 and 13 map to the HFD and are expectedly present in all HCP-3 and HCP-3L proteins, except for HCP-3L3 from *C. sp54*, which has the most divergent HFD.

**Figure 4.**
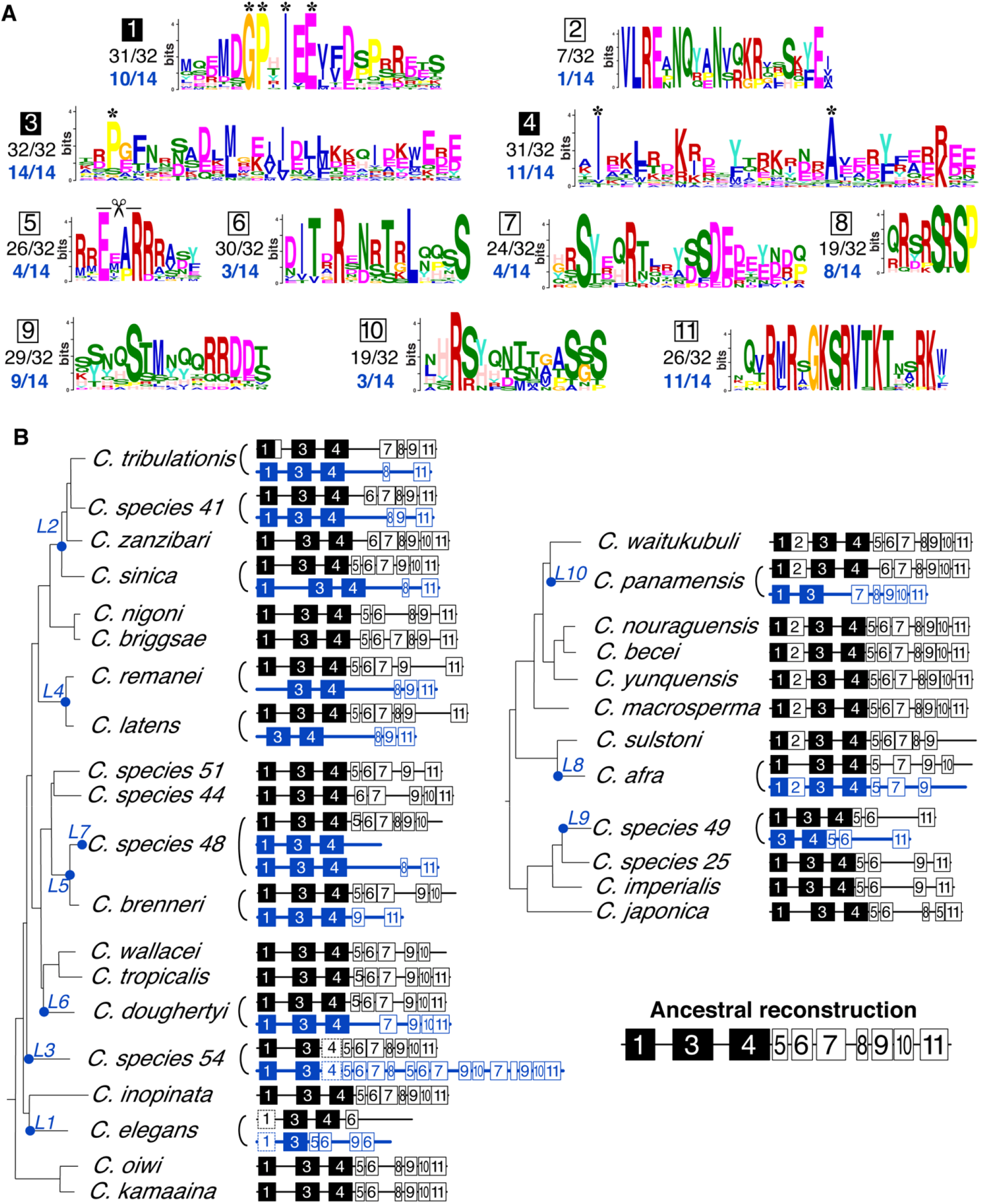
Loss or retention of N-terminal protein motifs in HCP-3 and HCP-3L paralogs. (A) Logo plots of eleven protein motifs within HCP-3 N-terminal tails discovered from an analysis of *Caenorhabditis* species without duplications. Asterisks above motif logo plots in Motifs 1, 3, and 4 indicate residues that are highly conserved within the motif. Proportion of all 32 ancestral HCP-3 proteins or 14 HCP-3L duplicates that have retained the motifs are shown. (B) *Caenorhabditis* species tree with schematics of protein motifs that are present (numbered boxes, empty boxes correspond to Motif 8) in ancestral HCP-3 (black) or HCP-3L (blue) in each species is shown. Motifs represented by dashed boxes indicates the motif’s presence was not detected by unsupervised MAST searches but was subsequently ascertained through manual alignments. All proteins contained a conserved, C-terminal HFD (not shown). Filled black boxes represent three motifs that show the highest retention in *Caenorhabditis* HCP-3 proteins. Light shading is used to indicate motif 8 which is too short to enumerate like the other motifs. A structure of the N-terminal tail of HCP-3 in the last common ancestor of *Caenorhabditis* is constructed based on the retention and loss of motifs in the N-terminal tail. L1-L10 on the species tree indicate *hcp-3* duplication events as in Figure 1.

We next examined how well N-terminal motifs 1-11 are conserved in species containing *CenH3* duplicate genes and found that these motifs varied in their evolutionary stability and conservation. Our initial unsupervised motif analysis found that motif 3 was universally conserved in all HCP-3 and HCP-3L proteins (Figure 4B) whereas motifs 1 and 4 were universally retained in at least one paralog in each species with only a few exceptions. Recognizing that this apparent ‘motif loss’ might be the result of indels or divergence of a critical conserved residue, we manually re-examined the sequences missing either motif 1 or 4. We were able to confirm the presence of motifs 1 and 4 in all species. We indicate motifs that were below the statistical threshold of the unsupervised motif analysis but subsequently identified by our manual curation by using boxes with dashed outlines (Figure 4B). Thus, following our manual curation, we found that three motifs (1, 3, and 4) are present in at least one HCP-3 paralog in all species. Notably, these motifs have not been identified in previous dissections of the N-terminal tail, highlighting the value of alignment-independent methods of important motif identification. These motifs include residues that are almost universally conserved in *Caenorhabditis* species (asterisks in Figure 4A). We predict that mutation of these residues may reveal important insight about the various functions of the HCP-3 N-terminal tail in future studies, including its interactions with kinetochore proteins such as KNL-2.

Motifs 5, 6, 7, 9, and 11 were slightly less conserved, being present in 78-94% of species. For example, motif 6 appears to be lost in both *CenH3* paralogs from *C. tribulationis* and *C. afra*, while motif 11 is not found in *C. wallacei, C. elegans* (both paralogs), and in sister species *C. sulstoni* and *C. afra* (both paralogs). Motif 5 includes a 4-amino acid segment, ExxR (Figure 4B, where x represents any amino acid), which constitutes a putative cleavage motif for the separase enzyme that initiates anaphase by cleaving the kleisin subunit of cohesin (Monen, et al. 2015). Although this previously identified motif is found in both HCP-3 and CPAR-1 in *C. elegans*, only the latter is cleaved by separase. This suggested that this motif is necessary but not sufficient for efficient separase cleavage; additional *cis*-acting determinants may be required. The separase cleavage site within motif 5 was not identified with high confidence from our motif analyses but individual alignments of HCP-3 sequences revealed that all CenH3 proteins, except the hcp-3L paralogs in *C. latens* and *C. sp48*, encode an ExxR motif (Supplementary Figure S3). Since CPAR-1 is not associated with centromeres (Gassmann, et al. 2012; Monen, et al. 2015), it is difficult to establish the significance of the separase cleavage site. For example, it is possible that the cleavage site mediates the removal of the N-terminal tail specifically for CPAR-1, thereby eliminating it from a role in germline re-establishment of centromere identity (Prosée, et al. 2021). Although it is unclear whether it is required for separase cleavage or some other function, the ExxR motif is nevertheless largely conserved in all HCP-3 proteins and most HCP-3L proteins.

In addition to many instances of motif loss in *hcp-3* genes, we found one instance of motif gain. Parsimony suggests that motif 2 was acquired in the ancestor of a clade of 8 species (Figure 4B). Importantly, after the acquisition of motif 2 in this species clade’s ancestor, the motif was not completely lost in any species. Two instances of motif 2 loss are seen: once in *C. panamensis* where it is lost in the duplicate gene but maintained in the ancestral gene, and once in *C. afra* motif 2 is lost in the ancestral gene but maintained in the paralog. Our evolutionary reconstruction suggests that some clade-specific HCP-3 protein-protein interactions or functions were acquired via motif 2 in the ancestor of these species.

In some instances, motif loss occurred in only one of the two *CenH3* paralogs from the same species. For example, most HCP-3L proteins lack motif 7 whereas ancestral HCP-3 in the same species usually contained this motif. Similarly, in species containing motif 5 and/or 6, the *hcp-3L* gene almost always lost these motifs while the ancestral *hcp-3* maintained them. In sister species *C. brenneri* and *C. sp48*, the converse is seen, where Motif 11 is maintained in the duplicate *hcp-3L* gene but lost in ancestral *hcp-3*. Overall, however, motif loss tends to occur more frequently in the *hcp-3L* paralog instead of the ancestral *hcp-3*. Thus, *hcp-3L* paralogs may be performing only a subset of the functions of an ancestral *hcp-3*. This asymmetric pattern of motif loss may also explain why ancestral *hcp-3* has been universally retained in all *Caenorhabditis* species, whereas *hcp-3L* paralogs are rarely present in more than two species.

### Selective constraints on *hcp-3* orthologs and *hcp-3L* paralogs

Our study represents a significant opportunity to evaluate the selective pressures imposed on CenH3 genes either as a result of holocentricity or due to their recurrent duplication. A previous analysis had concluded there was weak evidence of positive selection from an analysis of *hcp-3* sequences from 6 divergent *Caenorhabditis* species whose sequence was available at that time (Zedek and Bureš 2012). However, extremely large divergence and low number of sequences can be a source of artifact in such positive selection analyses. We, therefore, revisited this analysis using maximum likelihood methods (see Methods). We separately analyzed *hcp-3* sequences from the two deep lineages of *Caenorhabditis* species evaluated here, as well as analyzed two subsets of species from one of the lineages for which we had enough representation (Supplementary Table S1A). In every case, we found no evidence of positive selection acting on *hcp-3* genes. Since the presence of a paralog within the genome may affect the selective constraint on the ancestral *hcp-3* gene, we repeated the analysis by intentionally excluding all species that encode one or more *hcp-3L* paralogs and are therefore solely reliant on a single *hcp-3* gene (Supplementary Table S1A). Once again, we found no evidence for positive selection. Thus, in contrast to the previous study (Zedek and Bureš 2012) and in contrast to findings that *CenH3* genes from multiple other animal and plant taxa evolve under positive selection (Malik and Henikoff 2001; Talbert, et al. 2004; Schueler, et al. 2010; Finseth, et al. 2015), we find no evidence for positive selection acting on *CenH3* genes in *Caenorhabditis*. Lack of positive selection may suggest that holocentricity itself may have arisen as a means to defend against centromere drive (Malik and Henikoff 2009; Zedek and Bureš 2016), obviating the necessity for rapid evolution of centromeric proteins.

Based on their presence in few species, we infer that most of the *hcp-3L* genes we identified in *Caenorhabditis* species are relatively young. Our finding that *hcp-3L* genes bear the brunt of motif loss (Figure 4), together with previous studies that found no functional consequences of deleting *cpar-1* in *C. elegans* (Gassmann, et al. 2012; Monen, et al. 2015), raised the possibility that many *hcp-3L* genes are not functionally constrained. To address this possibility, we carried out three types of analyses. First, we examined selective constraints acting on *hcp-3* and *cpar-1* by investigating polymorphisms within natural isolates of *C. elegans* strains that have been previously sequenced (Cook, et al. 2017) (Supplementary Figure S4). We found only three synonymous (amino acid preserving) and zero non-synonymous (amino acid altering) polymorphisms in *hcp-3*. By contrast, *cpar-1* contained 2 synonymous polymorphisms (including one commonly shared between more than 25 strains) and 6 non-synonymous polymorphisms, four of which are shared among more than seven *C. elegans* strains. Some of these polymorphisms arise in otherwise conserved positions in the N-terminal tail (Figure 4, Supplementary Figure S4) or HFD, implying that they are likely deleterious for function. In addition to non-synonymous changes, we found at least two strains that may have disrupted *cpar-1* entirely, via either a frameshift or a splice site mutation. Based on this comparison, we infer that *cpar-1* is evolving under lower functional constraints than *hcp-3* in *C. elegans*, consistent with its non-essential function.

Second, we tested whether *hcp-3L* paralogs are generally evolving under less stringent functional constraints than *hcp-3* genes. For this, we calculated dN/dS values, which measures the ratio of the normalized rate of non-synonymous substitutions to synonymous substitutions. A lower dN/dS ratio is reflective of higher functional constraints, whereas a dN/dS ratio of close to 1 is reflective of lack of functional constraints for protein-coding function. We calculated dN/dS values in pairwise comparisons of the HFD of *hcp-3L* orthologs present in two distinct species: *hcp-3L4* in *C. latens* and *C. remanei, hcp-3L2* in *C. sinica* and *C. tribulationis*, and *hcp-3L5 in C. brenneri* and *C. sp48* (Supplementary Table 1B). We obtained dN/dS ratios of 0.02, 0.04, and 0.08 respectively. These values are considerably lower than 1, suggesting that all three paralogs have been retained under functional constraint for protein-coding function during the divergence of the respective *Caenorhabditis* species. Moreover, in all three cases, we found that dN/dS values for *hcp-3L* orthologs were comparable to or lower than corresponding *hcp-3* orthologs from the same species (Supplementary Table 1B). For comparison, the dN/dS values for pairwise comparisons of ancestral *hcp-3* from *C. latens/ C. remanei, C. sinica/ C. tribulationis* and *C. brenneri/ C. sp48* are 0.18, 0.02 and 0.03, respectively. Thus, unlike *cpar-1* in *C. elegans*, we find that *hcp-3L* paralogs have evolved under similar or even more stringent constraints than ancestral *hcp-3* genes at least in some *Caenorhabditis* species, which have co-retained *hcp-3L* paralogs for extended periods of time.

Given this finding, we revisited the age of the *hcp-3L* paralogs in *Caenorhabditis* species in a third analysis. Unlike dN or dN/dS values, dS values are relatively unaffected by selective constraints and provide a more reliable proxy for their divergence from *hcp-3* ancestors. We calculated the synonymous divergence (dS) between *hcp-3L* paralogs, whose closest relatives are *hcp-3* orthologs from the same species (Figure 2). These dS values range from 0.15 (for *C. afra*) to 0.74 (for *C. doughertyi*) (Supplementary Table 1B). These dS values are considerably lower than seen for *Drosophila* CenH3 paralogs from the same species (*e.g., D. virilis*). Although we lack reliable molecular clock-like estimates to convert these dS values to millions of years of divergence (Cutter 2008), the dS values are high enough to imply that a majority of these *hcp-L3* paralogs have been functionally retained for several million years, even though most of them have not been retained across multiple speciation events (Figure 1).

The overall selective pressure acting on *hcp-3L* paralogs is that of purifying selection or evolutionary constraint. However, our comparison of *hcp-3* and *hcp-3L3* from *C. sp54* revealed a dN/dS of 1.74, although this was not significantly different from the neutral expectation of dN/dS =1. Based on the phylogeny of *CenH3* HFD (Figure 2), we could infer that *C. sp44 hcp-3* is an outgroup to the two *C. sp54 CenH3* genes. We compared *C. sp44 hcp-3* to either *hcp-3* or *hcp-3L3* from *C. sp54*. These analyses revealed a lower dN/dS in a comparison between the two ancestral *hcp-3* orthologs (dN/dS =0.09) compared to that between *C. sp44 hcp-3* and *C. sp54 hcp-3L3* (dN/dS = 0.34). This implies that it is the unusual paralog, *hcp-3L3* that might have evolved under positive selection. HCP-3L3 also contains duplications of the N-terminal tail motifs and a rapidly evolving HFD (Figure 5A). Our findings suggest the possibility of incipient neofunctionalization of the *hcp-3L3* paralog in *C. sp54*.

**Figure 5.**
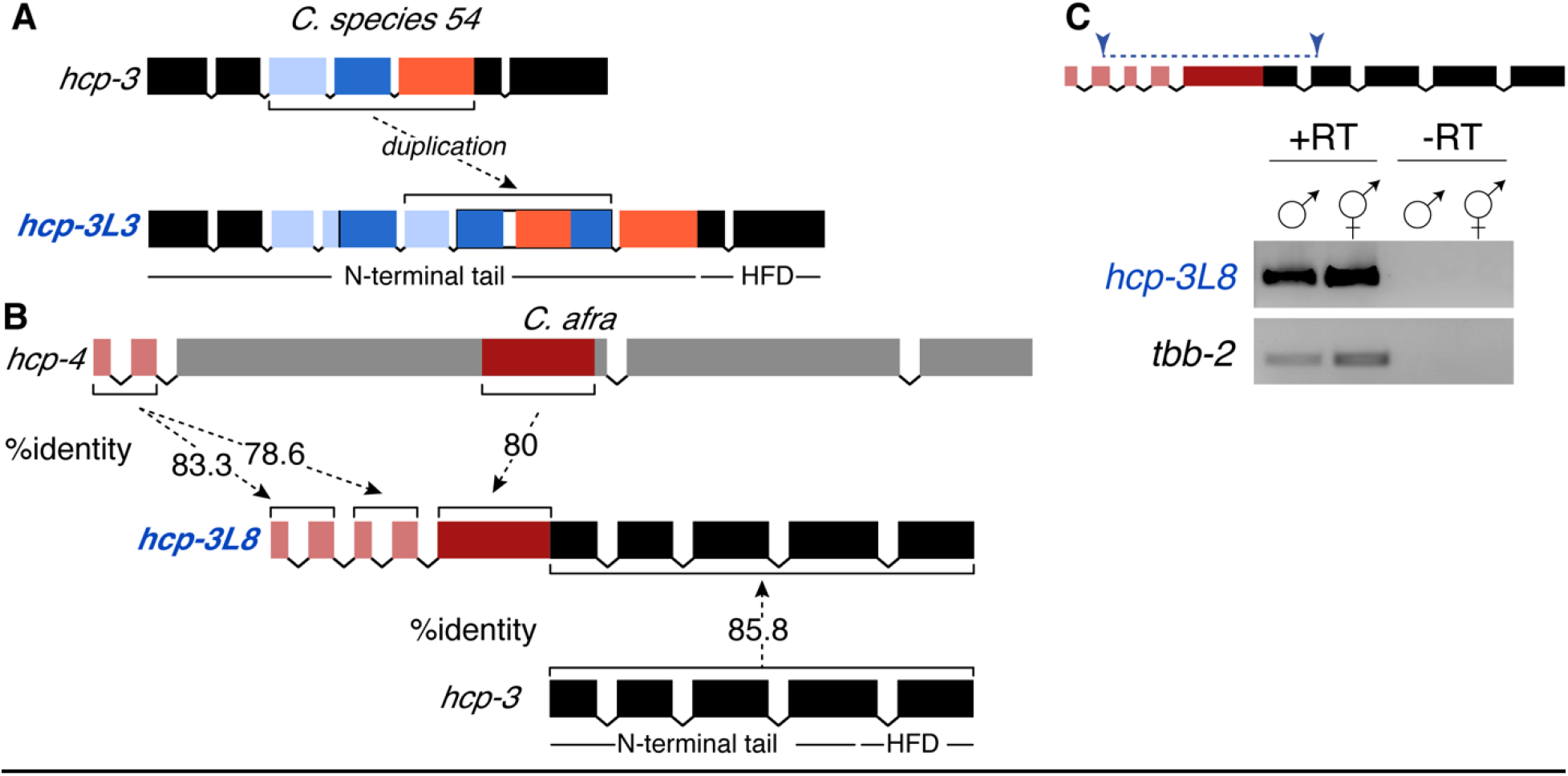
Gene structure and expression of unusual *Caenorhabditis hcp-3* paralogs. (A) Schematic of the exon structure of *C. sp54 hcp-3* (top) and *hcp-3L3* (bottom). Portions of *hcp-3* exon 3 (light blue), exon 4 (dark blue), and exon 5 (orange) are duplicated within the N-terminal tail of *hcp-3L3* (dashed arrow). The HFD is not within the duplicated region. (B) Schematic of the exon structure of *C. afra hcp-3L8* (middle) with homology to *C. afra hcp-4* (top) and *C. afra hcp-3* (bottom). The first five exons of *hcp-3L8* are homologous to *C. afra hcp-4* exons 1 and 2 (light red) as well as a portion of exon 3 (dark red). The last five exons of *hcp-3L8* are homologous to *C. afra hcp-3* (black). The HFD and the N-terminal tail of *hcp-3* are denoted. Percent amino acid identity between protein-coding exons are shown. (C) Primers designed to span exons that are homologous to *hcp-3* and *hcp-4* within *hcp-3L8* (top). Schematic of the gene shows primers used to amplify the *hcp-4-hcp-3* fusion region (top, blue) in RT-PCR of *C. afra* hcp-3L8 and tbb-2 in males and females (bottom) to confirm expression of a chimeric transcript. +RT and -RT indicate cDNA preparation with or without reverse transcriptase enzyme, respectively.

### Other centromere-localized proteins are also duplicated in *Caenorhabditis* species

In most cases, the protein sequence of HCP-3 paralogs can be confidently aligned to the ancestral HCP-3, indicating clear homology. However, aligning *C. afra* HCP-3 and *C. afra* HCP-3L8 revealed that the paralog contained an additional 198 amino acids on its N-terminus. This region was not homologous to HCP-3, but instead, was homologous to CENP-C (known as HCP-4 in *C. elegans*). HCP-4 and HCP-3 directly interact with each other in *C. elegans* (Oegema, et al. 2001) and in other eukaryotes where they are found. *C. afra hcp-3L8* contained two copies of *C. afra hcp-4* exons 1 and 2, followed by a partial copy of *hcp-4* exon 3 that is contiguous with *hcp-3-* homologous sequence (Figure 5B). We used RT-PCR to confirm that *hcp-3L8* was transcribed as a single transcript containing homology to both *hcp-4* and *hcp-3* sequences (Figure 5C). Therefore, *C. afra hcp-3L8* is a chimera of *hcp-4* and *hcp-3*. Such a fusion protein might abolish the requirement for HCP-4 to interact with HCP-3 via its C-terminal domain since the proteins are physically linked. In addition to this *hcp-4-hcp-3* fusion gene, *C. afra* also maintains its ancestral *hcp-3* and *hcp-4* genes.

Encouraged by this finding of *hcp-4* duplication and fusion with *hcp-3* in *C. afra*, we investigated whether other centromere-localized proteins have also duplicated and diversified like *hcp-3*. We performed similar paralog searches for seven kinetochore proteins (*hcp-4*, *knl-1*, *knl-2*, *him-10*, *ndc-80*, *spdl-1*, and *zwl-1*). We found an intact copy of each ancestral gene in every species (Figure 6) except for two instances where we were unable to identify full-length intact *zwl-1* genes (in *C. kamaaina* and *C. tropicalis*), likely because of sequencing error (‘#’ in Figure 6).

**Figure 6.**
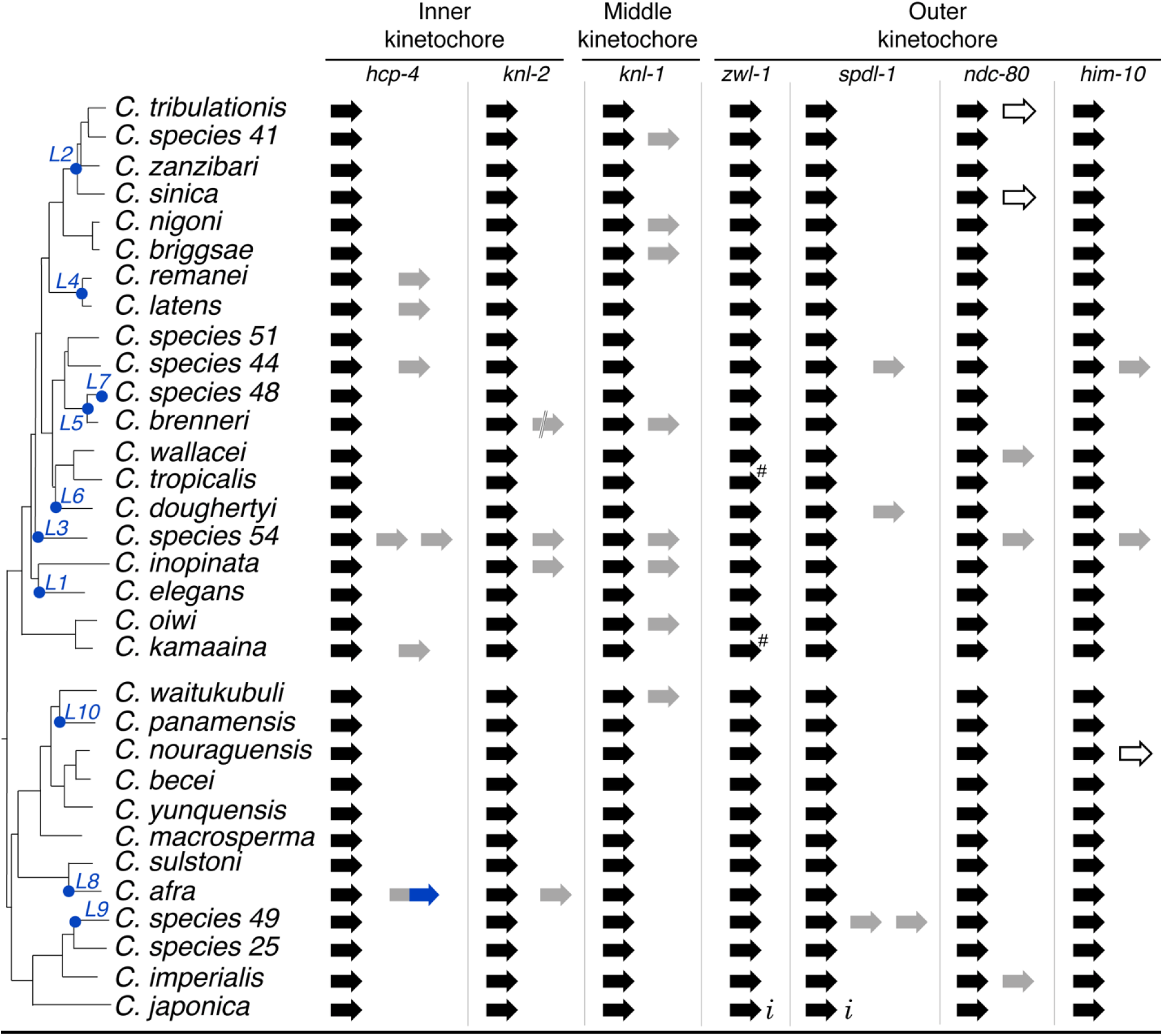
Ancestral and duplicate genes of inner and outer kinetochore proteins. A schematic representation of ancestral (black) and duplicate (grey) copies of seven kinetochore genes (*hcp-4*, *knl-1*, *knl-2*, *him-10*, *ndc-80*, *spdl-1* and *zwl-1*) alongside a *Caenorhabditis* species tree are shown. *hcp-3* duplication events are denoted as a blue dot on the species tree, as in Figure 1. The unique fusion between *C. afra hcp-4* and *hcp-3* duplicates is shown in grey and blue respectively. Incomplete sequence information in genomic scaffolds is denoted with *i* and potential pseudogenes are denoted as unfilled arrows. Double slash in *C. brenneri knl-2* duplicate indicates the sequence was split between two scaffolds. # indicates two potential pseudogenization events in *zwl-1* that are likely to represent sequencing errors.

We found instances of duplications for all kinetochore proteins except *zwl-1*. These duplications include those that appear to be functionally retained with an intact open reading frame (filled, gray arrows), or are interrupted (double lines) or show clear signs of pseudogenization (unfilled arrows) (Figure 6). In the 32 species examined, we found seven *hcp-4* duplicates, four *knl-2* duplicates (including a pseudogene in *C. brenneri*), eight *knl-1* duplicates, four *spdl-1* duplicates, five *ndc-80* duplicates (including two pseudogenes), and three *him-10* duplicates (including one pseudogene). Duplications of inner and middle kinetochore proteins were only marginally more prevalent than duplications of outer kinetochore proteins. However, we observed several instances of partial intron losses that occurred recurrently in genes encoding ancestral and paralog outer kinetochore proteins (Supplementary Figure S5). Such partial intron losses have been observed previously in plants (Roy and Penny 2007), fungi (Nielsen, et al. 2004) and in *Caenorhabditis* species (Robertson 1998; Cho, et al. 2004; Kiontke, et al. 2004). They are thought to be a result of partial retrotransposition where cDNA is used to partially overwrite the genomic locus. Overall, our analyses suggest that in addition to HCP-3, other kinetochore proteins are also undergoing duplication and diversification in *Caenorhabditis* species.

We next investigated whether any kinetochore protein paralogs have been co-retained with *hcp-3L* paralogs, which would suggest a concerted duplication and retention of multiple kinetochore proteins, consistent with significant specialization. We found that four of six independent *hcp-4* duplications coincided with retention of *hcp-3L* paralogs in the same species (Figure 6). These include an *hcp-4* paralog whose origin coincides with the *hcp-3L4* paralog in *C. latens/ C. remanei*, two *hcp-4* paralogs that cooccur with *hcp-3L3* in *C. sp54*, and the *hcp-4-hcp-3* fusion gene in *C. afra* (above).

Thus, 4 of 14 species expressing an *hcp-3L* paralog also encode a (complete or partial) *hcp-4* paralog whereas 2 of 18 species lacking *hcp-3L* paralogs encode a *hcp-4* paralog: *C. sp44* and *C. kamaaina*. However, there is no statistically significant evidence of co-retention (p=0.36), indicating that the duplication or retention of *hcp-3* and *hcp-4* paralogs may be independent.

Other kinetochore proteins analyzed also largely reflect this pattern of independent duplication. Even though KNL-2 is required to deposit HCP-3 proteins at centromeres in *Caenorhabditis* species (Maddox, et al. 2007; de Groot, et al. 2021; Prosée, et al. 2021), there does not appear to be a significant pattern of co-retention with *hcp-3L* paralogs. The one exceptional species is *C. sp54*, which encodes an *hcp-3L3* paralog, two *hcp-4* paralogs, a *knl-2* paralog, a *knl-1* paralog, an *ndc-80* paralog, and a *him-10* paralog. If the proteins encoded by these paralogs exclusively interact with each other, this species may represent an intriguing case of incipient kinetochore specialization.

## Discussion

Our comprehensive phylogenomic approach in *Caenorhabditis* nematodes uncovered two novel aspects of CenH3 evolution. First, we uncovered a detailed molecular architecture of the N-terminal tail of HCP-3 proteins (Figure 4). The HCP-3 N-terminal tail is dispensable for mitotic chromosome segregation and centromere maintenance during *C. elegans* development (Prosée, et al. 2021) but is essential in establishing a functional HCP-3 distribution in the germ line, which is maintained in the subsequent generation throughout development. At least part of this functionality of the HCP-3 N-terminal tail stems from its interactions with kinetochore proteins like KNL-2 (de Groot, et al. 2021; Prosée, et al. 2021). Thus far, however, the molecular architecture of the interactions of HCP-3 with other kinetochore proteins like KNL-2 has been only crudely defined. Like in other eukaryotic lineages, the N-terminal tail of HCP-3 proteins is much more divergent than the histone fold domain (HFD). Thus, comparisons of functional domains in CenH3 N-terminal tails between taxonomic groups or even within *Caenorhabditis* are practically impossible, exacerbating the difficulty in defining functional domains within HCP-3’s N-terminal tail. Our description of 11 motifs in HCP-3 N-terminal tails, including three that are nearly universally conserved, provide an important resource for the fine-scale dissection of the various protein-protein interactions mediated by the N-terminal tail and the functional role these interactions play in centromere biology. In particular, the three conserved motifs contain residues that are as well conserved across *Caenorhabditis* species as many HFD residues and are likely to be important for centromere re-establishment.

We propose that these N-terminal tail motifs are sites of previously proposed or novel protein-protein interactions. Consequently, motif gains or losses could indicate gains or losses of HCP-3 interactions with partner proteins. We observe one unambiguous case of motif gain — Motif 2 — in one clade of *Caenorhabditis* species. This likely represents a novel protein-protein interaction module important for *CenH3* function at least in those species. It would be interesting to test whether this interaction can be recapitulated even in species like *C. elegans*, which never acquired Motif 2. We also observe several cases of motif degeneration or loss. Unlike in *Drosophila CenH3* paralogs, we see no evidence for motif redistribution between the paralog and ancestral *hcp-3* genes, which would suggest sub-functionalization (Kursel and Malik 2017). Instead, we find that motif loss or degeneration preferentially occurs in *hcp-3L* paralogs rather than ancestral *hcp-3*, suggesting that the paralogs progressively lose ancestral functions and interactions. Since tail-less HCP-3 proteins can still function in mitosis (Prosée, et al. 2021), it is tempting to speculate that despite progressive loss of N-terminal motifs, HCP-3L paralogs could still function in mitosis.

The remarkable example of a chimeric gene in *C. afra*, where an HCP-3L protein is fused to an inner kinetochore protein, HCP-4 (CENP-C in mammals; Figure 5) exemplifies an instance where previously conserved motifs could be lost. HCP-3 and HCP-4 physically interact in many eukaryotes to format the kinetochore complex during mitosis. A fusion of these two proteins would therefore guarantee a protein-protein interaction, obviating the requirement to maintain motifs required for HCP-3L and HCP-4 interactions. This could lead to loss of HCP-3L N-terminal tail motifs required for HCP-4 association and of C-terminal HCP-4 motifs required for HCP-3 interaction.

The second major conclusion from our evolutionary analyses is the unusually rapid cadence of turnover of *hcp-3* paralogs in *Caenorhabditis* species. Nearly half of the species we analyzed contain an *hcp-3* paralog. Yet, in contrast to analyses in *Drosophila* and mosquito lineages (Kursel and Malik 2017; Kursel, et al. 2020), each *Caenorhabditis* paralog was typically acquired through independent duplication events. Most paralogs have only been retained in a single species with only one *hcp-3L* paralog being present in more than two species. Previous analyses suggest that *C. elegans* have a higher duplication rate than other species including *D. melanogaster* (Lynch and Conery 2000; Pan and Zhang 2007; Lipinski, et al. 2011), potentially as high a per gene duplication rate as 0.02 every million years (Lynch and Conery 2000). This high rate of gene duplication may account for the higher number of *hcp-3* duplications we observe in *Caenorhabditis*. However, these analyses also make clear that the vast majority of gene duplications that arise in *C. elegans* are efficiently purged by natural selection (Lipinski, et al. 2011). In contrast, our findings suggest that many *hcp-3L* paralogs are retained under purifying selection for significant periods of time.

Our evolutionary analyses thus reveal an unusual ‘revolving-door’ of *hcp-3L* paralogs in *Caenorhabditis* species. Under this regime, gene duplication is frequent, *hcp-3L* paralogs are retained under purifying selection for a significant evolutionary period, before eventually either degenerating (*e.g., cpar-1* in *C. elegans*) or being lost entirely (*e.g*., possibly *hcp-3L2* in *C. zanzibari*), returning to the ancestral state of the genome encoding only a single *hcp-3* gene. This cadence is completely unprecedented among all other taxonomic groups where *CenH3* duplications have been investigated. Even the high number of *hcp-3* duplications we have observed is likely an underestimate of the true number, since extant species represent only one evolutionary snapshot. Indeed, our study implies that many previously arising *hcp-3L* paralogs have been lost or degenerated beyond recognition during *Caenorhabditis* evolution. Although we have not evaluated all of them in the same level of detail, duplications of other kinetochore proteins also appear to evolve under a similar revolving-door dynamic.

What could account for this revolving-door *i.e*., the short-term evolutionary retention of *hcp-3L* paralogs and their long-term loss or degeneration? We consider several possibilities for what could provide the transient selective pressure to retain *CenH3* paralogs. First, this pattern could result from specialization or subfunctionalization of *CenH3* paralogs for tissue- or sex-specific expression, such as what has been seen in *Drosophila* species (Kursel, et al. 2021). However, we found no evidence that this is the case in *Caenorhabditis* regardless of whether they reproduce through male/female or male/hermaphrodite systems.

Second, specialized requirements for meiosis could explain the evolutionary pattern of *hcp-3* duplications. Unlike monocentric chromosomes, holocentric chromosomes have inherent challenges at undergoing meiosis, which have been overcome by different taxa via different means (Melters, et al. 2012). However, unlike most eukaryotes *C. elegans* chromosomes connect to the meiotic spindle by a CenH3-independent mechanism (Monen, et al. 2005). Therefore, at least in *C. elegans*, *hcp-3* is entirely dispensable for meiotic chromosome segregation (Monen, et al. 2005). This relaxes constraints to maintain meiotic functions on *hcp-3* genes but cannot explain the revolving-door pattern.

A third possible explanation for the transient retention of *hcp-3* paralogs is suppression of either ‘centromere-drive’ or ‘holokinetic drive’. Rapid diversification of centromeric DNA to exploit asymmetric meiosis (‘centromere-drive’) has been proposed to lead to both the rapid evolution of centromeric proteins such as CenH3 as well as the origin and retention of centromere-specific proteins, such as *Umbrea/HP6* in *D. melanogaster* (Ross, et al. 2013). Currently, it is unclear whether centromere drive can even occur in holocentric organisms. Indeed, transitions to holocentricity have been previously invoked as one possible means to prevent centromere drive from ever occurring (Malik 2009; Zedek and Bureš 2016). Although a previous study reported weak evidence of positive selection using an analysis of *hcp-3* from six highly diverged *Caenorhabditis* species (Zedek and Bureš 2012), our comprehensive reanalysis of *hcp-3* evolution across a much more densely-sampled series of closely-related species revealed no evidence of positive selection (Supplementary Table S1B). There is conflicting evidence of *CenH3* positive selection in some holocentric plant species (Krátká, et al. 2021), but not in others (Zedek and Bureš 2016). Asymmetric meiosis could also lead to another form of drive, leading to preferential inheritance of larger or small holocentric chromosomes (‘holokinetic drive’), which could explain the observed negative correlation between chromosome number and genome size in many holocentric lineages (Bureš and Zedek 2014). If either of these drive mechanisms occur in *Caenorhabditis* species, then *hcp-3L* paralogs could arise and be temporarily retained as drive-suppressors, but only while the driving elements were still present in the genome. As these driving elements are removed from the genome however, *hcp-3L* gene functions would be rendered superfluous, resulting in loss of these paralogs. Given the uncertainty about the status of centromere-drive or holokinetic drive in nematodes, or the role of *hcp-3L* paralogs might play in either process, we cannot comment further on the likelihood of this possibility.

Instead, we favor a fourth hypothesis, in which the holocentricity of *Caenorhabditis* species, with HCP-3 distributed along the length of the chromosomes, might itself lead to the revolving-door dynamics of centromeric proteins. CenH3 loading at holocentromeres is more plastic than at monocentromeres. Since CenH3 does not have to associate with specific sequences or chromosomal regions, holocentric chromosomes more easily tolerate chromosome breakage, fusion, or rearrangements. Indeed, even prior to clear cytological evidence, holocentric organisms were observed to maintain fertility despite radiation-induced chromosome breaks (Schrader 1935; Melters, et al. 2012). Moreover, even completely foreign DNA can form minichromosomes that assemble centromeres and be stably propagated (Zhu, et al. 2018; Lin and Yuen 2020; Lin, et al. 2021). Nevertheless, centromere distribution in holocentric organisms is not random. Although HCP-3 presence is partially linked to certain ‘hot sites’ in *C. elegans* (Steiner and Henikoff 2014), the overall pattern of centromere establishment in holocentric organisms appears to be predominantly linked to transcriptionally repressed genomic regions in the germline. This pattern of centromere definition via germline heterochromatin is seen in both *C. elegans* and in the CenH3-devoid *B. mori* (Gassmann, et al. 2012; Steiner and Henikoff 2014; Senaratne, et al. 2021).

Although such a mode of centromere definition is more tolerant of genomic rearrangements than monocentric organisms, it could also be subject to transient stress. This stress could be imposed by either chromosomal rearrangements or transposon invasion, which can quickly and dramatically alter the landscape of transcription and repression in the germline. In such circumstances, it might be advantageous to retain HCP-3L paralogs to temporarily increase the dosage of proteins required to correctly establish centromere identity, as has been proposed in some plant lineages (Evtushenko, et al. 2021). Alternatively, it may be advantageous to express HCP-3 proteins with slightly altered sequences and localization preferences, allowing restoration of optimal centromere distributions even after periods of such ‘genomic stress’. Under either scenario, eventual amelioration of the genomic stressor (*e.g*., decay or silencing of the invading transposable element) would render *hcp-3L* paralogs superfluous and these would be lost.

Thus, different *Caenorhabditis* species might represent different stages of the revolving-door process for kinetochore proteins. Species like *C. sp54*, which possess paralogs of five of seven kinetochore genes investigated, may be actively selecting for the retention and function of these paralogs. In contrast, species like *C. elegans*, with a nonessential *cpar-1* and no other kinetochore paralog, may have already overcome the need for such innovation. We, therefore, predict that functional consequences of kinetochore paralog loss in different *Caenorhabditis* species will differ based on their stage of genetic innovation. Our study underlines the need for the analysis of non-model organisms and the value of evolutionary comparisons to reveal novelties even in well-studied cellular pathways.

## Methods

### Strain maintenance

All strains were cultured on Nematode Growth Medium (NGM) plates seeded with 200 μl OP50 at 20°C using standard methods (Brenner 1974).

### Strains used

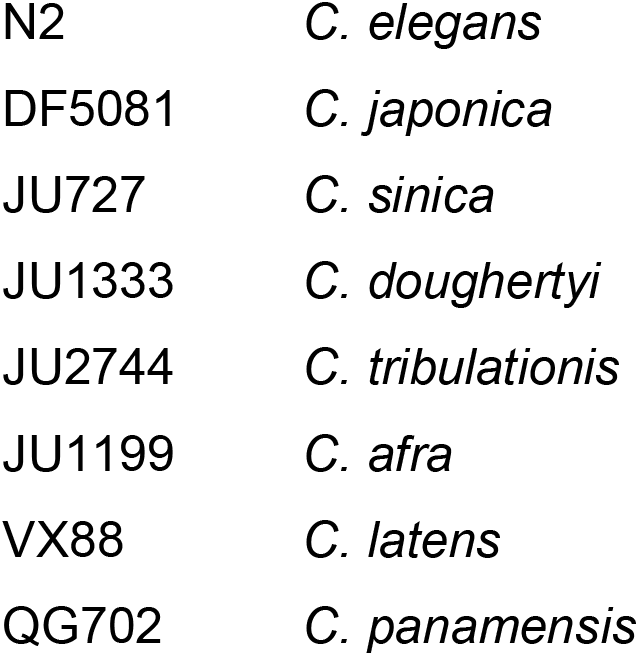

|  |  |
| --- | --- |
| N2 | *C. elegans* |
| DF5081 | *C. japonica* |
| JU727 | *C. sinica* |
| JU1333 | *C. doughertyi* |
| JU2744 | *C. tribulationis* |
| JU1199 | *C. afra* |
| VX88 | *C. latens* |
| QG702 | *C. panamensis* |

### Identification of *hcp-3* and kinetochore protein homologs in sequenced genomes

To identify *hcp-3* paralogs and orthologs we iteratively queried the assembled genomes of thirty-two *Caenorhabditis* species: *C. tribulationis*, *C. sp41*, *C. zanzibari*, *C. sinica*, *C. nigoni*, *C. briggsae*, *C. remanei*, *C. latens*, *C. sp51*, *C. sp44*, *C. sp48*, *C. brenneri*, *C. wallacei*, *C. tropicalis*, *C. doughteryi*, *C. sp54*, *C. inopinata*, *C. elegans*, *C. oiwi*, *C. kamaaina*, *C. waitukubuli*, *C. panamensis*, *C. nouraguensis*, *C. becei*, *C. yunquensis*, *C. macrosperma*, *C. sulstoni*, *C. afra*, *C. sp49*, *C. sp25*, *C. imperialis*, and *C. japonica* (Supplementary Data S2). We used TBLASTN (Altschul, et al. 1990; Altschul, et al. 1997) on each species’ genome (Stevens, et al. 2019) to perform a homology-based search starting with C. elegans HCP-3 (WBGene00001831) as our query. To ensure that we had not missed any *hcp-3* paralogs, we repeated our analyses querying each species hits on their own genome using TBLASTN and did not retrieve additional hits. To identify paralogs and orthologs of kinetochore proteins (Figure 6), we repeated this same homology-search procedure starting with *C. elegans* HCP-4 (WBGene00001832), KNL-1 (WBGene00002231), KNL-2 (WBGene00019432), ZWL-1 (WBGene00021460), SPDL-1 (WBGene00015515), NDC-80 (WBGene00003576), and HIM-10 (WBGene00001869). We used http://blast.caenorhabditis.org/ to perform all TBLASTN analyses.

Synteny was used to determine *CenH3* orthologs across *Caenorhabditis* species. We identified annotated genes immediately upstream and downstream of *hcp-3* and *hcp-3L* genes. We then used these neighboring genes as a query for TBLASTN searches of the *C. elegans* genome to identify the orthologous syntenic genes (Figure 1). Dissimilar flanking genes between *hcp-3L* orthologs provide support for their acquisition through independent *hcp-3* duplication events. In some cases, *hcp-3 or hcp-3L* genes were found in small genomic scaffolds or at the end of scaffolds, reducing our ability to identify upstream or downstream syntenic genes. In the latter case, we analyzed additional genes in the direction (upstream or downstream) that had sufficient genomic information available on the same scaffold.

### Phylogenetic Analyses

All protein and nucleotide alignments were performed using the MUSCLE algorithm (Edgar 2004) in Geneious Prime 2019.2.3 (https://www.geneious.com). We first inferred the phylogeny of the nucleotide sequences corresponding to the HFD of HCP-3 homologs using the Maximum Likelihood method and the JTT model (Jones, et al. 1992) for protein alignments and the General Time Reversible model (Nei and Kumar 2000) for nucleotide alignments. We inferred the bootstrap consensus tree from 100 replicates. Initial tree(s) for the heuristic search were obtained automatically by applying Neighbor-Joining and BioNJ algorithms to a matrix of pairwise distances estimated using the Maximum Composite Likelihood (MCL) approach, and then selecting the topology with superior log likelihood value. A discrete Gamma distribution was used to model evolutionary rate differences among sites (5 categories (+*G*, parameter = 0.8943)). The rate variation model allowed for some sites to be evolutionarily invariable ([+*/*], 26.05% sites). All positions with less than 95% site coverage were eliminated, i.e., fewer than 5% alignment gaps, missing data, and ambiguous bases were allowed at any position (partial deletion option). There were a total of 267 positions in the final dataset. Evolutionary analyses were conducted in MEGA11 (Stecher, et al. 2020; Tamura, et al. 2021).

### Motif Analyses

13 motifs were identified using MEME (Bailey, et al. 2015) on predicted, fulllength HCP-3 protein sequences from species lacking *hcp-3* paralogs (*C. zanzibari, C. nigoni, C. briggsae, C. sp51, C. sp44, C. wallacei, C. tropicalis, C. inopinata, C. oiwi, C. kamaaina, C. waitukubuli, C. nouraguensis, C. becei, C. yunquensis, C. macrosperma, C. sulstoni, C. sp25, C. imperialis, C. japonica*). E-values of all 13 discovered motifs were below 10^-5^ Motif logo plots were generated and downloaded from MEME. Presence or absence of these motifs in all HCP-3 and HCP-3L proteins was determined by using MAST (Bailey and Gribskov 1998). We considered a motif as present in a protein if the MAST P-value was below 10^-5^.

Since the N-terminal tails of HCP-3 and its paralogs are highly divergent, we were not able to identify the separase motif efficiently via motif analyses. To identify presence of the ExxR separase motif, we separately aligned each HCP-3 or HCP-3L protein sequence with *C. elegans* HCP-3 and CPAR-1 (HCP-3L1) either individually or together. This alignment was used to generate the predicted separase motifs shown in Supplementary Figure S3.

### Analysis of evolutionary selective pressures

To analyze selective pressures on *CenH3* genes, we compared rates of synonymous (dS) to nonsynonymous (dN) substitution among *hcp-3* and *hcp-3L* genes. dN and dS between all pairwise combinations of *CenH3* genes were determined using SNAP (Korber 2000) (www.hiv.lanl.gov) on a codon alignment of the histone fold domain (Supplementary Table S1A). dN/dS ratios were used to determine the selective pressures acting on *CenH3* genes.

For all tests, we generated codon alignments using MUSCLE (Edgar 2004), and manually adjusted them to improve alignments if needed. We also trimmed sequences to remove alignment gaps and segments of the sequence that were unique to only one species. We found no evidence of recombination for any of these alignments using the GARD algorithm at datamonkey.org (Kosakovsky Pond, et al. 2006). We used the alignment to generate a tree using PhyML maximum-likelihood methods with the HKY85 substitution model (Guindon, et al. 2010).

We analyzed selective pressures on *Caenorhabditis hcp-3* using the codeml algorithm from the PAML suite (Yang 1997) (Supplementary Table S1A). We generated codon alignments using MUSCLE (Edgar 2004) which we manually adjusted if needed to improve alignments. These adjustments also included manually trimming sequences to remove alignment gaps and segments of the sequence that were unique to only two or less species. These alignments were used to generate trees using PhyML maximumlikelihood methods with the HKY85 substitution model (Guindon, et al. 2010). To test whether any residues evolve under positive selection, we compared likelihoods between model 7 (which disallows dN/dS to be equal to or exceed 1) and model 8 (where there are ten classes of codons with dN/dS between 0 and 1, and an eleventh class with dN/dS > 1). To determine statistical significance, we compared twice the difference in log-likelihoods between the two models to a χ^2^ distribution with the degrees of freedom reflecting the difference in number of parameters between the models being compared (Yang 1997).

### *C. elegans* HCP-3 and CPAR-1 polymorphisms

To determine natural variation in *C. elegans hcp-3* and *cpar-1* genes (Supplementary Figure S4), we used the *Caenorhabditis elegans* Natural Diversity Resource (Cook, et al. 2017). The identified synonymous mutations in *hcp-3*, as well as the frameshift, synonymous, and non-synonymous mutations in *cpar-1* were identified by the CeNDR variant annotation feature. The *cpar-1* partial deletion was found manually by looking at whole-genome sequencing reads from *C. elegans* strain ECA740 mapped onto the N2 reference genome.

### RT-PCR

Total RNA was isolated using TRIzol (Fisher Scientific) from 50-100 L4 or young adult males, females, or hermaphrodites or from a near starved plate of mixed-stage animals. RNA was extracted by chloroform extraction, precipitated using isopropanol, washed with ethanol, and resuspended in 20μl of nuclease-free water. Next, RNA was treated with DNase I (New England Biolabs, 2 units/μl) at 37°C for 60 minutes followed by heat inactivation at 75°C for 10 minutes. DNase-treated RNA was purified using the RNA Clean and Concentrator-5 kit (Zymo Research) and converted to cDNA using SuperScript III Reverse Transcriptase (Invitrogen) using polydT primers as per manufacturer’s recommendations. RNA concentrations used to make cDNA were not kept the same between whole plate, male, and female/hermaphrodite samples except for samples from *C. afra* (in Figure 5C), *C. remanei* and *C. sinica*. PCR was done on cDNA using Phusion High-Fidelity DNA Polymerase Kit (New England Biolabs) guidelines according to the manufacturer’s recommendations using primers for *hcp-3*, *hcp-3L*, and *tbb-2*. All primer sequences used are listed in Supplementary Table S2.

## Data access

Supplementary Data files include all sequence alignments used to generate phylogenetic trees and to analyze evolutionary selective pressures and sequences of all HCP-3 and kinetochore protein orthologs and paralogs described in the paper.

## Acknowledgments

We thank Ching-Ho Chang for comments on the manuscript, the Bai lab at Fred Hutchinson Cancer Research Center for nematode resources, and Matthew Rockman for providing the *C. panamensis* strain. Some strains were provided by the *Caenorhabditis* Genetics Center (CGC), which is funded by NIH Office of Research Infrastructure Programs (P40 OD010440). This work was supported by grants from the National Institutes of Health (NIH) Institutional Training Grant (T32 GM007270 to L.C.), the Republic and Canton of Geneva (to F.A.S), the Swiss National Science Foundation (310030_197762 to F.A.S.), the National Science Foundation (NSF) CAREER Award (MCB-1552101 to M.A.), the NIH National Institute of General Medical Sciences (R01 GM074108 to H.S.M.), and from the Howard Hughes Medical Institute (to H.S.M.). The funders played no role in study design, data collection and interpretation, or the decision to publish this study. P.R. is a Washington Research Foundation Fellow and H.S.M. is an Investigator of the Howard Hughes Medical Institute.

