## Supplementary Figures and Legends for "Recurrent but short-lived duplications of centromeric proteins in holocentric *Caenorhabditis* species"

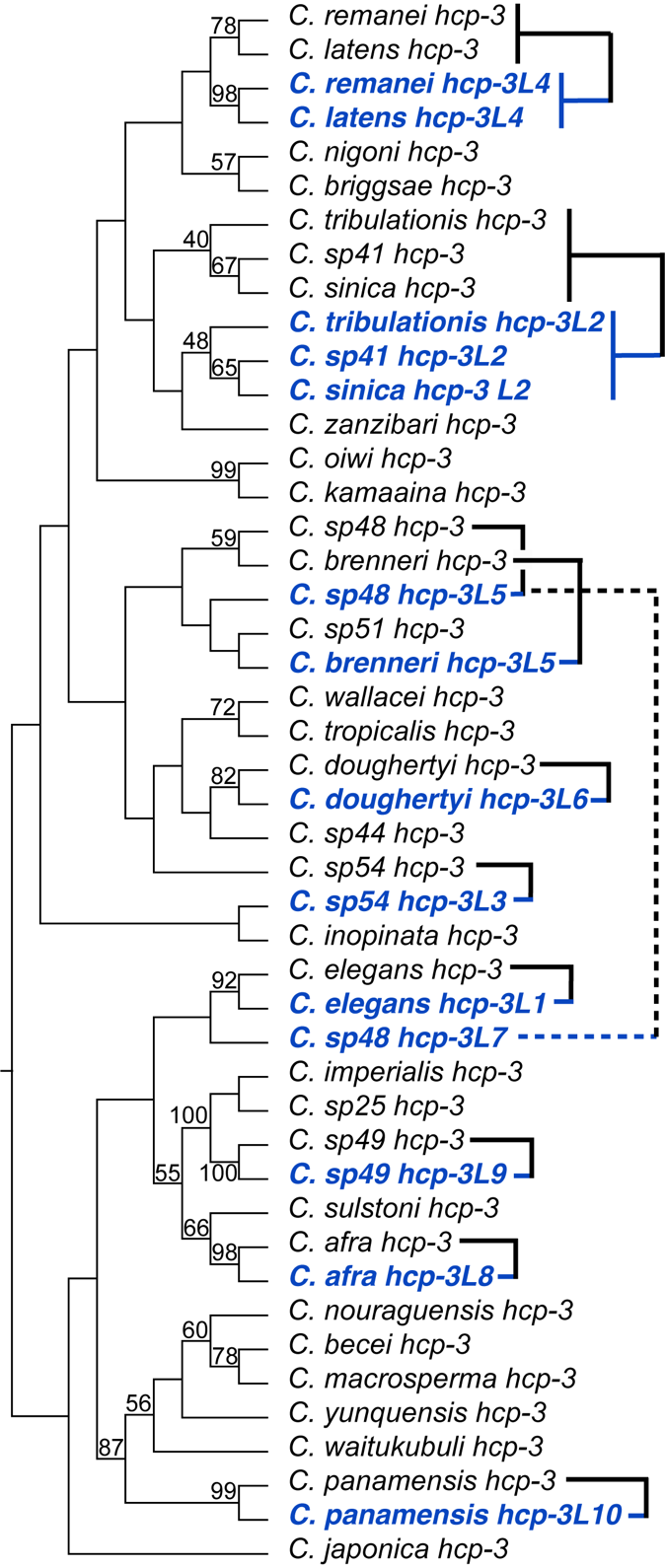

**Supplementary Figure S1. Maximum-likelihood phylogenetic tree showing relationship between *hcp-3* paralogs in *Caenorhabditis* species based on an amino acid alignment.**

A maximum likelihood tree based on an amino acid alignment of the histone fold domain (HFD) of ancestral *hcp-3* (black) and *hcp-3* paralogs (blue) is shown as a cladogram (branch lengths are not scaled to evolutionary divergence). Bootstrap values of 40 and above are indicated. Overall, this phylogeny is much more poorly resolved than one based on the nucleotide alignment (Figure 2) and does not fully recapitulate known relationships between *Caenorhabditis* species or relationships between *hcp-3* and *hcp-3L* genes from the same species. However, the well resolved nodes are in agreement in both phylogenies.

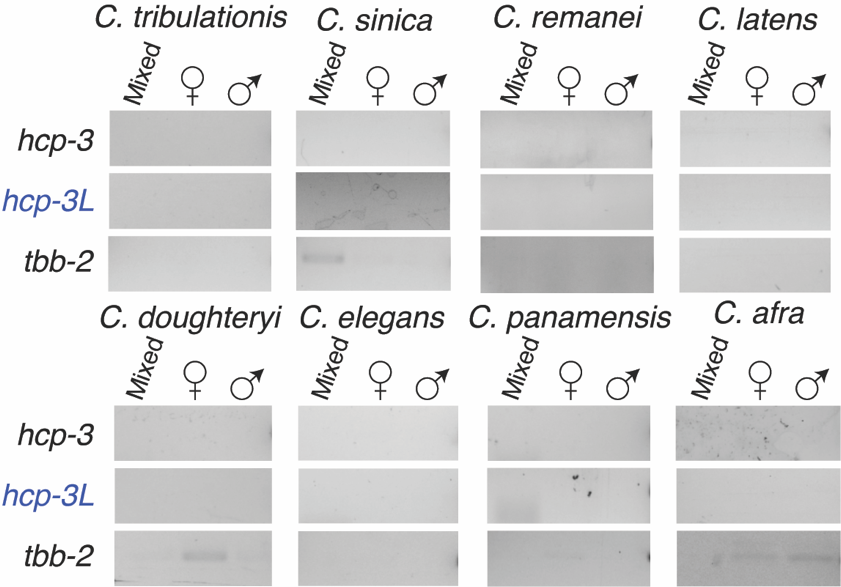

**Supplementary Figure S2. RT-PCR controls for expression analysis of ancestral and duplicate *hcp-3* genes.**

No reverse transcriptase (-RT) PCR control of ancestral *hcp-3* (top), *hcp-3L* (middle), or *tbb-2* (bottom; loading control) in species with *hcp-3* duplicates. RNA from a mixed worm population of various larval stages, L4 or young adult females/hermaphrodites or L4 or young adult males were used. In some cases we see bands in the *tbb-2* -RT samples, but these were significantly fainter than their +RT counterparts.

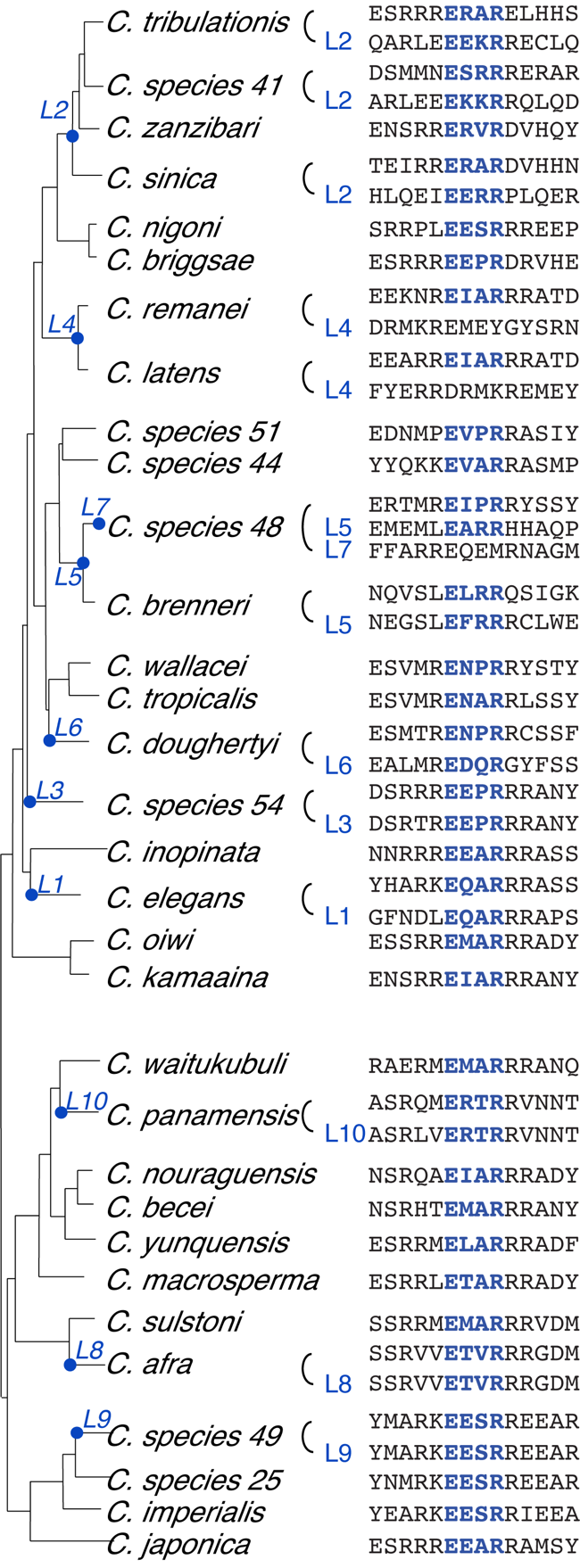

**Supplementary Figure S3.** **Alignment of ExxR motif of CenH3 paralogs.**

Alignments of ExxR residues (blue colored) and surrounding residues in CenH3 paralogs shown beside a *Caenorhabditis* species tree.

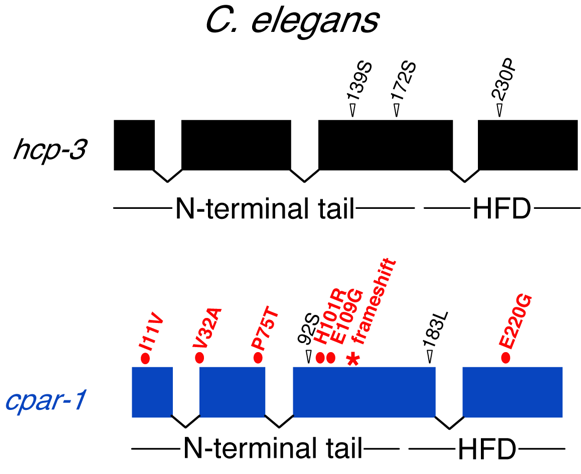

**Supplementary Figure S4.** **Natural variation in *C. elegans hcp-3* and *cpar-1*.**

Schematic of gene structure of *hcp-3* (top, black) and *cpar-1* (bottom, blue) in *C.* elegans where boxes represent exons and lines represent introns. The regions of each gene that code for the protein N-terminal domain and HFD are indicated below. Natural variation in *C. elegans* strains are indicated by arrowheads (black) and ovals (red) that represent synonymous and nonsynonymous mutations respectively. Three synonymous mutations and zero nonsynonymous mutations were found in *hcp-3*, while six nonsynonymous and two synonymous mutations were found in *cpar-1*. In addition, a single nucleotide insertion in *cpar-1* that causes a frameshift resulting in an early stop codon is indicated.

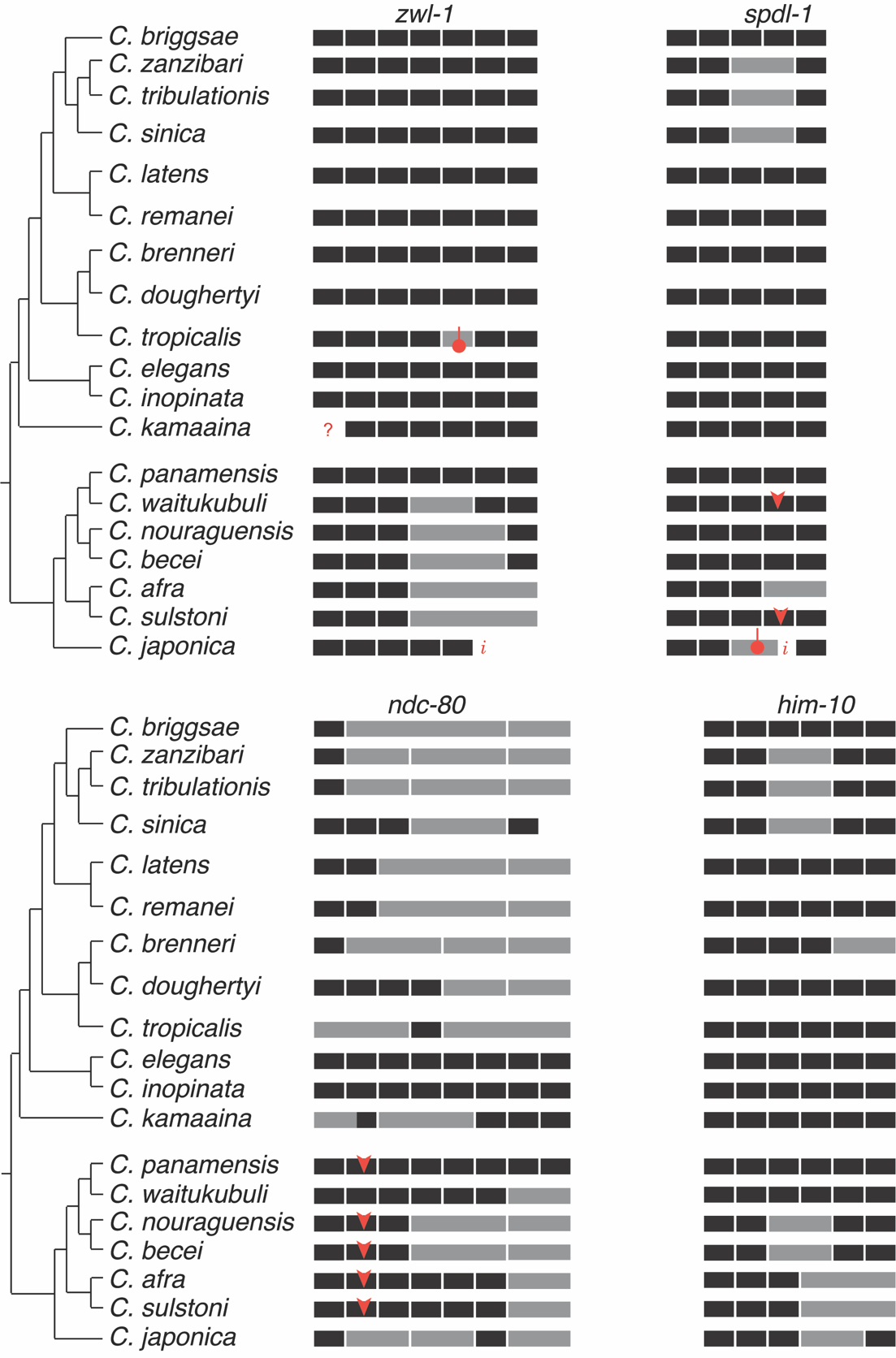

**Supplementary Figure S5.** **Schematic illustrating partial retrotransposition of outer kinetochore genes in representative *Caenorhabditis* species.**

Schematic of the exons of 4 outer kinetochore proteins (*zwl-1*, *spdl-1*, *ndc-80* and *him-10*) in a representative set of *Caenorhabditis* species. Each box represents an exon with black boxes showing ancestral exons and grey boxes showing fusion events between exons likely due to partial retrotransposition and overwriting of the genomic locus. Orange arrows indicate insertion events that likely create new introns and orange dos represent deletions in exons. Incomplete genomic sequence information is indicated with an *i.* While we were unable to identify exon 1 of *C. kamaaina zwl-1* using homology, it is unlikely to have been pseudogenized since *zwl-1* is an essential gene and has not been pseudogenized in related species.
