## Supplementary Data for "Recurrent but short-lived duplications of centromeric proteins in holocentric *Caenorhabditis* species"

**Supplementary Data File S1- Alignment of *Caenorhabditis* cenH3 paralogs used for Figure 2**

>C_tribulationis_hcp3_HFD

GGGCAGAAAGCGTTAAGAGAAATTCGCAAGTATCAAAAGTCTACGGATATGCTTATTCAGAAAGCACCCTTCGCTCGTCTCGTCCACGAAATCATACAGGAAACGACTTTGTTTAGT------------CATGACTTTCGTATTCGTGCCGACGCTCTGATGGCTCTTCAAGAAGCATCTGAAGCGTTTATGGTGGAAATGTTCGAAGGATCCTTCTTGATCTGCAATCACGCCAAACGCGTCACCCTTATGCCGACTGATATTCAGTTGTACCGTCGTTTGTGTCTTCGA

>C_sp41_hcp3_HFD

GGACAAAAAGCACTATCTGAAATCCGCAAGTATCAAAAATCTACGGATATGCTTATCCAGAAGGCGCCGTTCGTCCGCCTCGTCAACGAAATCATCCAGGAAGCGACGTCGTTCAGT------------AAAGAATTTCGTATTCGAGCCGACGCTTTGATGGCTCTACAAGAAGCCTCAGAAGCTTTTATGGTGGAAATGTTCGAAGGATCCGTGTTGATCTGTAATCACGCTAAACGTGTCACCCTTATGCCAACTGATATTCAGTTATACCGTCGCTTGTGTCTCCGA

>C_zanzibari_hcp3_HFD

GGACAGAAAGCGCTCTCCGAAATCCGAAAATATCAGAAGTCTACGGATATGCTCATTCACAAAGCTCCATTTGCCCGTCTTGTTCACGAAATCATACAGGGATCGTCATTGGAGAGT------------AAAGACTTTCGTATTCGTGCGGACGCTTTGATGGCTCTTCAGGAAGCCGCGGAAGCGTTTATGGTGGAAATGTTTGAAGGGTCCGCGTTGATCTGTAATCACGCTAAACGTGTCACCCTTATGCCGACCGATATTCAGCTATACCGTCGTTTGTGTCTACGA

>C_sinica_hcp3_HFD

GGACAGAAAGCTCTTGCTGAAATCCGGAAGTATCAGAAATCTACGGATATGCTCATTCAAAAAGCTCCATTCGCCCGTCTCGTTCACGAGATCATCCAAGAAGCAACTTCGTTTAGC------------AAGGAGTATCGTATTCGTGCTGATGCCTTGATGGCCCTTCAAGAAGCGTCGGAAGCCTTCATGGTGGAAATGTTTGAGGGATCTGTATTGATTTGTAATCACGCGAAACGAGTCACCCTAATGCCCACCGACATTCAGTTATATCGTCGTTTGTGCCTCCGA

>C_nigoni_hcp3_HFD

GGACAGAAAGCCTTAGCTGAAATTCGAAAGTATCAGAAGTCGACAGATATGCTGATCCAGAAGGCTCCTTTTGCTCGTCTTGTTCATGAAATTGTTCGAGAACAAACCAACCAAAGT------------AAAGACTATCGTATTCGTGCCGATGCTTTGATGGCTCTACAGGAAGCAGCAGAAGCATTCATGGTTGAAATGTTCGAAGGATCCGTTCTGATTTGCAATCACGCTAAGCGTGTCACACTCATGCCCACTGACATTCAGCTGTATCGTCGCTTGTGCCTCCGA

>C_briggsae_hcp3_HFD

GGACAGAAAGCCTTGGCTGAAATTCGAAAGTATCAGAAGTCGACAGATATGTTGATCCAGAAGGCTCCTTTTGTTCGTCTTGTTCATGAAATTATTCGAGAACAAACCTACAAAAGT------------CAAGACTATCGTATTCGTGCGGATGCTTTGATGGCTCTACAGGAAGCAGCAGAAGCATTCATGGTTGAAATGTTCGAAGGATCCGTACTGATTTGCAATCACGCTAAGCGTGTCACACTCATGCCCACTGACATTCAGCTGTATCGTCGCTTGTGCCTTCGA

>C_remanei_hcp3_HFD

GGACAGAGAGCGCTTGAGGAAATTCGAAAATACCAAAAGTCCACCGATATGCTGATTCAGAAAGCTCCCTTTGCACGTCTTGTCCACGAAATTATGCGCGAAGCAACTTCGGAAAGT------------CAAGATTTTCGGATTCGTGCAGACGCTTTGATGGCTCTTCAAGAAGCGGCAGAAGCGTTCATGGTGGAGATGTTCGAGGGATCCGTGTTGATTTGTAATCACGCGAAAAGAGTAACTCTCATGCCGACAGATATTCAATTATATCGTCGCTTATGTCTTCGG

>C_latens_hcp3_HFD

GGACAGAGAGCGCTCGAGGAAATTCGAAAATACCAAAAGTCCACCGATATGCTGATTCAGAAAGctccgtttgctcgtCTTGTCCACGAAATTATGCGCGAAGCAACTTCGGAAAGT------------CATGACTTTCGGATTCGAGCAGATGCTTTGATGGCTCTTCAAGAAGCGGCAGAAGCGTTCATGGTGGAGATGTTCGAGGGATCCGTGCTGATTTGTAATCACGCGAAAAGAGTAACTCTCATGCCGACAGATATTCAATTATATCGCCGCTTATGTCTTCGG

>C_sp51_hcp3_HFD

GGACAAAAAGCACTGGCTGAAATCAGACAATATCAGAAATCAACGGATTTTTTGATTCAAAAAGCACCGTTTGCACGGTTGGTCCATGAAATAGTTCGTGAGGCTACTTCAAGCAGT------------GGTGATTTCCGAGTTCGCGCGGATGCTCTCATGGCCCTTCAAGAAGGTGCCGAAGCGTTTATAGTAGAAATGTTTGAAGGATCTGTATTAATCTGCAATCATGCTAAGCGTGTCACCCTCATGCCAACTGATATCCAATTATACCGCCGATTGTGCCTCCGA

>C_sp44_hcp3_HFD

GGACATAAGGCCTTGGCAGAGATTCGACATTATCAGAAGACAACTGATCTTCTCATTCAAAAGGCCCCATTCGCTCGTCTCGTCCATGAAATCATCCGCGAAATGACCCCTAACAAT------------GCCGACTATCGAGTCCGTGCCGACGCTCTTCTCGCCCTCCAAGAAGGCGCAGAAGCTTTTATGGTAGAAATGTTCGAAGGATCCGCTCTGATTTGTAATCACGCAAAGCGTGTCACCCTTATGCCAGCGGATATTCAACTATATCGTCGATTGTGCCTTCGA

>C_sp48_hcp3_HFD

GGACAAAAAGCGTTGGCAGAGATCAGGCAATACCAGAAGTCAACTGATCTTTTGATTCAAAAAGCTCCATTTGCACGACTTGTCCATGAAATTATTCGGGAAGCAACTTCAAACAGT------------GGGGATTATCGCGTTCGCGCAGATGCTCTTCTAGCTCTCCAAGAAGGGGCAGAAGCGTTTATGGTGGAAATGTTTGAAGGATCTGTATTAATTTGTAATCACGCAAAGCGTGTAACTCTTATGCCCACCGATATCCAATTATATCGACGTCTGTGCCTCCGA

>C_brenneri_hcp3_HFD

GGGCAAAAAGCGTTGGCAGAGATAAGGAAATACCAGAAGTCAACTGATCTTTTGATTCAGAAAGCTCCATTTGCACGCCTTGTCCATGAAATTATCCGGGAAGCAACTACAAATAGT------------GGAGATTATCGCGTTCGTGCAGATGCTCTTCTAGCTCTCCAAGAAGGCGCTGAAGCATTTATGGTTGAAATGTTTGAAGGATCTGTATTAATTTGTAACCACGCGAAGCGCGTAACTCTTATGCCCACAGATATTCAATTATATCGACGTCTGTGCCTCAGA

>C_wallacei_hcp3_HFD

GGACAGAAAGCGTTGTCCGAAATCCGACAATATCAAAAATCAACGGATCTTCTCATTCAAAAAGCTCCGTTCGCACGTCTCGTTCATGAAATCATTCGTGAAGAAACCAAT------------------AAAGACTATCGAATCCGCGCTGATGCTATTCATGCCTTGCAGGAAGCGGCTGAAGCATTTATGGTTGAAATGTTCGAAGGATCTACATTAATTTCCAATCACGCAAAACGTGTCACCCTTATGCCAACTGATATCCAATTATATCGTCGCTTGTGTCTTCGA

>C_tropicalis_hcp3_HFD

GGACAAAGAGCTTTGGCCGAAATTCGGCAGTATCAAAAATCAACGGATCTTTTGATTCAAAAAGCCCCCTTCGCACGTCTCGTCCATGAAATCGTTCGTGAAGAAACCAAC------------------CAAGATTATCGAGTCCGCGCTGATGCTATTCTTGCACTACAAGAAGCTGCCGAAGCGTTTATGGTTGAAATGTTCGAAGGATCTACATTAATTTCCAATCACGCAAAGCGTGTTACTTTGATGCCCACTGATATTCAACTATATCGGCGGTTGTGCCTCAGA

>C_doughertyi_hcp3_HFD

GGACAGAAAGCGTTAGCTGAAATTCGACAGTATCAAAAATCAACAGATCTTCTCATTCAGAAAGCCCCGTTTGCACGTCTTGTTCATGAAATTATTCGCGAAGAATCATCACAG---------------ACTGATTTTCGCGTTCGCGCAGATGCTCTTCTCGCCCTTCAGGAGGCTGCCGAAGCATTCATGGTGGAAATGTTCGAAGGGTCTGTGCTTATCTGCAACCACGCGAAACGTGTCACTCTCATGCCAGCAGATATTCAACTATATCGGCGGTTGTGTCTTCGA

>C_sp54_hcp3_HFD

GGACACAAAGCGTTGTCCGAAATTCGAATGTATCAAAAGTCTACTGATCTGCTTATTCAGAAAGCTCCATTCGCTCGTCTTGTCCATGAAATCATTCGCGATACAACCTCCAACAGT------------CAAGATTATCGAGTCCGCGCGGATGCTCTTCTCGCCCTCCAAGAAGCAGCTGAAGCATTCTTAGTCGAAATGTTCGAGGGTTCCGTGCTGATTTGCAATCATGCCAAGCGTGTCACCCTTATGCCAACTGATATTCAACTGTATCGTCGCCTGTGCCTTCGA

>C_inopinata_hcp3_HFD

GGCCAAAAAGCGATGGCTGAGATTCGTAAATATCAAAAATCAACGGATCTACTTATTCAAAAAGCTCCATTTGCTCGTCTCGTGCATGAAATTGTTCGTGAGGCAACTTCTCATAGT------------AATGATTTTCGAGTTCGTGCAGATGCCATTATGGCCCTCCAAGAAGCTGCCGAAGCTTTTATAGTGAACATGTTTGAAGGATCTGCAATGATCTGCCATCATGCGAAACGTGTCACTCTTATGGAAACTGATATTCTGCTGTATCGTCGTTTGTGTCTTCCA

>C_elegans_hcp3_HFD

GGCCAGAAGGCATTGGAAGAGATCCGCAAGTACCAAAAAACTGAAGACCTTCTGATTCAAAAGGCTCCGTTCGCACGCCTCGTCCGCGAAATTATGCAGACTTCCACTCCATTTGGC------------GCCGACTGCCGTATTCGTTCTGACGCCATCAGTGCTCTTCAAGAAGCGGCGGAAGCATTTTTGGTCGAAATGTTCGAAGGATCGTCTCTTATATCCACCCATGCGAAACGTGTCACACTCATGACAACGGATATTCAGTTATACAGACGTCTCTGCCTTCGA

>C_oiwi_hcp3_HFD

GGACAGAAAGCCTTAGCAGAAATCAGAAAATACCAAAGATCGACGGATATGCTCATTCAAAAGGCTCCGTTTGCCCGCCTTGTACATGAAATCATCCGCCAAGAAACGACCCAGGAT------------AGT---TTCCGGATCAGAGCAGACGCTCTATGTGCTCTGCAAGAAGCTGCTGAAGCATTCCTTGTCGAAATGTTCGAAGGATCCGTGTTGATCTGTAATCACGCCAAACGAGTCACTCTTATGCCCACGGATATTCAATTATATCGCCGACTATGTCTCCGG

>C_kamaaina_hcp3_HFD

GGACACAAAGCCTTGGCAGAAATCAGAAAGTACCAAAGATCGACGGATATGCTCATTCAAAAGGCTCCGTTCGCCCGCCTCGTACACGAAATTGTTCGTCAAGAAACAACCAAAGAT------------TGC---TTCCGGATCAGAGCAGACGCTCTATGTGCTCTGCAAGAAGCTGCCGAAGCATTCCTTGTTGAAATGTTCGAAGGATCCGTATTGATCTGTAATCACGCCAAAAGAGTCACTCTCATGCCCACGGATATTCAATTATATCGCCGTCTATGTCTCCGG

>C_waitukubuli_hcp3_HFD

GGAGAGAAAGCAATGCGCGAGATTCGGCAGTATCAGAAGTCCACTGATCTGCTCATCCAAAAAGCTCCATTCTGCCGTCTGGTCCACGAAATCATGCAAGAAGTCACTTCGTTCAGC------------TCAGATTTCCGTATCCGAGCCGAAGCTCTCGGTGCCCTTCAAGAAGCGGCTGAAGCTTTCCTAGTCGAAATGTTCGAAGGCTCCGTTCTGCTAGCCAATCACGCGAAACGTGTCACTCTGATGCCAACCGATATTCAACTATATCGCCGTCTCTGTCTTCGC

>C_panamensis_hcp3_HFD

GGAGAGAAGGCGATGAAGGAGATTCGGAGATACCAGAAGTCAACTGATCTGCTAATCCAAAAAGCTCCATTCGTGAGACTCGTCCACGAAATCATGGCCGATGTGACGCCGCGCAGC------------TCTGAATACCGGATTCGTGCAGAGGCACTCGGCGCTCTCCAAGAAGCAGCCGAAGCATTCCTAGTCGAGATGTTCGAAGGATCAGTGCTCATCGCCAATCACGCCAAACGAGTCACCCTGATGCCCACGGATATCCAGCTGTACCGTCGTCTTTGTCTCCGT

>C_nouraguensis_hcp3_HFD

GGAGAGAAGGCTATGCGTGAGATCAGGCAGTACCAAAAGTCCACTGACATGCTCATCCAAAAGGCACCGTTCTGTCGTCTGGTTCATGAAATCGTGCAAGACGTCACCTCATCCAGT------------TCCGGTTTCCGCATACGCGCCGAGGCTCTCGGTGCTCTGCAAGAAGCCGCTGAAGCATTTCTTGTCGAGATGTTCGAAGGATCAGTGCTCATCGCTAACCATGCCAAGCGAGTAACTCTCATGCCTACTGATATCCAATTGTACCGCCGTCTGTGCCTTCGT

>C_becei_hcp3_HFD

GGAGAGAAGGCGATGCGCGAGATCAGGCAGTACCAAAAGTCCACTGATATGCTCATCCAAAAGGCACCGTTCTGTCGGCTGGTTCATGAAATCATGCAAGACGTCACCTCATCCAGC------------TCCGATTTCCGCATTCGCGCAGAGGCGCTTGGTGCTCTGCAAGAAGCCGCCGAAGCATTCCTTGTCGAGATGTTCGAAGGATCAGTCCTCATCGCCAACCATGCAAAGCGAGTAACTCTCATGCCCACTGATATCCAACTGTACCGCCGTCTGTGCCTTCGT

>C_yunquensis_hcp3_HFD

GGAGAGAAGGCGATGCGGGAGATCAGACAGTACCAGCAGTCCACCGATATGCTCATCAAAAAAGCTCCATTCTGTCGACTGGTTCATGAAATTATGCAAGAAGTCACTGGGTTCAGC------------TCCGATTTCCGGATTCGGGCAGAAGCTCTCGCCGCCCTTCAAGAAGCCGCGGAAGCGTTCCTTGTGGAAATGTTCGAAGGATCTGTCCTGATCGCGAGCCATGCCAAACGTGTCACTCTCATGACTTCTGATATCCGCCTGTACCGCCGTCTCTGCCTCCGT

>C_macrosperma_hcp3_HFD

GGAGAGAAAGCGATGCGAGAGATCAGGCAGTACCAGAAGTCCACCGACATGCTCATTCAAAAAGCTCCATTCTGTCGACTTGTCCACGAGATTATGCAAGATGTCACCTCCTTTAGT------------TCAGACTTCCGTATCCGAGCAGAGGCACTTGGTGCCCTTCAAGAAGCCGCAGAAGCGTTCCTTGTCGAAATGTTCGAGGGTTCCGTTTTGATTGCGAACCACGCCAAGCGAGTCACTTTAATGCCAACAGATATACAACTCTATCGTCGTCTTTGCCTCCGT

>C_sulstoni_hcp3_HFD

---GACCGCGCGCTTCAGGAGATTCGGCAGTACCAGAAGTCGACCGACCTGCTCATTCAAAAAGCTCCATTCTGCCGTCTCGTTCAAGAAATCGTCCGGGAGTCGAGTTCGTCCACT------------TCAGATTTCCGTGTCAGAGCCGATGCTCTCTCGGCCCTTCAGGAAGCCGCCGAAGCGTTCCTCGTCGAGATGTTCGAGGGCTCCCAATTGATTGCCGCGCACGCGAAACGTGTTACTCTGATGCCCTCCGATATCCAACTCTACCGTCGCTTGTGCCTGAGA

>C_afra_hcp3_HFD

---GACCGCGCACTTCTGGAGATCCGACAGTACCAGAAGTCTACCAACCTGCTAATTCAGAAGGCCCCATTCTGTCGTCTCGTTCAAGAAATCCTACGCGAGGTTACAAGTTCAAGCGATTACCGTTCAAGCGATTACCGCATCAGAGCAGATGCTCTCTCCGCCCTTCAAGAGGCCGCGGAAGCCTTCCTCGTTGAGATGTTCGAGGGCTCCCAGCTGATTGCCACTCACGCAAGACGTGTCACTCTTATGCACAGCGATATTCAACTGTACCGCCGTCTCTGCCTTCGG

>C_sp49_hcp3_HFD

GGACAGAAGGCGTTGAGCGAGATCAGAAAGTACCAGAAATCGACGGATATGTTGATTCAAAAGGCCCCGTTCCACCGCGTCGTCCAGGAGATCCTCTGCGAGACGTCCGGCTTCACC------------AACGCCCACCGCATCCGAGCCGACGCGATCTCCGCGCTCCAGGAGGCCGCCGAGGCGTTCCTCGTCGAGATGTTCGAGGGCGCCATGCTGCTTTCGAATCACGCGAAACGGGTCACTCTGATGGCGTCCGACATCCAGCTGTACCGTCGACTGTGCCTGCGG

>C_sp25_hcp3_HFD

GGACAGAAGGCGTTGGCCGAAATCAGAAAGTATCAGAAGACGAGTGATCTTTTGATACAAAAGGCGCCGTTCTACCGTGTCGTTCAAGAGATCCTTCGCGAAACGTCAGGCTTCACC------------AACGACCACCGCATCCGAGCGGACGCGATCGCCGCTCTTCAGGAGGCTGCCGAGGCGTTCCTCGTCGAGATGTTCGAGGGCTCAGCCCTTCTGTCATTGCACGCGAAGCGGGTGACTCTTATGCCGTCGGACATTCAACTGTACCGCAGACTGTGTCTCAGA

>C_imperialis_hcp3_HFD

GGGCAGAAGGCGTTGAGCGAGATTAGAAAGTATCAGAAGTCGACTGATCTTCTTATTCAAAAGGCTCCGTTCTACCGTGTGGTGCAAGAGATCCTCCGCGAGACGTCTGGCTTCACC------------AACGACCACCGGATCCGTGCTGATGCCATCGCCGCACTTCAAGAAGCGGCCGAAGCGTTCATCGTCGAGATGTTCGAGGGAGCGACGCTTCTGTCGACGCACGCGAAGCGAGTCACTCTGATGCCCTCCGACATCCAACTATACCGCAGATTGTGTCTCAGA

>C_japonica_hcp3_HFD

GGACAGAAGGCGTTGAGTGAGATTCGAAAATACCAAAATTCCACTGATTTGCTCATTCAAAAAGCCCCCTTCCGTCGATTAGTTCACCAGATTATTCAAGAAGCGACCGGCTTCGAT------------TCCGGATTCCGCATTCGCGCCGACGCGATGTCTGCCCTACAAGAAGCCGCCGAGGCGTTCATCGTCGAGATGTTCGAGGGATCTGTTCTCATCTCGAATCACGCAAAACGGGTCACTCTGATGACGGCCGACATTCAATTGTACCGTCGACTTTGCCTCCGA

>C_elegans_hcp3L1_cpar-1_HFD

GGCCAGAAGGCATTGGAAGAGATTCGCAAATACCAAGAATCTGAAGACCTTCTGATTCCAAAGGCTCCGTTCGCACGCCTCGTCCGCGAAATTATGCAGACTTCCACTCCATTTTCC------------TCAGACCTTCGCATTCGTTCTGACGCCATCAATGCTCTTCAGGAAGCATCGGAAGCACTGTTGGTCCAAATGTTCGATGGTTCATCACTCATTTCCGCCCATTCCAAACGTGCCACACTGACGACTACAGACGTGCAGCTTTACCGCCGTCTTTGCCTTCCA

>C_tribulationis_hcp3L2_HFD

GGACAAAAAGCGTTGGCTGAGATTCGCCGATACCAAAAATCTACGGATATGCTCATTCAGAAGGCTCCGTTTGCTCGTCTCGTACATGAAATCATCCAAGAATCAACAACCTTGAGT------------CGCGACTTCCGTATTCGTTCTGATGCGTTAATGGCTCTCCAGGAGGGTGCCGAAGCATTCATGGTAGAGATGTTCGAAGGATCGGCTCTCATCTGCAATCACGCTAAGCGCGTTACTCTCATGCCAACTGATGTGCAGCTGTACCGTCGCCTGTGCCTTCGC

>C_sp41_hcp3L2_HFD

GGACAAAAAGCGTTGGCTGAGATTCGCCAATATCAGAAATCTACGGATATGTTGATCCAGAAAGCTCCATTCGCTCGCCTTGTTCATGAAATCATCCAGGACTCCACAAACTTCAGC------------CGAGATTTCCGTATTCGTGCAGATGCCTTGATGGCTCTCCAGGAAGCGGCCGAAGCGTTCATGGTTGAGATGTTCGAAGGATCGACTCTCATCTGTAATCACGCAAAGCGCGTCACTCTCATGCCCACGGATCTCCAGCTGTACCGTCGCCTGTGCCTTCGA

>C_sinica_hcp3L2_HFD

GGGCAAAAAGCATTGGCTGAAATTCGCCAATACCAAAGGACAACAGAAATGTTGATACAAAAAGCTCCCTTCGCTCGTCTCGTCCATGAAATCATCCAGGACGCCACTTCCTTCAGC------------CGTGATTTCCGTATCCGTGCTGATGCTCTAATGGCTCTCCAGGAGGCAGCCGAAGCGTTTATGGTGGAAATGTTTGAGGGATCTGTGCTCATCTGTAACCACGCAAAACGTGTTACGCTCATGCCGACCGACCTTCAGCTCTACCGTCGCCTCTGCCTTCGC

>C_sp54_hcp3L3_HFD

GGACAAAAAGCGTTGTACGAGATTCAAAAGTATCAAAAGTCTACTGATCTGCTTATTCCGAAAGCTCCATTCGCTCGTATTGTTCATGAAATGATTCACAAAGCAACCTCAACCAGT------------CACGATTTACGAGTCCGCGCCAATACTTTTCTCCCCCTCCAAGAAGCAGCGGAAGCATTCCTAGTCCAAATGTTCCACGGTTCCATGAAGTATTGCAATTCTGCCAAGCGTGTCACCCTTATGCAAACTGATATTCAAAATTATCGTAGCCCGTGA------

>C_remanei_hcp3L4_HFD

GGACAGAAGGCTCTTCTCGAAATCCGAAAATACCAAAAGTCCACCGACATGCTCATCCAAAAAGCTCCATTCGCTCGTCTCGTCCAAGAAATCCTTCGAGAAACCACAAATGAGAGT------------CACGACTACCGTATCCGTGCAGACGCTCTCATGGCATTACAAGAGGGTGCCGAGGCGTTTATGGTGGAAATGTTCGAGGGATCCGTGTTGATTTCGAACCATGCGAAACGGGTCACGTTGATGCCCACTGATGTTCAATTGTATAGACGTTTGTGTCTCAGA

>C_latens_hcp3L4_HFD

GGACAAAAGGCTCTTCTCGAAATTCGAAAATACCAAAAGTCCACTGACATGCTCATCCAAAAAGCCCCATTCGCTCGTCTCGTCCAGGAAATCCTTCGGGAAACCACAAATGAGAGT------------CACGACTACCGTATCCGTGCAGACGCTCTCATGGCATTACAAGAGGGTGCCGAGGCGTTTATGGTCGAGATGTTCGAGGGATCTGTATTGATTTCGAATCATGCGAAACGGGTCACCTTGATGCCCACTGATATTCAGCTGTATAGACGTTTGTGTCTCAGA

>C_sp48_hcp3L5_HFD

GGTGAGAAAGCCTTGGCCGAAATCAGACAGTACCAAAGGTCAACGGATCTTCTGATTCAGAAAGCTCCATTCGCCCGTCTTGTCCATGAAATCGTGAGCGAAGCAACTTCTTCGAGT------------GGAGACTACAGGATTCGCGCCGATGCTCTCATGGCTCTGCAAGAAGGCGCCGAAGCATTCATGGTGGAAATGTTTGAGGGATCGGCGCTCATCTGCAACCACGCCAAACGAGTGACACTAATGGCATCGGACGTTCAGCTTTACAGGCGCCTTTGCCTGAGA

>C_brenneri_hcp3L5_HFD

GGACAGAAGGCCCTTGCGGAAATCCGCAAGTACCAAAAGTCCACGGATCTTCTGATTCAGAGAGCACCATTCGCACGTCTTGTACATGAAATTGTCCGTGAAGCAACTGCTTCAAGT------------GGCGACTACCGGGTTCGCGCAGATGCTCTCATGGCTCTGCAAGAAGGAGCTGAGGCCTTTATGGTGGAAATGTTTGAAGGATCGGTGCTCATTTGCAACCATGCCAAGCGTGTCACTCTCATGCCAACGGACATTCAGCTCTACCGCCGACTCTGCCTTCGT

>C_doughertyi_hcp3L6_HFD

GGACAAAAGGCATTGGCTGAAATCCGACAGTATCAAAAATCAACAGATCTTCTCATTCAGAAGGCTCCATTCGCACGTCTCGTTCATGAAATCATTCGCGAAGAAACTGCAGTC---------------GATGATTTCCGTGTCCGCGCAGATGCTCTTCTCGCCCTTCAGGAAGCTGCTGAAGCATTCATGGTGGAAATGTTCGAAGGATCTGTGCTTATCTGCAATCACGCAAAACGTGTCACTCTCATGCCAGCTGATGTTCAATTGTACCGACGTTTGTGCCTCCGT

>C_sp48_hcp3L7_HFD

GGCCAAAGAGCAATTGCGGAGATCAAACACTACCAGAAGACCACCGAACTTTTGATCCAAAAAGCGCCATTTGCTCGACTCGTCCAAGAAGTGGTTCAAGAAGCCACCTCAGAAAGT------------AGCAGTTACAGCATCCGCACTGATGCACTATCAGCGCTCCAGGAGGGTGCCGAAGCGTTTTTGGTGGAGATGTTCGAAGGCTCAAGCATGATTGCTAATCACGCCAAAAGAGCAACGTTGGGGTCGACCGATCTCAAATTGTATCGTCGCCTTTGCCTTAGG

>C_afra_hcp3L8_HFD

---GATCGCGCACTTCTGGAGATCCGACAGTATCAGAAGTCTACCGACCTGCTAATTCAGAAGGCCCCGTTCTGTCGTCTCGTTCAAGAGATCCTACGCGAGGAGGCTACTACAAGT---------TCAAGCGACTACCGCATCAGAGCGGATGCTCTCACCGCCCTTCAAGAGGCTGCTGAGGCCTTCCTCGTTGAGATGTTCGAGGGCTCCCAGCTGATTGCCACTCACGCAAGACGTGTCACTCTTATGCACAGTGATATTCAACTCTACCGCCGTCTTTGCCTTCGG

>C_sp49_hcp3L9_HFD

GGACAGAAGGCGTTGAGCGAGATCAGAAAGTACCAGAAGTCGACGGATATGTTGATTCAAAAGGCCCCGTTCCACCGCGTCGTCCAGGAGATCCTCTGCGAGACCTCCGGCTTCACC------------AACGCCCATCGCATCCGAGCCGACGCGATCTCCGCGCTCCAGGAGGCCGCCGAGGCGTTTCTCGTCGAGATGTTCGAGGGCGCGATGTTGCTGTCGAACCATGCGAAACGGGTCACTCTGATGGCGTCCGACATCCAGCTGTACCGACGACTGTGCCTGCGG

>C_panamensis_hcp3L10_HFD

GGAAAGAAGGCGATGAATGAGATTCGGAGATACCAGAAGTCAAATGATCTGCTAATCCAAAAAGCTCCATTCGTGAGACTCGTCCACGAAATCATGGCCGATGTGACGCCGCGCAGC------------TCTGAATACCGGATTCGTGCAGAGGCACTTGGCGCTCTCCAAGAAGCAGCCGAAGCATTCTTAGTGGAGATGTTCGAAGGATCGATGCTCATCGCCAATCACGCTAAACGAGTCACCCTGACGCCCACGGATATCCAGCTGTACCGTCGTCTTTGCCTCCGT

**Supplementary Data S2: All HCP-3 sequences used in this study**

>C_tribulationis_HCP3

MYAHTGPIIEEVEEAATDGQTVHRDWSKDPDVLRIRRELQKLMILPGFNNNADLMQRAITILVEQVDEWKLDQDIDGWNICRQEKIESFLPRIANFKNKRAQAIDQFYKERDSMINESRRRERARELHHSNDFGISGRELDHSSRLHQLSRRDSCASRVERYHSSDEDEENHPVPRYRSRSPGPSSSYNQSTMRQRRDDVPQPVRMRSGKSRVTKTRNAKWRPGQKALREIRKYQKSTDMLIQKAPFARLVHEIIQETTLFSHDFRIRADALMALQEASEAFMVEMFEGSFLICNHAKRVTLMPTDIQLYRRLCLRNLS

>C_tribulationis_HCP3L2

MEQMYENQHTPIIEELFDCSSVVEREVERVKHEIQALTSQSDFNKNYASMKEVINILTRQIAAWDADEDMGGSHPIRLRSIEKFHAKRVLFTEKLEAAERAYYEKKQARLEEEKRRECLQDRGKIAGQNNQLCHRQGHRYERDDSSDDSSDEENQRQRSRACSPQRRNHPSTSSQQYRVNRHVDYHKKHQNVSQKQQRLRAGINAVTKTKVRKFRPGQKALAEIRRYQKSTDMLIQKAPFARLVHEIIQESTTLSRDFRIRSDALMALQEGAEAFMVEMFEGSALICNHAKRVTLMPTDVQLYRRLCLRNM

>C_sp41_HCP3

MYAHTGPIIEEVEDGPAEGHTVHRDWRQDTDVQRLGREIQKFISLPDFSKNADLMQRAIHVLEKQVDEWKLNQELDGWDHHRQQKIELFQPKIAKFKEKREEAINHYYDVKDSMMNESRRRERAREVLHNTDQNITGFGNSTRLYPNSRRQSFAPRKERYQSSDEDEENDPVPRRRSRSPGPSSSYSHSAMYQLRDDSNAPQQQRMRSGKSRVTKTKNRKWRPGQKALSEIRKYQKSTDMLIQKAPFVRLVNEIIQEATSFSKEFRIRADALMALQEASEAFMVEMFEGSVLICNHAKRVTLMPTDIQLYRRLCLRNMS

>C_sp41_HCP3L2

MDQLYENHHTPVIEEILDTEFVVEREVERVKHDIQLITSQPDFNKNYDSMKEVIDILARQILDWEADEEMSGSHPMRRRIIDKFQAKKVLFTEKLEAAERAYYERKRARLEEEKKRRQLQDCGNIGGEQNRHARFQFMGHRNGRDDSSDDSSDGENRQVQRRRSRSRSPQHRNHSSQQHRRDDRHADGHRNYQSTSKNQPRLRAGINGVTKTKVRKYRPGQKALAEIRQYQKSTDMLIQKAPFARLVHEIIQDSTNFSRDFRIRADALMALQEAAEAFMVEMFEGSTLICNHAKRVTLMPTDLQLYRRLCLRNL

>C_zanzibari_HCP3

MYGHTGPVIQEVLENPNEGGQVVQRADQQNWRHDHDVLRLGREAQKLMSNPDFNTDPDLMEEAISLFENKVAEWRADQGFEGWDHRRQEKIEFFQQRVVKFKEQRKRACDRYYDERDSKIENSRRERVRDVHQYTDNNISNFGNSTRLNQHSHRQSYTTREDRYHTSEDEENIPVQHHRTRAPSSYSNYTQSTMQQRRDDSNVHYRSHQSAAGPSSSQQVRMRSGKSRVTKTRSRKWRPGQKALSEIRKYQKSTDMLIHKAPFARLVHEIIQGSSLESKDFRIRADALMALQEAAEAFMVEMFEGSALICNHAKRVTLMPTDIQLYRRLCLRNL

>C_sinica_HCP3

MYGHTGPLIQEIDETPTEGSTAVRDWTTDEDVRRLGKEVQKLTCQEGFTKNANLMRRLIELLEKQVDEWKEDQDIHGHDHCRQQKIQCFEKRIADYKKKCERSIRRYCDQRDSRTEIRRERARDVHHNSNYDITDRGDSIRLNQHYHRQSLAPREESYHSSDDDEENIPPRLYRSGRSTMYQQRQDDSNVHYRSHHSTLGASSSQQVRMRSGKSRVTKTHNRKWRPGQKALAEIRKYQKSTDMLIQKAPFARLVHEIIQEATSFSKEYRIRADALMALQEASEAFMVEMFEGSVLICNHAKRVTLMPTDIQLYRRLCLRNLS

>C._sinica_HCP3L2

MDHVPPPLTHQPPSLIAAFLPRILVFEVKMQAMQCTPIIEEIHEAPTALEMEIEVVKHNIKKLTSQPDFSKSYSLMEGAITILKKQIERWEYAERRDGPDGARQEHLAKFRLKMEQFEEKLKAAERAYYEKKKAHLQEIEERRPLQERQNAINRQSFEQRRRERRNSSDESSDDETEQFHHRSRSRSPRRNQLTTSNLQRHRQPTQPRLRQGVDRISKTKARKWRPGQKALAEIRQYQRTTEMLIQKAPFARLVHEIIQDATSFSRDFRIRADALMALQEAAEAFMVEMFEGSVLICNHAKRVTLMPTDLQLYRRLCLRNL

>C_nigoni_HCP3

MQRMYHHDSGPHIEEVFDPPPSQETMLREIASHPDVIALSKKVRKITKMPDSAFISSADRLVEIIDAFSEQIEKWKEDETLDDPCPYLSLKIEFFTEKRNQYQRKNSSAVDRYYDGKDSRDYSSRRPLEESRRREEPRDRGHETNIDITHRGDSTSLNHYSQRHYSQRQSQSSRFERDRESEEKDENRHPRQQYRSRSPQHTHSYNQSTMHQRDDTNVYHRSHQSTSQPSQVRMRSGKSRVTKTHNRKFRPGQKALAEIRKYQKSTDMLIQKAPFARLVHEIVREQTNQSKDYRIRADALMALQEAAEAFMVEMFEGSVLICNHAKRVTLMPTDIQLYRRLCLRNLS

>C_briggsae_HCP3

MYHHDSGPHIEEVFDPPSRRTMMQEIETHPDVIAFGKKLRKIKNQPESTFLSSADRMEEIIDAFRDQIAKWEEEEELNEPCEYRQLKIEIFTQKKIEYQRKNNLAVDEFYKKRNLKNHSNRKPLEESRRREEPRDRVHESNIDITHRGDSTSLNHYSRHHYSQRQSQSSRFERERESDEEEENSQPIQRYRSRSPKPSYSYNQSTMQQSQRDDTNVYHRSHQSTSQPPQVRMRSGKSRVTKTHNRKFRPGQKALAEIRKYQKSTDMLIQKAPFVRLVHEIIREQTYKSQDYRIRADALMALQEAAEAFMVEMFEGSVLICNHAKRVTLMPTDIQLYRRLCLRNLS

>C_remanei_HCP3

MYQMHHNGPRIEEMVDPPSRSTTNQLKNDTEYIKSEYRRISHLPDFNRDPELIQEVMNLTKRYIEKWLREERDEPNMERQGWIERFKTKLREWETKKETAEDEYYTRRDASSNEEKNREIARRRATDSQMNITGLHDSTRLNQQSYSRSYENRNRRYSSDEDDDENMAPQRRQRSRSPPSFAHHQRRDDTGSYYRSHHTQNSSNQRTHNTDFSSHYRGQYGPSTSQNVGMPSNAQNVRMRSGKSRVTKTRSRKWRPGQRALEEIRKYQKSTDMLIQKAPFARLVHEIMREATSESQDFRIRADALMALQEAAEAFMVEMFEGSVLICNHAKRVTLMPTDIQLYRRLCLRNLS

>C_remanei_HCP3L4

MAPLLIAVSSFVVCSAILIYLCSKKPKTIDLESEIGITSRKNFNRDSDSMQEVIDIMTRQINKWEQLEDDYGPDATRQRNIEAFQRKRDLWEEKKEQAERAFYERRDRMKREMEYGYSRNQIRRERRMDDDSDSDVMEDDRRQPLGNLDYSRRDNYVRLREAKPMALLNRDRSRSRSPLLLPRQDHPSTSLQLRRPNIPSTPPVRVRPGKSRVTKSKNRKWRPGQKALLEIRKYQKSTDMLIQKAPFARLVQEILRETTNESHDYRIRADALMALQEGAEAFMVEMFEGSVLISNHAKRVTLMPTDVQLYRRLCLRNLS

>C_latens_HCP3

MYQVYHNGPRIEEMVDPPPKSTTNQLKIDTEYIKSEYGRISSHSDFNRNPDAIQEVIDLARRYIEKWQREERDEPNMDRRGWIERFKTKLREWETKKETAVDDYVTRRNASSNEEARREIARRRATDSQLNITGLQDSTRLNQQSYSRSYENRNRRYSSDEDDDENMAPQRRQRSRSPSSFAYHQNTLNHQRRDDTGSYYRSHHTQNSSHQRTHNTDISSHYRRQNGPSTSQNVVMPSNTQNVRMRSGKSRVTKTRNRKWRPGQRALEEIRKYQKSTDMLIQKApfarLVHEIMREATSESHDFRIRADALMALQEAAEAFMVEMFEGSVLICNHAKRVTLMPTDIQLYRRLCLRNLS

>C_latens_HCP3L4

MDQIKEEIEAITSRKNFNRDSDAMQEVIDIMTRQINKWEQLEDDYGPDATRQRNIEAFQRKRDSWEEKKEQAERAFYERRDRMKREMEYDNGYSRSQVRRERRMDDDSDSDVMEDDGRQPLGNLDYSKRNNFVRGREAMPMKVVNRDRSRSRSPLPRPHHPSTSLQLRRPNIPSSPPVRVRPGKSRVTKSKNRKWRPGQKALLEIRKYQKSTDMLIQKAPFARLVQEILRETTNESHDYRIRADALMALQEGAEAFMVEMFEGSVLISNHAKRVTLMPTDIQLYRRLCLRNWS

>C_sp51_HCP3

MLQMDMDGPTIEELVDPQLDDTKQVIFKREVSDIEQECQKICSASNLPTNQQNRKILAVLQKAIDKWREDEEIEGEYEPRIKAIEKFEKKRRLWLRNIENAGANSRERRERRYEDNMPEVPRRASIYQETDITRRTNTTGLQQQSYQGSSSFRRNEYSSDEDTENIPYFRGSRYSSPPRKYNQSSMYQQRRATSPIHQSQHTMNGTSNTQVRMRAGKSRVTKTAVRKHRPGQKALAEIRQYQKSTDFLIQKAPFARLVHEIVREATSSSGDFRVRADALMALQEGAEAFIVEMFEGSVLICNHAKRVTLMPTDIQLYRRLCLRNL

>C_sp44_HCP3

MMLSDMDGPTIEELEDTQEDTRETIFNADIKKYENKMKAIAAVPGFLKDLDELRKIIDICDDAIEKWEAEEYRQGSYSRRREAIQQWGLKKRSFIDKLDRAEYVYYQKKEVARRASMPRDVDITRRNDSTGLYHHSQQGTSYRHRTQEYSSDEEIENRPRSDRDRYRHLRAFSQNKIFQYICFCRSPPRQYNQSSMYQQQRRDQSSMLQRSHQSTMNGSSTAQQVRMRSGKSRVTKTTTRKYRPGHKALAEIRHYQKTTDLLIQKAPFARLVHEIIREMTPNNADYRVRADALLALQEGAEAFMVEMFEGSALICNHAKRVTLMPADIQLYRRLCLRNL

>C_sp48_HCP3

MFRVTDGPTIEEVVETQLTEDTAEAEVRRDYDTILEELRAVLGVPGANRDQERLGRGLCILEKGIDKFEEDEDNQPLEIRRQYLGKLREKYSSCEAKLREAENAFHERKEREYEERTMREIPRRYSSYRDTDITRRNNTTGLYHHSQQSSSNYRQQGYSSDEEMENFPSSHRDRYRSPPRKFSHSTMLQQRRDISPVVNRSHQQSSASSQQVRMRSGKSRVTKTTRKYRPGQKALAEIRQYQKSTDLLIQKAPFARLVHEIIREATSNSGDYRVRADALLALQEGAEAFMVEMFEGSVLICNHAKRVTLMPTDIQLYRRLCLRNL

>C_sp48_HCP3L5

MMDEDSPRIEEIVEEDEAEKDVKKEFDEYRREIEAVTSLPGFNRDSGKMTQVLRIMEKAIGKWEEDEENLGSIQCRRRCLLEFKRRHDNYEKIIEKAEDDFYKRREMEMLEARRHHAQPESRQYPDGGAEQRQTIPKMRAGKSSVTKKPKKFRPGEKALAEIRQYQRSTDLLIQKAPFARLVHEIVSEATSSSGDYRIRADALMALQEGAEAFMVEMFEGSALICNHAKRVTLMASDVQLYRRLCLRNL

>C_sp48_HCP3L7

MLEDHDSPHIEELVDADDEKNAEEAVKKEFFEFKNEIEKISCLPNFTKDSEKLRQVLAVIGKAIDKWEEDEDNEGSIEVRRKYLKELKERYFKYDRIIEKAEQKFFARREQEMRNAGMKTTRYYPGHGSSNHQYPFSREEDVVNSRSPSRHRPNSGHPHQHAPKLSSCRQQHGYNYWDEKLCSRSPRRSNLPIRDQFHQSSTNAQRELKAGKSRVTKKVSHKYCSGQRAIAEIKHYQKTTELLIQKAPFARLVQEVVQEATSESSSYSIRTDALSALQEGAEAFLVEMFEGSSMIANHAKRATLGSTDLKLYRRLCLRNL

>C_brenneri_HCP3

MFHLSDGPTIEELVDTQQLENTAEAEFKEELDVIKKELAAVLAIPDIHRNREALEKSIRILEKAIDKWEEDEENQVSLELRRQSIGKFKEQRRSCKQKLRDAENAFHERREREYEERTMREIPRRYSSFRDTDITRRNNTTGLYHHSQQSSSNFRMQEYSSDEEIENIPSSHRDRYRLEKCLIIVFQNLYFSYPPKKISHSTMLQQRRDISPVVYRSQQQSSAGSQQERMRSGKSRVTKTTRKHRPGQKALAEIRKYQKSTDLLIQKAPFARLVHEIIREATTNSGDYRVRADALLALQEGAEAFMVEMFEGSVLICNHAKRVTLMPTDIQLYRRLCLRNL

>C_brenneri_HCP3L5

MMMMEDSPHIEEIVEVEEAEIEVKRQFEEYKKEIEEVARLPGFNKDSGKMNQVIRIMDKAIEKWEEDEENEGSLEFRRRCLWEFKHRHNNYKKIIEKAEEDFYKRREEAMKMVSILNRTPQSHSYSQTRDGTAGSYRTQATSSGPRHRAFESDEERDQRERMQVGRSTISKKTPKRKYRPGQKALAEIRKYQKSTDLLIQRAPFARLVHEIVREATASSGDYRVRADALMALQEGAEAFMVEMFEGSVLICNHAKRVTLMPTDIQLYRRLCLRNL

>C_wallacei_HCP3

MHSNTDGPTIEEIVDNHLSDEQIAQEEAFKREFQLVREEMTQMTRGGANNDSQQLEKVVNSLTRWIEKWEDDEDYGTKIKLRQDSIIKFRVRKDEYQRKIDRAEEEFFKRRQREQEESVMRENPRRYSTYREDEITRRSDRTGLNHNSYEGSSTRHRTQAYDSDDDYENRPISSYDRYRPSSKKSNQSTMYQQRRDESNSLHRSNFTMNGTSNSQQERMRSGKSRVTKTSRKYRPGQKALSEIRQYQKSTDLLIQKAPFARLVHEIIREETNKDYRIRADAIHALQEAAEAFMVEMFEGSTLISNHAKRVTLMPTDIQLYRRLCLRNL

>C_tropicalis_HCP3

MHSDMDGPTIEEIVDTSFTDEQKSEEDAFRREIDRVKREMYNITSRPGWNNNSDELGKIINIMTRYIDKWEEDEEYDSKINLRQDSIKKFREKRDDYQKKQERAEEEYFERRQRQHEESVMRENARRLSSYRDDDITRRSNRTGLNQNSYEGSSTHHRTHGYDFDEDSENRPISHHDRYRASSKKSNQSTMYHQRRDESSSLHRSHYTMNGTSHSQQQQQVRMRSGKSRVTKTNARKYRPGQRALAEIRQYQKSTDLLIQKAPFARLVHEIVREETNQDYRVRADAILALQEAAEAFMVEMFEGSTLISNHAKRVTLMPTDIQLYRRLCLRNL

>C_doughertyi_HCP3

MYDDSDRPTIEEIDDTENSGYRTAEAIFQERAEEVSKQIKTLLKNPTTDDLKQVIRIMSRSIDEWAEEEDNEGSIRARTTAIKTLTRKRNDYEMSLARKENEFLRKRGQRHEEESMTRENPRRCSSFRDTDITRRTDRTGLNQSSYQGPSSNQQTHYYSSDEDYENRPRSNRDPYRYNDSPQRSHQSSMYQQRRDASPPLHRSHTTLNGTSGSQQVRMRSGKSRVTKTTARKYRPGQKALAEIRQYQKSTDLLIQKAPFARLVHEIIREESSQTDFRVRADALLALQEAAEAFMVEMFEGSVLICNHAKRVTLMPADIQLYRRLCLRNLY

>C_doughertyi_HCP3L6

MMQMYDDGPTIEEILDNHHSGGPITAETIFKEDFDDVSKQIKTLTRDGSTKNADNLKQIISIMSRSIDKWAEDEDMEGSIRMRKDAIEAFTKKRNEFREKITHAEDEYMRRKRQRMDEEALMREDQRGYFSSRDNDMARHAGGTALDFNYQQGTSSNYRARYYSSDEDYENAPRSDRDRYRYPDSPQKSNQLATYHQRRDVSYSPSRSHQPMNGASSSTQVRMRAGKSRVTKKNSRKFRPGQKALAEIRQYQKSTDLLIQKAPFARLVHEIIREETAVDDFRVRADALLALQEAAEAFMVEMFEGSVLICNHAKRVTLMPADVQLYRRLCLRNL

>C_sp54_HCP3

MEQIYDDMRGRIEEIVDPPSRNTTVFRDLTQDQDSDVDRIRNQMKLIMQKPNFNTSIHEMQKVISILDDQINKWESEEELRDNLFEKMERAKADYYKRKEAEDDSRTREEPRRRANYTDMDITDRDNATRLNHLSYQRSYSHRNQIDNSDDDDIENMTRSRRDRSRSPPSHSYYQSTMNQQRRDESYAHQRSHYSMSVPSSSHQVRMRSGKSRVTKTNSRKWRPGHKALSEIRMYQKSTDLLIQKAPFARLVHEIIRDTTSNSQDYRVRADALLALQEAAEAFLVEMFEGSVLICNHAKRVTLMPTDIQLYRRLCLRNL

>C_sp54_HCP3L3

MRQIYDDMRERIEEIVDPPSRNSTVFRDLTQDQDSDLERVRNQIKRIIEKPGSNTSIHELQKVINILDDQIYKWESEEETRDNYLEKMERAKAEYYKRKEAQDDSRRREEPRRRANYTDMDITDRYNATRLNHLSSYQRSYSHQNQIDNSDDDDIENMIPRSRRDRSWSPPSHYKRKVAQDDSRTREEPRRRANYTDMDITDRGNSTRLNHLSSYQRSYSHRNQIDNSDDDDIENMTRKYNTIFKSPPSHGYHQSTMNQQRRDESYAHERSHYSKSAPSNSHQVRMSSYQRSYSHRNQSDNSDDDDIENMIPRSRCDRSRSPPSHGFHQSTMNQQRRDKSYAHQRSHYSMRAPSGSHQERMRSSKSRRVTKKNYRKWRPGQKALYEIQKYQKSTDLLIPKAPFARIVHEMIHKATSTSHDLRVRANTFLPLQEAAEAFLVQMFHGSMKYCNSAKRVTLMQTDIQNYRSP

>C_inopinata_HCP3

MHQMWNGMDGPQIEEIVDHPSRSSSVFRDVSQHMNRFFDNLKADLQRITSKPGFNKSFDEMQKVIDLFDKNIDKLEHYEDMHGSSEELRRHIRLLREKRHNYDEKKEKARMEFFERRADENNRRREEARRRASSPDMDITDRNNGTRLNLHSYRQSYNQQYNINSSDEENYDVARNWRDRSPSRRTQSYNQSTLNVQHRDHSLQRSHHNMNDPSTSQPVRMRFGKNRVTKSLVRKWRPGQKAMAEIRKYQKSTDLLIQKAPFARLVHEIVREATSHSNDFRVRADAIMALQEAAEAFIVNMFEGSAMICHHAKRVTLMETDILLYRRLCLPHY

>C_elegans_HCP3

MADDTPIIEEIAEQNESVTRIMQRLKHDMQRVTSVPGFNTSAAGVNDLIDILNQYKKELEDDAANDYTEAHIHKIRLVTGKRNQYVLKLKQAEDEYHARKEQARRRASSMDFTVGRNSTNLVDYSHGRHHMPSYRRHDSSDEENYSMDGTNGDGNRAGPSNPDRGNRTGPSSSDRVRMRAGRNRVTKTRRYRPGQKALEEIRKYQKTEDLLIQKAPFARLVREIMQTSTPFGADCRIRSDAISALQEAAEAFLVEMFEGSSLISTHAKRVTLMTTDIQLYRRLCLRHL

>C_elegans_HCP3L1

MADDGPIIEEIAEKNGRVARIMQRLQHDTQRVTSVPGFNTSATGYADLIALLDQYKNDLEAVGFNDLEQARRRAPSVDITVGSNSTNLVDYSHGRHDMPSHRRHDSSDEEITAANSHHQSPINVGNRNDTDGTNGRNGSRAGSSSSDRVRMIAGRNRISKTRRYRPGQKALEEIRKYQESEDLLIPKAPFARLVREIMQTSTPFSSDLRIRSDAINALQEASEALLVQMFDGSSLISAHSKRATLTTTDVQLYRRLCLPNL

>C_oiwi_HCP3

MQQQDMDGPYIEEIIEPPSREVSVLRNPGRAEFERSMKQFELAVNGWRNDPELSNTSQKMWGLVDIFNKQIDYCKFCCRQYGSSEDVEEAVKALTTKRDKMVKDINKSENRYFADKDERESSRREMARRRADYSELVVDRSNSTRLQQTSYRRPESHATRTHYSSDEENYHRRDRSRSPTYSRSYDQSRTLQNRQDDNTTHSRRHHDTTGPSGSQQVRMRSGKSRVTKTTSRKYRPGQKALAEIRKYQRSTDMLIQKAPFARLVHEIIRQETTQDSFRIRADALCALQEAAEAFLVEMFEGSVLICNHAKRVTLMPTDIQLYRRLCLRNL

>C_kamaaina_HCP3

MQQQDMDGPYIEEVTESPSREVSIFRDAGRAEFASNMKSFELEVTRRKKDPEFVHNSDKMWDLVGILDRQIEYCTRYSRENEYPDDVEEMTSAIRRKREKMAEEIKKLENRYFAEKDERENSRREIARRRANYSEVTVDRSNNTRLQQSSYRRPESHATRDHYSSDEENYHRRDKSRSPTYSRSYDQSRTLQNRQADNSTHYRRHHDTNVPSGSQQVRMRSGKSRVTKTSSRKWRPGHKALAEIRKYQRSTDMLIQKAPFARLVHEIVRQETTKDCFRIRADALCALQEAAEAFLVEMFEGSVLICNHAKRVTLMPTDIQLYRRLCLRNL

>C_waitukubuli_HCP3

MQQMEEEEMDGPRIVELFDSPRRDTSVLRENNQYENSQGPQLSRYEVRIRSLMALPGFNRDADLMGQAIELLKMHVAEMENEKRLNGRTPELDAKIEKTRNMAERCANKRNDALQRFHEEREESRRAERMEMARRRANQSDLDITERENRTRLQPSSFHRSYEQRTVNYDSDERENEDRRAQSNSRRQRSRSRSPAYSRPPTMNQTRRDDTRNHRSYQNSTMATSSNQRKQTRMRMGKSRVSKTHARKWKPGEKAMREIRQYQKSTDLLIQKAPFCRLVHEIMQEVTSFSSDFRIRAEALGALQEAAEAFLVEMFEGSVLLANHAKRVTLMPTDIQLYRRLCLRPV

>C_panamensis_HCP3

MLQMEDLDGPRIEELPASPEREPAALRDNNRNGNVQRALPAHFENELRRLMSDPNFSKDAELMTDAIELMKRQVNKMEDDQDMYGYEGGMAEMIRNLRIRIVSFTKKRDDAIARFREEREASRQMERTRRVNNTDFDMTDAENRTRLHPSMSSQRSYSYDRRDPYSSDENDDTYQPARHNAQQRMSRSPSPLMHQSHHSSINQRRDDSRQHRSYNNTATSSRPTTSNRSRMRVGKNCVTKTKNRKWKPGEKAMKEIRRYQKSTDLLIQKAPFVRLVHEIMADVTPRSSEYRIRAEALGALQEAAEAFLVEMFEGSVLIANHAKRVTLMPTDIQLYRRLCLRDK

>C_panamensis_HCP3L7

MLQMEDLDGPRNEEMPASPEREPAAEAQFENELRRLMNDPNFHRKADLMGKAIELMKRQVNRLEDDQDVYDDDASRLVERTRRVNNTDLDMTDAENRTRLRSSVSSQRRDPYSSDENNDTDQPARHNAQQRMSRSPSSLMHQGHGSTINQRRDDNRRQRSYNNTATSSRPTTSNQSRMRIGKNRVTKTKNRKWKPGKKAMNEIRRYQKSNDLLIQKAPFVRLVHEIMADVTPRSSEYRIRAEALGALQEAAEAFLVEMFEGSMLIANHAKRVTLTPTDIQLYRRLCLRDK

>C_nouraguensis_HCP3

MDEMDGPRIEEVFDSPRRETSVLRETNQRANIRKRAPHPYEIQLKNLMKRPGFNQNAELMGEAIETMKKLCDVREREDETFGPTEGSTEFIRELKDRIRLYTTQRNEAIEQYRQEKENSRQAEIARRRADYTNLDITDRENRTRLHPSSSSHRSYEQRNLRASSDEDEYDDQASRFRSQHHRSRSRSPAHSHTHHSTMNYQRQDDTRAHRSYQNSTAASGPNSRPKNQTRLRIGKSRVTKTNARKWKPGEKAMREIRQYQKSTDMLIQKAPFCRLVHEIVQDVTSSSSGFRIRAEALGALQEAAEAFLVEMFEGSVLIANHAKRVTLMPTDIQLYRRLCLRNI

>C_becei_HCP3

MDEMDGPHIEEVFDSPRRETSVLREANQQANVRKRTRSQYEAQLMNLVKRPDFNRNAELMGEAIEIMKQQSDEMEREEEMFGPNEDSAYIRKLKDRIKAYTKKREDAIEQFRQERENSRHTEMARRRANYTNLDITDRENRTRLQPSSSSYRSYEQRNLRASSDEDEFDDQTSRFRSQRQRSRSRSPAHSHTYHSTMNYQRQDDTRTHRSYQNSSAAAGSGSRAKHQTRLRIGKSRVTKTNSRKWKPGEKAMREIRQYQKSTDMLIQKAPFCRLVHEIMQDVTSSSSDFRIRAEALGALQEAAEAFLVEMFEGSVLIANHAKRVTLMPTDIQLYRRLCLRNI

>C_yunquensis_HCP3

MLQMEEMDGPHIEEVFDSPRKEASVLREANQYANVQKRVSSKFEIQLKKLMTRPDFNRDADLMSEAIELMKRQAAELEDNQLEYGSTAESSAYIRKLKDKIATYTRNRDEAMERYREEKEESRRMELARRRADFTDLDITERENRTRLQPASSSHRSYDQRTLRSPSDDEEYDDQPSRYRSQRQRSPSTSPVRSQSYHSTMNYQRRDDSRNHRSYQNTAAASSSNTRNTNQGRLRIGKSRVTKTKHPKWKPGEKAMREIRQYQQSTDMLIKKAPFCRLVHEIMQEVTGFSSDFRIRAEALAALQEAAEAFLVEMFEGSVLIASHAKRVTLMTSDIRLYRRLCLRNL

>C_macrosperma_HCP3

MLQMDELDGPHIEEVFDSPRKEPSVLREPNQYPNVQKRGFSKFEMQLKTLVTSPGFNRDADKMTEAIELMKAHTRELEREERMYGRSEQSSAAIAKLRDKTESFIEKRDTALARFHAERDESRRLETARRRADYTDMDITARENRTRLPSSSYRSYEQRTRTYDSDEEEYEDQTSRFRSQQRSRSRSPIQSQSYQSTMNIPRRDDTQNHRSYRNTTMASSSQPRQQTRVRIGKSRVTKTTARKWKPGEKAMREIRQYQKSTDMLIQKAPFCRLVHEIMQDVTSFSSDFRIRAEALGALQEAAEAFLVEMFEGSVLIANHAKRVTLMPTDIQLYRRLCLRNM

>C_sulstoni_HCP3

MLQEHNGPIITEMVEPPSRDPSILRDSGRQYNVERNRPTAIEKQIKELFERPGFNDNLEAMEQIISLMKKQIVEWQREQDLYGPTEERAKKIRAWQKNRETYIRNVNDANSRREELRRERDESSRRMEMARRRVDMTQLDITARHNSTRLQQSYSQRSYDQRTRYESDEENDYVPQSQRHRSRSPPRAHQSYHSHSSQSQQARRPDTSRHVDRSQNDFSSAENPTASSSSQQQQRKPKPRLRPGKSRVTKNLWRPRKDRALQEIRQYQKSTDLLIQKAPFCRLVQEIVRESSSSTSDFRVRADALSALQEAAEAFLVEMFEGSQLIAAHAKRVTLMPSDIQLYRRLCLRNI

>C_afra_HCP3

MLHQHVGPVITEMEEPASRDSSILRDSGRRHNVARSEPTAIERQIQDIFNQPRCNERPEAMAEAIRLMRKQIDQWEREQDLYGPTEERTKNIRIWKNNRRKFIAQLEVAKERQERARRERDESSRVVETVRRRGDMTETNVTAIHNSTRLQSSQRSYDQRTHFDSDEEENGYTSRAPRPLPRSPPRAHQSYHSNLSHQSRRLDASSQDNRSRNDYGSNNVTSVTSSSHQGKPKPRLRAGKSRVTKNMFRPRRDRALLEIRQYQKSTNLLIQKAPFCRLVQEILREVTSSSDYRSSDYRIRADALSALQEAAEAFLVEMFEGSQLIATHARRVTLMHSDIQLYRRLCLRN

>C_afra_HCP3L8

MNNSRNTRLKSDIVPGRRIIPDVIYRDAGIEEEPSYLAERTMDESVVNSTRNTRWNSNIVPGRRIIPDVIYRDSGIRGEPSYLAERTMDESVKSKRIGLGEREKRAADPTRSTMSSMIEDISSPGVQFFENNREKMRPTVATPRRSNLGANRLSTASDKEKTIDMLSIGGEPTIEANGSSYVEPVSVNGSRVSPTREDMLHRHVGPVITEVEESPSRDPSVLRDSGRRHNVARSKSTAIERQIKLIFIEPRFKERPEAMAEVIRLMRKQIDQWERDQDSYGPTEERTKNIRAWKDNRRRFIEQLEDAKERQERARRERDESSRVVETVRRRGDMTGMNVTAIHNSRSYDQRTHFDSEEEEKENGYTSRAPRPLPRSPPRAHQSYHSSLSHQSRRLDASSRDNRSRNDYGSNNVTSSSHQGKPKPRLRAGKSRVTKSMFRPRRDRALLEIRQYQKSTDLLIQKAPFCRLVQEILREEATTSSSDYRIRADALTALQEAAEAFLVEMFEGSQLIATHARRVTLMHSDIQLYRRLCLRK

>C_sp49_HCP3

MQRMDGPIIEEVVDHDARQEKRNRRINRIKDLMARPETKESKELLKKMLDLLQEHLNDLEDEQLEEGANHRDAIATLRHKIDIFTPLYNGAVADYMARKEESRREEARRAMSFGSSQNNITGRDNRSKLHQSSQRTYSSEDEENDEEYAGRNRQDARRYHHQSPEPQASSSARRDNTRRVDASSNNPRMRAGKSRVTKTNSRRWRPGQKALSEIRKYQKSTDMLIQKAPFHRVVQEILCETSGFTNAHRIRADAISALQEAAEAFLVEMFEGAMLLSNHAKRVTLMASDIQLYRRLCLRKF

>C_sp49_HCP3L9

MARPETKESKELLKKMLDLLQEHLNDLEDEQLNEGANHRDAIATLRHKIDIFTPLYNAAVADYMARKEESRREEARRAMSFGSSQNNITGRDNRSKLHQSSQRTYSSEDEENDEEYAGRNRQDARRYHHQSPEPQASSSARRDHTRRVENSSNNPRMRAGKSRVTKTNSRRWRPGQKALSEIRKYQKSTDMLIQKAPFHRVVQEILCETSGFTNAHRIRADAISALQEAAEAFLVEMFEGAMLLSNHAKRVTLMASDIQLYRRLCLRKF

>C_sp25_HCP3

MQRMDGPHIEEVFDSPRRSETRQSDRQWRIGRIQDLVHNPETGQSKDLLKEVLDLLKEHRQDYEDEQDIDGVDHRQTIAKLTDKIDMYTPCYYRAVDDYNMRKEESRREEARRAVSFGNSQHNITGRDNRSKLYQSSQRNYSSEDEENDEYVSRSHTRHADSRYESQDRDRDTRDYQSSSSQAHHHHHHQRDDSRRHQTSSFPKQPRMRAGKSRVTKTNSRKWRPGQKALAEIRKYQKTSDLLIQKAPFYRVVQEILRETSGFTNDHRIRADAIAALQEAAEAFLVEMFEGSALLSLHAKRVTLMPSDIQLYRRLCLRNF

>C_imperialis_HCP3

MQRMEGPYIEEVFDSPPRRHNSNRRELRVARLRELISDPATRESRDMLREVLDLLKEDKRHFQDEQDNENIDHQKTISTLANKINQFERMYDRAVDDYEARKEESRRIEEARRTASFGGSHHNVTARDNRSKLYQSSQRHYSSDEENEEYATRDRRQESRRDTRQDTRNYDSDGNDTRDHQSSSSAYQRHDDTSRRQQSTNQQSSSSAPIRMRPGKSRVTKTNSRKWRPGQKALSEIRKYQKSTDLLIQKAPFYRVVQEILRETSGFTNDHRIRADAIAALQEAAEAFIVEMFEGATLLSTHAKRVTLMPSDIQLYRRLCLRHL

>C_japonica_HCP3

MQRMIEMGGPHIEEIVDPPSPSNSVLQEADYRQNGPSRSRPKLIDQIRTLIRTPNFNKDAVKMGQAIDLMELQIAEWVEEQIRYGFTQEREDAIYQYRRKVIVFKKSVEEAEERYFEEREESRRREEARRAMSYSRGDISARDNRSKLHQSHTQRNYGDSLDSDDENERENGYQSYRPPPQRQQRLRSRSRSRSSSPMRSSYRHESPENSRRNASHQQTAQVRMRAGKNNVTKTKKWRPGQKALSEIRKYQNSTDLLIQKAPFRRLVHQIIQEATGFDSGFRIRADAMSALQEAAEAFIVEMFEGSVLISNHAKRVTLMTADIQLYRRLCLRNL

**Supplementary Data S3: All HCP-4 sequences used in this study**

>C_tribulationis_HCP4

MNRKRGIRSSIVPGRRPITKIVALHEVGMDEDMSYMNEKTLLDNSSQMDDTVDAEEREWRKKGLSENQIQRLFEARRQKQLKDAHDRLHKEKYGAKGFDEMLKLPQYASGRESEDDDDCHDENKSSSEIQVFAIPALPKHLNEKSIMGSPVAGSRGSGKAGLSCSTPKSGNDVSMRSLRALDISHVVSTDHLDATKTTTHTKNIMIPVVESREESSLQKTHTIVKDASSELKRTFTVVKDIEVSDADKEATKNKEDSSLQKTFNVIENSENDKTYVVVDSNEKEKESSLQATFVLSKDWNDSLMPTEEKRKFESSKKDESSDHVQQVYVPGTSNQPTPNISVQADLLIVQNSTQGGEIKKSNGVGAQKREKRGANMSTSLMTSMIEDVPSPGANYFKNPRKKLRNTAKSPSKSKAIARLSTESDKEKTIDMLSIAEETSINAESVSSSYIEPISSDGTRVSPILEVEETMEVVTTTPKSSRRGTAFSVRGSSMEKARGGIFCATPTAPTVDPTSPLVTPKLNYQKPTVSSMLKSKGPSDLNHDLCATRGKNRPSVNNVVETSLSPSPDNNSTAKLDEQETDAQSKVIEESISDENAVRNRTPGIEHQPSIGVDLPMDAMTIQSNNSNQHMNDFDMDYGDDVELHQYSEEKNRSGGGPSTRQRNRQRIGLLSDSIATINTPGVDRRPTRHLSRNETIPEGDWSDEEFEEMGRRRNGGKHQRDVGLQLKKREIIQPDDTSDGVRRSQRTRVKPVRSWLGEQPVYINSPSGGKRLTGVTDVIIKDKRLCKYKTADLRVATEREQKAKARRKEMAARRREQLARDHRRGRRLNESQEDIHTDDDDDDEMS

>C_sp41_HCP4

MNRKRGIRSNIVPGRRPITQFAALHELGMAENMSYVNEKTILDESSLMNDTVDAEEREWRKKGLSENQIQRLFEARKQKRLKDAQDRLNKEKYGARGFDEMLKLPQYYSGLESDDECHDENKNSSEIQVFAIPALPKHLSEKSIMGSPVAGSKGSGKAGLSCSTPKSGNDVSMRSLRSLDISHVVSTDNLDATKTTTHTKNVMIPIVESQEESSLQKTHTIGKDASSEVKRVFTVAKDTDISGSEKNMTKNKEDSSLQKTFTIAENCDNEEISVVVDSDQKDAESSLQGTFVLSRGWNDSLMSTGEKRKNDQLKNIESADHVQQVGSPGTSNQTIPNAGTSVEVDLPIAQNSTQGGEKRKSSGVGGEKREKRGANMSISMMSSMIEDVPSPGANFFKNPRKKLRNTAKSPSRTKAVARLSTESDKEKTIDMLSIAEEASIDAESVSSSYVDPISSNGTRVSPIPEAEETMGAVTTTPKSSRRGAGFSVRGSMMEKARGGILCTTPTAPTFDAISPLVTPKHHYQKPTVSSMLKSKGPPDIDQEMCITRAQNFTSVNNAVATPVRPLPSNNSSVDDEEQHARSKTAERDISEENEIRNRTPDIVHQPSNDIDLPVEAMTIQSSDSNQRFDDFDMDYGDEIAPHQYSEERESNASGNGPSTRRNRHRVGLLSDSIATINTPGIDRRPTRHISRNESIPEGAWSDDEFEEAGRRRNGGRHPRDIGLQLKKREIIQPEDTSDGVRRSQRTRVKPVRSWLGEQPVYVNSPSGGKRLTGVTDVIIKDKRLCKYKTADLRVATEREQKAKARRKEMAAKRREQLARDHRRGQRLNESQEDIHTDDEMS

>C_zanzibari_HCP4

MDRKRVIRSNIVPGRRPITEIAALHEVGIAQNMSYMNEKSLLDESSQVNDTVDVDEREWKRQGFTENQIQRLYEDRKQKQLKDARDRLHKEKYGAKGFDEMLQLPQYSSGRESEDDDEYHDENTNNIEIQVFAIPALPKHLSEKSIMGSPVAGSKGSGKAGLSCSTPKSGNDVSMRSLRSLDISHVVSTDKLNPTRTTTHTKNIMLPILESREKPSMQNTHTIRKDVSSELEGTCIVAKDDEVGGTRVESISQNKEISSLQKTHTIRKDASSELQGTFTVANDVEGAGSKTEADKNDPESSLQGTFVLSKDWSDCLISTQDKHGAELSKKKVDNVQLVDSPGTSQQTMSNVVVQVNPSIVEVSMQGGEKGRFNGVGAQKREKRGANMSNSLMASMIEDIPSPGANYFKNPRKKLRNTAKSPSKAKAINRLSTDSDKEKTIDMLSIAEEASIDTESVSSSYVDPVSINGTTQVSPIPEIDEPMDVATTTPKSSNSRTAFSIRRGLMEKARSEILCATPPAPTENNTSSLVSPKLHYQKPTLSSMLKNKGGLNSDVLLSATPRGKRAAVNNEIEVSQRPILNFDTSANLDHQETNDQTEPVEERAYNTNETRNSTFDVVNEPPNDIDLPIGEMTIQSNSNQHVNNNLDMDYDDDIEPNQGLEENESNASRGGPSRGRRHRVGLLSDSIATVNTPGIDCRPTRHLPRNEEISDDSSVDDESEEVGRRRNSGRHSREVGLLLKKREIIQPDDTSDGVRRSQRTRVKPVRSWLGEQPVYVNSPSGGKRLTGVTDVIIKDKRLCRYKTADLRVATEREQKAKARRAAIKREQLARDHQRGRRLDVSQDDIHTDEE

>C_sinica_HCP4

MKQXXXSGMDENLSYMNEKTVLDESSQMNDTTDVEEREWKKKGLSENQIQRLFEARKQKQLKNAQERLHKEKYGAKGFDEMLRLPQYASGRESDEEEEYQDENKTITEIQVFAIPALPKHLSEKSIMGSPVAGSKGSGKAGLSCSTPKSASDVSMRSLRSLDISHVISTDQLDVTRATTHTKNVMLPIVESREESSLQKTHTIKKDASSELQRTYTVAKDAEVVNTDTESASKNKDDSSLQKTFTVAGNPESEETTSSLQKTFTVEAAEGKERESSLQNTFVRSKDWDNSFASTKENHKSENSKKIESTDKEQVVVSPKGAKQPMPNASIHVEVEPLIVESSIQGGERRKFNKTGAREKRGANNSMMASMIEDVPSPGANYFKNPRKKLRSTAKSPSKVKAISRLSTESEKEKTIDMLSIAEEASIDAESVSSSYVDPVSNNGTRVSPVPEVEEPMEVVTTTPKSSLRGTSYPIRGSLMEKARGGLVCSSPTAPNLNNESLLGTPKLNYQKPTVSSILKSKGAPDCNDILGATRREHRTPANNAISIPSRQSPVHNSPSNLKDDHEPSGLRKTPEKDISEVPEVRNRTSEIFHQPPNDIDLSMGAMTIQSADSNQRVNYDYDMDYGDDFEPYQDSDIREEHESESGGPSRPGKIRHRVGLLSDSIATVNTPGVDRRPTRHVSRNETIPEDSWSEDELEEAGRRRNGGRRDRNVGLPLKPRELIKPGNTSDGVRRSQRTRVKPVRSWLGEQAVYINSPSGGKRLTGVTDVIVKDKRLCRFKTADLRVATEREQKAKARKKEMAARKREQLARDHRKGRRLNESQEDILTDDEMA

>C_nigoni_HCP4

MERKRAVRSTIVPGRKPITNIAALHEAGLSEDQSYLNEKTVLNETTQLNDTRDVEERKWRKEGLDENDIFKRQEARKQKQMRDAEDRLNREKYGAKGFEEMLALPQYSSERESDEDDHNEENQKAPEIPPIFAVPSLPKHMNEKSMMGSPVAGSRGSGKAGLSCSTPKSANDVSMRSLRALDLSHVINTDHLDANKTIVQTKNVLLPIVENSEESTLQRTYTIHEDGNNERNLPTPEENEIADNEVRKFATPEDTKIVHNTMEGGEQTRKATKVGAQERERRGANLSMSLMHSMIEDVPSPGANYFKHPRKKVRPETKSPQKPRVMGRLSTESDKEKTIEMGSIAEESSMTESIGSSYVDPVPENTTAFSPVPEEEEPMEIANTTPKSSRRGSSFVTPHSLMERSRGATLFSTPVPPVVDAITPKLNYQKQTVSSALKMKGAPNGCELLDVDRRCRSVPKKSANVARESPKDKGDGNGKPEDIVIDDGKVNDEEMCDVTVEMPQNAEQAASGVELDMNGLTLNGNSANVSYNLDHDGFDDFHGDPNLEDERNESEDAGPSTRRTARSRIGLLSDSIATVNSPGVDRRQTRRNFRNDTIPEDSWCSDEEVVPRRRNDARNAVKIGLQLKKREIIQPDDAANGVRRSQRTRVKPVRSWLGEKAVYVNSPSGGKRLTGVTDVIIKDKRLCKYKTGDLVLANEREQKTKARKKQLAAKRREQLSRDHQRGYRLNESQEDIVTDDELCDYT

>C_briggsae_HCP4

MSYLNEKSVLNETTQLNETGDVEEQVWRKEGLDENDIFKRQEARKQKQMRDVEDRLNREKYGAKGFEDMLALPQYSSERESDEDDHNEENQKAPEIPIFAIPSLPKHMNEKSMLGSPVAGSRGSGKAGLSCSTPKSANDVSMRSLRALDLSHVINTDHLDANKTNGEESDLQKTHTIQKMASSEGSTLQKTHTIHEDGSNERNLPTPEENDIADTDVRNPSNGGEQKRKAKKVGAQDRERRGANLSMSLMNSMMEDVPSSGANLLKHPLKKVRPETKSPQKPRVMGRLSTESDKEKTIEMGSIAEESSIAESMGSSYVDPVPENATAFSPVPEEEEPMEIANTTPKSTRRGSSFVTPHSLMERARGGTLFSTPVPPVVDAVTPKLNYQKQTVSSALKMKGAPNSCELLDVDRRCRSGPKKSVNVARESPKDNDDGNGKPEDVVIDDSKLNDEEMCDVTVEMPQNVEQAASGVELDMNGLTLHSANVSYNLDHDGLDDFHGDPNFEDERNESEDAGSSTRRTTRSRIGLLSDSIATVNSPGVDRRQTGKNYRNDTIPEDSWDSDEEVVSRRRNDVRNTVKIGLQLKKREIIQPDDATNGVRRSQRNRVKPVRSWLGEKPVYVNSPSGGKRLTGVTDVIIKDKRLCKYRTGDSF

>C_remanei_HCP4_paralog1

MKQKHGSRSTIVPGRKPITQIAALHDAGVTEDMSYCNEKSVLDESSNMNDTHDNDIVDERSLIQQGIPDRDINKILQRRKLEKLLKVSQAQGRLERQMLGAKTFKELMEKHEYSERESGDENNTAENISITNKPVFAIPALPKHFREKSMMGSPIAGSKGSGKAGLSCSTPKSVNDISMRSLRILDISHVVNTDQLYYDKVVVHNKNVLIPIVENTEQSSVMGETFTVRDDQCDNKKHGQVSANEASDSPIGSIKLDRNCISKELIDTTITENIVEGGKQRRRSNKVGVQERERRHADLNSSLMNSMIEEVPSPGAKYFKNPRKKVRPIAEVPPKVFNMLSVESDKEKTIEMLSIVEEVSIEAESNGPSFVDPLSVNGSHISPIPEVNELMNTANVTPKSSCPYIPILGNLMETVRVTQENDVSSVITPKLNYLKPTISSLRKNINAPECDDILFGTRRERCTPGKNTTTAKRGVQDAPTIDKTTVTVERITVKNDQRNIASNEDLSIGVPVETNRPSFNLELEMGEMSVRSSPKISNANLDSVEPADFDPTEDRERVENQPGPFNQKSSRNRVALLSDSIATVNTPVYDYNPARCLEPDTNNGNRRSTRTRVKPLRFFLGERAVYVNSPNGGKRLTGVTTVIIKDKRLCKYRTGDLKLATEREQRAKAFKKKSAAGKRKRLLRDQQAGRRMDESSYDIHTDDEQ

>C_remanei_HCP4_paralog2

MERKHGWRSTIVPGRKAITQIAALHDAGVTEDMSYLNEKSVLDESSNLNDTHDDDEERALRKKGFSEREINKRLQGQKVEKLAKAHGRLKEQMLGAKNFKELMEKHEYSERESDDENDTAENITITNKPVFAIPALPKHLSEKSMMGSPVAGTKGSGKAGLSCSTPKSGKDVSMRSLRVLDISHVVNTDQLDYDKVAVHNKNVLIPIVENTEQSSAKMGETFTVRDDQDDNQKHGHVSANDASLQKTFTVDPRTEGDKSESSLQKTFTVPGGDQDSTRHSSLQNTFVKSGANDSLLPNNGKRAQDDTKKIDESVNADQSGSVIDPSKLAGNCISNELIDTTITENTVEGGEQRIRSNKVGVQERERRHADLNSSLMKSMIEEVPSPGANYFKNPRKKLRPTVEVPPKIISRLSVESDKEKTIEMLSMVEEVSMEAESNGPSFVDPLSVNGSRISPIPEVDKLTNPANVTPKSNRRNLPIRGNSMETVRVTQANDVSSVITPKLNYLKPTISSLRKHVNEPECDDTLFGTRRERCTPGKSATTAKTVVQDAPIIEKTSVTGEGVTVKNDQRNDASNRDLSTGMPEETNRPSFNLELEMGDMSIRPSPKRLSANLDSVKPADFDLDLPTGRIENQAGPSNQRSSRNRAALLSDSIATVNTPGYNRTARCRVMNDTNVEESWQSDEDDVILSRRNPGKNGKNVGLQLKKREIIQPDTNNGNRRSARNRVKPLRSWLGEKAVYVNSPSGGRRLTSVSDVVIKDKRLCKYRTADLKLATEREQRAKAHKKELAARKREILLRDQQAGRRMDESHYDIHTDDEE

>C_latens_HCP4_paralog1

MKQKLGWRSTIVPGRRPITQIAALHDAGVTEDMSYLNEKSVLDESSNLNDTHDDDDERSLIKKGIPERdinkelqrrkleklmkvqaqVRLEGQLLGAKTFKELMDKHEYSERESDDENYTAENISITSKPVFAIPALPKHLSEKSMRSPIAGTKGSGKAGLSCSTPKSGKDVSMRSLRVLDISHVVNTDQSDYDKATVQNKNVRIPTVENSEHSSVARTKIGKTFTVRDDRDNNQKHGQVSANEASDSSIGSIKLAGNCISSELIDTTISGNTVEGGKQRRRSNKIGVQERERRHADLNSSLMKSMIEEVPSPGANYFKNPRKKLRPIAEIPTKVINRLSVESDKEKTIEMLSMVEEVSMEAESNGPSFVDPLSVNRSRISPIPEVNKLMNTANVTPKSRPPNLSIRANLMESVRVTQTNDVSSIITPKLNYLKPTISSLRKNVNAPECDDSLFGTRRERFTPGKIATTVKTVVQDVPSIEKTPVTVKGITVKNDQRNIARNKNLSIGVPVETNRQSFNLELEMGEMSTQSSPKKLNANLDSVKPADFVWDLPAEDRERIENQPSSSNQRSSRNRAALLSDSIATVNTPEYDRISDRCYSINNKNVVIPRRRNLEKNRKNVGLQLKKRVIIEPDDRNNGNRRSTRTRVKPVRSWLGEKAVYVNSPRGGKRLTGVSDVFVKDKRLCKYRTADLKLATEREQRAKAHKKELAARKREQRLRDLQAGRQMEESHYDIHTDDDE

>C_latens_HCP4_paralog2

MERKHGWRSTIVPGRKAITQIAALHDAGVTEDMSYLNEKSVLDESSNLNDTHDDDVERSLRKRGFSERDINKEIQRQRNAKLAKAQGRLEDQMFGAKTFKELMEKHEYSERESDDENDIAENIPTTNEPVFAIPALPKHLSEKSMMGSPTAGTKGSGKAGLSCSTPKSGNDLSMRSLRLLDISHVVNTEQLDYDKVTVHSKNVLIPIAENTEQSSAKMGETFTVRDDQDANQKHGHVSAYEASLQKTFTVDPRKEGDKSESTLQKTFTVQQRAESDKSESTLQKTFTIPGGDEDSTRHSSLQNTFVKSSSNDSLLPNNGKRAQDEKKKLDETVNAAQSDSEIDSSKLAGNCISNELIDTTITENTVEGGKQRRRSNKVGVQERERRHADLNSSLMKSMIEEIPSPGANYFKNPRKKLRPTVEVPPKIINRLSVESDKEKTIEMLSMVEEVSMEAESNGPSFVDPLSVNGSRISPIPEVDKLMNTANVTPKSNRPNRPIRGNSMETVRVTQANDVSSVITPKLNYLKPTISSLRKNVNGPECDDILFGTRRERCTPGKSATTAKIVVQDTPIIEKTPVTEEGITVENDQRNVASNSDLSTGMPEENNQPSFNLELEMGDMSIRPSPKRLNANLDSLEHVDFDGDLPTEDRERIENQPGPSNQRSSRNRAALLSDSIATVNTPGYGRNRRRFMNDTNVEESWQSDEDDVVLGRRNPGKNGKNVGLQLKKREIIQPDTNNGNRRSTRTRVKPVRSWLGEKAVYVNSPSGGKRLTGVTDVVIKDKRLCKYRTADLKLATEREQRAKAHRKELAVRKREQLLRDQQEGRRMDESHYDIHTDDEE

>C_sp51_HCP4

MENKRCFRSSIVPGRKMITNQVAYYQAGIIENMSYLAEKTIADDSSILNDTHDNEEREWRKQGLTEKQIYHRLQDKKLQKLRNAENRLNKEKFGAKNFQELLNKHEYSQRESEGEEEATTEKFDVFAIPALPKYLSEKSMLGSPVSGSKGCGKPGLSCSTPKSANDVSMRSLRSLDISHVVAVDQLDPKKVNIQSKTILIPIVQNSESSSLQKTHTSQKDQNSESLIGTKANSSLQESHTIPNSPSQSQTYTSVQKTYIIPSSSPHKKQDDHQDQTEPLHEKDKETSLQESSNVDIERETSNSDTLVVGTGLGNTLLDSVRKDQENEVQKLLNNNGDGLQEISSSNDVHIPVVQNLLQGGKKKSATNKIGLQEREKRSNDSLMRSMIEVVDSPGAGFFKNARKKQRPIKTPTKMKVNRLSTESDKEKTIDMLSIAEEASMDNETNGSSFVDPVNGSKISPIREVNEPIDHSKSETTPKSRPPTTPGSSIREKARAQRILEPLACNSFVVNNEIRLISTPKHHFQKPTFSSLVKNRNVDSNALLESSRRERINLPRRDITILKEAVPASLDEGEDQDDGTQNYFESTSDKTNRPDDDMSSKFDEISSHSTSDINDDLREMTVRDVPSMEIDYDPVDYVERDNQYDNSDVNTEHNERDSVNAGPSTKQKSFRKKIGLLSDSFASGSAPGIERRSTRHNFGNDTIVDDTWQPDEAPKRRNNKGRNGKNMGMQLKKREIIEPSTPNDEVRRSKRTRVKPTRSWLGEKPVYVNSPSGGKRLTGVTDVIINDKRLCKYRTADLKLATEREQKEKALKKMKAAERRRQLAADQKRGRRLNESQEDINTDDEYED

>c_sp44_HCP4_paralog1

MMKKHKERSKIAPGRRLITPQVAYHQAGIIENMSYLAEKTVVDDSSILNDTQDDEERQWRKQGLSEKDIFRRLENRKLKRLKNAENRLQREKFGAKDFQELLNKHEYSERESDDEEENKNDGNPKPIEVFAIPALPKHLSDKSMMGSPVYGSRGSGKAGFSCSTPKSANDISMRSLRALDISHVVSVDQLDVNKVTIQNMKITIPITDKCESSSLQNAKTTAEDQNSTPQNGEGEDLTLQKTHTVQSASLQKTYTIQDSSLQKTHTIQKPSLQETNTTSDSSLQETHTIQKPSPPKTIIITGLSLQETPTIQEPSLQKTITISNSYLQKTHIIHEEDNIRILKKKNSDSIVQDEPATVKVPLVMNQLQGGKKNQTTKGIGQQERGKRTANSSLMSSMIEDVQSPGAGLFKNTRKKLKPMVVTPQRMKNNRLSTESDKEKTIDMLSMAEEASMDNETNGSSYVDPISVNGSRVSPVAEVDEPMDVSKRSETPKTSRLTPRSSNIREKARAESIFATPSSDLKLNEDEVSLATPKHHYQKPTFSSIVKSKNSLERNDLLEVTKQGRRNRSCRNNLIDEHPQQGIPDEVVDTQRENNTGKEKENSNDSDVSGKESDNSSRSVDNIQAELEEMTVSGNPSTGFDYDSLYYNDQDNQVALCDPESPEEEEDGGPGTSTRQKAPRRNVLLLSDSIATINTPGERRSTRRYSDTLAEETWVPDLVSKRSNRGRNSKNTEMQLKKRVIIKPDTPADGVRRSTRTRVKPVRSWLGEQAVYANSPSGGKRLVGVTDVIIKDKRFCKYKTADLALATEREQKERAWKRNKAAEKRRQLAADQSRGYRLNESQEDIKTDDECDE

>c_sp44_HCP4_paralog2

MMGFPVYGSKGSGKAGLSCSTLKIANDTSMRSLKALNISHVVAGLDANKVTTQNRKITIPIMEKWESSSKQKAKTTTEDQYSTLPNAKGEDLTRQKTRTVRSASIQKTNTIQDSSLQKTRTIHEEDEMRILKKKDSDSIVRDEPATVKVPTVMNQLQGGKKNQTTKGNGRQEREKRTANSSLMSSMIEDIQSAGAGLLKNTRKRLKPMVVTSPGMKNSRLRTESDKEKTIDMLSMAEEASMDNKTNSSSYVDQISVNGSRVPPVAEVDEPMDVSKRPEKPKTRRLTSRGSNIREKARAGSIFAIPSSDWKLNEDEVSLAIPKHHYQKPTFSSIVKRKSSLERNDLLEEMNQSRRNRSCQNSLIDEYSQREIPDEEVNTRRENNITKEKENSNTTDVSGKESDNISRSLDNIQAELKEMTVSGNPSTGLDYDSIDYNDQDNQVALCDPESPGEEDGRPESSTPQKAPSPRFIANSVSSKTEHRAFVGEKKHREFVGEPMGDKEVTCIAGIGPTYGTKLTDAGFDKAYVLFEQYLFLKKDEDLFVDWLKETAGVTAKHAKSAFNCLNEWAEQFL

>C_sp48_HCP4

MVNKRYVRSSIVPGRKMITTQVAYHQAGISENISYLAEKTVVEDSSILHESHDNEEREWKKQGLSEREIYDRLRAKKLKKLRNAENRLNKEKFGAKNFLELLHKHEYSERESDDEENATVEMDVRNVEVFAIPALPKHLSEKSMFGSPVSGSKGSGKAGLSCSTPKSANDLSMRSLRSLDISHVVAVDQLDAHKVTIQSKNILLPIAENAESSSLQKTHTIQSSSPAKTFTIPDSSLMKTHTITRDLNTDSNNERKEESTLQKTYTIPSSSPQKSESIQDGSLMKTYTVQDSSLQKTFTVPDSSVQKNAVVSNTSPSKNDNAQNNQSCQLQEEGEDMSLQKTFDVDADEQNTKCEGTFVVQSGWRNTLLDSLRQDQEAQISKKGNGDLLSPSSSANIVLPVLENKLQGGEKKIGPKKTSLQEREKRAANSSLMSSMIEDIPSPGAGLFKNTRKKLRPTNATPQRMKANRLSTESDKEKTIDMLSIAEELSMENETNGSSFVDPVSINGSRVSPVIEVDEPIYISKNEATPKSRRHTLLGSSLRERARAQSILASPASAVGVVRDIAPADITPKHHYQRPTFSSLVKNKTSVERNDLLEATKKDRRNTPHRNTKTQEEAASLSGDEGVEDIHSETVPLSGEVDDCAEISSKPADISSHSITNIRLDLDGMTVRDVPSMDIDFDAVDYVDHNHSDQFVAESPEESEGESDEAGPSSRKKSSRRKFGTLNDSVTSLDARRSGSHSNFANDTVSNETWQPGETSRKNSRGGKNKPLEMQLKKREIIQPASPQGQVRRSERVRVKPIRSWLGEKAVYVNSPRGGKRLTGVTDVIIRDKRLCKYRTADLKLATEREQKEKAYKKSVAAEKRRKLAADQRRGRRLNESQEDIITDSEDDNEK

>C_brenneri_HCP4_paralog1

MVNKRYVRSSIVPGRKMITTQVAFHQAGISENISYLAEKTIVEDSSILHESHDNEEREWKKQGLSDKEIYDRLRDKKLKKLRNAENRLNKEKFGAKNFMELLNKHEYSERESDDEENATVEMDVRNVEVFAIPALPKHLSEKSMFGSPVSGSKGSGKAGLSCSTPKSANDLSMRSLRSLDISHVVAVDQLDANKVTIQNKTILLPIAENVESSSLQKTHTIQSSSPPKTFPAQDSLLFMKTHTITRDSNTDSNNERDEESTLQKTHTIPSSSLQKSESIQDGSLQKTYTIQDASLQKTFTVPDSTVHKNVIDSNTSPSKTDDAQDNQSCQLQEEGSDLSVQKTFDVDADEQNSTCEGTFVVQSGWRNTLLDSLRRDQEAQSSNIGNGDLLSPSSTANNVLPVSQNHLQGGEKRTGSKKTSLQERERRAVNSSLMSSMIEDIPSPGAGLFKNTRKKLRPTNVTPQRMKTNRLSTESDKEKTIDMLSIAEELSMENETNGSSFVDPVSVNGSRVSPVIEVDEPMDISKNETTPKSRRHTLLNSNLRERARALSILASPASAMKVVEDNAPTDITPKRHYLTPTFSSLVKKKTAVEINDLLEAKKKDIRNTPHRNATIQEEEAVSISAGEGVDDIHSESVPLSQQVDDCAENSSKPVDISSHSITSIRMELDEMTVNDGPSMDIDYDAVDYVDHNDRSDQSDAESLEESEGESDEAGPSSRKKSSRRKIGTLSDSMTSMDARISGNHSNFANDTISNETWQPEETSRKNNRGGRNKTSEMQLKKREIIQPASPQGQVRRSERVRVKPVRSWLGEKAVYVNSPRGGKRLTGVTDVIIRDKRLCKYRTADLKLATEREQKEKAYKKRVAAEKRKKLAADQRRGRRLNESQDDIFTDDDEQ

>C_brenneri_HCP4_paralog2

MVNKRYVRSSIVPGRKMITTQVAFHQAGISENISYLAEKTIVEDSSILHESHDNEEREWKKQGLSDREIYDRLRDKKLKKLRNAENRLNKEKFGAKNFMELLNKHEYSERESDDEENATVEMDVRNVEVFAIPALPKHLSEKSMFGSPVSGSKGSGKAGLSCSTPKSANDLSMRSLRSLDISHVVAVDQLDANKVTIQNKTILLPIAENVESSSLQKTHTIQSSSPPKTFTAQDSLLMKTHTITRDSNTDSNNERDEESTLQKTHTIPSSSLQKSESIQDGSLQKTYTIQDASLQKTFTVPDSTVHKNVIDSNTSPSKTDDAQDNQSCQLQEEGSDLSVQKTFDVDADEQNSTCEGTFVVQSGWRNTLLDSLRRDQEAQSSNIGNGDLLSPSSTANNVPPVSQNQLQGGEKRTGPKKTSLQERERRAVNSSLMSSMIEDIPSPGAGLFKNTRKKLRPTNVTPQRMKTNRLSTESDKEKTIDMLSIAEELSMENETNGSSFVDPVSVNGSRVSPVIEVDEPMDISKNETTPKSRRHTLLNSNLRERARALSISASPASAMKVVEDNAPTDITPKRHYLTPTFSSLVKKKTAVEINDLLEAKKKDRRNTPHRNATIQEEEAVSLSAGEGVDDIHSESVPLSQQVDDCAEKSSKPVDISSHSITSIRMELDEMTVNDGPSMVIDYDAVDYVDHNDRSDQFDAESLEESEGESDEAGPSSRKKSSRRKIGTLSDSMTSMDARISGNHSNFANDTISNETWQPGETSRKNNRGGRNKTSEMQLKKREIIQPASPQGQVRRSERVRVKPVRSWLGEKAVYVNSPRGGKRLTGVTDVIIRDKRLCKYRTADLKLATEREQKEKAYKKRVAAEKRKKLAADQRRGRRLNESQDDIFTDDDDQ

>C_wallacei_HCP4

MKLKGTARSTIVPGRKMITNQVAYHQAGIRENISYLAEKTIAETSSILNDTQDNEERDWRKQGLTEKQIYLRLEDRRLKKIRNAENRLKKEMYGAKDFQELLEKHEYSERESDGEDVTVTEDSQRKIEVFAVPSLPKHLEKSMFGSPAVGVRGSGKAGLSCSTPKSANDVSMGSLRSLDISHVVAIDQLDAHRVTVQSKNVLIPIVENGENSSLEKTFTDQKSQKSVEVSFHQNTPVIQNDMEICEMSSLQKTYIIEENKDAELSSNRKEGSLLQKAFDKDDTDGSKRNSSLQNTFTVVDNWKNTLLDSISEELENGHPKNTTDRAYPSSSSAGTKDATINDVVQPTDTSRIANDTLQGGEKSRSSKKVGLQEREKRANVSIMSSMIEDIPSPGAGFFKNTRKKLRPTNTTPQRTKNNRMSTESDKEKTIEMLSMAEEASVDLESIGPSFVDPISVNGSRVSPPRESEEPMDVTNAVTPKSTRRNNDNGMKAIRVQGTETSEAAESLKVRTPKYHYQTPTFSSLVKNKNANERNDLLESTKRDRRHAPVRSVASVETEKDREKTQLAAQEEEEDLGTQKINDTRHASPAKVVDTSNNSTRDINYELDVLSIRDQPELEDINEIPADREYENDNFDGFNDERRQTYAEEEDECSPSRVALMSDSIASVDFIDRRSTRRNMGNDTVVEDGWQPDEGSRRGNRGRNGSSSGLQLKKREIIQPATPTDEVRRSKRTRVKPVRSWLGEKPVYVNSPSGGKRLTGVTDVIIKDRRLCKYRTADLKLATEREQKEKALRKNLAAERRRQLAADHKRGRRLNESQEDIHTDDEDE

>C_tropicalis_HCP4

MKFKSSSRSTIVPGRRMITNQVAYHQAGIKENTSYLAEKTIAESSSILNDTQDDEERDWRKQGLTEKQIYLRLEDRRLKRIRNAENRLKKEMYGAKDFDELLEKHEYSERESDEEDVTIVAEGHRNIEVFAIPSLPKHLVSDKSMFKSPAVGAKGSGKAGLSCSTPKSANDVSMRSLRSLDISHVVAIDQLDVNRVNVHSKNVLIPIVEHAENSSLQKTFTIQKIQESIERSSTIQKDPETCEISSIQKTIELEEKKTISLQVTQTRVYCLKKHQILIVIQKKLRKVHRFSLRLLSLLMLLCSSQTFRKSWKMLCKVAKKPEVRRKLEGMSMLSIAEEASVEIESNGPSFVDPISVNGSRVSPTGEAEEATVEIESNSPSFVNPIPVNGSRVSPTREAEEATVEIESNGPLFSDPIPIGQAGESVDVSNSITPKSSRHNQDRSLQVASNSSTGDINLELDVPEHQHDDFDGYEGVRRVDLVEEAEEEEQIDHSPRSKQKSFKRRVALMSDSFASVDIPDLNRRPTRRYMGNETVEDDEWEPEKSEKKKSNRNTGLQLKKRVIIEPATPTDGVRRSKRTRVKPVRSWLNEKAVYVNSPSGGKRLTGVTDVIIKDKRLCKYRTADLKLATEREQKERAVKKKKAAEKRHQLALDHKRGKRLNESQEDILTDEEDGE

>C_doughertyi_HCP4

MVKENPKPVRSSIVPGRKMITKYVALHEGGMSENMSYLANKTVMEDSLDCTQDKEEREWKRQGLSEEQIFRNLENRKLKKINNAEHRLMREKYGAKNFLELLDRPEYSELESGNEEPSIAESKARDVVVFAIPALPKHLSEKSMFGSPIAGSKGSGKAGLSCSTPKSSNDESMHSLKALDFSHVIAVDTLEANKVTVHQKDIQIPMKDQANASSVEKTFIINVDHGIPETSSNSETVTVQKDRTSDVANEEDESPCQKTFTVENESKNSSLQKTFTVAPGWKNSFLDSDCEHQKDGISEPVDSTENVEEAGSSDKTRMDDSIMIARISTRSEITENDLQGGLKKNTSKLMGPQASKKRDANSTLMSSMIEDFPSPGAGLFKNTRKKMRPTNATQRKAINRLSNESDKERTIEMLSIVEEPSVHNDSIGPSFVDPCSERLVNTSQVSPVQEKDEQIESSIVDTPKSTKRAVDQSSSMERARARSILSTPNPTILDNSILITPKHNYQKPTISSIVKSKNIDTALFEATKKDCRTRTPGRIDALLEVVEEHTLTDDLEQGNHQDKKGRKNDKSGEDKIEFVPIADKTNSDLNEIRPEMDVSSVREQLEDFDFAADDRIERFGNEKVYDVNEFQEYHGEHERRHKTTTRRVALMSDSVAIENSLLTRPNRRHVANDSTTEEGWQSDEEPVRYNRGRNNKEIGLQLKKREIIQPATPTNEVRRSTRTRVKPVRSWLGEKPVYVNSPSGGKRLTGVTDVIINDKRQCKYRTADPRTATEREQKERAIKKKAAADRRRQLALDQSRGNRMNESQDDIHTDDEE

>C_sp54_HCP4_paralog1

MITHVAACYEAGMMENMSYLMEKTIANDSSLDETQDLEEREWKRQGLSDIEIFKRLENRKLKKLQNAEDRLKKEKYGAKDFDELLERPEYSECESFEKNDENVQNNYVFAIPSLPKHFSEKSMMGSPAIGARGSGKAGLSCSTPKNGHNLSMGSLRGLDISHVCSIDRLDENKVTVHNKNMLIPIIENYDSTCQKTHIIQKDQSSGLKGTFFVARDEEADNQKTNKIRKDTEESSLQDTFLVAEDKNDDKRNSSLQRKSKKVDLQERERRYTNITMSLSNMSNITVGDISSPGSYRFKQIPKKSKPTIKTPPRTKMMNRLSIESDKEKTIEMLSIVEEASIDADSNGPSFVDPLSVNGSCVSPVPELDELMDVSNVVTPKSNRRICNTSIREKFMEMERACSPAAKPTIMNVESNNTLSVTTPKHHYQKPTISSLVKSKNTIDINDLLEASRRDRRHTPSRTAIRTPNIYRTQTVKNATGSPIYEQEEEEEEVVLEDETLNKRNDEKMNKMNDESISNAADASNRSLNNIEFELDSMTVNDQQYSDDFVYDKPQSFSDDRGIDEDTDSEENVSNLRSRQKSTRRRVELLSNSMASTPGIDCRSTTRQGTSNETVTEDTWYPDIDYEVPQRKNKNRKNKVTVGLQLKKREIIKPERTSSNVRRSGRNRVKPVRNWLGERPVYEFSPSGGERLIGVTDVIVTNKRFCKYRTADPKLANQREQKERAIKREIAARRQRDLALDHRRGRRFERFIREYSHERD

>C_sp54_HCP4_paralog2

MEIRKRIRSSIVPGRRMITHVAACYEAGMMENMSYLAEKTIANDSSLDETQDLEEREWKRQGLSDIEIFKRLENRKLKRLQNAEDRLKKEKYGAKNFDELLERPEYSECESEFEKNDENVQKKFVFAIPSLPKHLSEKSMMGSPAVGARGSGKAGLSCSTPKSGHNVSMGSLRGLDISHVVSIDRLDENKVTVHNKNVLIPIVENDDSTCQKTYTIQKDQNSGLQGTFIVARDEEADNQRNNDARKDTEESSLQKTFTVAEDKNDDKRNSSLQDTFVVAEDKNDDKRNSSLQDTFVVAEDKNDDKRNSSLQETFTVAENKNGDKRNSSLQETFTVAEDKNDDKRNSSLEETYTVISVGDMPSPGSYRFKQLPKKSKPTIKTPPKTKMMNRLSTESDKEKTIEMLSIAEEASMEADSNGPSFVDPLSVNGSCVSPVPELDEFMDVSSAVTPKSSRRICNTPIREKSMEMARARSLAATPTIKNVESNNTLSATTPKHHYQKPTISSLVKSKITIDSNDLLEASRRDRRHTPGRTANRTPDRHRTQTVENATESPIDEQEEEEAALKDETLNKRNDEKMNEMNDESISNAAATSNRSLNNIEFELDAMTVSDQQYSDDFDYDKPQSFSDDRGIDEDTESEESVTRLRPRQKSTRRRVELLSNSMASTPGIDRRSTRRQGTSNETVAEDTWYHGVDYEVPQMNNKNRKNKITVGLQLKKREISK

>C_sp54_HCP4_paralog3

MEIEKKIRSSIVPGRRMITHVAACYEAGMMENMSYLMEKTIANDSSLDETQDLEERGWKRQGLSDIEIFKRLENRKLKILQNAEDRLKKEKYGAKDFDELLERPEYSECESFEQNDENMQKNNIFAIPSLQKHSNEIEKRIRSSIVPGRRMITHVAACYEAEMMENMSYLMEKTIANDSSLDETQDLEERGWKRQGLSDIEIFKRLENRKLKKLQNSGKAGLSCSTPKNGHNVSMGSLRGLNISHVCSIDRLDVNKVTIHNKNMLIPIIENDDYSCQKTHIIQKDHSSDLQGTFIVARDEEADNQKTNDVRKDTEESSLQDTFVVAEDKNDDKRNSSLQRKSKKVNLQERERRYANISMSLSNITIGDMSSPGSYRFKQIPKKSKPTIKTPPRTKMMNRLSIESDKEKTIEMLSIVEEASMEADSNSPSFVDPLSVNGSCVSPVPEFDELMDVSKAVTPK

>C_inopinata_HCP4

MHQRRNGRSTIVPGRRMITDQMAYHEAGLDVSNLSYLAEKTYMNESSVANDTVDQEEREWKKQGLSEKEIFNILERRKLKKLRDARNRLEKEMFGAKNFQELLERPEYSEIKNTDEDLNENAQKEPIFAVPAPPKHVSEKSMMSSPAAGSKGSGKAGLSCSTPKNSQDLSMRSLKLLDISHVVNIDRLDAKNVTVHNTKIVIPISKDTEKESTVLEQSSIDLQSTFTVETHKDPNAGKEPSLQKTFIVENEEEENQAESSLERTYVVKEGWDNAQNFDSAFEKKGTVNSIICDPVCGENTIEGGEKKKKSNKIGVQKRENRNVDLTTSLISSMIEDIPSPGAHYFKTTRKKFRPDPKTQSGTNKNNRLSTESDKEKTIEMLSIAEEASMEAELTGPSSIDPISVNEFQVSPIPEEIIAATTLRTPNSNRRENLPNNGISDETMRYKRMTETTPTRFPFESEDAATTPKPNYLTPTFSSLVKNRKITIADTNELLQANRRQGYLRNTPSQNYNARTKMTVSDTPFEVNTDPIEPQNDNQVVSSGEKDETTEVVNSSLDTALNALSVRERPDQPDIEDVDSDRYCQHDDDEDGFDNIGDLDYEEKVEEIPVSPKPRSIRRRKELLSDSVATVNTPGLDRRPTTQDTTYGTFVEDTWCPDIEKPKKTAGKNSTNEGMQLKKRELIMPGDAAGGVRRSSRVRVKPVRTWLGERAVYVNSPSGGKRLTGVTDVIIKDKRLCRFRTADLKLATEREQREIAYKKEIALRKRQQLAHDHRKGRRLNESQDDIHTDDEDDV

>C_elegans_HCP4

MNRKPRTRSTIVPGRKMITEIVALREAGLDDTRPSYLEEPTVIVDESMMNDSANLEEREWRKQGLSEKQMFAILEKRKQTQLNNARKRLEKEMYGAKTFKELIACREYSECESDSENTVNQNVRSASVFAIPALPKHISEKSMMGSPIANSRGSGKAGLSCSTPKSSSDTSMRSLRSLDISHVVNTDRLDAERVTVHTKSVVIPTILEERESTLQKTLTIEKNHSSQLQKTFTVAEDAPEELQKTVTIEKNQSSEMQRTFTVAKDASKEQSQLQRTVTIEKNQSSQLQGNFTVAKDAPEELQKTVTIEKNQSSEMQGTVTVAKDAEHSSLQKTFTKPTEKDESSLQRTFNVANRDENNDSTLQKTFIIEERDAYEQGTTSVGVKPPVLAQNTMEGGEKRRTSKVTSDERERRNASISNPLHNSMLEKEEIQSPGANFFKHPRKKIRPVVQTPPRIKATSRLSTESNKERTIEMESVAEERTMEADSIGSSYISPTFNASRVSPVPEVPEKVEPMHVSKAATPKSIRHTDNSIRNIGPTNNVECARAALLSTPTRMDIVDSVNRVTPARRYEQPTFASLVKRMNMKDANRLLEETSRKNTPAKTTATTSSAAVRMVLEDDEEDQATEVIEKRSENGGVIVDGEDEAADSSNRSLNIELNALTVNEEPAHDISAIDFPEDDTNEMRSSSDEEEMEARPDPRKKSSRRLGLLSDSIALGLASSSRRRPNDTFVDETWYPEPNSKGNRRPRTHRGMKLKEHQLMKPEDAPDGVRRSTRVRVKPVRSWLGEQPVYVNSPISGCKRLTGVTAVVIKDPRLCYYRTADVRTATERELKDKANKRALAQEKKQQRQNARSGRRHDSDDDEEDDM

>C_oiwi_HCP4

MHSASKIRRARSTIVPGRRMITAEVALHDCGMDVDQSYLNEKTIQSDESTIMNDTRDEEEREMRRRGLSEQEILKIQENRKREQRKNAERRLHEHKFGAKNFKELLGKHELSACESDGEEENRTVVIHRNQGEGGSKEIFAVPALPKHILGKSMLGSPVGARGSGKAGLSCSTPKGSKDVSMSSLRALDISHVVNTESLDSIKVTVQNKNIVIPAIPEESSLQKTHTIHRSDLEGTFIIKERSLDKTFTVESEKEMEQISEKNQENPSATEIVPAQEESSNSTLQKTFTAEKEWNDSLLDIVRKKQAPGPVEEEVPVVEASIQGGSKKSKKVNLQEREKRMADLTSSLLTSMNDDIPSPGAHCFRNTRKKIRPSLQTPPRAKVTSRLSTESDKEKTIEMLSIVEDGSMGGACSNGPSFVDAVSNNATQVSPIQEVPEPQTPKSSSHPLHIANDTLRLESSRARQHLDASSAITPRRNYERPTISSTIKSKNTAECESEVIVAIRSTERRHRRLPHDDDNGTLQAPEKPSEHVQLTEPEPEEEEMEQQSDVVETINQDSGESPRNQRPSVAAVEAKMDALTFEVTSPNMHHVMEFDYDNDQEEDDEVPEEPANDTTPRLKLSRRRTQLISNSMTSDESGQGPSRRRGFTDETVAEETWHPNLDEPSRRNRGGRKGKNDGLQLKKRELIVPDDEAPDGVRRSTRTRVKPVRSWLGEQPVYVHSPSGGKRLKGVTDVVIKDKRLCKYRTADLKLATEREQKERAMKKERAAQRRAQLAKAHKRGQRMNESQDDIHTDDEESD

>C_kamaaina_HCP4_paralog1

MHSASKIRRARSTIVPGRRMITSEVALHDCGMEVDQSYLNEKTIQSDDSSIMNSTRDEEEREMRRRGVSEQEIFKFQENRRREQLKNSERRLQEQKFGAKNFEELLARHEFSACESDGEEENRTIVIERNEEASKATFAVPALPKHILGKSMMSSPVGAKGSGKAGLSCSTPKGSKDVSMSSLRALDISHVVNTESLDSAKVTVQNKNIIIPVIPEESSLQKTHTIQRSDLEGTFIIKERSLERTFTVESEKEKEQSVEENQEGPSAEEDSESSLQKTFTAEHQLSDNSLLNIVRNKQAAAPVPVEAEEVPVAEASIQGGSKKPKKINLQDREKRIADLTSSLLTSMDDEIPSPGAHCFKNTRKKIRPSLQTPPRAKAISRLSTESDKEKTIDMLSIVEDGSMGGACSSGPSFVDAVSVNATQVSPIQEVPEPQTPKSSHHPLPSGNNTPCVESSRAKQHLDASAAITPRRNYERPTIASSMKNKNAADNEALLASLSKDRRPRRLPLDEDDVLEKSSVNVEMTEPEPEEVVETSHQDSGESPRHQRPSVAAVEVEMSALSFRGPSPNTNHVMDYDYHNDNGEEDADESEEPANANDTTPRLKLTRRQTQLMSNSMASEGPSRRRGFTDETVAEETWNPSLDEEPSRRTRGGRKGKNDGLQLKKRELIVPDDDAPAGVRRSTRTRVKPVRSWLGEQPVYVHSPNGGKRLKGVTEVVIKDKRLCKYRTADLKLATEREQKERAMKKERAAQKRADLAKYHKRGERMNESQDDIHTDDED

>C_kamaaina_HCP4_paralog2

MHSASEIRRARSTIVPGRRMITSEVALHDCGIEVDQSYLNEKTIQSDDSSAMNSTRDEEVREMRRRGVSEQESSTPKGSKDVSKRSLLALDISNVVNAESFESAKVKTFTAGLQLSDNTLRNIVRNKQAAAPVEVEEEEFPVAEASIRGGSKKPKKINLQIREKRIADLTSSHLTSMDDDILSPGAYCFQNTRKKIRPSLQTPPRAKAISRLSTESDKEKTIDMLSHAEDGSMRGACSSGPSFADAVSVNAKQVSPIQEVPEPQTPKSPHHHLPSGNHTPCVESLRAKQHLDVSSPITSRRNYKKPLIASLKIRVNTKKPEEIVETSNQDSEESPRLQRPSVAAVEVEMSALSFRFPSPTRNHVMHFDYENDNEEEDADESEEPANANDTIPRLRLTRGQIQQMSNSMASEAPSRRHEFTDDTVAEDEEPP

>C_waitukubuli_HCP4

MTRGSRKSNKIVPGRHMLSDQMQLHYLGMADTNHSYLAEKTADESSLMDCSREREERMWRKQGLTEKQIFRIQEDRVLKKHADAEARLEREKYGAKTFEEMLALSQYAIGALDVGEDDEPDVSKRENVFAIPALPKHVKPKSTMMSPAVRTTGKPGFGCSTPKGGKDLSMQALKTLDISHVINTDDLQSKKVNVQEKNVRISLKRTSDHANEEVSLEQTHTIRRDEDSELQGTFTVQKEGEGSLQGTFTVANDAEAEQSLQGTFVKEPWASMLCGTEDERDNGQSSGLQGTFVKESAGIEDQSQNNGAAENRNESESLQGTFVKAVDAPADAQQTKLNNTLDLMMLISNESNAAAQMTVSEGTVIVENTMEGGKSRRKSKKTDLNEREKRMADMTSSLINSMIDDAPSPGAALFKNTRKKIRPTVATPPRTRISRVSIESDKEKTIEMLSIAEEPTLEATGTSFVDPISVNGSRISPIPEHKESHPPVITATPKSSRRTGSLREQSVEKSRATGPSMRLSVASPFVGDSVIDTPVRTTPSRNYEKPTFASLAKSKNSKVVNLCETEKGHPEKQMSQKTLVRTPVLASKDFKDPEDDQLLGHVSVNTSDIPPRSEVQQVQDDDDPTVFPVTSNISIRNIEGDLNALSVNERNTVPVDYDMDFDDDDRQDPEMDRDEEEVEAEEEQVPARIGRQKPRSRRLGLLSDSIASVNTPEIAENRRPTRTHRPTDDTVVDDGWNSDEEYEDVPRRSRRNDQVGLQLAKRKLIKPADNDDGLRRSTRTRVKPVRSWLGEQAVYANSPSGGRRMTGVTDVYIKDKRLCKYRTADLGVATEREQRARAMKRARAEQKRRQLAHDQSMGRRLNESQDEIVTSEDEQ

>C_panamensis_HCP4

MEKKRSSIVPGRRMLSEQMALHQFGMADDNKSYLMEKTCDESTFMDCSQDREVRKWRKDGFSETQIFRIQEERTLKKHTEAEARLEREKCGAKTFEELLALQEHARRNQTMDTDEELDADPQGSVFAVPALPKHIKAKSTEMSPAGKAKGKPGFGCSTPVKGKNASMRSLETLDISLVVKPDALHPKNVNIQEKNIRVSLKRSTPHFHRIDDESERVEKDDTLQGTFTVAKDSASDSSLQKTHTIHKPDEDPERTFTVQKDDSLQGTFTVAKDSVNDSSLQKTHTIHKENDEPGGTFTVAKDASLQGTFTVAESSSQSLIKTFVRPPWDDEGLSAPSSQPAVVEDSAEDRSASLQGAFVKETDEAEATGKADQLQNTLDVFESNERTIVIEDVHLAQNTVEGGPSRRKSNKMDLEEREKRTADMTSSLLNSMVDDAPSPGQMFFKNANRKKIRPTTATPPRNKYVGRLSTDSDKEKTIEMLSIAEDPSMESNGSSFVEPVSVNGSRFSPVPEEHDGGQSMDLSVTPKSTRRSTVPLRQQSVEKDRAIGVSFQVSMCQSAVGSPCATPLRNYEKPTFSSLGKNKNATMFETEKAKQLEKMNKKTSIDHTPTRASTALTLAGSNEEEDVVVAPVSIPNPNRSPSSPLAEEHDEDTPTVFPVASNTSIGNVEADMNALAMNDNNDYGVNYDMDFDDDYRQDHRDVEEEDEQSPAPVQRPKPRSRRGVALLSDSMVSVNTPGIAGRRTTRRAMNDSAVDYNSNSDEEEDEGTSRNSRRNGNKKGLQLAKRKIIKPQANDDGLRRSTRTRVKPVRSWLGEQAVYVNSPSGGKRLKGVTDVFITDKRLCKYRTADLGIATEREQRERARKRARAEERRRQLAHDQSMGRRLNESQEDINTSDEEQ

>C_nouraguensis_HCP4

MERNRRTNNRIVPGRRMLTTEMAMYYAGMPDENRSYLAERTADESSLMDCSQDREERLLRKQGYSEKEILKMQEDRVQKKHADAEARLEREKYGAKNFEEMLNLPQYSMREPADDDDDDAGPQENIFAVPALPKHVNETSVVMSPAGQTIVKLGFGCSTPKGGKDMSMHSLKTLDLSHVVNTDKLHSQQINIQEKKVVISMKRTSNDDTESSLEKTHTIRNTFIVQNDDTLQHTFVIVKDPEEGLQATFTKEVEKNALSDSSQSTCLQGTFVKENDGQDQESVKEAGENPAEDQGSPTSLQNTFIKTVDGNENNNQNTLQLFEFTAHKQNSLMEETVSEQTELVHNTTEGGKNRRKSKKMDLGEREKRNADITSSLITRMIDDAPSPGARLFQNNRKKMRPTMPTPPRAKFGSRLSTDSDKEKTIDMLSIAEEPTMESNGSSFVDPMSVNGSRVSPIPEKDETQSMGISATPKSIRRTTGIRPISVEKSRASVLTPRISMATCLSDSVHIPAVTPTRNYEKPTLSSQLKNKNATEASVWEAEKEKEIGKMNRNTPKRTHATVAPVEASSDEEEEQVVVTPVNKNRSPVFERLDESPDEPTVYPGSSDTSIGNVESDLNALSVNDRFHAPAPEFDLGPDYDRSNQEEQSNGEEEEVEHREPSPAQRANPTSSRRGIALLSESMVSVNTPGIAGRRPTRNRRQVDDTVVDDGWNSEDDDSNRRGRRRNGKQEAGLQLAKRRIIEPKDAGDGLRRSTRTRVKPLRSWLGEQAVYVNSPSGGKRLKGVTDVIIKDKRLCKYRTADLRVATEREQKEKARKRARAEQRRRQLALDQSMGRRLNESQDDILTSDDEL

>C_becei_HCP4

MERNRRTKNSIVPGRRMLTTQVAMYNAGIHDENRSYLAEKTADESSLMDCSEDREERMLRKQGYSEREILKMQEERVQKKHADAEARLEREKYGANNFEEMLNLPQYSMREPADDDDDDVGPQENVFAVPALPKRVNETSIVLSPAGRVVGKPGFGCSTPKGGKDVSMRSLRTLDISHVVNTSKLHSERINIQGKNVVISMKRTSDADEESSLEKTHTIRNDGTFTVQKDDSVQRTLIIIKDPEEDLQATFTKETAQDAQSNSSRSSSLQGTFVKRTDGQDQESVKDTEENPAENRDSSASLQGTFIKPVDSDENKNQNTLQVFESSAQNQSSLMEETVTDQMEVVHNTTEGGKNRRKSKKMDLGEREKRNADITSSLISSMIDDAPSPGARLFQNNRKKMRPTMPTPPRAKFGSRLSTDSDKEKTIDMLSIAEDATLESNGSSFVDPISVNGSRVSPIPEKDETQPMEISATPKSSRRTTGIRPASVEKSRGPLLSPRISMAPCLSDSVTVAAVTPTRNYEKPTASSLAKNKNPAEAIRWEALKEKELGKINRQTPKKTPVAMPLAESSSDEEEEQAAVTPVNVNRSPAPDRLDESMDEPTVYADPVNSSIGNVESDLNALSVNDRFDAPAPDFDLGPDYDREPQSDGEDEVEDRMPSPVQRARPKSSRRGIALLSESMVTVNTPGIADRRPTRNRRQVDDTVVDDGWNSDDDDSNRRGRRRNGKQETGLQLAKRKIIEPEDAGDGLRRSTRTRVKPLRSWLGEQAVYVNSPSGGKRLKGVTDVIIKDKRLCKYRTADLRVATEREQKEKARKKARAEQRRRQLVLDQSMGRRLNESQDDIHTSDDEL

>C_yunquensis_HCP4

MDRNRRTNNKIVPGRNMLSAEMQLYYMGVEDDNRSYLAERTVDESTLMNCSQDQEERMWRKQGLSEKEIFRIQEERALKRHADAEARLEREKYGAVTFEDMLALPQYSVREPADDDDNDDVSPQENVFAVPALPKHANEKSIVMSPAARGIGKPGFSCSTPKGGIDLSMRSLRTLDISHVVNTDKLQSRKVNVQERNVVISMKRTSDRVDGTTSLQRTHTIRKDDEADLQGTFTVQKSPDRTFTVVREAEESLQGTFVKETDNNAQSDNSRSSSLQGTFVKQNEKSSSLQGTFVKETDNNAQSDNSRSSSLQGTFVKQNEKSSSLQGTFVKQTEESTSLQGTFVKEPSSDQSKEPNLQSAFVQETVGAPTEKRDSCAVLQGTFVKHANDSGNDNQNTLQVLQFQTHDESSVLVERVTEEVRVVHNTIEGGQNRRQSKKVDLDRREKRDADMTSSLITSMIDDAPSPGAQLFKNNRKKMRPTMPTPPRAKFGSRLSTDSDKEKTIEMLSIAEEPTLESNGSTFVDPISVNGSRVSPIPEKDETQMMDIASTPKSSRRTGSIRPSSVEKSRNNALSMRLSVAPSLTDSLIVTAITPARNYEKPTFSSLVKNKNSVEARIFLETQKDLGKMSKQTPKRSVAVHPVEPSSDEEKDQAVQAPNNRSISPERRDESMQDEPTVYPGSSNTSIGNVEADLNALSVHDRYDAPLDLDLDDFRPNDREAQSKGEDEVEERQAFPVRQAEPKSRRRVGLLSNSMITVNTPGIADCRQNRNRRQVDDTVVDDGWNSDEDDSNRRNRRRNRKQEVEMQLAKRKIIVPEDAGDGLRRSARTRVKPLRSWLGEQAVYVNSPSGGKRLKGVTDVVIKDKRLCKYRTADLALATAREQKERARKRARAEQKRQQLAHDQSMGRRLNESQDEIVTSDDEH

>C_macrosperma_HCP4

MKPDSRTSNKIVPGRKNITDLHGLHSAGMNDENRSYLAEKTADESAFMDCSQDHEERIWRKQGLSEKEIFRIQEQRALKKHADAEKRLENEKFGARDFKEMCALPQYSLREDNEEEDDDRFCSRENVFAVPALPKHVVEKSIVLSPAGRSLGKPGFSCSTPKGGKDLSMRSLKTLDISHVVNSEKLQVKKVNVQDKNVTITIKRTTEQAEKKKSIEKTHTVPEDETSELQGTFTVPKDGAESLQGTFVVIPGSGESLQGTFVKETGNCSGEQSASLQGTFVKESDGNGSIGQSQSTSLQRTFVKESDDAPNDDQEQSASLQGTFVKDVDASENDRTDSTLRVLQMNSHVQNELIDGTLTEELPTAQNTMEGGEKRRKSKKVDLGEREKRMANVSSFINSMIDDAPSPGAHFFKNSRKKMRPTVATPPRTKFGSRLSTESDKEKTIEMLSIAEEATMESHGSSFADPISVNGSRVSPIPEKDETQPMEICTTPKSSRRNNYSLRQSGTCEKSRASEVSMRLSVAPAASETLVSGIVTPSRNYQKPTASSLAKSKNAAVFSQSEREKANERSKTNKQTPRISASHTPVFKEDSHRETVVTPVCKDRSISPQGFEEDMQDEPTVYPGPSNDSIGNVEADLHALSVNDRYEAPFDLDFDADYGHDNAPESEEEGEEEQEEEEERRDLSPAQRPKPKSSRRRVALLSDSVASVNTPGLDRRSKRNTRVTNDTVVDDDWDSEDEEDQMRSSRRRNGKNEVGLQLAKRKIIEPNSTADGLRRSTRTRVKPLRSWLGEQAVYVNSPSGGKRLKGVTDVIIKDKRLCKYRTADLKLATEREQKEKARKKARAAQKREQLAADQSMGRRLDESQDEIVTSDDE

>C_sulstoni_HCP4

MENFRRPCVSRTIVPGRRIIPEVVYRDAGIAEDMSYLAEKTIDESIMNCTTDMEEKMWRKQGLSEKEIFRRQEERTLKKHAQAKARLEEGKFGAKTFAEMCALSEYSQRESEDEENEVPVQQKKKEDVFAVPAIPKHVNEKSILMSPSANTKGSGQPGLSCSTPKGNKDLSMRSLRTLDISHVVSTDRLDFRKITVQEKTVVIPVHLEEDSSSSLQKTHTIRREESSTQQGTFTVEKSGEKESLQETFTKEKDHSGSLLGTFVKESNQPESLQGTFVVESGKEQMESLQGTFVKVIEEVPQEVPQAVKEKTLVQVSEEVSTINETMLDGERRKSKKMGLKERERRVADLTSSMMSSMIEDIPSPGAHFFKNNRKKMRPTVPTPPRSKVGTSRLSTESDKEKTIDMLSIAEELTIESSGSSFVDPISVNGSRVSPIPEQNEAMEISVTPKSSRRPISSLRQDPASIEKSRRQGSTLRISMAPLALSIHSPMLEQTPRNNYLKPTFSSIAKHKTPAENKELLEAEKLNRRRTPCATPTKENGFVDFEKQEEQEETEECSQRSAIVEGLEILEDPTLAPSNRSIENMEDQLDALSVTDHRQFDIPEVDYGEDYPQYDDAPEPLMSEGEEEEPTQRPKKKEPKRRVGLLSDSMMTVNTPGLEDRRTTRSRRGGRGASNAEGPSYDDSYEQESRGYGRRKKQEVGLQLAKREIIEPEQQTDGLRRSTRTRVKKLRTWMGEKAIYVNSPSGGKRLVGVTDVFVKDKRLCKYRTADLNLAMERENRERAQKRARAAERRRQLALDHSFGRRLNESQEDIVTSDDES

>C_afra_HCP4_paralog1

MNDLRNTRFQSTIVPGRRIIPEVIYRDAGIEGEPSYLAERTMDESVVNCTVDMEERMWRKQGVPEAEIFRRQEERTLQKHAQARARLQKEKFGAKTFGEMCALSEYSARESEDEDDGARAQERKKKKENVFAIPTIPRHLNEKSILGPPSVHAKGSGKAGLSCSTPKGNKDLSMQSLRILDISHVVNTDRLDNRKTTSQEKTVVIPVCVDDESSLQKTHTIQKEESQGGATFTVAKNGEDGSLQGTFTLEKNGEEGSLQGTFTKGQNQSASLQGTFIKETDPTENQQGTFIVQSAKDQTESMHGTSVKTTEEDPEVVTQLAEEKTLVQISDDVPTANETIQGGERRKSKKIGLGEREKRVADLTSSLMSSMIEDIPSPGAHFFKNNRKKMRATVATPPRSKLGANRLSTESDKEKTIDMLSIAEELTIEANGSSYVEPVSVNGSRVSPIPEDNEAVPMEISVTPKSSRRPVSSLRQEAASSIEKGRRQASLMRISMNPMTSVHSPIAEQTPSKNYLKPTFSSIAKHKTPSESNELLEAEKMNRRRTPVSTPTKNRMFEDIDVPEEFKAEEGIQRPAVIEYPEGPEDPTLQTSNRSLGNVADDMNALSVTDNRPSASSDVQDHHHSCNPSDDYYPDDEAAAPMFEDEEEEEEDRTRKPKKKQTRRRVALLSDSIATVNTPGLEDRRAARSRRYGGAQSSSVANSPSFDYSYEQEPRYGRKKKQVEMMLAKRELIAPETPTDGIRRSKRTRVKPLRTWLGETAEYQNSPSGGKRLVGVKEVIIKDKRLCKYRTADLNLAMQRESREKALKRARAAERRRQLAVEQSFGRRMNESQEDIVTSDDE

>C_afra_HCP4_paralog2_HCP2L8

MNNSRNTRLKSDIVPGRRIIPDVIYRDAGIEEEPSYLAERTMDESVVNSTRNTRWNSNIVPGRRIIPDVIYRDSGIRGEPSYLAERTMDESVKSKRIGLGEREKRAADPTRSTMSSMIEDISSPGVQFFENNREKMRPTVATPRRSNLGANRLSTASDKEKTIDMLSIGGEPTIEANGSSYVEPVSVNGSRVSPTREDMLHRHVGPVITEVEESPSRDPSVLRDSGRRHNVARSKSTAIERQIKLIFIEPRFKERPEAMAEVIRLMRKQIDQWERDQDSYGPTEERTKNIRAWKDNRRRFIEQLEDAKERQERARRERDESSRVVETVRRRGDMTGMNVTAIHNSRSYDQRTHFDSEEEEKENGYTSRAPRPLPRSPPRAHQSYHSSLSHQSRRLDASSRDNRSRNDYGSNNVTSSSHQGKPKPRLRAGKSRVTKSMFRPRRDRALLEIRQYQKSTDLLIQKAPFCRLVQEILREEATTSSSDYRIRADALTALQEAAEAFLVEMFEGSQLIATHARRVTLMHSDIQLYRRLCLRK

>C_sp49_HCP4

MSNGESSNRIKPGRRMISQEAALFAAGYHEEEASYRAEQTIMSANSTINDTMDLQERAWKKQGVPDDEIFRRQEAKTLQKHTDHRTALDAGMDGANDFKELLMRPQYAVRRHFHDEEDEENHHPPKGQKNNVFAIPALPKHMGTPKVNGKAGFGCSTPIGGDVSMRSLRNIDISHVINVDQLDANKTTTHSKNVVIDMVATSANSTLQKTHTIRKDSEEESLQKTFVKQSDQSLQGTFVVAPATSDSTLQKTHTIRRDSETSTALQGTHVVEKDSTALQGIFVVEDKEPAAIQDTFVVEKEPQKEPGYSESSSALQETFVKHSGQLEQFEQPNDATISHETTQGGRRVKKVDLKERENRMADLTTSIMGSMMDTPSPGAHHFKNPRKKLRETVATPPRLTTARMSIESDKERSIHMESINNDSTRGRESVGSSYVDPVSVNASRMEIVVEEEEREEEKTHVPMDIATTPKSTRNTPRPWSRATAGESARAGRPSNYADPISQEDTTNLVTPVRNFQKPTISSLSKNRRASARMILETMSANRSTMKQSDDNTETSRKTSDAPVARKVQESPETADKVPDRQVSQNDSLGDVANHFRHISFTDPPPAPQEDVDFGDYGQEESDPEEGTSGLPPAFQRDSQKDAESHRTKPSRKGRLGLLSDSLTSTNTPGVDRRKNQVNDTAVDDWDSEEEEEEPTRRGKKAKQVGLQLAKRRIIEPEAAPDGLRRSGRTRVKPVRNWLGEQPIYEHSPSGGRRLKGVTTVFVSDKRMCKYRTADVTLATEREVESKRKRAKRRAEERRRQLELDQSRGHRMDESQDDIHTSDED

>C_sp25_HCP4

MSRSLSNGIRPGRRMISEEAALFAAGYREEEASYRAEQTIVSANSTINDTIDMQERAWRKQGISDEEIFRRLEARAVEKHTEHRQALRAAMDGAKNFEELLNRPQYAVRRHAEENEENQHSSGVQFAIPALPKHLGATPKAKGQPGFGCSTPIGSDVSMRSLRNIDISHVIHIEQLDPNAPTTHTKNVVIEVESANSTLPKTHTIRKDGESLQGTFTVAQPAASTSESSLQKTFVKESQEPLQGTFTVPPAAPSESTLQTTHTIQKDADETAIQGTFVVEKEKDKDDEQAAGDSEQSLQATFVVEEPPRNHETSQGGNGARRARKVDLKEREGRMADLTSSMMGSLMDTPSPGAHHFKNPRKKLRETAAPRLAATRLSTESSDQERSIQMLSINEDSTRGRESIGSSYVDPVSVNASQQMETVVEEEEAAIATRNTPRPLSPLTAAGESVRAAPRSSIIATTTTADTTGLLTPTRNFQRPTISSLSKSRNANAKEILETMSQKKSTAERKEAVEEEKEKEQVVPEVVVNSFRQISFHDPSPSPPPRSSPPRSPSPVAALEDVDFGDDAADQSDPEEGTSGPPPAFLMLHSHSHRGEAKCQRKPRIGLLSDSMTSINTPGIDKRRGRVNETVPDEWNSDEEEQEQEEMPRRRKPKQIGLQLAKRRIIEPEQAPDGVRRSTRTRVKPVRNWLGEKPIYVHSPSGGRRLKGVTDVFVTDKRMCKYRTADVRLCTEREVASRRKKAKRRAEEKRRQLEMDQSRGRRMDESQDDIHTSDEE

>C_imperialis_HCP4

MRDSDAASGIKPGRRMITQEAALYAAGYDEEASYRAEQTIVSANSTINDTMMMQEREWRKQGISDAEIFKRQEEMARKKLSDHRAALTAAMDGATNFAELLSRPQYAVRRIPDEENKFAIPALPKHLGTPKINGKAGFGCSTPIGSKDVSMRALRAIDLSHVIAVEQLDANKTTTHTMNVVIDMAGTSSAESSLQKTHTIQKEAEDSLQKTHTIQKDSENSSALQGTFVVSPASDSSIQKTSEASTAIQGTFVKDSEDVTLQGTFVVEKEAVPAPEDVTIANNTTLGGRRGRKVNLKEREGRMADLTTSMMGSMAETPSPGANHFKNPRKKLRETVAAMRMSIESDKERSIQMLSINEESTRGRESVGSSYVDPVSVNASQMETVVEEREQEVEEKEQEKEHGEAPMEITVTPKSSRNTPRPSIFSPGEAARGARPSNFAAIAPADLTLLTPTHHFQKPTVSSLSKNRNAGAKEALEALTQKKLAKKVPEPAPESPTSPKSVEQPSRLHNNSFGDVTNGLRQISMHDVPPIEDVDLGDHLERGEGTSGPPPEFQMTSPEIRSTRSSKSKKVGLLSDSMTSLNTPGVDRRRGPETTVVEEWNSDEEPEDVPRRRNKKQVGLQLAKRRLIEPEAAHDGLRRSTRTRTKPVRNWLGEQPIYEHSPSGGRRLKGVTTVFVTDKRMCRYRTADVKLATEREHEAMRKRAKRRAEERRRQLELDQSRGRRMDESQDDIHTSDEE

>C_japonica_HCP4

HQSRLVKGYRGAKNYHELLAAYNCLPDQPGPPDGDVEAEENVSNAQKKDDSVFAIPSMPKKKNAGPSCSTPISGKDASMRTLKGLDISQVFSNVESSFTTVKCAGGGGEPSLQKTFTMRNVEEEEQESIQKTFTVRNGEEESLQKTFVVAEADGGASSSSSSSLQKTFTVRNEEEPLQGTFVVEKEKEAEPETEHKPIPQPQEPPKTHNRPKKTNLEKREGRMADLTSSMMMGIDDTPSPGARHFKPNARKKLRESLQTPPRGLSNRMSLDSDKDKTISMLSVAESSGRESMGSSYADPVSIHHQTSHISSIPEEHEEEQEEVEVEEEAETTVQNSEASVAIKTPPRAPRLTQSSIERRRANQSALRISTAPKAAEGTFLTTPTAHNYQKPTFSSLVKSKDRAECTELLTELNKNRTPRRTVPPVEEPAQELGHVAEDLEALSISVGVLEKEQQEPRPLTEDVDFEDREEPVPDDESVKKMLRRVGLLSDSIATVNTPGHPRAAVHFNDTTAVDQWRSDEDDDEEEEEEGPSRRRQTRRGGKKQEVGLQLAKRRIIEPEQAPDGIRRSSRVRVKPLRSWLGERLDYAFSPNGTRRLKGVNDVFIKDKRMCKYRTADCRLAMEREQREKARKRERAAKKRARLALDQSQGRRMDESQEDIVTSSDEE

**Supplementary Data S4: All KNL-2 sequences used in this study**

>C_tribulationis_KNL2

MGDGDVVPARVQNVLDMDVVRLNLWSLQFNSSGVRLEGFVRSEDGNMMQKVHSGIICRRMSATMLFDVSGCFYELAGHIDREFQQKLGMPSRVIEEFMNGFPDNWAFLIKSCLPAEPKSALRPIQAAPEPSKARGEPIVTMPDETIVDNQSAEKERKRKELQKKKEQEERVRIQREQRAAAEAAAERKRQEDEEFKRREEEEERKRQEEENDAANYTFRVPQSQNGDAITPIRFTRGNGQKGANVRSIFEKTPRTKPAGPLASSTPQAPPQQHAPRISIAEAKQPDEPPHPHPKPQPPTRESIRETQYCSDSEFAVPKLPAPRNNRHASASSQPAPLDFLDEMDALFETANVDHTPGRGRKSRKPVSRSPSPGRFNSTARDRTGYDRYEPSRQSSSQRYYDDYNNVSRMSGRNHTLGGHDMRRDESRNSRKRGYNNSPEDYSRRFDDRSRRRDNYESDSRYDSKRSRPRDQSSSSGRSVRFEEDFPRNRRDESRDSRSYRHYEDHRNRESSGDREDKRKLDAIVRREKELVARLQNTQRSSSTLQRSGYSSEDDEMSDEWDRENQEMLDNSMMFGDGLSKKKGKSGGHKTMRQAKTRYPPKPKPAQKPAQPKKKKKDVEYESDEMNDSIASNRPRRACATPSTPAPKRITWPKRDLDRLKHTIELKKPTGAEADWAEVTRLLAKDGVDAEVVKQTAIAKLKWKEPSQKTIQQEEEEKKRRRGATARVKEGVRMHEELREGGNNRAESSQSGVEAVDDYEPDDVAADQSLLGLQTPIAVKKRGGTRASIMPQPVEDSPVVRGNNSTLNSPRLDQTKAKDVETTLKYVQHLSMMNARPSSRANSSYLNKSSSRGGDKKNTSLSVEQGARKALKIINRGRTIHEEDEDDEDEDITID

>C_sp41_KNL2

MGDGDIVPARVQNVLDMDVVRMNLWSIRFNGSGVTLEGFVRSEDGNMMQKVHSGIICRRMSATMLFDVSGCFYELAGHIDREFQLKIGMPSRVIDEFMNGFPDNWAFLIKSCSTSEPKSALRPIQAAPVEPLKSRSEPIVTMPDETVLAESQTAEKERRRKELQKKKENEERLRIQREQQEAVKAAEKKSQEEEKDKLQEEERKRQEEEDAANYTFRAPKSQSGEAITPIRFTRGNGQKGANVRPIFDKTPVRTKPAGPLASSTPQAPPPQHPHRLPNIETTRPDASVPPVPQPPIRETQYCSDSEFFAVPKLPAPKNIRGTSAKPLGFLEEMDALFENVNVDQTPGRVRNPRKVSRSPSPMRLNSSARNRDSGYDRYEPSRQSLSQRYYDDYTTSRMSGRDDTFGRNDTRRDESRNSRKRGYNNSPDEYRNRWDDRSRRHDNFESDPRYDSKRSRPRDQSSSSGRSVRFEEDHPRSRMDSRESMDSRNYRHYEDSRNRQSSGDREDKRKLNDILRREKELMARLQNSQRSSSTFQRTAYSSEDDEMSDEWDRENQEMLDNSMMFGDGLSQKKRRSGGRQPAKKPNYLQQAKTKRVPQPKPAPKPAQQKKKKKQDSEDERDEMNDSIASNRPRRACVTPSTPAPKRITWPKRDLDRLKHTIELKKPTGADADWAEVTRLLAKDGVDAEVVKQIAITKLKWKEPTQQTIQMEEEEQKRRRGATARVKEGVRMHEELRAGGNDRGNSSQGGVGAVEDYEPDDVAADQSLLGLQTPIAIKKRGGTRASIMPEPVEDSPMVRGNNSTLNSPRLDQTKAKEVETTLKYVQHLSMMNARPNSRANTSYFNKSSSRDKTNTSLSVEQGARKALKIINRGRTIEEEDEDDEDGDTTID

>C_zanzibari_KNL2

MGDGDVVPARVQNVLDMDIIRLNLWSIQFNGTGVKLEGFVRSEDGNMMQKVHSGIICRRMSATMLFDVSGCFYELAGHIDREYQLKIGMPSRVIDEFMNGFPDNWAFLINSCSEPKSAMRPIQAAPREPLRTRGEPIVTMPDETNIESQSSEKERKRKEREEQQKWEHEERLRARKEKEQREAAEIAEKNRREREIQKQREDEEKGRREEEEEAANYTFRAPKSQNGEAITPIRFTRGNGQKGANIRPIFDSTPVRTKPAGPLASSTPQAPPPQNRHSNTENRQPDALPTAGPAPPVQKPNRETQYCSDADLFAVPRLPAPRNNQNVSSSAQSAPLDFLDEMDALFESATVDKTPGRVKKVVRVNRSPSPDRFSSRDRDSGYDRYDSSRYSHSQRYNDEHHMSRMSGRNDTFRRNDGWRDESRNSRKRGYNNSPEYTRGWDDRSRHRDNYESESRYDSKRSRPRDQSSSSGRSVRFEDDHPGNHRDSRDPRNYRDYEDDRNRESSGDRADRRKLNDILRRERELEARLRNTQRSSTARRHAYTSEDDEMQDEWDRENQEMLDNSMLFGDGISQKKRRSGGRQPAKNSQSGRSQQPKPARKPAQPKKKKERSDETDELNDSIASNRPRRACVTPSTPAPKRITWPKRDLDRLKRTIDLKKPTGADADWDEVTRLLAKDGVDREIVKQTAIMKLKWKEPSQETVQQEEEEKTRRRGAAAMVKEGVRMHQELREGGDNKRGDDLQGGVEAVDDYQPDDVAADQSLLALQTPIAVKKRGGTRASIMPQPVEDSPVVKGNNSTLNSPRLNQTKVKEVETTLKYVQHISMMNARPSSRANTSCLNKSSSRGGGSKNTSLSVEQGARKALKIINRGRTIQEEDEDDDDESEAGDITIE

>C_sinica_KNL2

MGDGDVVPARVQNVLDMEIVRLNLWSIQFNGSGVKLEGFVRSEDGNMMQKVHSGTICRRMSATMLFDVSGCFYELAGHIDREYQLKIGMPSRVIDEFMNGFPDNWAFLIKSCSNPEPKSALRPIKAAPKMPLRSRGEPIVTMPDETVMAESQAAEKDRKRKEHEEQQKRDDEKRFRAEKEKEQREAADTAAKKRQEEEDRRRREEEDANYTLRAPESHNGEPVTPIRFIRGNGQKGANIRPIFEKTPVRTKPTGPLASSTPQAPPPPPTRRTSNIENKQPEASTSPKPQPPVQKPVRETQYCSDTEFFAVPKLPASRNKHVPASDHAAPLAFLDEMDALFNGADVDHTPGRERKPKKVSRSPSPERFDYSSRDRDSGYSRYDSSRYSHNQRYNDDYNMSRMSGRTDTSRRNDGRRDESRMSRKRGYNNSPDEYSRRYDDRSRRQDDYDSGSRYDSKRSRPREQSSSSGRSVRFEEDYSRHRQGSRDSRDPRDYRDYEDHRNRTSSGDREDTRKLNDILRRERELMARLQNSQRSSSTVQRTAYSSDEDEMADESSLWDRENQEMLDNSMMFGDGLSQKKRRGGGRHPAKQAKTRQPAQPKPARKPAQPKKKNRADETDDLNDSIASNRPRRACATPSTPAPKRITWPKRDLDRLRHTIELKKPTGADADWAEVTRLLAKDGVDSEIVKQTAITKLKWKEPSQETIQMEEEEKKRRKGAAARVKEGVRMHEELREGGDKRGDNSLGGVEAVEDYEPDDVAADQSLLALQTPIAVKKRGGTRASIMPQPVEDSPVVRGGNNSTFNSPRLDQTKAKEVETTLKYVQHLSMMNARPSSRANTSYHNKSSSRGGGSKNTSLSVEQGARKALKIINRGRTIHEEEEDDDEEGDTTIN

>C_nigoni_KNL2

MNDLFFVNYGSKFMNDYAQKKKASSHIEDYSLPEKKLRKVQEIVDKSPKCLFAKRAPPHTHFAANFSNLNSEGIKTVFLDIHLDFTISTPYFCYKASFCKKKMGDRNVVPARVQNVLDMDIVRLNLWSIQFVDSGVILEGFVRSEDGNMMQKVRSGMICKRMTATMLFDFSGCFFELSGQIDREYQQKMGMPSRVIDEFTSGFPENWVSLIKSFLPVNPISAVKHIQPAPREPLRPISEPIVTLPDETTLESEKDQKRREKEERLEKKRLEEERLEKERREQKLRAQKEKERREAVAAAEAEKKRLEEEEDAANYTIRVPKSQSGEAITPIRFTRGNGQKGANVRPIFDKTPVREKTAGPLASSTPQAPPPQQRLSNVEKKPASPEPQPRNPQRDREIQYASDSDLFAVPKLPAPKTTKPSTTSEASSGGVLDFLDEMDTLFDTVVVEQTPTRDRRPPRRYSRSPSPRRRQHSSSRDMGKGYDNFESSRYSQRYNDNYNTSRMSRRDDTFRRNDERRDESRMSRKRVYNNSPDDFEYNNRYDDRSRRPDYYDSDSRYDSKRSRPRETSSSSGRSVRFEDDYRRNHRDPRDRSDSKDYRNYDESRRNPEDRREEKRKLNDILRREEELVTRLQNRKKPSASYRREPSSDEDDTADEWDRENQEILDNSMMFGDGLSQKKRRSAGRPSKPSKKERQVQPKPVRKPPQPKKKQKSPPDELNDSIASNRPRRACVTPSTPAPKRIVWPKRDLDRLKHTIGLKKPTGSDADWAEVTRLLAKEGVDAEVVKQVAITRLKWKEPAQDPETIQREEEEEKKRRRGVAARVKEGVKLHEELRQPGVKRGDNSQTGVEAMEDYEPDDVAADQSLLALQTPVGAKRKGGTRASIMPEPVEDSPLVRRNNSTFNSPRLDQTKAKEVETTLKYVQHLSMMNARPSSRANTSYYNKSSSRGGGSKNTSMSLEQGTRKALKIINRGRTIHEEDEDEDDDDDDEDQYEDGVVY

>C_briggsae_KNL2

MGDRNVVPVRVQNVLDMDIVRLNLWSIQFVDSGVILEGFVRSEDGNMMQKVRSGMICRKRSETKKQEERLGKKRFEEERLEKEQREQKLRAEKEKERREAVAAAEAEKKRLEEEEDAANYTLRVPQSQSGEAITPIRFTRGNGQKGANVRPIFDKTPVRGKPVGPLASSTPQAPPPQQRLSNVEKKPASPEPRPRIPQRDRDVQYASDADLFAVPKLPAPKTTKPSTTSEPSTGGGFDFLDEMDELFDTVVVEQTPTRNRRPPGWYSRSPSPRRRQHSPPRDSFESSRYSQRYNDNYNTSRMSRRDDTFRRNDERRDEFRMNRKRVYVCDRGLVRKRKYVFQNNSPDDFEYSNRYDDRSRRPDYYDSDSLYDSKRSRPRETSSSSGRSVRFEDDYRRNHCDPRNRSDSRDYRSYDESRRKPEDHREDKRKLNDILRREQELVSRLQNRKKSSVSYRPEPSSDEDDMADEWDRENQEILDNSMMFGDGLSQKKRRSAGRPSKPSKKERQVQPKPVRKPPQPKKKKQKSPPDELNDSIASNRPRRACVTPSTPAPKRIVWPKRDLDRLKHTIGLKKPAGSDADWAEVTRLLAKEGVDAEVVKQVAITRLKWKEPAQDPETIQREEEEEKKRRRGVAARVKEGVKLHEELRQPGVKRADNSQTGVEAMEDYEPDDVAADQSLLALQTPVGAKRKGGTRASIMPEPVEDSPLVRRNNSTLNSPRLDQTKAKEVETTLKYVQHLSMMNAHPSSRANTSYHNKSSSRGGGSKNTSMSLDQGTRKALKIINRGRTIHEEDEDDEEDDEDQYEDAVVY

>C_remanei_KNL2

KRQEEEDADYTFRAPQSQNGEPITPIRFNRGNGQRQGVTRSVFERTPQRGQSGPLAASTPQAPPPPPPQQRLSNIENREPPPRVVAPPSPVRQPPAPQPPIREPQFANDDDLFAVPKIPPPKIPRGSAGNSGGNIDFLDEMDALFDTVYIDKTPKRDVKPKRPSSPSPERRRYSPMPRDREMGYNDDFESSRRGGRYPSESSNMSRMSRRDETNRRNDGGMGRDESRMSRKRGYDQSPDDMEYQRRREDHYRRPDYYPPRDSRQDSKRYRPRENSSSSGRSASVRFADDYQRNRGDSRDPRDFRDPRDSFSRDPRFYYENNQRGESSKDRDTRKLNNILRQERELVARLQNIKSNTTTNTNTTRRVTYSSEDDEMADEWERENQEIMDNSMMFGDGISKKGRRSGPGRPPQRKPKEQPKPKSQSQPKRRNQNQSKPRRSQYDPVETDDLNDSIASNRPRRACVTPSPVVPKKIIWPKKDLDRLKRTIELKKPTGSEADWTEIARLLAK

>C_latens_KNL2

MPSRVIDEFTNGFPENWGFLIKSCFGNESRSAMRPIQAAPREPLRQRNEPIVTLADETELTNRKNSDNDSESENNRKRREREEEAERERQERYRADEMERERIAAEKKRQEEEDADYTFRAPQSQNGEPITPIRFKRGNGQRQGVTRSVFEKTPQRGQSGPLAASTPQAPPPPPPQQRLSNIENREPPPRVAAPPSPIRQPPAPQPPIREPQFANDDDLFAVPKIPPPKIPRESVGNGGGSIDFLDEMDALFDTVYIDKTPKRDVKPKRPLSPSPERRRYSPMPRDRELGYNDEFESSRRGGRYESSNMSRMSRRDDTYRRNDGGMGRDESRMSRKRGHNYSPDDMEYERRREDHYHRPDYYPDSRYDSKRYRPRENSSSSGRSASVRFADDYQRKRGELRDPRDFEDSFSRDPRFYHESNRKRESSQDREDTRKLNDILRREKELMARLQNNKTTTSKNTTTRRVTYSSEEDEMADEWERENQEIMDNSMMFGDGISKKGRRSGPGRPPQRKPKEQSKPKSQSQPKPRNKNQSKPRRSQHDPVETDDLNDSIASNRPRRACVTPSPVVPKKIIWPKKDLDRLKRTIELTKPTGSEADWTEVARLLAKNGCDGEAVKQTAITKLKWKEPVERDEETIQREEEEEEETKRRRGVNAKVKEGVKLQEELRRGATHQKRAEDVRVEVTAEEIEPDDLAADQSLLAMATPVAVKKRGGTRASIMPMPVEDSPIVRGSRNNSTFNSPTLNQTKAKEVETTLRYVHQLSMVQARPASRNNTSYMNKSSTRGGGSKNTSMSLEQGVKKAMKIINRGTTIHEDDEEEEESESEEEDNSEDLEEH

>C_sp51_KNL2

MGDTDILPTRVQNVLEMEIVRLNLWSIKFNALNIKLEGYVRSEDGNMMQKVCSEIICKRMNSTMLFDVSGCFYELSGQIDREYQSRMGMPSRVIDEFVNGFPENWAFLIKSCLSMGQRSALRPIQDTPREPIRTRVEPIVTLADETELVENERKKAESEKDKKRKEREEDRARENERNLAAQREKERKDTVAAEERKRKKEEEEAHAVAERKRQEEEEAANYTLRAPQSQTGEAITPIRFTRGNGGNGAHVKSIFQKTPVRGKSNGPIASSTPQQTRVLPTLEKPVETTVSKSTKSKSPPPRSIRQTEYASDADLFAVPRNPAPKNNRKPAPTTSSDIGFDFLDQMDSLFDNPDFEQTPTRDRKPRRQFSPSPEPRHRSYSRDRDRYAHLESTRYKQRYTDDYDMSRISRRDVPFRRNDGGRDESRMGRKRGYYGSPDGSEYGRRYDDRDRHMDYYEPDFKYDMKRSRPRETSSSSGRSVRFENDYRRRREDSREPNSSRNHREYDDYRGRTDSEDNRRLEFLEKRENELMARLQNHKKPSSSSSRRLTNSSDDEMADMWEQENQEILDNSMMFGDGISKKKRRSGGGRQPKKQPTRKTQPKPTRKPAEPRKKTQRSPVQRDELNDSIASNRPRRACATPSTPAPKRITWPKRDLDKLKHVIQLKKPTGADADWIEVARLLAKDGVDSELAKQTAVSKLKWVEPVQHEERIDEEEEENKKRRRGAAAKIKEGVRMHEELMSGRQEDDIRGGGEIMEDYQPEDMAADQSLLALGTPLAAKKKGGTRGSIMPKPVEDSPVTRGGNSTYNSPRLDRTKAKEVETTLRYVQHLSNMQARPSSVSNTSTLNKSSGRTSKNTSMSLEQGTRKALKIINRGRTIHEEDEEESGDEESELSEEEEEGESFNY

>C_sp44_KNL2

MGDTDILPARVQNVLDMEIVRLNLWSIKFNASNIKLEGYVRSDDGNMMQKVCSEIICKRMNATMLFDVSGRFFELSGQIDREYQAKMGMPNRVIDEFVNGFPENWAYLINSCLTISQRSALRPIQAKEPIRSKGGPIVTLADETELVENEQRNSDSEKEKKKKEGEQRASENKRNQEAQRERERKEALAAAERERKRHEEDAERKRRKEEEDAANYTFRVPESQLGEPLTPIRFTRGNGGNGANIKPIFEQTPVRGKTTGPLASSTPQAPAPQQPRVLKDLEETVKPTAPQSSKAKPPPQKPQTEYASDSDLFAVPRPPADKNSRTTTSASSSDIGLGFLDQMDLLFDSAAVERTPTRHPKPRRIAVSPPNPRYRSHSKDRNRYGDFNSTRYTQRPSDDYDMGHMSRRDATFRRNDGGRDESRMSRKRGYYGSPDESDYGRRRDDRDRRDNYYEHEYDMKRSRPRETSSSSGKSVRFEDNYRRRREDPRETNYSRSYRDFDDFNTRGTSGDREDNRKLNDILRRENEVKARLQNHHKSSFSRRHVQSSDEESDDMADEWDRENQELMDNSMLFGDGIPKKKRGSDGGRPAKKQPVRKPQPKPTRKPAEPRKRAPRSPVDTDELNDSIASNRPRRACATPSTPAPKRITWPKRDLDKLKHVIQLKKPTEAEADWVEVARLLAKEGVDSELVKQTAISKLKWKPPVQEERNDDEDDEDKRRRRGAAAKIKEGVRMHEELMSGRNPVDDIRSGVEFVDDYQPEDVAADQSLLAMGTPLAAKTKGGTRGSIVPKPVEDSPIIRGRNSNYNSPRLDQTKAKEVETTLRYVQHLSNIQARPSSRANTSTMKKTSSHASKNTSMTLEQGTRKAMKIMNRGRTIREEDEEEDESDREDESEQSGEEEEEDDIYY

>C_sp48_KNL2

MGDTDILPARVQNVLDMEIIRLNLWSIKFNASNIKLEGYVRSEDGNMMQKVCSEMICKRMNATMLFDVSGRFYELSGQIDREFQAKMGMPGRVIDEFVNGFPENWAYLINSCLTVGQRSALRPIQSAPREPIRSRAEPIVTLADETELVENERKNSEAEKEKKKNERDEQRARENERNQEAQRERERKEALAAAAETKRRKEEEEERAEAERKRREEEEAANYTFRAPESQQGEPITPIRFKRGNGANGGYKSIFEKTPVRGKSNGPLASSTPQAPPPQQPRILSSLEREKKTEPDAPKSPIADRTVQKPIRNTEYADDADLFAVPKLPPIRNNRPSAAAPSSEIGYDFLDQMDSLFDTVVIDQTPNRNRMPRRAASSSMEERLRSSPPRNRDRYDDRESTRYSQRYGDDYNTSRMSRRDATFRRNDGGRDESRMSRKRGYYGSPDEFDHRRRDDRDRRGDYYEPDYKYDMKRSRPRENSSSSGRSVRFEDERFGDYRRHREDSRERKYSRNHREYDDNRGRRGSSGEDDRKLNDILRRENELMARLQNHRPSSSRRDASSSGEDEDDLANEWDRENQEILDNSMMFGDGLPKKSRRSAGGKIGRQPKKQPTRNPPPKPARKPEPRKKAPKSPVETDDLNDSIASHRPRRACATPSTPAPKKITWPKRDLDKLKHVIQLKKPTADDADWAEVTRLLAKDGVDSEVVKQTAISKLKWKEPVQEQERKEEEEEEERKRRRGAAAKIKEGVRMHEEMRRGRREEDTQMSADSMEDYQPEDVAADQSLLALGTPLVAKKKGGTRGSIMPKPVEDSPITRGRNSTLNSPKLDQTRVKEVETTLRYVQHLSNMQARPSSRANTSTVSKSGRGSKNTSMSLDQGARKAMKIINRGRTIDEEDEDEESNEEEEDHSEDEAFDY

>C_brenneri_KNL2_paralog1

MGDTDILPARVQNVLDMEIIRLNLWSIQFNASNIKLEGYVRSEDGNMMQKMCSEMICKRMNATMLFDVSGRFYELSGQIDREFQAKMGMPGRVIDEFVNGFPENWAYLINSCLTVGQRSALRPIQNAPREPIRSRAEPIVTLADETELVENERKNSEAEKEKKKKELEEQRAKENERNLEAQRERERKEALAAAERKRREEEEEAANYTLRAPESQPGQPITPIRFRRGNGGNGGHMKPVFEKTPVRGKSNGPLASSTPQAPPPQQPRILSSLEKPKSPIAERSAQKPMRRTEYADDADLFAVPKLPPIRNSRPSASAPSSDMGFDFLDQMDSLFDTVVIDQTPTRNRMPRRAASSSMESRIRSPPRDRERYDDRESTRYSQRYGDDFNMSRVSRRDATFRRNDGGRDESRMSRKRGYVSTPDEFDHRRRDDRDRRGDYYEPDYKYDMKRSRPRENSSSSGRSVRFEDERFGDYRRHREDSREPKYSRNHREYDNYRGGRGSSGEDDRKLNDILRRENELMARLQNHRPSSSRRDAYSSDEDQDDLANEWDRENQEIMDDSMMFGDGLPKKQRRSGGKIGRPPKKQRTRKPPPKPARKPEPRKKAPRSPVETDELNDSIASHRPRRACATPSTPAPKKITWPKRDLDKLKHVIQLKKPTAAEADWAEVARLLAKDGVDSEIVKQAAISKLKWKEPVQEQEKKEEEEEEERKQRRGAAAKIKEGVRMHEEMRKGRRENETQMSAESMEDYQPEDVAADQSLLALGTPLVAKKKGGTRGSIMPKPVEDSPITRGRNSTFNSPRLDQTKVKEVETTLRYVQHLSNMQARPNSRANTSTVSKSGRGSKNTSMSLEQGARKAMKIINRGRTIDEEDEDEESNGEEEEDHSEDEAFDY

>C_brenneri_KNL2_paralog2

IDREFQQNGMPVRSYDEFVNGFPETGLILLILASQEPIRSRAEPIVTLADETELVENERKNSEAEKEKKKKELEEQRAKENERNLEAQRERERKEALAAAERRRRQEEEERADAERKRREEEEEAANYTLRAPESQPGQPITPIRFRRGNGGNGGHMKPVFEKTPVVSEGQ

>C_wallacei_KNL2

MGDNHVLPARVQNVLDMEIIRLNLWSIKFNASNIKLEGYVRSEDGCSMQKVCSEIICKRMNATMLFDVSGRFFELAGQIDREFQLKMGMPSRVIDEFVNGFPENWGFLINSCLTMNQMSVPRPIQAAPREPLRSRNEPIVTLADETELADSGRKTSESERDKKRREREEQRMKENEKRLETEKEKERQKAQAAAAAEKKRQEEEEDAANYTLRGFQMESGDPITPIRFKRGQANGVHAKPTFVQTPVRGKPSGPLASSTPQAPLPEHPRRSLDLGKRSTPLASPSPKVQQPVSKPARETEYASDSELFAVPKLPAPKAARPSATSSAPSDTAFDFLDDMDVFFDTAVIDQTPVRARKQPRRVSPDVRRDRSSSRDPERFGDYDSTRNNQRYNNDYDMSRMSRRDATFRRNDERRDDSRMNRKRGYYNSPDDYDNRRSDYYESDSRYDTKRSRHRETSSSSGRSVRFQDDHRRSENDDYRRREGFRDPEYQRNYRDFEERRVRGSSGDEDKKKLNDILRREKELMARLQNNQKPSSSRRVSCSSDDDIDMADEWDRENQEIMDNSMMFGDGISKQKRRSGGSRQEKKKQPTRTVQPKPKRVASKKKSPVPSDDLNDSIASNRPRRACVTPSTPAPKRITWPKRDLDKLQHVIDLKKPSAEDADWVEVTRLLAKEGVESEVVKQIAISRLKWKEPVKNKNNEKKEEEEDETRRRRGAVARVKEGVRMHEELIRGRDDEERDVQVESMEDYQPEDVAADQSLLALGTPLAAKKKGGTRGSIMPKPVEDSPMARRQNSSFNSPRLDQTKAKEMETTLRYVQQISMMQARPSSRANKSKLNKTSARGSKNNSMSVEQGTRKALKIINRGRTIHEESEEEEEESDDDEFDEREEGDNSIY

>C_tropicalis_KNL2

MEDNHILPARVQNVLDMEIVRLNLWSIKFNSSSIKLEGYVRSEDGCSMQKICSEIICKRMNATMLFDVSGRFFELAGQIDREFQSKMGMPSRIIDEFVNGFPENWGFLINSCLTMNQMSVPRPIQAAPREPLRSRNEPIVTLADETELNDGGRKNSESEKDKKQKEREQQRMKENERKLQAEKDKKDKEAVAAAEKKRQEEEDAANYTLREFQTGSGDPITPIRFKRGTQSKGANNIKPTFLQTPVRGKPNAPLASSTPQAPPPEHPRRSLDLGKPPIQLASPSPKVQKPETVRETQYASDSDLFAVPRLPAAKNNRPPPSSSSETVLDFFDDMDVFFDTAVIEKTPVVARRQRRLSPEARRDRSSSRDPERFGDYDSTRYTQRYNNDYDMSRMSRRDAVFRRNDERRDDSRMSRKRGYYSSPENDFDRRRPEYYESDSRYDTKRPRPRENSSSSGRSVRFHEDHMGRGDEFRGSRYERGYREFDDQRGRGSSADEDKRKLNAILIREKELMARLQNKHKPSASRRVSYSSDEDSDDMAAEWERDNQEIMDNSMLFGDGIMQQKRRSARGHPAKKKQPIRTEQHKPAQKRAPPKKKSPAPRDDLNDSIASNRPRRACATPSTPAPKRITWRKRDLDKLQHVISLKKPTAADVDWVEVTRLLAKEGVEPEVVKQAAITKLKWKEPVRNDENMREEEDESRRRRGVVARVKEGVRMHEELMNGGNEVEEDLQVQSVEDYQPEDVAADQSLLALGTPLLAKKKGGTRGSIIPKPVEDSPIARGKNSSFNSPRLDQTKAKEMETTLKYVQQISMMQARPSFRANKSQMNNSTARGSKNTSMSVEQGTRKALKIINRGRAIQEESEEDEEDDDDDEFEEQEEGDNSIY

>C_doughertyi_KNL2

MGDTNILPSRVQNVLDMEIVRLNLWSINFNASNIKLEGYVRTDDGSMMRKVCSEIICKRMNSTMLFDVSGRFYELAGQIDKEYQLKMGMPSRVIDEFSNGFPENWAFLIKSCLSLPPRSTLRPIQDVPREPLRSRAEQPIVTLADETELVENERKNSDSERDKKRRDREEQRVRENEKRVEDQREKEHQEAIAAAAAEKKRKEEEESAADYTFRVPGTDLGNPITPIRFTRGGQLNGHIKSVFQQTPVRGKSSGPLASSTPQAPPPENPRRLSHLERASVPAAPLPEKDKTPAQRPVRSTDYASDSDLFAVPQLPPAKNIRPSVSSDSAFDFLDQMDSLFDSAVIEQTPGRDQKARKIAPPSPDARRRFSSREPERRSDYESSRYINRYNDDYNMSRMSRRDEPWNRRHDEGRDESRMSRKRGHFNSPPHDSDYGRRDNRGRRPEYYESNSKYDTKRYRPRETSSSSGRSVRFEDDYKRRDDFREPSYSRSYRMYDDRRGRESSRDLEDKNKLDAILRREKELVDRLENTSKPSSSRRHDLYSSDEDPIDLADEWDRENQEIMDNSMMFGDGISKKKRRSGAGRPQKKQPTGSTQPKPARKPAEPRKKARRSPPPADNLNDSIASNRPRRACVTPSTPAPKRITWPKRDLDKLKHVIELKKPTGSDADWAEVARLLAKEGVEAEVVKQAAITKLKWKEPVQIDNKEQEEEEEKKRRRGVAARVKEGVKMREELMSGKQEEDRALRVDSMEDYQPEDVAADQSLLTMGTPLAAKKKGGTRGSIMPKPVEDSPMSRGRNSTFNSPRLDQTKAKEVETTLKYVQHLSMMQGRPSSRANKSYLDKSSRGSKNTSLSLEQGARKALKIINRGTTIHEDDEDDDEEEDDDDEFEDQEVGDTSNY

>C_sp54_KNL2_paralog1

MGDTGIIPFRRKDVLERVEAVLNMEIVRLKLWSIKFNTSVIMLEGFVRSEEGTMIQKVCSENICKRISSTVLFDESGRFFELAGQIDREYQQKMGMPSRMIDEFINGFPENWADLINACVLNQQRSALRPIQAAPKEPLRTRAEPIVTLADETELIGEESKKDMKRREREEQREREQEKKLATEKERKRQEAAAVEKKRREEEEAEEAANFTFRVPQSQSGEAITPIRFKRGNGKKGENTRTIFDKTPVRQNSSGPLASSTPQAPPPQHPRRLSNIENNQPLRSPSQKTQLAASIPERETKFASDADLFAVPKLPPSKTTRPSTTSAVSSEGALDFFDEIDALFDTAIIEQTPTRDRKSRKPIATSPSPDPRRRSLSRDRKSERYGSYESSRYTQRYNDDYDNSRMSRRDVTFKRHEGGRDDSRMSRKRGYYRSPDDFEYSRRRDDREKDYYAYDSRHDTKRSRPRETSSSSGRSVRFEDDYRRRGGEFRESKDSRDFRDYDDHRNRKNSGEHEDKKKLKDILRREKELLARLQKSQKSSSSRRDSYSSEEETFDMADEWERENQEIMDNSMMFGDGLSKKKNRRSGGGGAAKKQLARTSQSKPHRKPAQPRKKTLRDPLETDDLNDSIASHRPRRACVTPSTPAPKKISWPRRDLDRLKHVIELMKPTTTDKYWIEVTRLLAKEGVDPEAVKQIAVAKLKWKEPVQNEEVLKKDEEEEKKRRRGAVARIKEGVKMHEELREGGDKKADDLESGVEAMEDYQPEDVAADQSLLALGTPIAAKKKGGTRASIMPKPVEDSPMAARGSNSTFNSPILDQTKAKEVETTLKYVQHLSMMQARPSSRANKSYLNKSSTRGSRNTSMSLEQGTRKAMKIINRGRTIHEDDEEDDEEDEDDEFDEQDGNTSIY

>C_sp54_KNL2_paralog2

MSEDPDEFLNLRSKVTDPWSVELIGLDAIQQQLDVMGENVAEVILKAEQSFNKKSLPFYEKRKKFTSKIDNFWKKAFLNHHFLSKAIPEEQEDLIEVLRDLEVQEFDDLRSGFKIIMTFDVNEFFENKVITKSYHLKSKPPKTQITEIQWKENQKPHMTSEDGDTTITFLEWLTHAVPPKSDEIAHVIKDDLFINPLQYYMKPDIQENGQPIRRIPKINSSMMEVDDTAEGEDSDEKEEEATNCTYRVPQSQNGEAITPIRFKRGNGKKRENTRTIFDQIPMRGNSSGPLASSTPQAPPPQQPRRFSNIENNRPAGPIPERETQFASDADMFAVSKRAPSKTTSSTSTSAVLFEGEGAYHFLDEVDDFFDTLVIETRERKPQRRRIATSPSPDPRRRRSSSRDRERYGSYESSSRYTRRYNDDDYDDSRMSRRDIATFKRRHEGGRDNSRMSRKREYYRSPDDFEYRRRRNDDRDKDYYNAYDSRHIAKRSRLGETSSSSGRSVRFEDDYRRSKDPRNFRDHRNRTISEKQLLARLQELRKPSTSSHRDSYSSEEEETFNMADEWERENQEIMDNSMMFDDGLSKKKIKRRSGRGAVKKQLSRTSQSKKPHRKPAEPRKKTIRDPLETDDLNDSIASHRPRRSCVTPSTNPVPKKISWPKRDLDRLKHVIELMKPTTSDKYWIEVTRLLAKEGVDSEAVKQIAITKLKWEEHVQNEEMMKMDEEEEKKRRRGAVARIKEGVKMHEELREGGDKKADDLKESGVEAMEDYQPEDVAADQSLLALGTPIAEKKKGGTRASIMPKPVEDSPMAARCSNSTFNSPRLDQTKAKEVETTLKYVQHLSMMKARPSSSKTNKSYLNKSSTRGSRNTSMSLEQGTRRAMKIINRDRTIHEDDEEDEADEDYEFEEQDGDTSIY

>C_inpoinata_KNL2_paralog1

MGDTNILPARVQNVLDMEIIRLNLWSIKFNSSNIKLEGFVKNEDGTAMQKVCSEIICKRMNSVMLFDVSGRFFELVGQIDREYQQKMGMPSRIVDEFLNGFPENWPYLIKSCISVDSRSSLRPIQAAPREPLRSRAEPIITLADETEVVVGDNKNSDTGNGRKHREQVERNSNDQVQMPREANMKNNIQEEEDDIANYTMRAPVSQNGEAITPIRFTRGNGKKGANTRSIFESTPARGTSSGPLQSSIASVPPPQQSRRLSTNQNSQLQSQCQKIQVAPSQNEKPPFREPQFSNDTDLFAVPRLQENKNNRLPSLSAASSDGAHDFFDEMDTLFDTANIEHTPARDRNIIRRSRVASPSPPLRRYRTSSRDCERDIYEPTRHSNRPDNGFNINRNRSDAFRKNEGRNDGSRMSRKRGYFYSPDKYRDNFSRRIKNIDYEQMHDNKHSRERENSPYSGRSIQFEDVYRKRNYGDYDDFRKRSRDEEEDRKLNEILRREKEMMARLQQHRKSVSSFPKSYLSDDDEFDMADQWERENQELLENSMMFGDGLPKKRKMSRDCKSERLKNPKESVKKPVQSQKSNRVKRLETDNANDSIALSRPRRLCATPSTPAPKKIVWPKRDLDRLKRIIELKKPTAANADWAEVARLLAKDGVELEAAKQTAITKLKWKEAVQNEELLNCEEEDKKRRRGLAARVKEGVKMHEEIREGGENKGGDLQNGVESMEEYRPEDMAADQSLLAQKTPIVVKRKGGTRASILPKPVEDSPLARGSNSALNSPKLDQTKAKEVETTLKYVHHLSQLHANPTCRGNKSFANKSLPGGRKNTSMSLEQGTRKAMKIINRGQTFYEDEEDEDDECSENDEDTSMY

>C_inopinata_KNL2_paralog2

MGDTNIVPARVQNVLDMEIVRLNLWSIKFNSSNIKLEGFVKNEDGTAMQKVCSEIICRRMNSVMLFDVSGRFFELAGQIDREHQQKLGMPRRIIDEFLNGFPENWPYLIKSCISVDSRSSLRPIQAAPREPLRSRVEPIITLADETEVVVGDNKNSDTGNGRKHREQVERNSNDQVQMPREANMKNNIQEEEDDIANYTMRAPVSLNGEAITPLRFTRGNRKKGANARSTFESTPAEGTSSGPLLSSITSGPPSQQSRQLSINQNSQLPSQCQKIRVAPSQNKKPPFREPQFSNQDLFDEMDSLFDTANIEHTPVRERSINRSRVASPSPPRRRYRTSSRDCGRDIYEHTRYSSRSDNGFNISGNRSDAFRKKEERNDESRISRKRGYFISPDKYRENFCRRIKNIDNDQMPDNTHSREGENSLYSGRSARFEDVHHKRDYGEYDDFRKQSRDEGKDRKLNEILRREKELMARLEQSRKSASSFQKSYSSDDDELHMTDQWERENQELVENSMLGDDLSKKRKVSRDGKTKQSERLKNHNESVKKPVQSEKRNRVKRLETDNANDCIALSRPRRLCATPSTPAPKKIVWPKRDLDRLKRIIELKKPTAADADWAEVARLLAKNGVELEAAKQAAITKLKWKEDVQNEELLNCEEEDKKRRRGLAARVKEGVKMHEEIREGGGLQNEVESMEEYRPEDQSLLALKAPTVVKRKGGTRASVLPKPVEDSPLARGNNSALKSSKLDQAKAKELETSLKYVHHLSQMHANPSSRGNKSHANKSLPCGKKNTSMPLEQGTRKSMKIINRGRTFCEDEKDDSTDNDEDT

>C_elegans_KNL2

MGDTEIVPLRVQNVLDSEIIRLNLWSMKFNATSFKLEGFVRNEEGTMMQKVCSEFICRRFTSTLLFDVSGRFFDLVGQIDREYQQKMGMPSRIIDEFSNGIPENWADLIYSCMSANQRSALRPIQQAPKEPIRTRTEPIVTLADETELTGGCQKNSENEKERNRREREEQQTKERERRLEEEKQRRDAEAEAERRRKEEEELEEANYTLRAPKSQNGEPITPIRFTRGHDNGGAKKVFIFEQTPVRKQGPIASSTPQQKQRLADGANNQIPPTQKSQDSVQAVQPPPPRPAARNAQFASDADLFAVPKAPPSKSVRNLAASNVDIFADVDSVLDTFHFESTPGRVRKPGRRNVSSPSPEPRHRSSSRDGYEQSRYSQRYEHDNSRWSRHNATYRRHEDESRMSRKRSIVRDDFEYSRRHDDGARRRDYYDADIQGDSKRYRGRDASSSSGRSVRFEEEHRRHGDEYRDPRGPRDYNDYGRRRNHANSRSGEDEEKLNAIVRREKELRNRLQKSQKASSSSYRHRSNSSDAEESLNEWDIENQELLDNSMMFGDGIPKRSNARKDKFVKKQATRSKPANSTKSPAQARKKKRASLEDNRDLNDSIACNRPRRSCVTPVAKKITWRKQDLDRLKRVIALKKPSASDADWTEVLRLLAKEGVVEPEVVRQIAITRLKWVEPEQNEEVLKQVEEVEQKRRRGAVARVKENVKMHEELREGGNHRAEDLQSGVESMEDYQPEDVAADQSLLALRTPIVTKKRGGTRASIMPKPVEDSPMSRGNNSTFNSPRLEQTKAKDIETNFKYVQHLSMMQARPSSRLKKSSSMNNSTYRGNKNTSISLEKGTQKALKIINRGTTIHEDDENEDNDDDDDMREEDTSIY

>C_oiwi_KNL2

MGDVEILPQRVQNVLDMEIVQLKLWSIKFNSTNIRLEGFVRNEEGTMMQKVCSENLVKRMNSTMLFDVSGRFYELSGQIDREYQQKLGMPSRIIEEFANGFPANYAILINSCLNSDQIRSIRRPIEAAPRDPLRPIVTLANETENPKIPEKIPEKSILEEEKEENEADFTLKAPVSTDGNAITPIRFTRGNGRALARTVFEQTPVRGKASTSGPLASSTPQPAPPPIPRRSSTTENRNPPEAPPPKVQYATDSDLFAVPKLPAPRQQAPPISLGFLDDFDDIFADAVIQKTPKRGRPPKREEEDSRRFYEDESRMSRKRGYYRSRSRSPDYDRRHRDRDYSDFYGREDSKRPRRRDESSSSGGGGKSVRFEREDRHQRHHPHRRDRSNSRESQRHHHRRHHDDDYNVSRSRSHEKEEKRRLQELLRKEKELEARLQSFRRPPTSSSSSSEDSDEDEMAGEWERENQEIMDNSMMFGDGISKKRRRQSGEKKRKQPSRPKPKSAAPAPSATRKPAKKNKRSPAETDSLNDSIASHRPRRACATPSTPAPKKVTWPKRDLDRLRRVIELKKPSGREEEWVEVARLLQKEGVNPMEVKEAAIGRLKWKEPVEKEHTEEEEEEEKKRRRGVAARIKEGVKMHEELREGRQQNREDALKNRVEEVEEFQPEDVEADQSLLALTTPVALKKKGGTRASIMPKPVEDSPMSRGNNSTFADSPRFDQTKAKNLETTAKYVHHLSILQGRPSSSAANRTQMNRSTTRGGGSKNTTLSLEQGTRKAMKMISEGRTIHEDSEEDDEDDEDMDDDVFN

>C_kamaaina_KNL2

MGDTEILPLRVQNVLDMEIVRLKLWSIKFNSTNIRLEGFVRNEEGTMMQKVCSENLVKRMNSTMLFDVSGRFYELSGQIDREYQQKLGMPSRIIDEFANGFPENYAILINSCLNSDQIRSIRRPIEAAPREPLRQKQEPIVTLANESEIQNPQKTKIVPSKSPKNSKILEKSIQEEEEEEEDPANFTLKVPISTNGDAITPIRFTRGNGRGMARTVFEQTPVRGKPSSSSGPLASSTPQPPPPPRRSSTTENRNPKTEQAPAPKPPQNPQYATDSDLFAVPKLPAPKQQAPPISLGFLDDIDDIFADAVINKTPRRGRIGRPTSPSKQDVTSDSRGRFYESRRDEMVDESRMSRKRGYYRSRSRSPDYDRRRRDHRDDHRYSDYYERDSRSDSKRPRQREESSSSGKSVRFERREDRHPDRRRHRSTSRESHRHRRRDDDYDVSRSRSRESRKNEKRKLQKIMKREKELMSRLQNSRRHSPSSSSSSSSSEGSDDEDEMAGEWERENQEIMDNSMMFGDGISRNRKRKSEGKKRKQPSRPKQIKSEAPVPAKKSAPKRNKRTPAETDFLNDSIASNRPRRACATPSTPAPKKITWPRRDLDRLKRVIELKKPSGKEEDWVEVARLLAKEDVKPMEVKEVAIGKLKWKEPVEKVQTEEEEEEEKKRRRGAAARIKEGVKMHEELQGGNPLKREDALKNRVEDVEEFQPEDVEADQSLLALTTPVALKKKGGTRASIMPKPVEDSPMSRGNNSTFANSPHFNQTKAKELETNYKYVQHLSMMHARPSSSAANKSQMNRTTTRGGSKNTSMSLEKGARKAMKMISEGRTIQEDSEEDDDDDKEEELEDDVFN

>C_waitukubuli_KNL2

MGDTDILPVRVQNVLDMEIIRLNLWSIKFTATNIKLEGYVRNEEGTMMQKVCSEVICKRMNATMLFDVSGRFFELSGQIDREFQIKHGMPSRIVDEFTNGFPENWAFLISNCLTTEQRSAIRPIQAAPVQPLRLREPIVTLADETELGDRTVVRKENEKDQRKKNREEEEKKHRLLEEKRKKEQEDKIAAGKRKESREEAERKQKEEEDAANYTFRAPPSFGPDAITPIRFTRGTQGRGGIRIFEDTPQRTTSGPIDSSTPKPPPVHEQLPIKVQQKPEVHQQPEPRQTEKAPQTFASDSDLFAVPRLPARAPISSELPTGFDFYDEMDAIFDNAKIDKTPVMESRRKQIPRRFQYEPSPPPILHGQSSSSSQAYEPDFDSRSYYNGRYDHERSRMSRRDDRYVSERDDSRISRKRILYSPDHHDDSDHSRRRAYGSRSRYEDDFPKRSRTRETSSSSGRSVRFEDDYRSNRYHNDREHRDRDRTREQQERDMYESQKLKEIMRREKELEAKLRLKKTATRRVSYSSDDSADDTMNMTEEWERENQEILDNSIMGDRKTSRKRNNAPKREAVKPKKVDKPARAPAKRTKKEPRQAPDLNDSIASNRPRRACATPATPAPKRVTWPKRDLDKLRHVIDLKKPSASIEDWTEVARLLKKDGVEPADVKLIAETKLKWKEPVQDEDVLLQEEEEEKKRRRGVAAKVKEGVRMREEMREGGAKEDELRNRVESLEDYQPDDMDADQSLLALTTPVAAKKKGGTRASIMPQPVEDSPAVKGNNTSSFMNSPKLDSTKVKEVETTLKYVQHLSTMQARPGSSMNKSYMNASSSRGRNASISVEQGTLTTHIVVSIFLRMQMTFHLTPSDTVLFESWKVTDGWTMMGACALVVVAGVFVEAIKRYRKKINQDQMIREQLVYEPFSNRLFASML

>C_panamensis_KNL2

MGDTDILPVRVQNVLDMEIIRLNLWSIKFSATNIKLEGFVRNEEGTMMQKVCSEVICKRMNSTMLFDVSGRFFELSGQIDREFQIKNGMPSRIVDEFMNGFPENWAFLISNCLTTEQRSAIRPIRAAPVQPLRPREPIVTLADETELPGDRTMSKKETERERKNKERDEEERKQALLEENRQRDRERELQKEKEREAVERRKKEEREEAERKRKEDEDAANYTFRAPPSFGSDAITPIRFTRGKKGAVVTRIFDDTPVRSTGQPLASSTPQRSLPVKEQPPQLPKQLKEDSKEQQPPPRQVERPMQSYASDADLFAVPRLPSRAPAPTGLSGGFDVLDEMDSLFDNAKIEKTPVMGSRRKPIPRRYQYESSPPPFLRGQSSSQQAYERDLSSASYYSSRYDDNRSRMSRRDDTSRRYDSGRDESRMSRKRGHYSPPDRRDDYAYNRRREDESRQRDSFRYDEDYNYDSKRSRPREASSSSGRSVRFEDDYCSNRSRENKDRRNHERVRDQRDRNMYESRELKEIMRKEKELEAKLRSKRSSVRRVSYSSDDSDNDTMADEWERENQEMLDNSMIMDRKPSNKRKNVPKKETSKPKRVEKPTRAPAKKNKKDPVETDDLNDSIASNRPRRACVTPSTPAPKRITWPKRDLDKLRHVIELKKPSGNLDDWAEVTRLLKKAGVEPTDVKQIAETKLKWKEPVQNEEALLLEEEEEKKRRRGAAAKVREGVRMHEEMRNGGKKEDDLRSGVESMEDYQPDDMDADQSLLALATPVAAKKKGGTRASIMPQPVEDSPAVKGNNTSSFMNSPKLDSTKVKEVETTLKYVQHLSTLQARPGSSMNKSHMNRSTSRGKNTSISVEQGARKAMKIINRGTTIHEGDEDDEDDDDEVTEDDEENVIY

>C_nouraguensis_KNL2

MGDTDILPVRVQNVLDMEIVRLNLWSIKFTATNIKLEGFVRNDEGTMMQKVCSEVICKRMNATMLFDVSGRFFELAGQIDREFQVKHGMPSRIVDEFINGFPENWAFLISNCLTTEQRSAIRPIQAAPVHSLRPKEPIVTLADETELAGDRTMARKESEKDRRNREREEEKRKKLLEEKRQREEELQRERDREEERSRQEEEEKRRQEEEEAEKLRKEEEANNTFRAPKSFGTDAITPIRFTRGNSKKGLGAIKIFDNTPTPKRKNGGPLASSTPQPQRLPLVKEQSQKEKTPQPEKPPQQQQESSRPAERPKQTFANDADLFAVPRLPTKSTASSGFSSGFDMFDEMDTLFGTAKIDKTPVMESKRKPIPRRYEYESSPPPFRGQSSSSQTYQRDHDSRFNFNDRYDDDRSRISRRDGTFSRYDSGRDESRMSRKRGLYSSPDQRHDYEYSRRKEDESRYRDRSRYENRYDPKRSRARETSSSSGRSVRFEDDYRSKKYREDRREHDRTRDHRERDMYEDRKLKEIMRREKELEAKLHSQRTRRVSSSSDDSADETVNMADEWDRENQEMLDNSMMMDTKTSRKRKNVPKREFVKPKSVEKKPARASAKKNKRDPIETVDLNESIASNRPRRSCVTTAPKRITWPKRDLDKLRHVIDLKKPSADLDDWAEVTRLLKKDGVQPAYVKLIAETKLKWKEPTQDADVLRLEEEEETKRRRGAAAKVKEGVRMRQEIRGGGEKEDDLRRGVEAVEDYQPDDMDADQSLLALATPVAAKKKGGTRASIMPQPVEDSPLVKGNNTSSFMNSPKLDSTKVKEVETTLKYVQHLSTMQARPGSSMNKSSLNRSSSRGKNTSISVEQGTRKAMKIINRGTTIHEDDEEEEEDENSEDDEENAVY

>C_becei_KNL2

MGDTDILPVRVQNVLDMEIVRLNLWSIKFTATNIKLEGFVRNDEGTMMQKVCSEVICKRMNATMLFDVSGRFFELAGQIDREFQIKHGMPSRIVDEFINGFPENWAFLISNCLTTEQRSAIRPIQAAPVQPLRPKEPIVTLADETELAGDRTVVRKETEKDRRNREREEEKRKKLLEEKRQREEEIQREKDREEEKRRQEEEAERKRKEEEEANNTFRAPKSFGTDAITPIRFTRGNAKKGLGAIKIFDNTPTPKRINGGPLASSTPQRPPPVKEKTPQPEKPPQKQQEAPKPAEISKQTFANDADLFAVPRLPTKSTASSGFSSGFDMLDEMDTLFETAKIDKTPVVESRRKPIPRRFEYESSPPPFRGQSSSSQAYQRDLDSRSYFNERYDDDRSRMSRRDGTFSRYDSGRDESRMSRKRGLYSSPDQRHDYDYSRRREDESRDRDRSRYDYRYDPKRSRARETSSSSGRSVRFEDDYRSKKHREDRRDREDRREYDRTRDQRERDMYEDRKLKEIMRRERELEAKLQAQRKSARRVSSSSDDSADETVNMADEWDRENQEMLDNSMMMDTKTTRKRKNVPKKEFVKPKSVAKPARASAKKNMREPLETVDLNESIASNRPRRSCVTAVPKRITWPKRDLDKLRHVIDLKKPSANIDDWVEVTRLLKKDGVQAAYVKLIAETKLKWKEPSQDADVLRLEEQEETKRRRGAAAKVKEGVRMRQEIRGGGEKEDDLRRGVEAVEDYQPDDMDADQSLLALATPVAAKKKGGTRASILPQPVEDSPLVRGNNTSSFMNSPKLDSTKVKEVETTLKYVQHLSTMQARPGSSMNKSSLNRSSSRGKNTSISVEQGTRKAMKIINKGTTIHEDDEEEEDEDDENSGDEEENSIY

>C_yunquensis_KNL2

MGDTDILPVRVQNVLDMEIVRLNLWSIKFTATNIKLEGFVRNEDGTMMQKVCSEVICKRMNATMLFDVSGRFFELAGQIDREYQIKHGMPSRIVDEFMNGFPENWAFLISNCLTTEQRSVIRPIQAAPVQPLRPKEPIVTLADETELGDRTMAKKETEKERRNREREEERKQALLEEKRQRDHELRQEKDREEERKRKEEEDAANNTFRAPKSFGADAITPIRFTRANGKKGHGAFKIFDNTPKRNTNEPLASSTPQRPLPVKEPEQPPQKQPETSRPAEPSRTYASDADLFAVPKLPTKSTGSSGISSGLDFFDEMDTLFETAKIDKTPVMESKRKQIPRRFEAASSPPAFRGQSSSSHASNRDYDSRSYLDDRFNDDRSRMSRRDGTFNRYDSGRDESRMSRKRGHYSPDQRRDYEYSHRRDDESRYRDRSRYEDDYRYDPKRSRAREASSSSGRSVRFEDDYRPNYHREDHRQRDRTRDGNDVYENRKLEEILRRERELEAKLRAKKKPARRVSCSSDDSEDENVDMADEWDRENQEMLDNSMMLNRKTARKQKNVPKRDVAKPKRVEKPIRAPIKKKTRDPVETDDLNDSIASNRPRRACVTPSTPAPKKITWPKRDLDKLRHVIDLKKPSANIEDWVEVTRLLKKVGVEPADVKGIAETKLKWKEPTQNAEVLRLEEEEETKRRRGVAAKIKEGVRMREEMRNGGEKEDDLRSGVEALEDYQPDDMDADQSLLAFATPVAATKKKGGTRASIMPQPVEDSPLVNGNNTSSFMNSPKLDSTKVKEVETTLKYVQHLSTMQARPGSSMNKSYMNNSSSRGKNTSISVEQGTRKAMKIINRGTTIHEDDEDEEEDDDEVSEDDEDNAIY

>C_macrosperma_KNL2

MEYIVMGDTILPVRVQNVIDMEIVRLNVWSIKFTATNIRLEGFVRNEEGTMLQKVCSENICKRMNATMLFDVSGKFYELAGQIDREFQIKHGMPGRVVDEFLNGFPENWAFLIGTCLTTEQRSVLRPIQAAPVQPLRPKEPIVTLADETELTGDRTMAKMETEKERRAKQREEQRKLEKEREEEAVAERKDKEKREREEAERKRKEEEDAANYTIRAPNPLGNEAITPIRFHRGKGGGPRFFNVFDKTPKATSSAPLASSTPQARPPVRKEPEQPVSSKPVERPVAFASDADLFAVPRLPAKGPTTSSSSAGAPLQDDFDFLDEMDTLFDTAKIEKTPVMDTRRKPIIRRFEHRSYSPQHGQSSSSQAYEMDYNDRREPYYHDRYEDDRSRVSRRDFGGDASRMSRKRGYYQSPENRDRDEYEDRRRREHESRYSQQIYSREFEMNGWPRRETSSSSGRSVRFDDDYHYSDRHRNPRDKTRDERDREMYERQKLKEITRREKELEAKLKSRKVSRSSEESADDSMDMAEEWERENQEMLDNSMMMMPDRKSSRKQKVVPKREKPKKAAEKKPAARAPRKKNIREPLLEEEDDPNDSIASSRPRRACATPSTLAPKRITWAKRDLDKLNHVIELKKPTANLDDWVEVTRLLKKNGVEAADVKHAAETRLKWKEPVQNDERAQEDEIEEERKRRRGVAAKVKEGVRMREEMREGGEREDEMETRVGAVEDYQPDDMDADQSLMALGTPAVKKKGGTRASILPQPVEDSPIVRGNNTSSFMNSPRLDQTKAKEVETTLKYVQHLSTMQARPGSSMNKSHANKSYMNKSTSRGGKNTSISVEQGTRRAMKIVNRGTTIEEDEEEDEEENDDDEEDTSVY

>C_sulstoni_KNL2

MEENGILPTRLQNLADMDVIPLELWSIKFYGTRYKVEGIVRPEDFTTAQKFTSEFIIKRITSTALVDASERFIELVGQIDRTFQIRNGMPEEIVGKFQNGFPRDWWELLKPCLAAGQKTSTMRAIEAPPLKTSTTSSTVQNVQKTSSEPIVTFADETEVQSDREKERRHRQEEREVQKVLEKDDDDDADDADYTIRQPRAIDGGENTPLRYKRGHSGGPQNRNRVFADTPQRTAIPLAASTPNAPPKIAKPPVKFASDADLFAVPVPPTRRPAERKERSESPEGCDGLFDKISEVFSNVNFPVTPLPAARVRAHFSPAERSEGSYYYRDDRYDEDRNRTSRRDRSRSREYDESRTSRKRGYYRSPEGYEYSRRRDDRRRDRYEASSQRHGSSSSGRSVRFEDPYEYDVEQRRRRREDQKLREIERREREIEARLERSKRSRRYSSDEQSEDMADEWERENRQILDDSMLFRSAAGKATNKKRAGRPKKEVKTRSPPVQKPQKARKEPRKRRSEPAETVDLNESIASNRPRRACVTPVAPPPPKRVTWPKRDLDKLRRVIEVKKPTGDEHDWVEVARLLAKDGVTPTDVRTIAETRLKWREPTRDPKVLEAAEEEETKRRNGLAARVKESIRMREEMREGRQEEDVELKPGGENYEPADVSADESLLAIGTPRVPPKKKGGTRASLIAPPVEDSPMTRRSGAAATVVASPKLDATEVAATQKHVHHLSMIQARPGSSMMNTSRANKSIRNGKNTSISVEQGARRAMKMVTRGRSTIAEEDESEEDDDENEEDVTAEEDEIY

>C_afra_KNL2

MEENAILPTRLQNLAEMDVIPLELWSIKFYGTGYKLEGIVRPEDFTSAQKFSSELIIKRITQTALIDASERFIELVGQIDRTFQLRNGMPEEIVNKFQNGFPKNWYELLKPWASQNQKTSTWRAIEAPPVAPPNPQIPPIPQKAPCEPIVTFDDETEVQSDKEKEEARRRRQLEVEEAEKQRKILEDLNDDADYTLRNPRSIDGVQNTPLRFKRGNPNFERNPRVFNDTPVRTGGGPPLAASTPQAPKVSDPIPKAANYASEGDLFSFAVPTQPPSQRPTRTTDEDRKHQADRRRRSESLGGSDDGGGGSGLDLFSQMSEVFSNAKFPITPLPAAPPRSSRFERERAERSDYYREDRDRERYRDPSRESRERDESRMSRKRGYYRSPEGYEYSRRREDRHRDRYESSYQRYGESSSSGRSVRFDDGYDDPYEREEEERRLRRHRDNQKLREIMRREREVEARLERSKRSRRYSSEDDGPEENDDDLADEWDRENREILDNSMSFARGGSGRAGKKRGAGRPKKTSAPKSRSPPTQQKPRKQQQAAKKRRSEPAETDDLNDSIASHRPRRACTTPKAAAPKRVTWPKRDLDKLRRIIEVKKPTGDADDWVEVARLLAKDGVEPADVKTCAETRLKWKEPTRDPEVRQKEEEEEKKRRNGLAARVKQGIRMREELREGYEEDVAPPPTSSSQEPDYEPDDVSADESLLAMGTPLPPPKKKGGTRGSIVVLPVEDSPMVRRSGVAASPKFDASDLAATQRHVQHLSMIQARPGSSMNTSRASKSILGGKNTSISIEQGARRAMKMMNRGRVAIAEEDEDSDSDDSDDDENDSNDNAEYIKIKVVGQDSNEVHFRVKFGTSMAKLKKSYADRTGVSVNSLRFLFDGRRINDEDTPKSLEMEDDDVIEVYQEQLGGGI

>C_afra_KNL2

MEENAILPTRLQNLAEMDVIPLELWSIKFHGTGYKLEGIVRSEDFTSVRKFSSELIIKRITQTALIDASERFIELVGQIDRTFQLRNGMPEEIVIKFQNGFPKEWYELLKPWASQNQKTSTWRAIEKPPVVPPNPQIPPLTQKTPCEPIVTFDDETELRSDKEKEEARRRRRLEVEEAERQRKLLEEGSDGAEKRGYNRSPAEKNNRERSERSRRCSSEDNRSDGELADEWERVNSEILDNSMSRVAGRPKKMLAFGRALTQADVEKFAIRLEIAGARIPAVVASEKYVVTTLKNIPEASRKLGFALSAFDHLEQRIELKIHGMHRHSDFVVLRRDEGEFSEAPVYRNEEVGDKYFVVKRSIDGKKIVCHGKVTSVGATGFAFVSEPYEGFPVANGDGAFAFLDGKLLGVVKDTKEQGRVLEFVSIYLVHFAIDIIFHHQPFSV

>C_sp49_KNL2

MTERTIVPVRVQNVMDMEILRMNVWSIKFNSSSVMVSGFVRTPDGNSMNLVTSEAICKRMNSTLLFDISGRFYELLGQIDKSTQLKAGMSSRVMEEFLNGFPDNWVFLVGQCLTTEQRSAIRPIQAAPREPIRPKSPIVTLADETELREQGKAQKEQKEDADATFRAPTAFFHGAVTPIRFTRGNGATGRSQRVFESTPSREPSSSRPLASSTPQPQKLPVLPPVAPRDVPAAQPRTFATDDDLFAVPKLPASSSGLDFLEDMEAIFNGAHINRTPSHAAPRARVPREFRSRSPSYEQRYQDRRYQDESRVSRKRGMDRSPEYYEMERRAEESRRSHESKRARPRETSSSSGRSVRFEETYRRDRHHQNDRSHRSHHYEQEKEHKRRLREIERREREVEERLKRNRKVSSDSNSSSGEEDGSFNEWDVENREILDRSLMEASGRERRPKREPTSKKSTSKKPAAPKPKPTKNIRKKRDPIETDDPLNTSIASSRPRRACATPATKKITWPRRDLDRLKRVLEMKAPTGRSEDWVEVARLLAKDGVNSEEVRQIAESKLRWKEPEHHPEEHQEDPLNDSQLIEKKRRNGIAARVKESVKMAEELQTRPPNSDEMRGRVEEVEEYQPADVSADESLLALSTPQVIKKRGGTRKSTLPEPIEDSPMARGANSSVNSPRLNATKAKEQETTLRYVQHLTTMQTRPASRMNKSAMNKSTASAKNTSMTVEQGTRHVQKMMKKAMAIDEEEEDDEESDENQEETSFY

>C_sp25_KNL2

MTEQTVVPVRVQNVMDMEILRMNVWSVKFNSSSVMVVGFVRTPDGNSMNLVTSEAICKRMNQTLLFDVSGRFYELLGQIDKDTQLKQGMTSRVMEEFINGFPENWVFLVGQCLTTEQRSAIRPIQAAPRQPIRPKSPIVTLADETEVAAQAQAQREQKEQKENEQRRRREDKEKEDADATFRAPASFANGAITPIRFTRGASVASRNQRVFDATPTRQAPLASSTPQQKVQKVPTIAADEDLFAVPKLPVKKGVAPASDLDFLDDMDALFTGARINRTPSHRVPPKPQQRRRSRTRSRSPSIDRSRSRRYEEEFDSRMSRKRGYDRSPDYDDYRREESRRRFKESKRSRPRETSSSSGRSVRFESDYHRQRHDDRRRHEDREELERQERRRLKEIERREREVEERLKRSRRKNYSSSSSSDEDQDGSFNEWDAENREILDQSLLASTGGSSRSASGGNSRGRLPKREPSLKKAKPAAAAPKPKPRKKKEVVDLNDSIASNRPRRACATPATPAVKKITWPRRDLDKLKRVIELKKPSASEEHWTEVTRLLAKEGVAWTDVKSIAETKLKWKEPRQEMVEVEEEEEEERERKRRRGIAAMVKESVKMCEEIQTRNVQEDEMEGRVEEVEEYQPADVSADESLLAMSTPQITKKRGGTRKSTLPEPIEDSPMAKGTNSSWNSPRLNATKAKEQETTLRYVQHLTTMQTRPGSSMNKSYMNKSTASTRGKNTSLSVEEGTRKVQKMMKKAMAIEEEDEEEEDSDDEGETSLY

>C_imperialis_KNL2

MTESTVVPVRVQNVMDMEILQMNVWSIKFNSSSVMVVGFVRTPDGNSMNLVTSEAICKRMNSTLLFDVSGRFYELLGQIDKTSQLKQGMSARVLEEFLNGFPENWVFLVGQCLTSEQRSAIRPIQAAPRQPLRQKSPIVTLADETDLGDRHKREEVENEARKREERKMIERENEEKRRAEERRQREEQEKEDANCTLRAPASFASGDLTPIRFTRGNGPCNRKNRVFEDTPSRETTSRPLAASTPQQKPKPVDPPREVPRDVPARPKTNFATDDDLFAVPKLPANRAAKPSSDLDFLDDMDALFAGAHINRTPSHAPKPVSRIRQSPPRRRSPSENRSRHEYSQRSNRYDDDYDVRFNRNGKEESRLSRKRGYYRSPDEYYETRRDWGRDRSRDRSRQRYDSERYDSRYDSQVSRPRETSSSSGRSVRFEDDYSQRNNRRYEHEVRERKQLKDIERREREVQERLQRSKKSAPARQATYSSSEDDDDASFNEWDAENREILDQSLLASTAGTSRRRSAPKREPKSKKPKPAPAPAPKRKPSKPRKQRDPVETDDLNDSIASNRPRRACATPATPAPKKITWPRRDLDRLKRVIEMKKPTASDGAWAEVARLLAKDGVTPADVKQIAETKLKWKEPTQDPVEIEEQEEVELKRRRGIAAKVKESVAMREEMQRRAVDEDELNGRVEEVEDYQPADVSADESLLAMGTPQVTKKRGGTRKSTLPEPIEDSPMARGNNSSFMNSPRLDATKAKEQETTLRYVQHLSTMQNRPGSSMNKSQLNKSSTSRGKNTSMSVEQGTRKAQKMIRKAMAIEEEDEEDDEDDEDENREEEETSFY

>C_japonica_KNL2

MAETGVLPLCVQNVMDMEIVRLNLWSIKFYAESIKLEGFVRNSDGNMMQKVCSEAICKRMTSVLLFDVSGRFFELVGQIDHEFQVKNGMPSRIVAEFMNGFPENWLFLIKQCLPPTDEQIPQRQIQIQSAPAPPKEPIVTLADESEHVSDHPKKIVENGRIRDEESADCTFRVPHPVAIDAITPIRFTRGVRKENAQRIFDSSTPKPQVEKQKKPLAAFDADLFAAPAPARAPPPTRAPPPNPPGPSSEFVLLDEMDELFNNAEFDKTPGRTVAQPRRINETSRMDRKTSRTYSEDCDRKYDDRRVECYSRSARENVEPNRNEFRMNRKRPSVNNYADEMNGGYNYRSNKYGRREVFESNGYHSDEFRSREAPSSRSYADRDYYSRKSYHRGEEQRNEKMYGDILRREMALETRLKNGRSQRHVSESSEDSMDDSYMVDEWDRENQRILDSSIVSSFGRFETSPKKSRIYVKKERTAKKARPKPVPPRRKKNSPDSPEMDDLNDSIASNRPRRACVVPSTPVPKRVTWPKRDLDRLSRVIELKKPSSSDSDWTEVARLLAKEGVEGADAMKIAMAKLKWKEPVKDCETVQEDLEMVDKKRKRGFAAKVKESVQMMEEMRDGGEQSGEMGGQGVEQAEDYQPADLSADESLLALATPCAGKKKGGTRRSVMPAPVEDSPVIKLNECSYANSPKLDVTKCKEVETTLRYVHHLSTVQARPGSSSSRSYKNRSVCLGKSLSLEQGARKAQKLISMAVGIEEEDEEEEDDDDAELSIF

**Supplementary Data S5: All KNL-1 sequences used in this study**

>C_tribulationis_KNL2

MGDGDVVPARVQNVLDMDVVRLNLWSLQFNSSGVRLEGFVRSEDGNMMQKVHSGIICRRMSATMLFDVSGCFYELAGHIDREFQQKLGMPSRVIEEFMNGFPDNWAFLIKSCLPAEPKSALRPIQAAPEPSKARGEPIVTMPDETIVDNQSAEKERKRKELQKKKEQEERVRIQREQRAAAEAAAERKRQEDEEFKRREEEEERKRQEEENDAANYTFRVPQSQNGDAITPIRFTRGNGQKGANVRSIFEKTPRTKPAGPLASSTPQAPPQQHAPRISIAEAKQPDEPPHPHPKPQPPTRESIRETQYCSDSEFAVPKLPAPRNNRHASASSQPAPLDFLDEMDALFETANVDHTPGRGRKSRKPVSRSPSPGRFNSTARDRTGYDRYEPSRQSSSQRYYDDYNNVSRMSGRNHTLGGHDMRRDESRNSRKRGYNNSPEDYSRRFDDRSRRRDNYESDSRYDSKRSRPRDQSSSSGRSVRFEEDFPRNRRDESRDSRSYRHYEDHRNRESSGDREDKRKLDAIVRREKELVARLQNTQRSSSTLQRSGYSSEDDEMSDEWDRENQEMLDNSMMFGDGLSKKKGKSGGHKTMRQAKTRYPPKPKPAQKPAQPKKKKKDVEYESDEMNDSIASNRPRRACATPSTPAPKRITWPKRDLDRLKHTIELKKPTGAEADWAEVTRLLAKDGVDAEVVKQTAIAKLKWKEPSQKTIQQEEEEKKRRRGATARVKEGVRMHEELREGGNNRAESSQSGVEAVDDYEPDDVAADQSLLGLQTPIAVKKRGGTRASIMPQPVEDSPVVRGNNSTLNSPRLDQTKAKDVETTLKYVQHLSMMNARPSSRANSSYLNKSSSRGGDKKNTSLSVEQGARKALKIINRGRTIHEEDEDDEDEDITID

>C_sp41_KNL2

MGDGDIVPARVQNVLDMDVVRMNLWSIRFNGSGVTLEGFVRSEDGNMMQKVHSGIICRRMSATMLFDVSGCFYELAGHIDREFQLKIGMPSRVIDEFMNGFPDNWAFLIKSCSTSEPKSALRPIQAAPVEPLKSRSEPIVTMPDETVLAESQTAEKERRRKELQKKKENEERLRIQREQQEAVKAAEKKSQEEEKDKLQEEERKRQEEEDAANYTFRAPKSQSGEAITPIRFTRGNGQKGANVRPIFDKTPVRTKPAGPLASSTPQAPPPQHPHRLPNIETTRPDASVPPVPQPPIRETQYCSDSEFFAVPKLPAPKNIRGTSAKPLGFLEEMDALFENVNVDQTPGRVRNPRKVSRSPSPMRLNSSARNRDSGYDRYEPSRQSLSQRYYDDYTTSRMSGRDDTFGRNDTRRDESRNSRKRGYNNSPDEYRNRWDDRSRRHDNFESDPRYDSKRSRPRDQSSSSGRSVRFEEDHPRSRMDSRESMDSRNYRHYEDSRNRQSSGDREDKRKLNDILRREKELMARLQNSQRSSSTFQRTAYSSEDDEMSDEWDRENQEMLDNSMMFGDGLSQKKRRSGGRQPAKKPNYLQQAKTKRVPQPKPAPKPAQQKKKKKQDSEDERDEMNDSIASNRPRRACVTPSTPAPKRITWPKRDLDRLKHTIELKKPTGADADWAEVTRLLAKDGVDAEVVKQIAITKLKWKEPTQQTIQMEEEEQKRRRGATARVKEGVRMHEELRAGGNDRGNSSQGGVGAVEDYEPDDVAADQSLLGLQTPIAIKKRGGTRASIMPEPVEDSPMVRGNNSTLNSPRLDQTKAKEVETTLKYVQHLSMMNARPNSRANTSYFNKSSSRDKTNTSLSVEQGARKALKIINRGRTIEEEDEDDEDGDTTID

>C_zanzibari_KNL2

MGDGDVVPARVQNVLDMDIIRLNLWSIQFNGTGVKLEGFVRSEDGNMMQKVHSGIICRRMSATMLFDVSGCFYELAGHIDREYQLKIGMPSRVIDEFMNGFPDNWAFLINSCSEPKSAMRPIQAAPREPLRTRGEPIVTMPDETNIESQSSEKERKRKEREEQQKWEHEERLRARKEKEQREAAEIAEKNRREREIQKQREDEEKGRREEEEEAANYTFRAPKSQNGEAITPIRFTRGNGQKGANIRPIFDSTPVRTKPAGPLASSTPQAPPPQNRHSNTENRQPDALPTAGPAPPVQKPNRETQYCSDADLFAVPRLPAPRNNQNVSSSAQSAPLDFLDEMDALFESATVDKTPGRVKKVVRVNRSPSPDRFSSRDRDSGYDRYDSSRYSHSQRYNDEHHMSRMSGRNDTFRRNDGWRDESRNSRKRGYNNSPEYTRGWDDRSRHRDNYESESRYDSKRSRPRDQSSSSGRSVRFEDDHPGNHRDSRDPRNYRDYEDDRNRESSGDRADRRKLNDILRRERELEARLRNTQRSSTARRHAYTSEDDEMQDEWDRENQEMLDNSMLFGDGISQKKRRSGGRQPAKNSQSGRSQQPKPARKPAQPKKKKERSDETDELNDSIASNRPRRACVTPSTPAPKRITWPKRDLDRLKRTIDLKKPTGADADWDEVTRLLAKDGVDREIVKQTAIMKLKWKEPSQETVQQEEEEKTRRRGAAAMVKEGVRMHQELREGGDNKRGDDLQGGVEAVDDYQPDDVAADQSLLALQTPIAVKKRGGTRASIMPQPVEDSPVVKGNNSTLNSPRLNQTKVKEVETTLKYVQHISMMNARPSSRANTSCLNKSSSRGGGSKNTSLSVEQGARKALKIINRGRTIQEEDEDDDDESEAGDITIE

>C_sinica_KNL2

MGDGDVVPARVQNVLDMEIVRLNLWSIQFNGSGVKLEGFVRSEDGNMMQKVHSGTICRRMSATMLFDVSGCFYELAGHIDREYQLKIGMPSRVIDEFMNGFPDNWAFLIKSCSNPEPKSALRPIKAAPKMPLRSRGEPIVTMPDETVMAESQAAEKDRKRKEHEEQQKRDDEKRFRAEKEKEQREAADTAAKKRQEEEDRRRREEEDANYTLRAPESHNGEPVTPIRFIRGNGQKGANIRPIFEKTPVRTKPTGPLASSTPQAPPPPPTRRTSNIENKQPEASTSPKPQPPVQKPVRETQYCSDTEFFAVPKLPASRNKHVPASDHAAPLAFLDEMDALFNGADVDHTPGRERKPKKVSRSPSPERFDYSSRDRDSGYSRYDSSRYSHNQRYNDDYNMSRMSGRTDTSRRNDGRRDESRMSRKRGYNNSPDEYSRRYDDRSRRQDDYDSGSRYDSKRSRPREQSSSSGRSVRFEEDYSRHRQGSRDSRDPRDYRDYEDHRNRTSSGDREDTRKLNDILRRERELMARLQNSQRSSSTVQRTAYSSDEDEMADESSLWDRENQEMLDNSMMFGDGLSQKKRRGGGRHPAKQAKTRQPAQPKPARKPAQPKKKNRADETDDLNDSIASNRPRRACATPSTPAPKRITWPKRDLDRLRHTIELKKPTGADADWAEVTRLLAKDGVDSEIVKQTAITKLKWKEPSQETIQMEEEEKKRRKGAAARVKEGVRMHEELREGGDKRGDNSLGGVEAVEDYEPDDVAADQSLLALQTPIAVKKRGGTRASIMPQPVEDSPVVRGGNNSTFNSPRLDQTKAKEVETTLKYVQHLSMMNARPSSRANTSYHNKSSSRGGGSKNTSLSVEQGARKALKIINRGRTIHEEEEDDDEEGDTTIN

>C_nigoni_KNL2

MNDLFFVNYGSKFMNDYAQKKKASSHIEDYSLPEKKLRKVQEIVDKSPKCLFAKRAPPHTHFAANFSNLNSEGIKTVFLDIHLDFTISTPYFCYKASFCKKKMGDRNVVPARVQNVLDMDIVRLNLWSIQFVDSGVILEGFVRSEDGNMMQKVRSGMICKRMTATMLFDFSGCFFELSGQIDREYQQKMGMPSRVIDEFTSGFPENWVSLIKSFLPVNPISAVKHIQPAPREPLRPISEPIVTLPDETTLESEKDQKRREKEERLEKKRLEEERLEKERREQKLRAQKEKERREAVAAAEAEKKRLEEEEDAANYTIRVPKSQSGEAITPIRFTRGNGQKGANVRPIFDKTPVREKTAGPLASSTPQAPPPQQRLSNVEKKPASPEPQPRNPQRDREIQYASDSDLFAVPKLPAPKTTKPSTTSEASSGGVLDFLDEMDTLFDTVVVEQTPTRDRRPPRRYSRSPSPRRRQHSSSRDMGKGYDNFESSRYSQRYNDNYNTSRMSRRDDTFRRNDERRDESRMSRKRVYNNSPDDFEYNNRYDDRSRRPDYYDSDSRYDSKRSRPRETSSSSGRSVRFEDDYRRNHRDPRDRSDSKDYRNYDESRRNPEDRREEKRKLNDILRREEELVTRLQNRKKPSASYRREPSSDEDDTADEWDRENQEILDNSMMFGDGLSQKKRRSAGRPSKPSKKERQVQPKPVRKPPQPKKKQKSPPDELNDSIASNRPRRACVTPSTPAPKRIVWPKRDLDRLKHTIGLKKPTGSDADWAEVTRLLAKEGVDAEVVKQVAITRLKWKEPAQDPETIQREEEEEKKRRRGVAARVKEGVKLHEELRQPGVKRGDNSQTGVEAMEDYEPDDVAADQSLLALQTPVGAKRKGGTRASIMPEPVEDSPLVRRNNSTFNSPRLDQTKAKEVETTLKYVQHLSMMNARPSSRANTSYYNKSSSRGGGSKNTSMSLEQGTRKALKIINRGRTIHEEDEDEDDDDDDEDQYEDGVVY

>C_briggsae_KNL2

MGDRNVVPVRVQNVLDMDIVRLNLWSIQFVDSGVILEGFVRSEDGNMMQKVRSGMICRKRSETKKQEERLGKKRFEEERLEKEQREQKLRAEKEKERREAVAAAEAEKKRLEEEEDAANYTLRVPQSQSGEAITPIRFTRGNGQKGANVRPIFDKTPVRGKPVGPLASSTPQAPPPQQRLSNVEKKPASPEPRPRIPQRDRDVQYASDADLFAVPKLPAPKTTKPSTTSEPSTGGGFDFLDEMDELFDTVVVEQTPTRNRRPPGWYSRSPSPRRRQHSPPRDSFESSRYSQRYNDNYNTSRMSRRDDTFRRNDERRDEFRMNRKRVYVCDRGLVRKRKYVFQNNSPDDFEYSNRYDDRSRRPDYYDSDSLYDSKRSRPRETSSSSGRSVRFEDDYRRNHCDPRNRSDSRDYRSYDESRRKPEDHREDKRKLNDILRREQELVSRLQNRKKSSVSYRPEPSSDEDDMADEWDRENQEILDNSMMFGDGLSQKKRRSAGRPSKPSKKERQVQPKPVRKPPQPKKKKQKSPPDELNDSIASNRPRRACVTPSTPAPKRIVWPKRDLDRLKHTIGLKKPAGSDADWAEVTRLLAKEGVDAEVVKQVAITRLKWKEPAQDPETIQREEEEEKKRRRGVAARVKEGVKLHEELRQPGVKRADNSQTGVEAMEDYEPDDVAADQSLLALQTPVGAKRKGGTRASIMPEPVEDSPLVRRNNSTLNSPRLDQTKAKEVETTLKYVQHLSMMNAHPSSRANTSYHNKSSSRGGGSKNTSMSLDQGTRKALKIINRGRTIHEEDEDDEEDDEDQYEDAVVY

>C_remanei_KNL2

KRQEEEDADYTFRAPQSQNGEPITPIRFNRGNGQRQGVTRSVFERTPQRGQSGPLAASTPQAPPPPPPQQRLSNIENREPPPRVVAPPSPVRQPPAPQPPIREPQFANDDDLFAVPKIPPPKIPRGSAGNSGGNIDFLDEMDALFDTVYIDKTPKRDVKPKRPSSPSPERRRYSPMPRDREMGYNDDFESSRRGGRYPSESSNMSRMSRRDETNRRNDGGMGRDESRMSRKRGYDQSPDDMEYQRRREDHYRRPDYYPPRDSRQDSKRYRPRENSSSSGRSASVRFADDYQRNRGDSRDPRDFRDPRDSFSRDPRFYYENNQRGESSKDRDTRKLNNILRQERELVARLQNIKSNTTTNTNTTRRVTYSSEDDEMADEWERENQEIMDNSMMFGDGISKKGRRSGPGRPPQRKPKEQPKPKSQSQPKRRNQNQSKPRRSQYDPVETDDLNDSIASNRPRRACVTPSPVVPKKIIWPKKDLDRLKRTIELKKPTGSEADWTEIARLLAK

>C_latens_KNL2

MPSRVIDEFTNGFPENWGFLIKSCFGNESRSAMRPIQAAPREPLRQRNEPIVTLADETELTNRKNSDNDSESENNRKRREREEEAERERQERYRADEMERERIAAEKKRQEEEDADYTFRAPQSQNGEPITPIRFKRGNGQRQGVTRSVFEKTPQRGQSGPLAASTPQAPPPPPPQQRLSNIENREPPPRVAAPPSPIRQPPAPQPPIREPQFANDDDLFAVPKIPPPKIPRESVGNGGGSIDFLDEMDALFDTVYIDKTPKRDVKPKRPLSPSPERRRYSPMPRDRELGYNDEFESSRRGGRYESSNMSRMSRRDDTYRRNDGGMGRDESRMSRKRGHNYSPDDMEYERRREDHYHRPDYYPDSRYDSKRYRPRENSSSSGRSASVRFADDYQRKRGELRDPRDFEDSFSRDPRFYHESNRKRESSQDREDTRKLNDILRREKELMARLQNNKTTTSKNTTTRRVTYSSEEDEMADEWERENQEIMDNSMMFGDGISKKGRRSGPGRPPQRKPKEQSKPKSQSQPKPRNKNQSKPRRSQHDPVETDDLNDSIASNRPRRACVTPSPVVPKKIIWPKKDLDRLKRTIELTKPTGSEADWTEVARLLAKNGCDGEAVKQTAITKLKWKEPVERDEETIQREEEEEEETKRRRGVNAKVKEGVKLQEELRRGATHQKRAEDVRVEVTAEEIEPDDLAADQSLLAMATPVAVKKRGGTRASIMPMPVEDSPIVRGSRNNSTFNSPTLNQTKAKEVETTLRYVHQLSMVQARPASRNNTSYMNKSSTRGGGSKNTSMSLEQGVKKAMKIINRGTTIHEDDEEEEESESEEEDNSEDLEEH

>C_sp51_KNL2

MGDTDILPTRVQNVLEMEIVRLNLWSIKFNALNIKLEGYVRSEDGNMMQKVCSEIICKRMNSTMLFDVSGCFYELSGQIDREYQSRMGMPSRVIDEFVNGFPENWAFLIKSCLSMGQRSALRPIQDTPREPIRTRVEPIVTLADETELVENERKKAESEKDKKRKEREEDRARENERNLAAQREKERKDTVAAEERKRKKEEEEAHAVAERKRQEEEEAANYTLRAPQSQTGEAITPIRFTRGNGGNGAHVKSIFQKTPVRGKSNGPIASSTPQQTRVLPTLEKPVETTVSKSTKSKSPPPRSIRQTEYASDADLFAVPRNPAPKNNRKPAPTTSSDIGFDFLDQMDSLFDNPDFEQTPTRDRKPRRQFSPSPEPRHRSYSRDRDRYAHLESTRYKQRYTDDYDMSRISRRDVPFRRNDGGRDESRMGRKRGYYGSPDGSEYGRRYDDRDRHMDYYEPDFKYDMKRSRPRETSSSSGRSVRFENDYRRRREDSREPNSSRNHREYDDYRGRTDSEDNRRLEFLEKRENELMARLQNHKKPSSSSSRRLTNSSDDEMADMWEQENQEILDNSMMFGDGISKKKRRSGGGRQPKKQPTRKTQPKPTRKPAEPRKKTQRSPVQRDELNDSIASNRPRRACATPSTPAPKRITWPKRDLDKLKHVIQLKKPTGADADWIEVARLLAKDGVDSELAKQTAVSKLKWVEPVQHEERIDEEEEENKKRRRGAAAKIKEGVRMHEELMSGRQEDDIRGGGEIMEDYQPEDMAADQSLLALGTPLAAKKKGGTRGSIMPKPVEDSPVTRGGNSTYNSPRLDRTKAKEVETTLRYVQHLSNMQARPSSVSNTSTLNKSSGRTSKNTSMSLEQGTRKALKIINRGRTIHEEDEEESGDEESELSEEEEEGESFNY

>C_sp44_KNL2

MGDTDILPARVQNVLDMEIVRLNLWSIKFNASNIKLEGYVRSDDGNMMQKVCSEIICKRMNATMLFDVSGRFFELSGQIDREYQAKMGMPNRVIDEFVNGFPENWAYLINSCLTISQRSALRPIQAKEPIRSKGGPIVTLADETELVENEQRNSDSEKEKKKKEGEQRASENKRNQEAQRERERKEALAAAERERKRHEEDAERKRRKEEEDAANYTFRVPESQLGEPLTPIRFTRGNGGNGANIKPIFEQTPVRGKTTGPLASSTPQAPAPQQPRVLKDLEETVKPTAPQSSKAKPPPQKPQTEYASDSDLFAVPRPPADKNSRTTTSASSSDIGLGFLDQMDLLFDSAAVERTPTRHPKPRRIAVSPPNPRYRSHSKDRNRYGDFNSTRYTQRPSDDYDMGHMSRRDATFRRNDGGRDESRMSRKRGYYGSPDESDYGRRRDDRDRRDNYYEHEYDMKRSRPRETSSSSGKSVRFEDNYRRRREDPRETNYSRSYRDFDDFNTRGTSGDREDNRKLNDILRRENEVKARLQNHHKSSFSRRHVQSSDEESDDMADEWDRENQELMDNSMLFGDGIPKKKRGSDGGRPAKKQPVRKPQPKPTRKPAEPRKRAPRSPVDTDELNDSIASNRPRRACATPSTPAPKRITWPKRDLDKLKHVIQLKKPTEAEADWVEVARLLAKEGVDSELVKQTAISKLKWKPPVQEERNDDEDDEDKRRRRGAAAKIKEGVRMHEELMSGRNPVDDIRSGVEFVDDYQPEDVAADQSLLAMGTPLAAKTKGGTRGSIVPKPVEDSPIIRGRNSNYNSPRLDQTKAKEVETTLRYVQHLSNIQARPSSRANTSTMKKTSSHASKNTSMTLEQGTRKAMKIMNRGRTIREEDEEEDESDREDESEQSGEEEEEDDIYY

>C_sp48_KNL2

MGDTDILPARVQNVLDMEIIRLNLWSIKFNASNIKLEGYVRSEDGNMMQKVCSEMICKRMNATMLFDVSGRFYELSGQIDREFQAKMGMPGRVIDEFVNGFPENWAYLINSCLTVGQRSALRPIQSAPREPIRSRAEPIVTLADETELVENERKNSEAEKEKKKNERDEQRARENERNQEAQRERERKEALAAAAETKRRKEEEEERAEAERKRREEEEAANYTFRAPESQQGEPITPIRFKRGNGANGGYKSIFEKTPVRGKSNGPLASSTPQAPPPQQPRILSSLEREKKTEPDAPKSPIADRTVQKPIRNTEYADDADLFAVPKLPPIRNNRPSAAAPSSEIGYDFLDQMDSLFDTVVIDQTPNRNRMPRRAASSSMEERLRSSPPRNRDRYDDRESTRYSQRYGDDYNTSRMSRRDATFRRNDGGRDESRMSRKRGYYGSPDEFDHRRRDDRDRRGDYYEPDYKYDMKRSRPRENSSSSGRSVRFEDERFGDYRRHREDSRERKYSRNHREYDDNRGRRGSSGEDDRKLNDILRRENELMARLQNHRPSSSRRDASSSGEDEDDLANEWDRENQEILDNSMMFGDGLPKKSRRSAGGKIGRQPKKQPTRNPPPKPARKPEPRKKAPKSPVETDDLNDSIASHRPRRACATPSTPAPKKITWPKRDLDKLKHVIQLKKPTADDADWAEVTRLLAKDGVDSEVVKQTAISKLKWKEPVQEQERKEEEEEEERKRRRGAAAKIKEGVRMHEEMRRGRREEDTQMSADSMEDYQPEDVAADQSLLALGTPLVAKKKGGTRGSIMPKPVEDSPITRGRNSTLNSPKLDQTRVKEVETTLRYVQHLSNMQARPSSRANTSTVSKSGRGSKNTSMSLDQGARKAMKIINRGRTIDEEDEDEESNEEEEDHSEDEAFDY

>C_brenneri_KNL2_paralog1

MGDTDILPARVQNVLDMEIIRLNLWSIQFNASNIKLEGYVRSEDGNMMQKMCSEMICKRMNATMLFDVSGRFYELSGQIDREFQAKMGMPGRVIDEFVNGFPENWAYLINSCLTVGQRSALRPIQNAPREPIRSRAEPIVTLADETELVENERKNSEAEKEKKKKELEEQRAKENERNLEAQRERERKEALAAAERKRREEEEEAANYTLRAPESQPGQPITPIRFRRGNGGNGGHMKPVFEKTPVRGKSNGPLASSTPQAPPPQQPRILSSLEKPKSPIAERSAQKPMRRTEYADDADLFAVPKLPPIRNSRPSASAPSSDMGFDFLDQMDSLFDTVVIDQTPTRNRMPRRAASSSMESRIRSPPRDRERYDDRESTRYSQRYGDDFNMSRVSRRDATFRRNDGGRDESRMSRKRGYVSTPDEFDHRRRDDRDRRGDYYEPDYKYDMKRSRPRENSSSSGRSVRFEDERFGDYRRHREDSREPKYSRNHREYDNYRGGRGSSGEDDRKLNDILRRENELMARLQNHRPSSSRRDAYSSDEDQDDLANEWDRENQEIMDDSMMFGDGLPKKQRRSGGKIGRPPKKQRTRKPPPKPARKPEPRKKAPRSPVETDELNDSIASHRPRRACATPSTPAPKKITWPKRDLDKLKHVIQLKKPTAAEADWAEVARLLAKDGVDSEIVKQAAISKLKWKEPVQEQEKKEEEEEEERKQRRGAAAKIKEGVRMHEEMRKGRRENETQMSAESMEDYQPEDVAADQSLLALGTPLVAKKKGGTRGSIMPKPVEDSPITRGRNSTFNSPRLDQTKVKEVETTLRYVQHLSNMQARPNSRANTSTVSKSGRGSKNTSMSLEQGARKAMKIINRGRTIDEEDEDEESNGEEEEDHSEDEAFDY

>C_brenneri_KNL2_paralog2

IDREFQQNGMPVRSYDEFVNGFPETGLILLILASQEPIRSRAEPIVTLADETELVENERKNSEAEKEKKKKELEEQRAKENERNLEAQRERERKEALAAAERRRRQEEEERADAERKRREEEEEAANYTLRAPESQPGQPITPIRFRRGNGGNGGHMKPVFEKTPVVSEGQ

>C_wallacei_KNL2

MGDNHVLPARVQNVLDMEIIRLNLWSIKFNASNIKLEGYVRSEDGCSMQKVCSEIICKRMNATMLFDVSGRFFELAGQIDREFQLKMGMPSRVIDEFVNGFPENWGFLINSCLTMNQMSVPRPIQAAPREPLRSRNEPIVTLADETELADSGRKTSESERDKKRREREEQRMKENEKRLETEKEKERQKAQAAAAAEKKRQEEEEDAANYTLRGFQMESGDPITPIRFKRGQANGVHAKPTFVQTPVRGKPSGPLASSTPQAPLPEHPRRSLDLGKRSTPLASPSPKVQQPVSKPARETEYASDSELFAVPKLPAPKAARPSATSSAPSDTAFDFLDDMDVFFDTAVIDQTPVRARKQPRRVSPDVRRDRSSSRDPERFGDYDSTRNNQRYNNDYDMSRMSRRDATFRRNDERRDDSRMNRKRGYYNSPDDYDNRRSDYYESDSRYDTKRSRHRETSSSSGRSVRFQDDHRRSENDDYRRREGFRDPEYQRNYRDFEERRVRGSSGDEDKKKLNDILRREKELMARLQNNQKPSSSRRVSCSSDDDIDMADEWDRENQEIMDNSMMFGDGISKQKRRSGGSRQEKKKQPTRTVQPKPKRVASKKKSPVPSDDLNDSIASNRPRRACVTPSTPAPKRITWPKRDLDKLQHVIDLKKPSAEDADWVEVTRLLAKEGVESEVVKQIAISRLKWKEPVKNKNNEKKEEEEDETRRRRGAVARVKEGVRMHEELIRGRDDEERDVQVESMEDYQPEDVAADQSLLALGTPLAAKKKGGTRGSIMPKPVEDSPMARRQNSSFNSPRLDQTKAKEMETTLRYVQQISMMQARPSSRANKSKLNKTSARGSKNNSMSVEQGTRKALKIINRGRTIHEESEEEEEESDDDEFDEREEGDNSIY

>C_tropicalis_KNL2

MEDNHILPARVQNVLDMEIVRLNLWSIKFNSSSIKLEGYVRSEDGCSMQKICSEIICKRMNATMLFDVSGRFFELAGQIDREFQSKMGMPSRIIDEFVNGFPENWGFLINSCLTMNQMSVPRPIQAAPREPLRSRNEPIVTLADETELNDGGRKNSESEKDKKQKEREQQRMKENERKLQAEKDKKDKEAVAAAEKKRQEEEDAANYTLREFQTGSGDPITPIRFKRGTQSKGANNIKPTFLQTPVRGKPNAPLASSTPQAPPPEHPRRSLDLGKPPIQLASPSPKVQKPETVRETQYASDSDLFAVPRLPAAKNNRPPPSSSSETVLDFFDDMDVFFDTAVIEKTPVVARRQRRLSPEARRDRSSSRDPERFGDYDSTRYTQRYNNDYDMSRMSRRDAVFRRNDERRDDSRMSRKRGYYSSPENDFDRRRPEYYESDSRYDTKRPRPRENSSSSGRSVRFHEDHMGRGDEFRGSRYERGYREFDDQRGRGSSADEDKRKLNAILIREKELMARLQNKHKPSASRRVSYSSDEDSDDMAAEWERDNQEIMDNSMLFGDGIMQQKRRSARGHPAKKKQPIRTEQHKPAQKRAPPKKKSPAPRDDLNDSIASNRPRRACATPSTPAPKRITWRKRDLDKLQHVISLKKPTAADVDWVEVTRLLAKEGVEPEVVKQAAITKLKWKEPVRNDENMREEEDESRRRRGVVARVKEGVRMHEELMNGGNEVEEDLQVQSVEDYQPEDVAADQSLLALGTPLLAKKKGGTRGSIIPKPVEDSPIARGKNSSFNSPRLDQTKAKEMETTLKYVQQISMMQARPSFRANKSQMNNSTARGSKNTSMSVEQGTRKALKIINRGRAIQEESEEDEEDDDDDEFEEQEEGDNSIY

>C_doughertyi_KNL2

MGDTNILPSRVQNVLDMEIVRLNLWSINFNASNIKLEGYVRTDDGSMMRKVCSEIICKRMNSTMLFDVSGRFYELAGQIDKEYQLKMGMPSRVIDEFSNGFPENWAFLIKSCLSLPPRSTLRPIQDVPREPLRSRAEQPIVTLADETELVENERKNSDSERDKKRRDREEQRVRENEKRVEDQREKEHQEAIAAAAAEKKRKEEEESAADYTFRVPGTDLGNPITPIRFTRGGQLNGHIKSVFQQTPVRGKSSGPLASSTPQAPPPENPRRLSHLERASVPAAPLPEKDKTPAQRPVRSTDYASDSDLFAVPQLPPAKNIRPSVSSDSAFDFLDQMDSLFDSAVIEQTPGRDQKARKIAPPSPDARRRFSSREPERRSDYESSRYINRYNDDYNMSRMSRRDEPWNRRHDEGRDESRMSRKRGHFNSPPHDSDYGRRDNRGRRPEYYESNSKYDTKRYRPRETSSSSGRSVRFEDDYKRRDDFREPSYSRSYRMYDDRRGRESSRDLEDKNKLDAILRREKELVDRLENTSKPSSSRRHDLYSSDEDPIDLADEWDRENQEIMDNSMMFGDGISKKKRRSGAGRPQKKQPTGSTQPKPARKPAEPRKKARRSPPPADNLNDSIASNRPRRACVTPSTPAPKRITWPKRDLDKLKHVIELKKPTGSDADWAEVARLLAKEGVEAEVVKQAAITKLKWKEPVQIDNKEQEEEEEKKRRRGVAARVKEGVKMREELMSGKQEEDRALRVDSMEDYQPEDVAADQSLLTMGTPLAAKKKGGTRGSIMPKPVEDSPMSRGRNSTFNSPRLDQTKAKEVETTLKYVQHLSMMQGRPSSRANKSYLDKSSRGSKNTSLSLEQGARKALKIINRGTTIHEDDEDDDEEEDDDDEFEDQEVGDTSNY

>C_sp54_KNL2_paralog1

MGDTGIIPFRRKDVLERVEAVLNMEIVRLKLWSIKFNTSVIMLEGFVRSEEGTMIQKVCSENICKRISSTVLFDESGRFFELAGQIDREYQQKMGMPSRMIDEFINGFPENWADLINACVLNQQRSALRPIQAAPKEPLRTRAEPIVTLADETELIGEESKKDMKRREREEQREREQEKKLATEKERKRQEAAAVEKKRREEEEAEEAANFTFRVPQSQSGEAITPIRFKRGNGKKGENTRTIFDKTPVRQNSSGPLASSTPQAPPPQHPRRLSNIENNQPLRSPSQKTQLAASIPERETKFASDADLFAVPKLPPSKTTRPSTTSAVSSEGALDFFDEIDALFDTAIIEQTPTRDRKSRKPIATSPSPDPRRRSLSRDRKSERYGSYESSRYTQRYNDDYDNSRMSRRDVTFKRHEGGRDDSRMSRKRGYYRSPDDFEYSRRRDDREKDYYAYDSRHDTKRSRPRETSSSSGRSVRFEDDYRRRGGEFRESKDSRDFRDYDDHRNRKNSGEHEDKKKLKDILRREKELLARLQKSQKSSSSRRDSYSSEEETFDMADEWERENQEIMDNSMMFGDGLSKKKNRRSGGGGAAKKQLARTSQSKPHRKPAQPRKKTLRDPLETDDLNDSIASHRPRRACVTPSTPAPKKISWPRRDLDRLKHVIELMKPTTTDKYWIEVTRLLAKEGVDPEAVKQIAVAKLKWKEPVQNEEVLKKDEEEEKKRRRGAVARIKEGVKMHEELREGGDKKADDLESGVEAMEDYQPEDVAADQSLLALGTPIAAKKKGGTRASIMPKPVEDSPMAARGSNSTFNSPILDQTKAKEVETTLKYVQHLSMMQARPSSRANKSYLNKSSTRGSRNTSMSLEQGTRKAMKIINRGRTIHEDDEEDDEEDEDDEFDEQDGNTSIY

>C_sp54_KNL2_paralog2

MSEDPDEFLNLRSKVTDPWSVELIGLDAIQQQLDVMGENVAEVILKAEQSFNKKSLPFYEKRKKFTSKIDNFWKKAFLNHHFLSKAIPEEQEDLIEVLRDLEVQEFDDLRSGFKIIMTFDVNEFFENKVITKSYHLKSKPPKTQITEIQWKENQKPHMTSEDGDTTITFLEWLTHAVPPKSDEIAHVIKDDLFINPLQYYMKPDIQENGQPIRRIPKINSSMMEVDDTAEGEDSDEKEEEATNCTYRVPQSQNGEAITPIRFKRGNGKKRENTRTIFDQIPMRGNSSGPLASSTPQAPPPQQPRRFSNIENNRPAGPIPERETQFASDADMFAVSKRAPSKTTSSTSTSAVLFEGEGAYHFLDEVDDFFDTLVIETRERKPQRRRIATSPSPDPRRRRSSSRDRERYGSYESSSRYTRRYNDDDYDDSRMSRRDIATFKRRHEGGRDNSRMSRKREYYRSPDDFEYRRRRNDDRDKDYYNAYDSRHIAKRSRLGETSSSSGRSVRFEDDYRRSKDPRNFRDHRNRTISEKQLLARLQELRKPSTSSHRDSYSSEEEETFNMADEWERENQEIMDNSMMFDDGLSKKKIKRRSGRGAVKKQLSRTSQSKKPHRKPAEPRKKTIRDPLETDDLNDSIASHRPRRSCVTPSTNPVPKKISWPKRDLDRLKHVIELMKPTTSDKYWIEVTRLLAKEGVDSEAVKQIAITKLKWEEHVQNEEMMKMDEEEEKKRRRGAVARIKEGVKMHEELREGGDKKADDLKESGVEAMEDYQPEDVAADQSLLALGTPIAEKKKGGTRASIMPKPVEDSPMAARCSNSTFNSPRLDQTKAKEVETTLKYVQHLSMMKARPSSSKTNKSYLNKSSTRGSRNTSMSLEQGTRRAMKIINRDRTIHEDDEEDEADEDYEFEEQDGDTSIY

>C_inpoinata_KNL2_paralog1

MGDTNILPARVQNVLDMEIIRLNLWSIKFNSSNIKLEGFVKNEDGTAMQKVCSEIICKRMNSVMLFDVSGRFFELVGQIDREYQQKMGMPSRIVDEFLNGFPENWPYLIKSCISVDSRSSLRPIQAAPREPLRSRAEPIITLADETEVVVGDNKNSDTGNGRKHREQVERNSNDQVQMPREANMKNNIQEEEDDIANYTMRAPVSQNGEAITPIRFTRGNGKKGANTRSIFESTPARGTSSGPLQSSIASVPPPQQSRRLSTNQNSQLQSQCQKIQVAPSQNEKPPFREPQFSNDTDLFAVPRLQENKNNRLPSLSAASSDGAHDFFDEMDTLFDTANIEHTPARDRNIIRRSRVASPSPPLRRYRTSSRDCERDIYEPTRHSNRPDNGFNINRNRSDAFRKNEGRNDGSRMSRKRGYFYSPDKYRDNFSRRIKNIDYEQMHDNKHSRERENSPYSGRSIQFEDVYRKRNYGDYDDFRKRSRDEEEDRKLNEILRREKEMMARLQQHRKSVSSFPKSYLSDDDEFDMADQWERENQELLENSMMFGDGLPKKRKMSRDCKSERLKNPKESVKKPVQSQKSNRVKRLETDNANDSIALSRPRRLCATPSTPAPKKIVWPKRDLDRLKRIIELKKPTAANADWAEVARLLAKDGVELEAAKQTAITKLKWKEAVQNEELLNCEEEDKKRRRGLAARVKEGVKMHEEIREGGENKGGDLQNGVESMEEYRPEDMAADQSLLAQKTPIVVKRKGGTRASILPKPVEDSPLARGSNSALNSPKLDQTKAKEVETTLKYVHHLSQLHANPTCRGNKSFANKSLPGGRKNTSMSLEQGTRKAMKIINRGQTFYEDEEDEDDECSENDEDTSMY

>C_inopinata_KNL2_paralog2

MGDTNIVPARVQNVLDMEIVRLNLWSIKFNSSNIKLEGFVKNEDGTAMQKVCSEIICRRMNSVMLFDVSGRFFELAGQIDREHQQKLGMPRRIIDEFLNGFPENWPYLIKSCISVDSRSSLRPIQAAPREPLRSRVEPIITLADETEVVVGDNKNSDTGNGRKHREQVERNSNDQVQMPREANMKNNIQEEEDDIANYTMRAPVSLNGEAITPLRFTRGNRKKGANARSTFESTPAEGTSSGPLLSSITSGPPSQQSRQLSINQNSQLPSQCQKIRVAPSQNKKPPFREPQFSNQDLFDEMDSLFDTANIEHTPVRERSINRSRVASPSPPRRRYRTSSRDCGRDIYEHTRYSSRSDNGFNISGNRSDAFRKKEERNDESRISRKRGYFISPDKYRENFCRRIKNIDNDQMPDNTHSREGENSLYSGRSARFEDVHHKRDYGEYDDFRKQSRDEGKDRKLNEILRREKELMARLEQSRKSASSFQKSYSSDDDELHMTDQWERENQELVENSMLGDDLSKKRKVSRDGKTKQSERLKNHNESVKKPVQSEKRNRVKRLETDNANDCIALSRPRRLCATPSTPAPKKIVWPKRDLDRLKRIIELKKPTAADADWAEVARLLAKNGVELEAAKQAAITKLKWKEDVQNEELLNCEEEDKKRRRGLAARVKEGVKMHEEIREGGGLQNEVESMEEYRPEDQSLLALKAPTVVKRKGGTRASVLPKPVEDSPLARGNNSALKSSKLDQAKAKELETSLKYVHHLSQMHANPSSRGNKSHANKSLPCGKKNTSMPLEQGTRKSMKIINRGRTFCEDEKDDSTDNDEDT

>C_elegans_KNL2

MGDTEIVPLRVQNVLDSEIIRLNLWSMKFNATSFKLEGFVRNEEGTMMQKVCSEFICRRFTSTLLFDVSGRFFDLVGQIDREYQQKMGMPSRIIDEFSNGIPENWADLIYSCMSANQRSALRPIQQAPKEPIRTRTEPIVTLADETELTGGCQKNSENEKERNRREREEQQTKERERRLEEEKQRRDAEAEAERRRKEEEELEEANYTLRAPKSQNGEPITPIRFTRGHDNGGAKKVFIFEQTPVRKQGPIASSTPQQKQRLADGANNQIPPTQKSQDSVQAVQPPPPRPAARNAQFASDADLFAVPKAPPSKSVRNLAASNVDIFADVDSVLDTFHFESTPGRVRKPGRRNVSSPSPEPRHRSSSRDGYEQSRYSQRYEHDNSRWSRHNATYRRHEDESRMSRKRSIVRDDFEYSRRHDDGARRRDYYDADIQGDSKRYRGRDASSSSGRSVRFEEEHRRHGDEYRDPRGPRDYNDYGRRRNHANSRSGEDEEKLNAIVRREKELRNRLQKSQKASSSSYRHRSNSSDAEESLNEWDIENQELLDNSMMFGDGIPKRSNARKDKFVKKQATRSKPANSTKSPAQARKKKRASLEDNRDLNDSIACNRPRRSCVTPVAKKITWRKQDLDRLKRVIALKKPSASDADWTEVLRLLAKEGVVEPEVVRQIAITRLKWVEPEQNEEVLKQVEEVEQKRRRGAVARVKENVKMHEELREGGNHRAEDLQSGVESMEDYQPEDVAADQSLLALRTPIVTKKRGGTRASIMPKPVEDSPMSRGNNSTFNSPRLEQTKAKDIETNFKYVQHLSMMQARPSSRLKKSSSMNNSTYRGNKNTSISLEKGTQKALKIINRGTTIHEDDENEDNDDDDDMREEDTSIY

>C_oiwi_KNL2

MGDVEILPQRVQNVLDMEIVQLKLWSIKFNSTNIRLEGFVRNEEGTMMQKVCSENLVKRMNSTMLFDVSGRFYELSGQIDREYQQKLGMPSRIIEEFANGFPANYAILINSCLNSDQIRSIRRPIEAAPRDPLRPIVTLANETENPKIPEKIPEKSILEEEKEENEADFTLKAPVSTDGNAITPIRFTRGNGRALARTVFEQTPVRGKASTSGPLASSTPQPAPPPIPRRSSTTENRNPPEAPPPKVQYATDSDLFAVPKLPAPRQQAPPISLGFLDDFDDIFADAVIQKTPKRGRPPKREEEDSRRFYEDESRMSRKRGYYRSRSRSPDYDRRHRDRDYSDFYGREDSKRPRRRDESSSSGGGGKSVRFEREDRHQRHHPHRRDRSNSRESQRHHHRRHHDDDYNVSRSRSHEKEEKRRLQELLRKEKELEARLQSFRRPPTSSSSSSEDSDEDEMAGEWERENQEIMDNSMMFGDGISKKRRRQSGEKKRKQPSRPKPKSAAPAPSATRKPAKKNKRSPAETDSLNDSIASHRPRRACATPSTPAPKKVTWPKRDLDRLRRVIELKKPSGREEEWVEVARLLQKEGVNPMEVKEAAIGRLKWKEPVEKEHTEEEEEEEKKRRRGVAARIKEGVKMHEELREGRQQNREDALKNRVEEVEEFQPEDVEADQSLLALTTPVALKKKGGTRASIMPKPVEDSPMSRGNNSTFADSPRFDQTKAKNLETTAKYVHHLSILQGRPSSSAANRTQMNRSTTRGGGSKNTTLSLEQGTRKAMKMISEGRTIHEDSEEDDEDDEDMDDDVFN

>C_kamaaina_KNL2

MGDTEILPLRVQNVLDMEIVRLKLWSIKFNSTNIRLEGFVRNEEGTMMQKVCSENLVKRMNSTMLFDVSGRFYELSGQIDREYQQKLGMPSRIIDEFANGFPENYAILINSCLNSDQIRSIRRPIEAAPREPLRQKQEPIVTLANESEIQNPQKTKIVPSKSPKNSKILEKSIQEEEEEEEDPANFTLKVPISTNGDAITPIRFTRGNGRGMARTVFEQTPVRGKPSSSSGPLASSTPQPPPPPRRSSTTENRNPKTEQAPAPKPPQNPQYATDSDLFAVPKLPAPKQQAPPISLGFLDDIDDIFADAVINKTPRRGRIGRPTSPSKQDVTSDSRGRFYESRRDEMVDESRMSRKRGYYRSRSRSPDYDRRRRDHRDDHRYSDYYERDSRSDSKRPRQREESSSSGKSVRFERREDRHPDRRRHRSTSRESHRHRRRDDDYDVSRSRSRESRKNEKRKLQKIMKREKELMSRLQNSRRHSPSSSSSSSSSEGSDDEDEMAGEWERENQEIMDNSMMFGDGISRNRKRKSEGKKRKQPSRPKQIKSEAPVPAKKSAPKRNKRTPAETDFLNDSIASNRPRRACATPSTPAPKKITWPRRDLDRLKRVIELKKPSGKEEDWVEVARLLAKEDVKPMEVKEVAIGKLKWKEPVEKVQTEEEEEEEKKRRRGAAARIKEGVKMHEELQGGNPLKREDALKNRVEDVEEFQPEDVEADQSLLALTTPVALKKKGGTRASIMPKPVEDSPMSRGNNSTFANSPHFNQTKAKELETNYKYVQHLSMMHARPSSSAANKSQMNRTTTRGGSKNTSMSLEKGARKAMKMISEGRTIQEDSEEDDDDDKEEELEDDVFN

>C_waitukubuli_KNL2

MGDTDILPVRVQNVLDMEIIRLNLWSIKFTATNIKLEGYVRNEEGTMMQKVCSEVICKRMNATMLFDVSGRFFELSGQIDREFQIKHGMPSRIVDEFTNGFPENWAFLISNCLTTEQRSAIRPIQAAPVQPLRLREPIVTLADETELGDRTVVRKENEKDQRKKNREEEEKKHRLLEEKRKKEQEDKIAAGKRKESREEAERKQKEEEDAANYTFRAPPSFGPDAITPIRFTRGTQGRGGIRIFEDTPQRTTSGPIDSSTPKPPPVHEQLPIKVQQKPEVHQQPEPRQTEKAPQTFASDSDLFAVPRLPARAPISSELPTGFDFYDEMDAIFDNAKIDKTPVMESRRKQIPRRFQYEPSPPPILHGQSSSSSQAYEPDFDSRSYYNGRYDHERSRMSRRDDRYVSERDDSRISRKRILYSPDHHDDSDHSRRRAYGSRSRYEDDFPKRSRTRETSSSSGRSVRFEDDYRSNRYHNDREHRDRDRTREQQERDMYESQKLKEIMRREKELEAKLRLKKTATRRVSYSSDDSADDTMNMTEEWERENQEILDNSIMGDRKTSRKRNNAPKREAVKPKKVDKPARAPAKRTKKEPRQAPDLNDSIASNRPRRACATPATPAPKRVTWPKRDLDKLRHVIDLKKPSASIEDWTEVARLLKKDGVEPADVKLIAETKLKWKEPVQDEDVLLQEEEEEKKRRRGVAAKVKEGVRMREEMREGGAKEDELRNRVESLEDYQPDDMDADQSLLALTTPVAAKKKGGTRASIMPQPVEDSPAVKGNNTSSFMNSPKLDSTKVKEVETTLKYVQHLSTMQARPGSSMNKSYMNASSSRGRNASISVEQGTLTTHIVVSIFLRMQMTFHLTPSDTVLFESWKVTDGWTMMGACALVVVAGVFVEAIKRYRKKINQDQMIREQLVYEPFSNRLFASML

>C_panamensis_KNL2

MGDTDILPVRVQNVLDMEIIRLNLWSIKFSATNIKLEGFVRNEEGTMMQKVCSEVICKRMNSTMLFDVSGRFFELSGQIDREFQIKNGMPSRIVDEFMNGFPENWAFLISNCLTTEQRSAIRPIRAAPVQPLRPREPIVTLADETELPGDRTMSKKETERERKNKERDEEERKQALLEENRQRDRERELQKEKEREAVERRKKEEREEAERKRKEDEDAANYTFRAPPSFGSDAITPIRFTRGKKGAVVTRIFDDTPVRSTGQPLASSTPQRSLPVKEQPPQLPKQLKEDSKEQQPPPRQVERPMQSYASDADLFAVPRLPSRAPAPTGLSGGFDVLDEMDSLFDNAKIEKTPVMGSRRKPIPRRYQYESSPPPFLRGQSSSQQAYERDLSSASYYSSRYDDNRSRMSRRDDTSRRYDSGRDESRMSRKRGHYSPPDRRDDYAYNRRREDESRQRDSFRYDEDYNYDSKRSRPREASSSSGRSVRFEDDYCSNRSRENKDRRNHERVRDQRDRNMYESRELKEIMRKEKELEAKLRSKRSSVRRVSYSSDDSDNDTMADEWERENQEMLDNSMIMDRKPSNKRKNVPKKETSKPKRVEKPTRAPAKKNKKDPVETDDLNDSIASNRPRRACVTPSTPAPKRITWPKRDLDKLRHVIELKKPSGNLDDWAEVTRLLKKAGVEPTDVKQIAETKLKWKEPVQNEEALLLEEEEEKKRRRGAAAKVREGVRMHEEMRNGGKKEDDLRSGVESMEDYQPDDMDADQSLLALATPVAAKKKGGTRASIMPQPVEDSPAVKGNNTSSFMNSPKLDSTKVKEVETTLKYVQHLSTLQARPGSSMNKSHMNRSTSRGKNTSISVEQGARKAMKIINRGTTIHEGDEDDEDDDDEVTEDDEENVIY

>C_nouraguensis_KNL2

MGDTDILPVRVQNVLDMEIVRLNLWSIKFTATNIKLEGFVRNDEGTMMQKVCSEVICKRMNATMLFDVSGRFFELAGQIDREFQVKHGMPSRIVDEFINGFPENWAFLISNCLTTEQRSAIRPIQAAPVHSLRPKEPIVTLADETELAGDRTMARKESEKDRRNREREEEKRKKLLEEKRQREEELQRERDREEERSRQEEEEKRRQEEEEAEKLRKEEEANNTFRAPKSFGTDAITPIRFTRGNSKKGLGAIKIFDNTPTPKRKNGGPLASSTPQPQRLPLVKEQSQKEKTPQPEKPPQQQQESSRPAERPKQTFANDADLFAVPRLPTKSTASSGFSSGFDMFDEMDTLFGTAKIDKTPVMESKRKPIPRRYEYESSPPPFRGQSSSSQTYQRDHDSRFNFNDRYDDDRSRISRRDGTFSRYDSGRDESRMSRKRGLYSSPDQRHDYEYSRRKEDESRYRDRSRYENRYDPKRSRARETSSSSGRSVRFEDDYRSKKYREDRREHDRTRDHRERDMYEDRKLKEIMRREKELEAKLHSQRTRRVSSSSDDSADETVNMADEWDRENQEMLDNSMMMDTKTSRKRKNVPKREFVKPKSVEKKPARASAKKNKRDPIETVDLNESIASNRPRRSCVTTAPKRITWPKRDLDKLRHVIDLKKPSADLDDWAEVTRLLKKDGVQPAYVKLIAETKLKWKEPTQDADVLRLEEEEETKRRRGAAAKVKEGVRMRQEIRGGGEKEDDLRRGVEAVEDYQPDDMDADQSLLALATPVAAKKKGGTRASIMPQPVEDSPLVKGNNTSSFMNSPKLDSTKVKEVETTLKYVQHLSTMQARPGSSMNKSSLNRSSSRGKNTSISVEQGTRKAMKIINRGTTIHEDDEEEEEDENSEDDEENAVY

>C_becei_KNL2

MGDTDILPVRVQNVLDMEIVRLNLWSIKFTATNIKLEGFVRNDEGTMMQKVCSEVICKRMNATMLFDVSGRFFELAGQIDREFQIKHGMPSRIVDEFINGFPENWAFLISNCLTTEQRSAIRPIQAAPVQPLRPKEPIVTLADETELAGDRTVVRKETEKDRRNREREEEKRKKLLEEKRQREEEIQREKDREEEKRRQEEEAERKRKEEEEANNTFRAPKSFGTDAITPIRFTRGNAKKGLGAIKIFDNTPTPKRINGGPLASSTPQRPPPVKEKTPQPEKPPQKQQEAPKPAEISKQTFANDADLFAVPRLPTKSTASSGFSSGFDMLDEMDTLFETAKIDKTPVVESRRKPIPRRFEYESSPPPFRGQSSSSQAYQRDLDSRSYFNERYDDDRSRMSRRDGTFSRYDSGRDESRMSRKRGLYSSPDQRHDYDYSRRREDESRDRDRSRYDYRYDPKRSRARETSSSSGRSVRFEDDYRSKKHREDRRDREDRREYDRTRDQRERDMYEDRKLKEIMRRERELEAKLQAQRKSARRVSSSSDDSADETVNMADEWDRENQEMLDNSMMMDTKTTRKRKNVPKKEFVKPKSVAKPARASAKKNMREPLETVDLNESIASNRPRRSCVTAVPKRITWPKRDLDKLRHVIDLKKPSANIDDWVEVTRLLKKDGVQAAYVKLIAETKLKWKEPSQDADVLRLEEQEETKRRRGAAAKVKEGVRMRQEIRGGGEKEDDLRRGVEAVEDYQPDDMDADQSLLALATPVAAKKKGGTRASILPQPVEDSPLVRGNNTSSFMNSPKLDSTKVKEVETTLKYVQHLSTMQARPGSSMNKSSLNRSSSRGKNTSISVEQGTRKAMKIINKGTTIHEDDEEEEDEDDENSGDEEENSIY

>C_yunquensis_KNL2

MGDTDILPVRVQNVLDMEIVRLNLWSIKFTATNIKLEGFVRNEDGTMMQKVCSEVICKRMNATMLFDVSGRFFELAGQIDREYQIKHGMPSRIVDEFMNGFPENWAFLISNCLTTEQRSVIRPIQAAPVQPLRPKEPIVTLADETELGDRTMAKKETEKERRNREREEERKQALLEEKRQRDHELRQEKDREEERKRKEEEDAANNTFRAPKSFGADAITPIRFTRANGKKGHGAFKIFDNTPKRNTNEPLASSTPQRPLPVKEPEQPPQKQPETSRPAEPSRTYASDADLFAVPKLPTKSTGSSGISSGLDFFDEMDTLFETAKIDKTPVMESKRKQIPRRFEAASSPPAFRGQSSSSHASNRDYDSRSYLDDRFNDDRSRMSRRDGTFNRYDSGRDESRMSRKRGHYSPDQRRDYEYSHRRDDESRYRDRSRYEDDYRYDPKRSRAREASSSSGRSVRFEDDYRPNYHREDHRQRDRTRDGNDVYENRKLEEILRRERELEAKLRAKKKPARRVSCSSDDSEDENVDMADEWDRENQEMLDNSMMLNRKTARKQKNVPKRDVAKPKRVEKPIRAPIKKKTRDPVETDDLNDSIASNRPRRACVTPSTPAPKKITWPKRDLDKLRHVIDLKKPSANIEDWVEVTRLLKKVGVEPADVKGIAETKLKWKEPTQNAEVLRLEEEEETKRRRGVAAKIKEGVRMREEMRNGGEKEDDLRSGVEALEDYQPDDMDADQSLLAFATPVAATKKKGGTRASIMPQPVEDSPLVNGNNTSSFMNSPKLDSTKVKEVETTLKYVQHLSTMQARPGSSMNKSYMNNSSSRGKNTSISVEQGTRKAMKIINRGTTIHEDDEDEEEDDDEVSEDDEDNAIY

>C_macrosperma_KNL2

MEYIVMGDTILPVRVQNVIDMEIVRLNVWSIKFTATNIRLEGFVRNEEGTMLQKVCSENICKRMNATMLFDVSGKFYELAGQIDREFQIKHGMPGRVVDEFLNGFPENWAFLIGTCLTTEQRSVLRPIQAAPVQPLRPKEPIVTLADETELTGDRTMAKMETEKERRAKQREEQRKLEKEREEEAVAERKDKEKREREEAERKRKEEEDAANYTIRAPNPLGNEAITPIRFHRGKGGGPRFFNVFDKTPKATSSAPLASSTPQARPPVRKEPEQPVSSKPVERPVAFASDADLFAVPRLPAKGPTTSSSSAGAPLQDDFDFLDEMDTLFDTAKIEKTPVMDTRRKPIIRRFEHRSYSPQHGQSSSSQAYEMDYNDRREPYYHDRYEDDRSRVSRRDFGGDASRMSRKRGYYQSPENRDRDEYEDRRRREHESRYSQQIYSREFEMNGWPRRETSSSSGRSVRFDDDYHYSDRHRNPRDKTRDERDREMYERQKLKEITRREKELEAKLKSRKVSRSSEESADDSMDMAEEWERENQEMLDNSMMMMPDRKSSRKQKVVPKREKPKKAAEKKPAARAPRKKNIREPLLEEEDDPNDSIASSRPRRACATPSTLAPKRITWAKRDLDKLNHVIELKKPTANLDDWVEVTRLLKKNGVEAADVKHAAETRLKWKEPVQNDERAQEDEIEEERKRRRGVAAKVKEGVRMREEMREGGEREDEMETRVGAVEDYQPDDMDADQSLMALGTPAVKKKGGTRASILPQPVEDSPIVRGNNTSSFMNSPRLDQTKAKEVETTLKYVQHLSTMQARPGSSMNKSHANKSYMNKSTSRGGKNTSISVEQGTRRAMKIVNRGTTIEEDEEEDEEENDDDEEDTSVY

>C_sulstoni_KNL2

MEENGILPTRLQNLADMDVIPLELWSIKFYGTRYKVEGIVRPEDFTTAQKFTSEFIIKRITSTALVDASERFIELVGQIDRTFQIRNGMPEEIVGKFQNGFPRDWWELLKPCLAAGQKTSTMRAIEAPPLKTSTTSSTVQNVQKTSSEPIVTFADETEVQSDREKERRHRQEEREVQKVLEKDDDDDADDADYTIRQPRAIDGGENTPLRYKRGHSGGPQNRNRVFADTPQRTAIPLAASTPNAPPKIAKPPVKFASDADLFAVPVPPTRRPAERKERSESPEGCDGLFDKISEVFSNVNFPVTPLPAARVRAHFSPAERSEGSYYYRDDRYDEDRNRTSRRDRSRSREYDESRTSRKRGYYRSPEGYEYSRRRDDRRRDRYEASSQRHGSSSSGRSVRFEDPYEYDVEQRRRRREDQKLREIERREREIEARLERSKRSRRYSSDEQSEDMADEWERENRQILDDSMLFRSAAGKATNKKRAGRPKKEVKTRSPPVQKPQKARKEPRKRRSEPAETVDLNESIASNRPRRACVTPVAPPPPKRVTWPKRDLDKLRRVIEVKKPTGDEHDWVEVARLLAKDGVTPTDVRTIAETRLKWREPTRDPKVLEAAEEEETKRRNGLAARVKESIRMREEMREGRQEEDVELKPGGENYEPADVSADESLLAIGTPRVPPKKKGGTRASLIAPPVEDSPMTRRSGAAATVVASPKLDATEVAATQKHVHHLSMIQARPGSSMMNTSRANKSIRNGKNTSISVEQGARRAMKMVTRGRSTIAEEDESEEDDDENEEDVTAEEDEIY

>C_afra_KNL2

MEENAILPTRLQNLAEMDVIPLELWSIKFYGTGYKLEGIVRPEDFTSAQKFSSELIIKRITQTALIDASERFIELVGQIDRTFQLRNGMPEEIVNKFQNGFPKNWYELLKPWASQNQKTSTWRAIEAPPVAPPNPQIPPIPQKAPCEPIVTFDDETEVQSDKEKEEARRRRQLEVEEAEKQRKILEDLNDDADYTLRNPRSIDGVQNTPLRFKRGNPNFERNPRVFNDTPVRTGGGPPLAASTPQAPKVSDPIPKAANYASEGDLFSFAVPTQPPSQRPTRTTDEDRKHQADRRRRSESLGGSDDGGGGSGLDLFSQMSEVFSNAKFPITPLPAAPPRSSRFERERAERSDYYREDRDRERYRDPSRESRERDESRMSRKRGYYRSPEGYEYSRRREDRHRDRYESSYQRYGESSSSGRSVRFDDGYDDPYEREEEERRLRRHRDNQKLREIMRREREVEARLERSKRSRRYSSEDDGPEENDDDLADEWDRENREILDNSMSFARGGSGRAGKKRGAGRPKKTSAPKSRSPPTQQKPRKQQQAAKKRRSEPAETDDLNDSIASHRPRRACTTPKAAAPKRVTWPKRDLDKLRRIIEVKKPTGDADDWVEVARLLAKDGVEPADVKTCAETRLKWKEPTRDPEVRQKEEEEEKKRRNGLAARVKQGIRMREELREGYEEDVAPPPTSSSQEPDYEPDDVSADESLLAMGTPLPPPKKKGGTRGSIVVLPVEDSPMVRRSGVAASPKFDASDLAATQRHVQHLSMIQARPGSSMNTSRASKSILGGKNTSISIEQGARRAMKMMNRGRVAIAEEDEDSDSDDSDDDENDSNDNAEYIKIKVVGQDSNEVHFRVKFGTSMAKLKKSYADRTGVSVNSLRFLFDGRRINDEDTPKSLEMEDDDVIEVYQEQLGGGI

>C_afra_KNL2

MEENAILPTRLQNLAEMDVIPLELWSIKFHGTGYKLEGIVRSEDFTSVRKFSSELIIKRITQTALIDASERFIELVGQIDRTFQLRNGMPEEIVIKFQNGFPKEWYELLKPWASQNQKTSTWRAIEKPPVVPPNPQIPPLTQKTPCEPIVTFDDETELRSDKEKEEARRRRRLEVEEAERQRKLLEEGSDGAEKRGYNRSPAEKNNRERSERSRRCSSEDNRSDGELADEWERVNSEILDNSMSRVAGRPKKMLAFGRALTQADVEKFAIRLEIAGARIPAVVASEKYVVTTLKNIPEASRKLGFALSAFDHLEQRIELKIHGMHRHSDFVVLRRDEGEFSEAPVYRNEEVGDKYFVVKRSIDGKKIVCHGKVTSVGATGFAFVSEPYEGFPVANGDGAFAFLDGKLLGVVKDTKEQGRVLEFVSIYLVHFAIDIIFHHQPFSV

>C_sp49_KNL2

MTERTIVPVRVQNVMDMEILRMNVWSIKFNSSSVMVSGFVRTPDGNSMNLVTSEAICKRMNSTLLFDISGRFYELLGQIDKSTQLKAGMSSRVMEEFLNGFPDNWVFLVGQCLTTEQRSAIRPIQAAPREPIRPKSPIVTLADETELREQGKAQKEQKEDADATFRAPTAFFHGAVTPIRFTRGNGATGRSQRVFESTPSREPSSSRPLASSTPQPQKLPVLPPVAPRDVPAAQPRTFATDDDLFAVPKLPASSSGLDFLEDMEAIFNGAHINRTPSHAAPRARVPREFRSRSPSYEQRYQDRRYQDESRVSRKRGMDRSPEYYEMERRAEESRRSHESKRARPRETSSSSGRSVRFEETYRRDRHHQNDRSHRSHHYEQEKEHKRRLREIERREREVEERLKRNRKVSSDSNSSSGEEDGSFNEWDVENREILDRSLMEASGRERRPKREPTSKKSTSKKPAAPKPKPTKNIRKKRDPIETDDPLNTSIASSRPRRACATPATKKITWPRRDLDRLKRVLEMKAPTGRSEDWVEVARLLAKDGVNSEEVRQIAESKLRWKEPEHHPEEHQEDPLNDSQLIEKKRRNGIAARVKESVKMAEELQTRPPNSDEMRGRVEEVEEYQPADVSADESLLALSTPQVIKKRGGTRKSTLPEPIEDSPMARGANSSVNSPRLNATKAKEQETTLRYVQHLTTMQTRPASRMNKSAMNKSTASAKNTSMTVEQGTRHVQKMMKKAMAIDEEEEDDEESDENQEETSFY

>C_sp25_KNL2

MTEQTVVPVRVQNVMDMEILRMNVWSVKFNSSSVMVVGFVRTPDGNSMNLVTSEAICKRMNQTLLFDVSGRFYELLGQIDKDTQLKQGMTSRVMEEFINGFPENWVFLVGQCLTTEQRSAIRPIQAAPRQPIRPKSPIVTLADETEVAAQAQAQREQKEQKENEQRRRREDKEKEDADATFRAPASFANGAITPIRFTRGASVASRNQRVFDATPTRQAPLASSTPQQKVQKVPTIAADEDLFAVPKLPVKKGVAPASDLDFLDDMDALFTGARINRTPSHRVPPKPQQRRRSRTRSRSPSIDRSRSRRYEEEFDSRMSRKRGYDRSPDYDDYRREESRRRFKESKRSRPRETSSSSGRSVRFESDYHRQRHDDRRRHEDREELERQERRRLKEIERREREVEERLKRSRRKNYSSSSSSDEDQDGSFNEWDAENREILDQSLLASTGGSSRSASGGNSRGRLPKREPSLKKAKPAAAAPKPKPRKKKEVVDLNDSIASNRPRRACATPATPAVKKITWPRRDLDKLKRVIELKKPSASEEHWTEVTRLLAKEGVAWTDVKSIAETKLKWKEPRQEMVEVEEEEEEERERKRRRGIAAMVKESVKMCEEIQTRNVQEDEMEGRVEEVEEYQPADVSADESLLAMSTPQITKKRGGTRKSTLPEPIEDSPMAKGTNSSWNSPRLNATKAKEQETTLRYVQHLTTMQTRPGSSMNKSYMNKSTASTRGKNTSLSVEEGTRKVQKMMKKAMAIEEEDEEEEDSDDEGETSLY

>C_imperialis_KNL2

MTESTVVPVRVQNVMDMEILQMNVWSIKFNSSSVMVVGFVRTPDGNSMNLVTSEAICKRMNSTLLFDVSGRFYELLGQIDKTSQLKQGMSARVLEEFLNGFPENWVFLVGQCLTSEQRSAIRPIQAAPRQPLRQKSPIVTLADETDLGDRHKREEVENEARKREERKMIERENEEKRRAEERRQREEQEKEDANCTLRAPASFASGDLTPIRFTRGNGPCNRKNRVFEDTPSRETTSRPLAASTPQQKPKPVDPPREVPRDVPARPKTNFATDDDLFAVPKLPANRAAKPSSDLDFLDDMDALFAGAHINRTPSHAPKPVSRIRQSPPRRRSPSENRSRHEYSQRSNRYDDDYDVRFNRNGKEESRLSRKRGYYRSPDEYYETRRDWGRDRSRDRSRQRYDSERYDSRYDSQVSRPRETSSSSGRSVRFEDDYSQRNNRRYEHEVRERKQLKDIERREREVQERLQRSKKSAPARQATYSSSEDDDDASFNEWDAENREILDQSLLASTAGTSRRRSAPKREPKSKKPKPAPAPAPKRKPSKPRKQRDPVETDDLNDSIASNRPRRACATPATPAPKKITWPRRDLDRLKRVIEMKKPTASDGAWAEVARLLAKDGVTPADVKQIAETKLKWKEPTQDPVEIEEQEEVELKRRRGIAAKVKESVAMREEMQRRAVDEDELNGRVEEVEDYQPADVSADESLLAMGTPQVTKKRGGTRKSTLPEPIEDSPMARGNNSSFMNSPRLDATKAKEQETTLRYVQHLSTMQNRPGSSMNKSQLNKSSTSRGKNTSMSVEQGTRKAQKMIRKAMAIEEEDEEDDEDDEDENREEEETSFY

>C_japonica_KNL2

MAETGVLPLCVQNVMDMEIVRLNLWSIKFYAESIKLEGFVRNSDGNMMQKVCSEAICKRMTSVLLFDVSGRFFELVGQIDHEFQVKNGMPSRIVAEFMNGFPENWLFLIKQCLPPTDEQIPQRQIQIQSAPAPPKEPIVTLADESEHVSDHPKKIVENGRIRDEESADCTFRVPHPVAIDAITPIRFTRGVRKENAQRIFDSSTPKPQVEKQKKPLAAFDADLFAAPAPARAPPPTRAPPPNPPGPSSEFVLLDEMDELFNNAEFDKTPGRTVAQPRRINETSRMDRKTSRTYSEDCDRKYDDRRVECYSRSARENVEPNRNEFRMNRKRPSVNNYADEMNGGYNYRSNKYGRREVFESNGYHSDEFRSREAPSSRSYADRDYYSRKSYHRGEEQRNEKMYGDILRREMALETRLKNGRSQRHVSESSEDSMDDSYMVDEWDRENQRILDSSIVSSFGRFETSPKKSRIYVKKERTAKKARPKPVPPRRKKNSPDSPEMDDLNDSIASNRPRRACVVPSTPVPKRVTWPKRDLDRLSRVIELKKPSSSDSDWTEVARLLAKEGVEGADAMKIAMAKLKWKEPVKDCETVQEDLEMVDKKRKRGFAAKVKESVQMMEEMRDGGEQSGEMGGQGVEQAEDYQPADLSADESLLALATPCAGKKKGGTRRSVMPAPVEDSPVIKLNECSYANSPKLDVTKCKEVETTLRYVHHLSTVQARPGSSSSRSYKNRSVCLGKSLSLEQGARKAQKLISMAVGIEEEDEEEEDDDDAELSIF

>C_tribulationis_KNL1

MENNPRPKRNSILKVRQEKNLLEMLDETKAAAAPENATRRVSFHQMKHVKEYDRHLGKIIDATPTKEKAFDTISSDGTATPHVTRTDMDITGETTTTSRLFENAQATPTSSAHFGHHHHHQQQHDGTMDMSMDTTIENHRPIVQNEGSADETARLFDVTREKTVMYEEVTKTETQTTQTRTLISVYHERAAPDDTMALFDMTNRDEVDMSMDGGGVAGVAAARVDDTMAVFNTTNTEQIDMDITQDQKAPSVPLDDTMAVFRSPAPPRLQKASGIQKTSNENSESERVDMDITRNSSVLAPTDSDDPMDITGIQKAPEDVEPGSEAMDITMAAPNDTMDVFATPKRIQKTSGHQKTPREEDDDDDAMELETPMKVKTGSNETFDLLQSPARAKIQNPSHQILVVEEEEEHVDMDITQREAVEPSDDTMAVFRSPAKTSHETTRQVFDNESMDIESTLRPEDVVPEESPEDVKRSDSHTMMSMATETSIVVVEENRCVSTMLQMSPIGRSEDVEIQKMSTTTTTMASDSQMSLSSSIHNSSVTLNVSKTGDVGEEDGGASERSQIQTMVQEDADVTNPLQNLTTKEEETEADSEDVTMKEESRISAMNVSSVSRRRRSLLMDPLCRESPRRLALGNSMLSMSTAAATTLQGGGGALAEYRHSKRMLNESVGSVGSAGGNQTLANLTATGRDIFQLNTSVRSPAHRPIVTHHHHLPNTTVTSPDATLQQQQHVVPPKSPESFRLPQFDAAIVNVIYLTPEDDAQEPIPEAFEFQKLLAQQEAYAHREIEEAVNGDETLQKVVDEIGGDVVKIRGHLERDVIRIAIGQAEKKFLELRGEFAEEHVAKSDALVVELEAENARLAQRVQDARNVETLRAELHELQRRPTREEARRIRREHHEAKMESMRMQLASTRARYEAEMEMRAQRQRVEDALEEKREQIARLEDAREKIKTEMAGRVRAIVPEE

>C_sp41_KNL1_paralog1

MENNRKQKRNSILKVRQETNLIDETTVVAPNANRRVSFHQVKHVKEYDRDHGKIIDATPIKEKAFDTMSSDGASTAHTTRIDMDVTGDTTMPSRLFEGGGSAHFAHHNGHNGTMDMSMDQDHTMSLENNQNETARLFDVTREKTMVYEKKTTTTRTTTRITVVGGGSSSCSSSGVNDTMAVFNMTNRDGVEMEIEGGGGRVDDTMTVFNTTNTDQIDMDITQQQQQQEQREEQKALDDTMAVFRSPAPMRGIQKTDIQKTSRRVAENSDMTKIADDTLSLFKSPQKKEDLMDITGIQKTSSSEEGAEPGSEAMDITVAPPDDTMEVFATPKRLQKAQNAEDMMLLETNETFDLLQSPARAKIQGGKQHLEHQEHHDDMDITQKEVEPADCTMALFRSPEKIPAQNQTRQVFDDDDENMDIESTLLRPSEDTPEDVKPSESSIMTMSMATEASMVVVEEERCTVSTVLQMSPIEDVEIQNPLQNPLSMTSSSEMSLSSSIHNSSVTLNVSKTAESPLSSQIQTMILETDSTNTIQNLTTTPIKEDSEDSEDVTMKEDESRVERVQEDSVMDVSKETNESRISNVSRRRRSLLLDPSMCRESPRRLALENSMLSTVGAYPGAGGAGGGALAEYRQAKLNESVESSGGAGNRTLLANLTATGRDIFKMNTSIRSPAHRAAATTMISTPEDTSSHIMSHAPKSQPPESASFHLPQFDAAIVNVIYLTPEDVVAAQEEPIPEAFEFQKVLAQEEANAHRDIEKAVNGEESLQKMVDEIGGDVVRMMGHLERDVIGIARGQAEQKFLEIRGRFAQEQVLKSNERIIELESENSKLAQRIQDARNVNVLRAELQELQSRPTKEEASRIRKEHHELKMESMRMHMASIRARYELEVELREQRKKVEDALEEKRETLARLEEQEEKRREEMTTRVRAIVPGLQ

>C_sp41_KNL1_paralog2

MPENLNHSILKSPSAPDSHDENRHVHFDKMKEVQVYDRNTGSWMGQTPIKEKAFDTLSSGGILTPQLSDMDISQSPESSSVSSINKSSYNFETSVNDTLSIFEVTDEPAPEEFKFPEFDKNVANPTFLCPKIAQEFEKLLVAELTETQSEIQRLFEKSDGEYRKKLEFLKNSPAGEWAKTEQDVIVISSQLAESLLLDLRLKFFAEKNEKCEKEIEVLKKENSKISGSIEQKKAKRKNEDFHTIQDDYYKMKKLEMKCRLDSVMKSYEEQLECEKERITLEMEIEELKVRLEFVAEKEKEAKESLEKLITSIDY

>C_zanzibari_KNL1

MENNTNTRQKRNSILKVRQEVNLIDETTVVTAPNANRRVSFHQVKHVKEYDRDHGKIIDATPVKEKAFDTMSSDGASTAHTTRVDMDVTGESQMASHVFESAPVTPGPLEMSLENQNLDETARLFDVTREKTLVYTETTKTKTTVTRITTTTEAAADDTMALFNMTNRDGVDMSIANHDDTINVFNTTNTDRIDMDITQQKGPLDDTMALFRSPMPPPPPPPSSRILQNPLVENSESIDMDITTPQMAVVKDSEDMDITEAQNPEVEEEEEEPGSEAMDITVASAAARGGSAASDDTMSVFLATPKKIQMTSEIQKNHEDMEMTLKNETLDLFQSPAVAAVARGGKTMLQEQEVDMDITQREGGEPSDNTISVFRSPEKIPTRPVFNDNDSMDISSTFRPESEVAVEENRCVSTVLQMSAMEMASPEVSIQNSLPTTTSSATSHSSSINNSSVTLNVSKNDGDAGASERSQHATMMMDVTNPVLEEDSEDVTMTKEDSRRSLNVSSVSRRRRSLLMDSARESPRRLALENSMLSMTAVVGGGAGGGGGSEALAEYRQNKTMLNDSVASGGNQTFGNVTATGRDIFKMNTSIRSPAHHRPTTTIVTSPESIPPPKSPESFRLPQFDAAIVNVIYLTPEDVTQEPIPEAFEFQKLLAQHEADAHRDIEKSVAGDESLQKMLDELSEDVVKMRGGHLERDVIGIARGLAEQRFLEIREGFAEEQVAKSEAKIVELEAENARIAQRIQDARNVDLLRAEIHELQRQPTREEASRIRREHHEVKMEALRMQVASIRSRYDAELKLREQQKRMQDAMEEQRERIARLEEQEAMRKEEMVKRVRAIVPGVL

>C_sinica_KNL1

MDDKNKKRNSILKQRQEVNLIDETVVTSSNNGNRRVSFHQVKHVKQYDRDHGKIIDATPVKEKAFDTLSSDGTSTAHTTRIDMDVTGESAPATPRFPSHHHNGTMDMSMDFSTIENQNRDETARLFDVTREKTTVVYEETTVEKTTKMTKIVTTTNDTMALFNVTNRDDVDMSVAADDTMTVFNSTNAGAVDMDITMPQQVQNPKNPNLLDDTMAVFRSPAPPRIQKTSSGIQNPQNQILENSESVDMDITRNQNPIDNTMSVFDTPKKDSKSKDSESKNSEDVKPSDDVTKDSEDMDITEEIQKASEDVEEEAEPGSEAMDITMTPKKIQKASDDTMQVFETPKKIPGIQKTLTSSSGVLDNLEPVDMEMTLKNETFDLLQSPARIQKTSGRILENLEIQKTSGGILEQRDDMDMDITQREPMQPGDDTMAVFRSPEKIPTKNTTRQVFTDDDDDDNMEIETTLRVSEEPSEDVKPSESSMMMSMETTIIEENRCAIVGSAGNLEISPFGAFGAQEEEPEDDMEIEKTSMMTMTMTSEVQNLNPPTSSASESQLSPEKTPLKSEQNRSPIATMMLQKSPQDSEDAPEDVTMVKEQKEESRMSQDSENVTKDSEDAPEDVTIPDDSRISLNVSSVSRRRRSLLLLDPSLCRESPRRLAMENSLLSMTAIQGAGAASEALAEYRMNKTLQMHQASFNETSGGGGNQTTLNNVSATGRDIFMMNTSVRSPAHRPITITNTITSPESHIANVTAPKSPESIQILLPEFDAAIFNVIYLTPEDLFEEPIPEAFEFQKVLAQHEALAHQEIQKSVDGDERLQKIFDELNSNSEDSGANLEHFEPKIAKNFGLLERDVIGIAHGLAEQKFLEIREKFALEQVGKSEAAIAELEAENAKMAQRIQNCKNADVIRAELQELRAQPTREECSRIRREYHEMRMEKMRIQQVQLRRRYELEVAYREQREKMECVAKEQRERIARIEDAEKERKAEMARRVRAMVQGV

>C_nigoni_KNL1_paralog1

MDNQRKKRNSILKVRQETNLMDVLEDTTVATSSSATNRRVSFHQLKQVKNYDRAGGQIIDATPIKEKAYDTMSSDGNSTSHTTRLDMDITGLGTPVTPKTPAPFNGSMDMSMENQDETARLFDITRDKTICVYEKTVETTTKIVERVVRVPEGSSGANDDTLALFNETNRAEVDMSIDGALKVDDTISVFNQTNVEPVDMDITVQKPLDDTMGVFRSPAIPASSRIQKTFSDSQDMSMDMDITSNETMAAFKSPKIMSSGLVDAVKDADGMDLTGLVNTRAEDVADDTMAVFRTPTRAQTTIQKTSGEISESVDMEMTGIENSDVPDDTMAVFKTPTRAQMTIQKTSGEIPESVDMEMTLQNETLALVQPVSAVKDNSDDVAMDITQQTLVGASDDTMAVFKSPAAEKKTPAKPLFDDSMEIESTIVRPDDVTSSEIAQPKNPENHSSMMMMSMASEVSEDVVVQKTSESLQQSSMMMSTTITEDVTASKNPESSMISEVQKIPEVVQETSNAVEDIQKAPETPEDVSMEITSEVVEGERGTMYQMSMMDMDSLQKTSLASPKIQMTSFTDTSEKMGGASETSMIQTMMIEASESMECSEAPEDVTVSPEGLTMAPEDVTVASNVSRIVSVSSISRRRRSQLQESLHRESPRRMALEKNLSMMSQMGGASEALADFRQNKLNNQTTLLNDSVNTTIGANTSGSIGRDIFKMNTSIRSPAHRSSTAPSSMVSKTLPESPKFHVTPFDAAIVNVIYLTPEDAETQEPIPEAFEFEKVLSAEESNVHKEIDTANWSISGAIKSNLDAEVMNIARGQAEMKFLELRGEFAKERNVEIAQKNQELESQNLELAEKMRDSQNLPELQKQIEELQKPQFSLEEAERIENEYHETKVALLRAQAASIRRQHELQMAIREERRRLIQEIEEKDELLARLEEEDRKKREEMVERVRDVMRA

>C_nigoni_KNL1_paralog2

MSSEHNHSILKNPDESFPHDENRHVHFDKMKEVQIYDPTTGSMTGATPTKEKAFETVSSEGTPSSYMDVVEDSDDSLNKTKNVSHESIQEETLDIFDISEEFKLPTFDPEVVKTIFSSEEEFQNLLKSKMAESQKEMEKTYEESDGEYHDRLKFLKSNSISELTKIEQKVIEVAYKQAEIRFLELRLKFAAEQRVKSEETIVEMQKVQKSRDSIPEMFDRIQDEYFEAKKEIMRIRLEKIQKSYEEQLDSERKRFQLRLEIEERRKKLERLEMKENESKERMEQLISEMEV

>C_briggsae_KNL1_paralog1

MDNQRKKRNSILKVRQETNLMDVLEDTTVATSSGATNRRVSFHQLKQVKNYDRAGGQIIDATPIKEKAYDTMSSDGNSTSHTTRLDMDITGLNSTPVTPKTQTPFNGSMDMSVENYDETARLFDITRDKTICVYEKTVETTTTKVVERVVRVPEGSSGANNDTLALFNMTDRAEVDMSVDGGSEGALKVDDTISVFNQTNVEPVDMDITVQKPLDDTMGVFRSPAIPTSSRIQKTSASTMDSQNMSMSMDMDITSNEMMAAFKSPKIMSSVLVAAVKDSDDMDLTGLVNTAAEDVADDTMAVFRTPTRAQTTIQKTSGEIPESVDMEMTGIGNSDAPDDTMAVFRTPTRAQQTVQKTSGEIPESVDMEMTLLALLQPVSDVKDNFDDVAMDITQQTLVGVSDDTMAVFKNPAAEKKTPGKPLFDESMEIESTIVCPDNVTFSETAQPENPAYHSSMLMSMASEVSEDVVVQKTSESLQQSSMRMSTTITEDVTASKNPESSTISEVQKIPEVVQKTSNGVEDIQNASETPEDVSMEITSEVVEGERGTMYQMSTMDVDSLQKTSLASPKIQMTSFTDTSEKMGGVSETSLIQTMMIEASESMECSEAPEDVTASPEGLTMAPEDVTVASNVSGIVSVSSISRRRRSQLQESLHRESPRRMALEKNLSMMSQMGGASEALAEFRQNKLKNQTTLLNDSVNTTIGANTSESIGRDIFKMNTSIRSPAHRSSTAPSPMVSKTLPESPKFHVTPFDAAIVNVIYLTPEDAETQEPIPEAFEFEKVLSAEESNVHKEIDTANWSISGAIKSNLDAEVMNIARGQAEMKFLELRGKFAKESNVEIAQKIQELESQNLELAGKIRDSQNLRVLQKQIEELQKPQFSLEEAERIENEYHETKVELLRAQAASIRRQHELLMTIREERRRLIHEIEEKDELLARLEEEDRKKKEEMVERVRGVMRA

>C_briggsae_KNL1_paralog2

MFSEHNHSILKNPDDSFPHDENRHVHFDKMKEVQIYDPTTGSMIAATPTKEKAFETVSSEGTPSSYMDVVEDSDDSLNKTKNVSHESIQEETLDIFDISEEFKLPTFDPEVVKTIFFAEEDFQNLLKSEMAASQKEMEKSYAESDGAYHDRLKFLKSNSISELTKIEQKVIQVAYKKAEIRFLELRLKFAAEQRVKSEKTIFELQKVQKSRNSIPEMFDRVQDEYFEAKKEIMRIRLEKIQKSYKEQLDSERKRFQLRLEIEERRKKLERLEMKENESKERMEQLISEMEKINIGHLINHRQNQLSQQNMKASKVFITFDDISKVEHPFQVIVVVIRIGGCCHA

>C_remanei_KNL1

MEGNRRRSILRQPRAMSEEITDENVVEGAQTAIVTNRRVSFHATKQVKEYDREYGKIINGTPVREKAFDTMSSDGGAMTPRVASVDMDVSESSTSTTPFKIFDTNRQDTDGSLMDMSLDRTLNNETARLFDITREKTLVYEKTTEVTTRTTERILTVPTSGADGGGGQDDTMALFNQTDRQEVDMSVDQGGANNETLNIFSTTNRDVTDMDITSDHRVAPQFKTPRLPADAKKAPITDNMEPIDMDMDTTLNAANDTMAVFKSPARIKRNDSEPIDMDVTRDQLNNETLALFQTPENHRKNPMSVLQKTPEIGSEAMDISMAAGMTPKPAPAPVDMELSSDDTMALFKITTPVTKKNFGGEEDMDITQRPASVADDTMALFKSPARVEAAPTAVQQQQLFEESMEMEDQSALSEKVAEPEDVATSETPEAPSNHSSMMMETETSIVEEDRAAVQMSMMDVTAPLDDVMESMLLNQTNSEKTLTTTSTTEESPRPKSSVRDVTSSVHVSSVTLNVSTNRKENDSMIRTMQYTEVDTTNTLQNTSQRMEDSDEEEIKEESTAKETTLIREESEDMTIQNPQEHQQEMSISSYNLSVFNQTTNSTGSVPISRRRRSLLREVCESQRRHAMEKSMNISTAGGETALEEYRKEKMNASGVMNQSLDQSAQGRDIFTMNTSIRSPNVRLAQGTPPPKSPRFEMPLYDPAVVNIVYLTPENVNNPTPLPEAVEFQKVLAEESQRVQKEIQRKQKESGVPPEKLDWIQKNEMSKLTRDEREVVTIAREEAEIRFLRLRLKFAKEQREKGDEMIEKMTAENEKLALKMQGVKNIPALREEVEHLRNQPTLAECHRIEAEHHEMKRLMMQLSLDWIRFMIQKVMEQKELKKKIQMDIELQQERIYQLEEEEKKAMDEIKRRVEEMEIQ

>C_latens_KNL1

MDGNRRRSILRQPRATSEEVTDENVLEGAQTAIVTNRRVSFHATKQVKEYDREYGKIINGTPVREKAFDTMSSDGGAMTPRVASVDMDISESSTSTTPFKVFDTNRHVDTDASSMDMSLDRTLNNETARLFDITREKTLVYEKTTEVTTKTTERILTVPTADADGGGGSGQDDTMALFNQTDRQEVDMSVDQGGANNETLNIFNTTNRDVTDMDITSAPQFKTPRLPVDAKKAPLTDNVEPIDMDMDTTLNAANDTMAVFKSPARIKKNELEPVDMDVTRDQINNETLALFQTPENRKKPMSVLQKTPEIGSEAMDISMAATPKVMTPKRAPAPVDMDMSMSSDDTLAVFKSTTPARKNFGGEEEMDITQRPASVADDTMALFKSPARVETAPTVIQPQQQLFEESMEMEEEDQSTVAPESEKPADSEGVVTSEAPEAPEAPTDIRSNHSSMMMETETSIVEGERAAVQMSMMDVTAPLDDVMESVFLNQTDSEKTLTTTSTTEESPRPKSSARDVTSSVHVSSVTLNVSTNRKENDSMIRTMQYTEVDTTNTLQNSSQRMEDSDEEEIKEETTARETTLIREESEDVTVQNPQDQNQQENQQEMSISSYNLSVFNQTTNSTGSVPISRRRRSLLREVCESQRRHAMMENSMNISTAGGETALAEYRKEKMNASGVMNQSLNQSAQGRDIFMMNTSIRSPNVRLAQETPPPKSPRFELPIYDPAVVNIVYLTPEDVSNPTPLPEAVEFQKVLAEESQRVQMEIQRKMKESGVPPEKLEWIQKNEMSKLTRDEQEVVTIAREEAEIRFLRLRLKFAKEQREKGDEMIEKMTAENEKLALQIQGVKNIPALREEVEHLRNQPTLAECGRIEAEHHEMKRLMMQLRLDSIRFMIQTVMEHQELKKKIQMDIELQQERIYQLEEEERKAMEEIVRRVKEMETQ

>C_sp51_KNL1

MENKKGRRNSILKTRTTTVDVLSETVDNGAGPSGPVTNRRVSFHNMKQIQEYDRHHGQMIEGTPIKEKITDTLGSDGILTPRGGNMENTNILEHDATFQVFNGAQKEHGAADMSLETTIGGNENNETARLFNVTRDKTVYYEEVIETTKTTKTVVKMFGATAGADDTMALFNQTNSEEVDMSMDQRQNDTMAMFNQTNPDGIDMDLDATRTFLVPSAPAPRAVGGGNDRTMNMDVVEEESRRANDTMNLFNMTNADNTDMDISQYPAPTTGPNDTISMFKSPAPPAAKKKSVFQHVDMDITGEEPVEPGSEAMDISQAPPPTPKVNDLRNVTLDMDITNEHHEEKTKETSRMDISAMKEEEETFSETMALFGSPARGKMPAPTTTATPNRGSKSPNETLQMFQSPARNEAKVPVSAQKTRIFDESMEIEESTLRPLDPAPEDAPEDAPDDVVTENSQNSESMVLEEKSVLEEERGATVVLQNSMIMDPREDMEVEMDQTIQKYLQMSMDISMDQSLQRTFQKTPKVHSDFQMSMLHRPKSVQDASTIHNESITLNVSQGAKTSESMMIKTMQISTDAADVTHTLQDTEQDDRTERDTERDEVTGEMDMEEDMEETLEEKEVSLRKKNDTTTSDTQEAEGESTKLQETFTTKFHVNQSIEDVTIQKNEEENHVDMSISMSSAILNQSCSMIRNSSRRRRSLLRDASIRESPRRVALENSMHISIHGGENGRMTALEEYRKNQSLLNATEQMNESGMNVSATSSSSNGARDIFVMNTSIRSPSAAKFAITNMNTPNAPSPISIPSAPSPGFQMPDYDPAVTNVVYLTSEDGEPLSEATEFQRLIMEEKKKVQKTLEQDSEIIEQNTVSKQDRQEIMKLAREEAEIKFLELRLQFAQEISRKQAAAIVSIESENSRIAESIKDAQSIEAEIATLRAYSPPVQYTRGRRENCANGGGDEEGERNDFGGVEGCYGAAMIFHIIDTSPMPYRLHHIIFLKIYFIIYFFFFVKFILRLQFHVRNHQKDLSDNPPHTYSLDLGQLESPVLPPLQLTTMPSLKNSNPTEVHISSGRDLVQRTLVFRNLTGKDFLLKLIASNESITFPTNVFRFPPLSHRVIQFRVATSKISQWDKSKLSIKGYVMPVYAKNLKQFINQKTTAGTACQEAFSLAIKFTDQFSAPQTVINLPGGATCIEATDHPVDVEELDTMTALNIERDVATAAPIGSMLGFVKEYQRGQKAKGCWLSNYICGGTAEKAEKEKPSMRSRRSSSASVSGKSQSSCRTQSRKKNKDVNVCLEATPCGGSTIQA

>C_sp44_KNL1

MENKKGRRNSILKTRTTVDVLCDTVENNAVRPGGNRRVSFQNVKHVQEYDRHHGQMIEGTPIKERITDTLDSDGILTPRGGNMDNTKVLEADRTFQLFNQREEHGAADMSIDTTLGGEADQTARLFGITRDKTLVYEEVVEETTKTTKRVTTKIFEPAGGEVAAGGNDTMALFNLTNTEEVDMSMDPKHDDTMAIFNQTNTDEVDMELEGTFVVPRLPSRVKGVNKTINQTVAMDLEDDQDQNMIRANETVNLFNITNTSCVDMDISRQPTSVCPNDTLAMFQSPNPKKKILTVVEEHVEEPVEMGSEAMDISVAPPTPKRENQKNLTMALFDVTRDEDQDMDMDITKDATYLKKDENHLDMDISVTVAAPTPISPPEDTHSETLKLFQSPARGGKSGNPQISKLFNESMEIENTNIHPDPAPEDPKSSESLADPDTSVVEEKRAALQNSCMDVTMPREDEQEEVAPEETTIQKSIHHTSMDMSMETTLRQNLFQNPEKNQNMGISTGVSQSVAQDASSIHNESVTLNVSSESKNKEPTKAANDSMVKTMQISTDTADMTRTLQETEIEEIQEKTDDEMDIEKEEETLKDSNDSSKIVEKTFNLSKVHIESTTSQEDVTFPNGQQVESTIHVETSHMDFFTSSHVNNKSGTPSTSILNQSCSMIRSSSRRRRSLLRDPSMRESPRRMALENSMHISMQMPNVKAGGSDGMMMTALEEYRQNQSLLNENSGMNESGLSVNNGTRDIFAMNTSLRSPRAAMANNRSFNLNSPRVPSPISINYTAPPAASPGFSMPTYDPVVVNVVYLTSTDGSTLPEADGFQKMILQEKKVLEKDLHDVDVPQANLSKRDEQEVMYIAREEAEIRFLELRLKFAENTNSSQIAEIEDLESENAKMAEELRDAQNLPAIRKELEAMKAQYTKAEIREIRQEYKRCKQMQFDAQLAALKHNYNVGLEIANKRSEIRMEMEEIDEIMARQDEKARKEREVLMKELKNIMEQ

>C_sp48_KNL1

MENKKGRRNSILKTRVTVDVLSETVDNNVPAGPAQNRRVSFHNMKQIQEYDRHHGQIIDGTPIKEKITDTLGSDGILTPRGGNLDATNILDPNSTFQVFNNGGGAADMSLETTENHDETARLFNVTRDKTIYYEEVVETTKTTKTKITKIAGSSGAAAAPGSSDDTMSLFNQSNTEEVDMSVDRPNDTMALFNQTNPDEIDMDISGATFAVPKMPRHQNRTMNRTLPMDLEEEEDQTKKANETMNLFNVSNLENVSMDISQHPAQQSILHAPNDDTLSLFRSPAPKKQVFQEEMDISQVQEPEEPGSEAMDISQAPPTPIHAKKNATMIALFEENDDMDITQNTTIVKETVEEKSCHMDISTSVRPDVQSETMALFESGGVSPAREKMTVPTANTSPQDQEKTADATLKMFQSPAGKQAENEMKMPPSAQKTMVFEESMDIEESTIHVESATSPKAPGSSTLQDSESTLHKSLQESSLMLEQSVVEEERAAVLQNSMMEVLPTDDQVEEEELGEIQKTSSFVRREVEEESMDVSMDISHQKTPQRSMHRPKSAQDASTIQNEASANLVDEEEPKNESTVKTMQISTDLDVTRTIQEEESEEDKTSGHPDGQMEELQETVQLFTSKLQVDSTELGEDVTIQMAMEESSIQVAQNNQSEISQFHSSLNESYSIRNTSRRRRSLLRDTMVRESPRRVALENSMHISMNMPNHYGKEDGRLTALAEYRQNKSLLNASQMNESGGSNVTMTASPGGNATRDLFAMNASIRSPQPTTINKSMNLNTTPRVPSPISVDQSVVIPPPPESPFQPMPIYDPAVVNVLYLTADDGTAHPEAADFQKVLMDEKQGLEEKDSEEAIQNSNLSKRDLRDMTLLAREEAEIRFLELRLKFATEHNEKQTRSIEQLTSENAEMAEQIRDAQNLPELQKEIEKLKIEKTQYTKAEVRQIRQDYMYWRKKLFDAQMTALKHNYEVAVAIADQRAELRMEVEELEERAGRSDSQDYGGMIFDTAIHHISLSPRHHPHINLFFCKINPNYPDFLMSKTY

>C_brenneri_KNL1_paralog1

MENRNGRRNSILKTRVTVDVLSETVDNNVPAGPAQNRRVSFHNVKQIQEYDRHHGQMIEGTPIKEKITDTLGSDGILTPCGGHMDVTNMLEHNSTFQVFNNGGADMSLETTVAGGEEENHHDQTAHLLNVTRDKTVYYEEVVETTKTTKSKITKICGSSAGGASDDTMTLFNQTNLEEVDMSMDRPANDTMALFNQTNSDEVDMDMSQSATFAVPRMPRHQNKTLNRTLPMDLEEEEEESQMKKANETMNLFNVSNLENTDMDISNHPAQNSILQAHNDTISMFRSPAAKKQVFQKQVEMDIYQAQEPEEIGSEAMDISQAPPTPSHVTKNATLAMFEDDDDNMDITQNTTIIKETTVEKINENCHMDISSVTVTNPDVQSQTMALFESGASPAREKMAVPTAISSPKDEDQSADATLKMFQSPARQAGVEMKMPTSTQKTMVFEESMDVEESTIHVESPKSPASEEQLATETTESSESPGSPESSQIQDSESTFHKSLNENSIMLEQSLVEEERAAVLQNSMMEVLPRDEQEEELVDVRKEEAMEQDSMDVSMDISHQKTPQRSMHRPKSVQDASTIQNETSANLVEEEERKNESMVKTMQILTDTADVTHTIQEEETLKNRTSGHSDGQTEELQVKSTELGEEVTIQMAMEESVAQNQSQMSFNESYSIRNTSRRRRSLLRDTLVRESPRRLALENSMHISIHMPNHYEKEDGRLTALAEYRQNKSLLNASQEQMMNESGSNLSMTTGAAASPGNGTRDIFAMNASIRSPRNTTINKSIDLNTPRVPSPISKDQSMVIPPESPFQPMPIYDPAVVNVLYLTADDGTAHPEATDFQKALMVEKLGLEKIQEVTIQNSNLTKRDVRDLMLLAREEAEIQFLEFRLKFAMEQSQKQTRCIEELTSENSKMAEQIRDAQNLPEIKKEIEKLRIETTRYTKTEVHQIRQEYMYWRKKMFDAQMAALKHNYEVGRAIADQRAELRMEVEQLEERAAQIKEKNRQERRELVDQILKIMEEW

>C_brenneri_KNL1_paralog2

MENKKGRRNSILKTRVTVDVLSETVDNNVPAGPAQNRRVSFHNMKQIQEYDRHHGQMIEGTPIKEKITDTLGSDGILTPRGGNMDITNVLEHNSTFQVFNNGAADMSLETTIAGGEEENRHDQTARPFNVTRDKTVYYEEVVETTKTTKTKITKICGSLGAAPGSSDDTMALFNQSNMEDVDMSMDRPANDTMALFNQTNSDEIDMDMSQSATFAVPRMPRQQNKTLNQTMLMDLEEEEGQTKKANETMNLFNVSNLDNTDMDISNPAQNSILQAPNDTISMFRSPARKKQVFQDQEPEEIGSEAMDISQAPATPSHVTKNATLAMFEDDDNMDITQNTTIIKETTVEKTNESCHMDISSVTVTLANPDVQSKTMALFESGASPVSEKMAVPTAISLPKDQEQTADATLKMFQSPARKTGEEVKLPMSTQKTMVFDESMDVEQSTIHVESPKYPAPEEPESSESAESPESSQIQDSESTFQQSLHEHSVMLEQSVLEEERAAVLQNSMMEVLPSDEQEEEQLVDVGEIQKTSSFVRREEAMEQDSMDVSMDISHQKTPQRSVHRPKSVQDAATIQNMTSANPVEEEERKDESMVKTMQISTDTTDVTHTIQEEEEETLKNKTSAHPDERTEELQQTVQMFTSTLQVDSTELGEDVTIQMAIEESTIHVAPQNQSQISQSSFSESYSIRNTSRRRRSLLRDTLVRESPRRVALENSMHISMHMPNHHGKVDGRLTALAEYRQNKSLLNASQDQMMNESGSNLTMATGAAASPGNGTRDIFAMNTSIRSPRNVTINKSINLNTPRVPSPISMDKSVVVPPESPFQTMPIYDPAVVNVLYLTADDGTAHPEAADFQKALMVEKQGLEKIQESDEEGVIQNSNLPKQDVRDVMLLAREEAEIQFLELRLKFAMEHSQKQTRCIEQLTSENSKMAEQVRDAQNLPEIKKEIEKLKIEKTRYTKTEVHQIRRDYMYWRKKMFDAQMAALEHNYEVALAIADQRAELRMEVEELEERATQIEEKNRQERRELVDQILKIMEEW

>C_wallacei_KNL1

MDRKKRNSILKIRQTVDVLSETVETGTTTNRRVSFHNVKHVKEYDRDHGKMVDATPVKEKISETVGSDGILTPRGGNMDVTNVQLEDGTFRVFNGAGAADMSLESTIGGAGENKNLTAGLFGITREETVMIEERINTNDTINIFNQTNTEAVDITVDGTVINDTMAIFNQTNSSEVDMSMDQSVFAVPKLPSKNKMKAELENVEEEKEMDISKGNETMKLFNITNPDVIDMDMSQMPTEARDETIAMFRSPAPPKKKEMDDVEMDITQTEQEEMGSEAMDISAAPVQQNSTLAMFVENEIKEKDMSVESTMDITASKEQEMDISVASPPSPEASETLALFNSPARSKKDQTLKMFQSPAQKTQIFDESMEMEEEEEDFVPQDPEVTPEVIPEVTLEKSTSQIESLKETLVVEEERAVLQNSMMEVEAQGEEEALLKGEEDIQKTTPSFMDFTHNMTVETSEKTNNESIVKTMQLNETGTIREEEEDVEKTITKNESLSMEDTSRKMVESSDDVTIRMHTKIVEEERYTQMENTTESRMSISTFSQNMSISTSTTLNLPPISSVSKKRRSLIRDQSARESPRRMAIENSMLSMHNGGGVALAEYRQNKSMMNTSVGQMNESGNMSIGSNHGRDIFIMNTSVRSPRHQTNSSINLNIPRQPSPIAQKTHRFEMPDFDPAVTNVVYLTSDEQTIPEAVEFHKMISEEKQRIQEEIQNSDELIKEKIEKTDAKLTNDEQEAVMVARKEAELVFLQRRLQFATNQNQIQEKQIQNLESENRETAERIRDAQNLPIIKKELKEMKNHPTKSECRRIRQEYMKMKKEWFEIQMRSLKNTYEAAVELADLRAEARREFDEREYEIKQLEEENKKKMSVFHGKVNEMIMQIEN

>C_tropicalis_KNL1

MERPKKRNSILKVRQTVDVLSETVETGTTTTRKGRVSFHNTKHVKEYDRDHGKMTDATPVKEKVSVSVGSDQALTPRGGNMDITNVQLEEGTFRVFGEADFGGAADMSLESTVIGENRNETANMFGITREEPMTFEERTTSGSRTNETLNIFNQTNTEVVDMTVDGEGIVNDTMALFNQTNSSEMDMSIDQTRFVVPKLPSKSKVNKNMTVNKTIPMELEDDEEAPVKEKVSLSVGSDQALTTSGSNMDITNAQLEDGTFRVFGEAVSGGAADMSLESTVIGENRNETANMFGITREEPMTFEEKTTVGSRTNETLNIFNQTNTEVVDMTVDGEGIVNDTMALFNQTNSSDVDMSIDQTRFVGPKSPSKSKMDKTVNKTIPMDVEEDEEVNVHIGNDTMKMDVNEDTEKTLTRNDSQLMEDTSRKMVNSSDDVSVRMHTKTVEEELYTQVENTITNQSRMSISTYSHNMSITTSSTLNLPPISSVSKRRRSLLRDQSIRESPRRMALENSILSMHFPNGGEEALTEYSQNKINTSTGKLNESSNVSCGSNGGRDIFLMNTSIRSPLHQTNVSINLNTLPQQSPIAQESARSSFHMPDFDRAVTEIVYLTSEDAEISLPEAVEFHKMISNEKSRIENEIQTKIQVSDVVLVEKMENLKKDQVSRMTNDEQKAVMMARQEAELQLLKLRLQFATSQRQNQEQIIRKLESENMETAEKIRDAQNLPSIQKELKEMKSHPTRTECRRIRQEYMETKKEWFNIQMKALKNTYDTAIELADFRAEARREFEERDYEMKRLEEENRKRMNIFRGKVNEIIQKVGE

>C_doughertyi_KNL1

MENRKKRNSILKVRQTVDVLAEIEDNSGPSGTTTTTNNRRVSFQNVKHVKQYDRDLGKMCDATPIKEKITDTLGSDGILTPRGGNMDNTNVQLDEHGSFQVFGAGGGAHNTHGEADMSLESTQGGGAGGGGAPNDTARLFNVTREKTIVYERITTERIVVVPQDSGAPNVASGASGAPSGAPSGALNDTLSLFNQTNHEEVDMSMDHHQQRKDDTMAIFNQTNTEEIDMSMDETHGFAVPKLMPRHHLTKNQMNKTMNETMPMDVVEEEEGLEIGKKGNETMSLFNVTNPEAMDMDITQTAALAPTSANDTLALFRSPAPKKKTMTSSVAEDVEPIDMDITTPQRPPSELGSQVMDISVAPTPRGNNSTLKLFESHLEPEKDTKEQDMSLEKTIQAPPPQSTEAIDMDISQLPKSPEEHPQNQSIDMDITVEKAAASETMALFQQSPARSKTPSANETLKMFQSPARGNEKMRMPGTIEKTKMFDDDSMDISIAPPTPKAASEDVSKSKTPKPAEKTKVFEDDSMDISVAPPTPKAVSEPKTPGTTEKTKVFEDDGMDISVAPPTPKAVSKDVSEPKTPGTTEKTKMFEDDGMNISVAPPTPKAVPEDVHEPKTPKSAEKTKVLEDDGMDISVQPPTPKSTPEDSEDVEDSDDVEELQKTLHDNSSMMISTETSVLEAGRAVHQTSMMEVYQEDVVETEGLLKHQTSFVVQNTSICRPSSGARDVSSLNNDSITLDVSAKNDSMLKTMQLTDDVDVTRTLNEDSDDVEKMEEEEEEKTLTEKSEPSEVPEDVTQKIQEETVVEERTIGFDNMSSTRFSYNMSMSTTTQVLNQSASASRRRRSLLREPSLRESPRRLALENSMMSMLQQPSYGGDALSEYRQNKMNTTLEHQQQMMLNESVQNTSTGSIGGGGRDIFAMNTSIRSPHHITQKNASFNMNTPRVPSPLAFEDSPAPPHSTVFQMPDFDPAVVNVIYLTPEDPDAAPFPESLEFQKLVDVEASRMQKEIQTILENSEDVGGLLEKKLNALRAKDMSKLTSDERDVVMEAREEAELYFLELRLKFAEKTLQNQRLQIEKREAENAEKAEQIRDAKNLPAIKKKLDEMKACPTRAERRQIRKEYMEVKKAVFDLQMERLKKVQNTIFEIGNQREELQDELYGLGEQLAQLEEQKRREKAAIMELVMEIYQQW

>C_sp54_KNL1_paralog1

MENRKKRNSILKVRQAVDVCDETVQTGGPSATTTITNRRVSFHNVKHVKHYDRDHGKLMDATPAKEKISDTLDSDGILTPRGGNNMSINDAASFQLFDGAGSQDKSMNMSLETTTNENNETARLFDITREKTVVYEKTVETTTKITERIVSVPPPPPSVTAADDTLAMFNTTNTEEMDMSINQSAVFAAPKLPRNNRTLNKTMPMDLDETTTANANETVNIFNMTNMDATEMDITLAQNTSQFKHPAAINDTLAMFQSPAPKLQQQRQQKDDVEHVDMEITQPIFEEPASEAMDISAVQIPMNASMNMSKVAIGNETMAMFQSPKKPEESFMDLEKTMENHLNSNSSSMDITQTPPEVMEKVEKEEPEVKGNDSMDISIVPPMTGANDTMAMFQSPARNNAQNMTAAPPASLQKKQQLFEESMDIEDEATLVAPEAPEAPEAPEDVPTVPEDVTPTPEEAIEDVTQESMMITETSVLEEERAAVLQTPVMELTPSEEDPGTLLLSAYHQIQKASSSSLMQEQERPQSSTSRHSTIFNDSSVGLINTERKLNDEDTMVKTMQFTETDATNTLQDTESISPMGKATEDVDVQMDMEEEEEETLQRNDTQQVEDVKYEEKLHETLQNSVNIQENINESIDTEVTIQKEMSIHQYTFNQSINQSITSCSSKYCSPAISSKRRRSLLNASNHRESPRRVALENSMLSLHLTTEALSGYRQNKMNLSSDSVANTSGGRNIFAMNTSVRSPHHLNTSMSSSTTPSPHHVTPIPTEVPPSPGFDMPDFDPAVYNVIYLTPEDVTKEPLPEAVEFSSLVAEEKAKMQKEIQESIETSGVSMEKIDYLRNNQISKLNSVERDAVIMAREQAEFRFLELRLKFATEQRKKNEQQIMEIQSENEKIAEQLRDAENLPALKKELETLKSQPTQEECRVIRKEWRKMKQWAFDMQMKALSDTHETIMEIREQKTDLLMEIEEREEKLAQLEDLEKKKKATIVEKVMEVMQ

>C_sp54_KNL1_paralog2

MENRKRRTSILKKRQIVETGESSGRRVSFHNVKHVKQYDRDTSEAMDISADQIPMNSSMNMSKVAIGNETMTIFQSPKKPEEKQEESFMDITQTPETLENMEKESEVKSNDSMDISIVPPVTGANDTMAIFQSPARKNLQKMTAAPSASLQKKQQLLEESMDIEDEETLLAPAASEAPEDVTQESMMITETSVFEEEAVLQTSMMELTPSEEDPGALLTSACRQIQKASSSSLMEALTSSSPGFDMPNVIYLTPEDVTKEPLPEAVEFSSLVAEEIQKSIETSKNSEVSMEKIDEAEFRFLELRLKFATEQRLKNEQQIKEIQSENAKIAEQLALKKELEMLKSRPPPTREECRVILKEWREMKQWAFDIQMKALSDIHETIMKIREQKTNLLMEIEEREEKLAQLEDWEKKKKATILEALEDFRQNVNFSDKLVFFGCGYAGRRPLFLTLRKTIFGSKPGLGRMTENRQAVKVRNKYLKRFKEVKKWELDSKKSIIDSVNLPWLMIFYTLETLIDKLTIEGSCERVNTKMYVFSGFYLFMILWFIFAYRWLKDVNEEREQMDSGEHQMMNCDEDKFDENIQSVRESINSLNKKSIAVCILYGTIVVLLLCVSSLS

>C_inopinata_KNL1_paralog1

MDNGRPGLPTTNRRVSFQNLKQVKQFDREHGKIIDGTPMKEKITDTLGSDGILTPRGGNMDISDSQSACVTLQLFDGINPTKETVNMSLETITNENIVNGRMLDISKGKTLVYENTVETTTKMTERIFMVPQSDETLKMFNSTICQHKDMDISLDGSAVSAAPKIIKHNVTLNETVPMDLDHSKALDGSYNETLNMFNVSNLGNCAMDITLVQNASFASKPASFSHRRPNETLEMFGSDPPSTLSQLNAIQLTDMKPGVPENPIENDYLMDISVQQGSPVAVNDTMAMFKSPSRKDMSISFDRSAVSAAPEMLKHNVTLNETVPMDLDCSRAAAGSYHETVNMFNVSNLENSAMDITLVQNADFASKPAPFSHRRPNETLEMFESNPPSTLPQLNTIQLADINPGLPENPIENEGLMDISVQQGSPVAVNDTMAMFKSPSKLNMKDKVKILLNPEIISKNETNRTVADDSMEIEETFNLEDRKIVVKQLTLETTSVTEKFPNLQAPPNENSDTVRISRAYQTEEDLENKRAHLNSSAMDVTQFGEDIDILNNLQELQLRPQFAASTEQSLAFHNSRVTVDVSGKLTENEETPKVMMMETMQISTTDPTNTLMEINADDNRTKMEGDNRKVIQVENSERTILEQRMNQTVTEIEKNIQNVQNASVNLSRTAIERQEDIDVENEHSRMLTTSFNLSIHQGISKRRRSLLRESVISRESPRRCIAEKSLLSMNLGTCTTPLNMDELSEQRQNAIKTSEDDDVSLNSSLNKSARDIFTLNIMVRSPRNTSINGKSPATPYSLSHKSPLDSLRSQDSKSPEFELPDFDPSVFNVIYLTPNNGKIIPEAAEFQRLVEDERKKVQLEIEDAVKSGEFFAEHWEILKNKQFSKLTYDEREVVRIAREMAEFRFLELRLKFAKEQRVKSEQKILELEEEIAKMAELKRELENLPAIRAELDELKRRPTFAECQRIRQEYSEMTKLVFEQQMELMRRNYEIALQIREHRATLLRELDEREEILNQLEDKERQNKAEIEKVVKNVMAHMN

>C_inopinata_KNL1_paralog2

MDNGRPELSTTNRRVSFQNLKQVKEFDRDHGKIINGTPMKEKITDTLDTDGILTPRGGNMDISDSQSACVTLQLFEGIDPAKETMDMSLETITNENNVTARMLDISAEKTLVNENTVEQTTKMTERIFSVPQSDETLRMFNSTICQHQDMDISVDGSTVSAAPKILKRNVTLNETVPMDLDYSRAVAGRYNETVNMFNVSNLENPAMDITLVENANLASKPASDIKPGLPENPIENEDLMDISVQQGSPVAVNDTMAMFKSPSRLNMKILPDPGIISKNETSGTIADESMEIEETLNLEDRKIVVKQLTLETTSVTERFQNLQDECKTLDENSDTVRISRTYQTEEDDVENERAHLNSSAMDVTQLGEDIDILNNREERQLRPQFSVSTEQSLAFHNSSVTVDVSGKLTEHEKTPKVMRMEAMQISAVDPTNTLMEINVDDNSTKMEGDNRKTIQVENSERTILENSMNQTATDIEKNIKNVENASVNLSETAIERQESMDVENEHSRMLTTSFNLSIHQRISKRRRSLLRESAVSRESPRRCIAEKSLLSMNIGTCTTPWNIDELSEQRQNAMKTSEDDVTLNSSLNKSARDIFTLNIAVRSPRNTSINGKLPATPYSLSHKSPLDSLGSPASKSPEFELPDFDPSVFNVIYLTPNNGEIFPEAAEFQKLVENEKKKMRLEIENGVKSGEFSAEQWEILKNNQLSKLTRDEREVVRIAREKAELRFLELRLKFAIEQRVKSEQQILELEEENAKMAELKRELENLPAIRAELDELKRRPTLAECKKIRQEYMEMTTLVWEQQMEAMRNTYKLALQIRDQRAALLREIEEKEEILNQLEEKEQQNKAEIENVVKNVMAHMI

>C_elegans_KNL1

MSMEPRKKRNSILKVRQAVETIEETVMNSGPSSTTTNRRVSFHNVKHVKQYDRDHGKILDATPVKEKITDTIGSDGILTPRGGNMDISESPACTSSFQVFGGGNLDKTMDMSLETTINENNETARLFETTRDPTLLYEKIVETTTKVTERIVSMPLDDTLAMFNTTNQEDKDMSVDRSVLFTIPKVPKHNATMNRTIPMDLDESKAAGGQCDETMNVFNFTNLEAAEMDTSKLDENNTMNAIRIPINSNVMPVDMDITEHHTLIEEKKNDTFGPSQLMDISAPQVQVNDTLAIFNSPRDICNKGLGVPQNLINIASNVVPVDMDITDQAVLNAEKKNDQFETSQLMDISIPKVLVNDTMAMFNSPKHVSKSSMDLEKTIEAADKSTKYPSIADEVEDLDMDMDITEQQPCEAGNQQNDGLQLQKEDLMDISVIRDSPAVNDTMAVFQSPARVKIGANNSIIDSQKSIVFGDEMSIDETQNDGTLTLPKSNVEVTTTNDVYTSLERQEENASENVSMINESSVHSEIDKKSFMLIEEERAFMHSSMIDVAQKLEDDGSSKTPVILASQSASLATKEPSALHNSSATLNNSMELDNNTLLKTMQITTCEDISMVHESIAVELNSNKEQEQFGDETLQKNDTSNTGANFTFQGHNETSQIMNNVDSEAVNTSKISTYSAFNLSINQSISKRRRSLLNSARESPRRVALENSIMSMNGQTMEALTEYRQNKTMQTSQDSMPSMSLNDSGRDILAMNTSVRSPHLNSSKTAAPGTPSLMSQNVQLPPPSPQFEMPDFDPAVVNVVYLTSEDPSTEQHPEALKFQRIVENEKMKVQHEIDSLNSTNQLSAEKIDMLKTKELLKFSHDEREAIMIARKDAEIKFLELRLKFALEKKIESDQEIAELEQGNSKMAEQLRGLDKMAVVQKELEKLRSLPPSREESGKIRKEWMEMKQWEFDQKMKALRNVRSNMIALRSEKNALEMKVAEEHEKFAQRNDLKKSRMLVFSKAVKKIVNF

>C.oiwi_KNL1_paralog1

MENRNNRRNSILKVRQHVELLDETSADGPQASSTTTNINRRVSFHNVKHVKEYDREHGKIIDATPQREKINETVASDGILTPRGTNMMIKEESTIPENVNGTVDMSVENNETLKMFETIVYEKTVETTTRSTVNAPAPADDTLAMFNVTTSEEQDMSIDQSRVFAVPELPGARAVSVPAAGEQDMSIDQSALLEAQKTMNQTVPMDLSGSKAANDTMSMFNATSNDTFDMDITAPNPPPTPTMTRRLPKNVFSGEEDMEIEETVVNISAPQNDTMSIFQSQSPARNKMPVVPEDPREEEEEVREPMDISMVPGNETIGIFQSPARQKIGNVSEMIPMAQVSSNDMDITNVTVSEEKNSEEVDMEIEDPTVQKIQNDTLSMFQSQSPARNRIVPVVPELQDESMDISAMPTIEPSDHQIPEDSTLILDKKDTSQDLLESPARQNMSEELADRSEKDLEESKNTSAIEEDRAVLLHSMMEVTAVQEEPDLDQKMPERRPQSAASTEQDSTINNASITLATSASAKSMMTSPQKTMQIEEDATHTLQETSLVQDQKDQEDQEEMILEKNESRSLIPEVQDPSDQEEMIVEKNESRSLIPEVPEARDPQEMIVVESRMSISYHEHLRIRTPSSSATRRRRSLLHSTDSPRRLALENAMAEKTALEEYRHQKSLHTSHNLSNLNVTAAANNMSISMSMSAAGRDIFDMNTSVRSPKCMVNSPSPANVFPSEAPPSPVFQMPDFDPAVTNVVYLTSESGTTHPEASEFRQMIISEAARLEREIQNTVGSGGVDSGKLEALRSGHFSRLSHDERDVVMKARDEAEIRFLKLRLKFAQEQRAKQDREILEVERKNEQLAEQLKNAENVEVLRAEIEELKRIPTRAEYEQIMKEWREMKKLEIDVQIQTIREAHHKLLQIQYETAQRTMELEGADESRRDERNCLIIPISHLQPK

>C_oiwi_KNL1_paralog2

MFLNQQEAPIVNEHCEYVKKILNHDVVLKARGEAEIRFLELRLKFAQQQRAKQDREISEIERKNEQLAEQLKNAENVETLRAEIEELKRIPTRAEYEQIMKEWREMKKLEIDVQIQTIRDAHHKLLQIQYETAQRTMELEGADESRRDERNCLIIPISHLQPK

>C_kamaaina_KNL1

MDNRNNRRNSILKVRQHLEVLDETSADGPQATTTNVNRRVSFHNVKHVKQYDREHGKIIDATPQREKINETVASDGILTPRGTNMMIKEESTIPHNGTMDMSMESAAANENNETMKMFDTTRETVVYEKTVETTTRTTEKIVIAPADDTLAMFNMTNSEEQDMSIDQPRVFAVPAVPSRAVPTTGGVQDMSIDQSAILEAPKNDKTVNQTVPMDLSSSKAANDT

>C_waitukubuli_KNL1_paralog1

RRSSILKVRQHVDIDIDETTVAGSGAPGASTTTTNRRVSFHNVKHVKQYDRDHGKLIDASPLREKITDTMDSDGILTPRGRNLDASEVAGNQTIHVFGNANNANQTADMSLETVNENVGSETMQLFNFTREQTMVFEKTTTTTTVTERRVEGPVMDDTMALFNMTNTEEKDMTVDQTAFAVPVSRAARPNATVNQSVLMDFDESTIVPQPNNDTMSLFNMSSVSVKDMDISNATVNETFKVPKSPSRSVMPRVFEENEDEVDMELTNVEGEKSIVANESMDISVMPAATPKKVNADQSDMKSIVANESMDISVMPAATPKKVNADQSDMVLEKTNLIQKTRPVLNESMDISVLQTESPKNFVGPNDTLALFQSPARQANQSYQHQEQPKNIVFRDQSVDMDITQTKPPAAAIKLNDSMDISEMPSEAHQSPARVQQSFVAPVSAPEPFDDNSMEVEEDVVREGTAQVIPTAEQTEMMVEEPSIVEQHEEERAMAAVSMMDVSGIQESLDYQQKLSTSVHASPALGSSPKESEVHIMDDESSVSILKDTTIASNAESTISPQKTMQIETDMTATPREDESMQMEDVENQEKTFNEFVEQETMTMTMSFSSAKKSTSHLSRRRSLLRESVLESPRRVAIENSMITMNMAEMTALSEHRRRKSIQLNQQNLTNTSALNDSANMSKGRDIFALNTSVRSPHCSSTPRSGNVSAAAEIQIPAPESFQMPDFDPAVTNIVWLTSDNGDPIPEAEEFAKLVNAEKIKAEQENRMRLLDQNIDKEKAEAVRNGQWSKLSHDEKEVVLIARQDAEIRFLELRLAFAEKTREKYEKEEKELTEANARFAKQLEDVESMNALRQEIEELKSQPTEEDFERIEKEWQEAKNEIFKWKRAALQHNLDILLQMSEAKRQLEMEIEVREEKLAQLEAMETAEKEELMKIVATL

>C_waitukubuli_KNL1_paralog2

dettvagsgapgastttMNRRVSFHNVKHVKQYDRDHGKLIDASPLREKITDTMDSDGILTPRGRNLDASEVAGNQTIHVFGNVNNANQTADMSLETVNENIGSETMQLFNFTREQTMVFEKTIKTTTVTERRVEGTVMDDTMALFNMTNTEEKDMTVDQTAFAVPVSRAARPNATVNQSVIMDLDESTIVPQANNDTMSLFNMSSVSVKDMDISNATVNETFKVPKSPSRSVMPRVFEENEDEVDMELTNVEGEKSIVANESMDISVMPAATPKKVNADQSDMVL

>C_panamensis_KNL1

MENRNRRSSILKVRQHVDIDIDETTVAGNGLPGASTTTMNRRVSFHNVKHVKQYDRDHGKLIDASPLREKITDTMDSDGILTPRGRNLDASETANNQTIRVFGNVDNGNQSVDMSLETVNENVGGETMQLFNITREPTMVFEKIVTTTTTVTERRMGPQGVLQSQAAPLDDTMALFNVTNAEEKDMTVDQTRMFAAPEKRLVHSQTTMNDSMDISVAPSDTPIRVNDTMALFRSPATRANESDMVMEKTNVFEQEEGKVLHKQTMPPVMNDSMDISVMPDDSPKNLRALLPNATLNQSVLMDLDESNAAPANNDTMALFNISTASVRDMDISNATIHETFKVPKSPARSVLPLVREENDDDVDMELTSMEGEKSIVQNQKTVALNDSMDISVAPVNQFLPVFPVVEQEEANQQNTLAPIVTDDAMDISVMPSDLPKNIDSNDTLALFQSPARQVNQSSQQPVHNVFNEPKDESVDMDITQTKEVATPQRMNVANKSVVVAPTVLKGPKDDSMEIEEAGAGEGTFVVPTTESPETSPEDILIPNIEQANMGGDLCDKIIVEEPSTVEKHEENRAVAEVSMMDVSGIEESPSHQKFPNLQASPTEESKSNSTLHESSVSHNDSSPKDTSTRGNESTLSPQKTMLVETDMITTLHDMSLVQKENDSEEMEEQAEVEEPEKTVTEIVEQNTTTTAQMIMSSLNTSSNRKPTSHASRRRSLLRESVLESPRRIALENSMITMNVAEMTALSEHRRRKSLQLNQQSVLNASNISMLNESLNMSKGRDIFALNTSVRSPHCHETTQSGNMSIVESGAAPETTSFSMPNFDPAVTKIVWLTPEVGEDSIPEAEDFARLVAAEKIKAKQHAQKTSENLDKEKAEAVRTGQWSKLSHDEKDAVLIARQDAEIRFLELRLTFAEKNRLKYEHEAKELLESNARIAEQLEDIENMESLQKEIEELKNRPTEADYERIGEEWRVAKEELFQLKHALLQQTLHIIMQMGAQKQQLKSEIEDRQKNLASLEAKETSEMQEMLKMMETL

>C_nouraguensis_KNL1

MTDETTVADNGLPGASTTTNRRVSFHNVKHVKQYDRDHGKLIDASPLREKITDTMDSDGILTPRGRTLDASAAAANQSIHVFGNVEHGAADMSLETINENAGSETMQLFNITREQTVVVEKSITRTVTERVVEQHVARGSNDTMSLFNMTDCDEKDMSVDQSAVFAVPRVLQQNTTINQSVLMDLDESSVETGTAPGPSNNDTMSLFNMSTASAKDMDISNATMNETFKEPKSPARAFMPKVIEEDNEEIDMELTQMEVEESIVQNQNTIAFNKSMNQMAAKSAVFAVPCVLKQNATINQSMLMDLDESSVGIGPAAVPSNNDTMSLFNMSTASAKDMDISNATINETFKEPKSPARVFMSEVIEEDNEEVDMELTQMEVEESSVQNQKTIAFNSNMSTASAKDMDISNATMNETFKESKSPSHVFMPNVVEEDNEEIARDLTQMEAEESIVQNQNTIPLNESIDQMTATSAVFTVPCVLNENATINQSMLMDLNESSVGTEPAPVPSNNDTMSLFNMSTASVKEKVISNVTMNQTLKEQKSPARALMPKVIEEDNEQIDSDLTQLGAEESIVQDQKTIPLNFDMSTASVKDKDISNTTMNETFKEPKSPARAFVPKVIEKDNDEINMELTQMEAGESIVQDQKTIPLNFDMSTASVKDKDISNATMDETFKEARSPAPVFIPKVIEEANEQIDRDLTKMETEESIVQDQKAIPLNSNMSTASVEEKDISNATTNETFKEPRSPAHVFIPKVIEEDNEQIDRDLTQMKAEESIVQNQNAIAFSELLDQMAATLLQMKELKTEVQMQIEEREEMLARLQTLEDSTTEMKRKTEEIRDSIRDY

>C_becei_KNL1

MDRNRRSSILKVRQHIDIDETTVAGNGLPGASTTTNRRVSFHNVKHVKQYDRDHGKLIDASPLREKITDTMDSDGILTPRGRTLDASVAAANQSIHVFDNIDQGTVDMSLETVNENAGCETMQLFNITREQTVVIEKSVTRTVTERVVERHVAPGSNDTMSLFNMTDCDEKDMTVDQSAVFAVPRALKQNATINKSMLMDLDESNFGSGPAPGPSNNDTMSLFNMSTASAKDMDITNATMNETFKEPKSPARVFMPRVVEEDNEEIDMELTQMEAEKSIVQNQKTMALNESMDISVAPSPRRPNANDTMALFRSPARADQSSMVLETKKTIFEAEPEDVDMEITQMDEQKPMEQQPAMNETMDISVAQNAIVNNTMALFQSPANQANQSYQQMKIVAEDDNGVEMEIMQTEEPERMSQTMDISVAPTGSPALPGNNDTMAMFQSPATVNQSAVVIPVNTPKPFDVSMEQEEEGAGEGTFIAQNDISLDDQSPEAVTTPITQPTNFEENKNEFIIAVEPSSIEKEEERAVGQASMMDISNMQDSTLDQKSLILQASPAQESGNHSILHESHVSETNNESVMSPQKTMQLETDMTNTIHEMTPIENEDVSMQVEEQVEIQEKLVDDVVEKETTMTSQLTMSFTSANKSKSHLSRRRSLLRESILESPRRIAIENSMITMNIGEMTALSEHRRRKSLQKSQQNILNSSAASLLNESLNVSKGRDIFALNTSVRSPRVTGTPQSGNDSTIAAIPETSSFHMPNFDPAITNMVWLTPEEGEEPLPEAAEFASLIAAETLKAEQEVQRTLQNENLDKEKAEAVRTGQWAKLSHDEKDAVLLARQDAEIRFLEMRLAFASKTRLKYEKVVQEQLEKNTRMAEQLNDIEDMDVLRQEIEELKKRPTEADFRRAKNELKATRAALVERQAHHLAQMATGVMQMRNQITELRMEIEEREEMMARIETTVDSEMEKIRNQTEDILESMRDY

>C_yunquensis_KNL1

MDNRNRRSSILKVRQHIDITDETTVAGNGLPGASTTTNRRVSFHNVKHVKQYDRDHGKLMDASPLREKITDTMDSDGILTPRGRNLDASETAVNQSIHVFNNIDHGAADMSMETTVNENAGSDTMRLFDITREQTVVFEKSITTTVTERVVERHVAPASNDTMSLFNMTDCEEKDMSVNQSAMFAVPRISKQNATINQSMLMDLNESNVGSGPAPGPSNNDTMSIFNMSTASANDMDISKAINETVKEPKSPSRSFMPRVVEEDNEEVDMELTQMEAEKSIVPNQKTMALNVSMDFSLAPSPRQIGANDTMSLFRSPARADQSNMILETTKKVFEDEDKTVYMDITQAEEEKTLVQPAIEESMDSSMPQNVSISKTLALFQSPALESNQTHQPTQAVSDVVKMEVEMEEQEKMSQSMDISVAPMDSTDLLGSNDSLALFQSPARINQSNVTVTIDKPKLFDESVEMEQEAGAGEGTFLVQNELSSDGRSPETLPAPVAAQITHLANMETDQSEMKVAVEPSASEREEERAVTHTSMMDVSGMHDSPTLDQKSFVLPASPAQESASQSILQQSHVSETNDESVMTPQKTMLLETDLTNTIHESTPIDKENLSMKMEEQVEIHEKTVDDVIEKETTMTRELTMSFMNTSANRSKSHLSRRRSLLRDSIVESPRRIAIENSMITMNAAEMTALSEHRRRKSLQITQQNLLNSSTASMLNESLNASKSRDIFALNTSVRSPHFPGTPQSAHLSKDATIEVVPEASSFHMPNFDPAITNIVWLTPEEGEGPVPEAAEFASLIAAETAKAEQEIQRTLQNENLNKEKAEAVRTGQWAKLSHDEKDAVLIARQDAEIRFLELRLVFAEKTRQKYEKQAQEQLEANARLAEQLEDIENMDALREEIEELKNRPTEADYQRVKAEWRAAKREYYRLKNAEIVQILVTLQQIKEQKAQFKMEIEEREEKLARLQALEDSELETMKHIVLDIHESLRDN

>C_macrosperma_KNL1

MDKRNRRSSILKVRQHIDITDETIAGNGMPGASTTTNRRVSFHNVKHVKQYDRDHGKLIDASPLREKITDTMDSDGILTPRGRNLDASEMAANQSIRIFGNVENGNQTADMSLETVNENEGSETLKLFDITREQTVVFEKTVKTTTLVTERVTERVMAAGSLGDDTMALFNMTDCEERDMSVDQSAVFAAPRAPRNNLTINQTVPMDFDESRVGPANDTMSLFNMSIASAKDMDISNATVNETLAKSPARSVMAKVVEENDEDVDMELTQMEAEKTLVQNQKTIGNESMNISVMPTEPFDSPKRAAVNDTMSLFRSPARVNQSDMGMETTKKVFEQEEFVAEDVNMEITNVEEPKTSEPLVLNETMDISQMRSESPKTTVVNETLALFQSPARGVNQSLQQVDSVKKVFHQEEQHHVDMEVTQTEEQEEVPMSQPMDISIAPSDSPKLAKNNDTLALFQSPARLNQSALIASLKKKELFDDSMEVEEEQGAGEGTFILPTESPASVPPEDVVIPFNQAINAEDHQSEMMLTEEASALLVREEERAMAHTSMMEVSGVQESVDINQKSLALPSDSAQETTDTSTLQEKDVSVDKVRDVSINETTSHEQESAMSPQKTMQIEADLSTLHEMTPVQNEAVSMHLEEDGKMEVEVHEKTSGEFVETETTISSHMTMSFSGSSRKPAAKSHLSRRRSLLRESVLESPRRIALENSMITMNVGEMTALSEHRRRKSLQLNQNNTTNTSVLNESANLSVARDIFALNTSVRSPHFPSAPQSTNDSTVDAHSTTESTSFQMANFDPAVTNIVWLTAENEKPVPEAQEFARLIDDEKIRMEQEIEKSLLDDTVDQGKSEAVRDGQWSKLSHDEKEAVLIARQAAEIRFLELRHSFAQTHRQKYEREAQELLQSNAQMAGKLSEIENMETLQAEIDELKNRPTEADYHRIKAEWKAAKAESFKRKLAALGQIHDTLLLMKEQKSALKMEIEEREEKLTQLQITENTEKEAMKTIVEDIMKSLKN

>C_sulstoni_KNL1

MEHQKKRLSILKVRQPLETLDETNALGQGAPGESVTTTTRDRRVSFHNVKHVRQYHRDHGNLIDASPLREKITDTMDSDGILTPRGRNLDASETGENATINVFGKQNGTMDMSLETENGGNDTMKMFEVTTHEKTITTVTERIVERHVVGIPKQNTAADDTMVLFNTTNNEEKEMTIDESAIFAVPRLPNPNATSNQTVRMDLDESQVPQSNETMNMFNVSTASVHEMDISNAQNSTNAPKLEDVDMELTQVEQPEEATSRVMNQSTMNQSMDISIAPGAAVGGVNETLAMFQSPARGALINESMDISMAPEQKSPAREVVAPAREVLAPTANNETLALFQSPASVRRNEHSTMILERTQRVFEDDPNDVEMEITQVEQPEESPRAINQSIMNQSMDISVAPAVNETLKMFESPARNRTIPIELFNESVEMNEETAIPIDDSVPMVEQVSEQSPLVAEEERAQVQSSMMMDISSVRESLVDQSMQKSSQETEETLTEKSVQETLEESMQASSPQQKTMQIEKDATITPMPEVFEETMENQKTFQENTFTKDISITMNSSSIRNTTLPMSNRRRSLIRECSSRAAVLESPRRVALENSMISMNVGNMTALVEHRRRKSIQMNQSQKELTNLNESIANMSAGRDIFALNTSIRSPKPMGTPSDNVSLIHIAPESPPSFQMPNFDRAVTNIVWLTPDEHEDVAPPEAAEFARLIAMEKLKAEEEVRMTLLDRKLDPKKVVSGGKVSSKLSNDEKEVVLIAREEAEIRFLELRREFAAQTRRKYEEQVQCLETENARMAEQLRDVEQMEALKEEIEELRNRPTHADYERTRAEWKTAKREAFEMQMEALRAAQETLIALREERLSLLIDIEERQERLAQAEEREEQEKTEMKAILEGLMSGI

>C_afra_KNL1

MENQKKRLSILKTRQPLEILDETNALGQAGPGDSVTTTTRDRRVSFHNVKHVRQYHRDHGNLIDASPLREKITDTMDSDGILTPRGRQLDASQLGENATINVFEKQNGTMDMSLASENAGNETMKMFEVTTHEKTVTTVTERIVERHVFGIPKNTAADDTLALFNMTNNEEKDMTIDQSTVFAVPRLPEPKSASNQSMQMDVDESHVPQDSEKSPARGVLHQSMDISVASVQKSPARQRVPETNNETLAMFASPARGVLNESMDISMVPESKSPAREAILGPTGINETLALFQSPAPVRQNQTASMILDTPKRVFEEDPSDVDMEITQVGCQEEEELPRVVNQSMLNQSMDISIVPPVNETMKMFESPARIKTIPVEPFNESVEMETDDVVPANDPEPEQGSEQSPLMAEEERAQVQSSMMMDMSSIQESLVDGSALKTFQETDDTVTEKSLHESLGESMQIQSPQQKTMQIEQDTTITRMSEPLEETMENVKSFQEETFVKDTSVSMNSSSVRNTTVPMSARRRSLIRDCSSRGAVLESPRRVAIENSIVSMNAANMTALVEHRRRKSIQLNSNQKDLSNLNESVANMSAGRDIFALNTSVRSPMPSAASTDPAFSFHIAPESPATFLMPNFDPAVTNVVWLTPAENEDVPPPEAAEFARLVAAEKLKAQNEIEKALLDGKLDPKKIVSRGKMSSKLTDDEKEVVLIARQDAEIRFLELRRDFAAQTRQKYDLKIRELETENARIAEQIREFEQLEELKKEVEAMRNRPTYADYEKIRAEWKAAKKEMFEMRMEALRSAQETLVSLEEEKMALLVDIEERQERLAQMEDRAQREKAEMKKILEGMMSSF

>C_sp49_KNL1

MDKKARRSSILKVRQPVDVLDETTAPSAATTTMNRRVSFHNVKHVKQYDREHGKIIEESPLREKITDTLDSDGILTPRGRTDVPQYNESFSVFGQGKSNNVTMDMSLETVKTEDETIGMFEGLQTICEAPEKENATIHGERDMSIDQSAMFAVPHQPNATINQSVLMDIDESDRTLKANDTMNMFAPRNNSPGRQNVTKVFSDDDMEMSQLEKTHMEEKNQEKELDMDISEAPTNKNDTMQMFADKKTATFAEDKKVFGEEEMELTLVETETPEGVHESMDISMTPRKTIVRENDTIGMFHSLEQRQEHVDMDISQSFRADDTRQMFKSPGAKKGGDDMELTLQESPARATKNISMSKQKVFEEEGAGEEVPEENVSPGEVDRDQSSLHITLDKKEEERAEMQTCLMEMDGTNTTFNHSQKTDLPSNESISPGKTLCEDTLASPQKTMQIASDLTNTFHETSPIREIVEEVFVEHQSFHNDSCDMSVDMEVTSRLSVSYNRSGVLNRSSSSARKRRSLVRESPRRMAIENSMLSGAAEMSALGEYRRRKSLQTSQQHLLNSTNPTNTSLGRDIFAMNTSIRSPGRPGMLKKVDDSPAPEKFSFELPEFDAAVANIVWLTSENGHPIPEAETFANLLNVEKSKTESEIQKTLSHVDGAKFQAVRSGELSQLSHDEKEVVLFARGHAETRFLHLRQDFAQEHSQKYEEKISNIQLENAQIASQLLDLEDEEPLRHQLEEKRKHEVLMEADKIREEWRERERLETERRSERLAASAALILKMHQEKLAMQKEKEELDERMARLEDEERAVTEQISEEIKMVLAAAAAH

>C_sp25KNL1

MDQKARRSSILKIRQPVDVLDETAAPSATTIPANRRVSFHNVKHVKQYDREHGKIIEESPLREKITDTLDSDGILTPRGRTDGPQYDHSFSVFGQGTSSNATMDMSVETVHAEDDTIRMFDGAKTFCAPSEKENVTIHEERDMSIDQSVVFAVPHPMNATINQSVLMELDESERTLRACDTMDLFAPLTSTNNDNSVAPNDTLTMFQSPARKNATKVFSDDDMELTHVENQEHVDMDISEAPNNKTDKNDTLQMFSEVKQEKVVAETKVFEDEMELTLVEKAGDDSMDISMTPCKKTILPGDDTIGMFKSPEQKNNLQHDDMDISQSAEIHQMCKSPETKGGDDMEITLQESPARAVKEVSMCSMQKVFDEEPKEKTLPEEIRDHSSTTIDGREKERSMLQTSLMEVVAAETTLNNSEKTEIHSVSPGITLGSPQKTMQIASNLTSTFNETSPLQEPIQEVVEHIVEHQSFHNESYDMSLDMEVTSRLSVSMNRSLTSRRRRSMLRESPRRVALENSMISFGAAEVTALEEYRRRKSLQKSQQNLLNSTNQADLSLGRNIFTMNTSIRSPKTIGKLTIPEEPAPVPFELPEFDAAVANIVWLTSEDGNPIPEAKQFTNKALDLIDGAKFEAVRNGKLSQLSSDEKEVVVFARGHAESKFLHLRHAFAEEHLQKYGEKIHELQMENAQLAGQLLDLEDEELLRREVEVKRKLVVLADADKIRAEWRENQRLETERRIEQLTTSAALMLELHQERLAMLKEKEQLEEQIARLNIEQQAFDDQLLENLEMINAKALE

>C_imperialis_KNL1

MDKKSRRSSILKVRQPVDVLDETAAAPSTTNAVMNRRVSFHNVKHVKQYDREHGQLIEESPLREKIADTLDSDGILTPRGRSVNVSYHNETLVDMSLETVVAENDTIRIFDTIHENELEHKENGHQFAVPQLPNATINQSVLMELDESERASDTMNMFNPPANVSMDISSYTQNATKVFADDDMDVTSSRIQHEDTMKMFETPKKVNESMELTLVEEAPIQKTSDDMDISMTPSRIQKAPATQSASDDTVMMFQSPIQKANDDMDISITPSRILKTPKAQSAPEDTMVMFQSPIQSQKSSDEMDISMTPSRTLQRTPRAQNAQNAPDDMDISMTPIQKTPRVQKTSDDVDMEITLLEKTLDQNDTRQLFKSPARTQKMFVEEPEHAVAVNAPESPIAEAVAEIDRDESSMMITEQPSAMDLTDDMDQKTLDENQKTSEFQKTLTPQKTMRIDTEMSDIQNTLHETSDIQKTPQKTSMCNDVTSDHQNPAQKTLTPNQAVESPQKTMRIESETSELQKTFNETCEMSISRSHLSISFNTSITSIRRRRSIQKRESPRRLAMENSMITMSMAPEQMSALGEYRRRELLQTSQQHLLNSTNLADTSRDIFKMNTSIRSSPHTFSKQKPNEPAEPLKLPEFDAAIANVIWLTSDDAGALVPEVEHFQLLLEQEAQKTKALIHNQDSPNSLSNQDEQEVLLVSRELAEQRFLDLRLKFAQEQSHELQIQKIQLENAQMASNLLDLEDEHQLLGELEQLEKQKVLANADQIRAEWREKKKAESEQRVAAISGAVELMLQLHQKKMEMKKEMEELAERIAQLEEAEAKFKEQFIEQINTNVPY

>C_japonica_KNL1

MENKKKRLSILKVRQHVDLLDEATEVKAPGATTPTMNRRVSFHNMKQVRQYDRDHGKMIEESPLREKITDTMDSDGVLTPRGRTDAGNSTINVFGNVNKIDQLANGTMNMSLETVNVNDDTLKMFSGLRDRTISESTMMTEGVVLQKQNMTIREVRDMSIVDESEIFAVPQMPKPNATINKSVFMDLDESQSHHGQASSQNDTMALFLDATNNANISMDISIGSARQNVTKTFDDDDMEITQMDEKTLIRKEVDETVDMFESPARNHIAKKESDDVDMEITLVQMTPKSSTSKKEEDVNETVGLFKSPARNQQQVFKEVEEENNDMDITQPQMTPKNKSIQQINTVSPFDSPKIIEPNKSSEDVDMEITLPHNQMTPKLTTRQRTGEINETFDMFKSPAPKQIDTSKFLQNDENVDMEITQEPANVTSDSRSEKSSYDVVETNSEESLAILEKSHSLVEVAEQDRAMGNCSVMNISDVQETLNLQNNSPKLHGFADISQESMNRSTTSMSSHKLPQDDTDSSPQKTMQIESTTTLHEISERIEEVIEEKVENVEIEIEVEVEVEQKTFQTENYDVTVTESTTKKQIHALFNHSLNGSQNRTHSARRRRSLLRSSVLESPRRIALENSMVSTSVARERTALEEYRRRKSLQTSQQSILDSSNLHDVSASARDIFALNMSVRSPHQPDTPKSKIAQEVVSPAPQFVPFQMANFDPAVTNIVWLTPDEDSATSIPEALEFSNILAAEKTAVGDEIQKALEYGQIDEVKWEAVRNDLMPQMSQDEKEAVLIAREEAEIRFLQLRLRFATEHHQNYELKVNEIQTENKTISDKVMDLQNIDSLSAAINEFERREVVREKEKIVAEWREAKQMSWDRVSKKMNAAMKLLLEINQQREADRKELEEIEEKIARLDDASKKERAEIMDTISRIKMAAQAGINVS

**Supplementary Data S5: All ZWL-1 sequences used in this study**

>C_tribulationis_ZWL1

MSIKIEDLQNYNQAVSARGEKEQAPTPVLLLNKYRVRLIPISSLPFVSNYSNVSQLGLNSDDALVIDSPIESSEKQKKSSLVPRRQAARVNEDEDIMETESVEEITEDKENSVTENDRGPLVTSFLTLEKLEKNGDVEIVGLDCEVEFFDANPVPFEDGVSLQRLRLESSNHFSAAVNKYPIWISTFGSKFPSLCWLAAGRTAKNVQIGGATRVLGYFKDDSEKLVKQLNEACGVAQINRYRAVFDEIRRVATPTRDSPGEVIVDMRWTTKSSLVLLEQPDNAADCTIKIDLGWRDNRFFIDDAIFEQLFFVLNLADVLANPEKEVEFPPEFDKFDYLVQEMNDLVEACSREDNVFASNEFRSEEVTDKVWNIVRQCGDVKRATMLFKNFLQALTYGKIKSHVQEGNKSHLASLIRASKTCDFRMPILERLSTIEMMMEIGVESLRRIINKFSNTLQFPSDELTFILKTCENDLSSGEGAINASVISLLPITMALATVYQIFGLLNVKDHVILPDLARRVLTKFTSSMVEKAKRGETETDYTFETTLPLLRINKEIFMDKRPRVWTCENANTVGANVQTRIMTALELDVSLEHVSRLVNASRPVRLIDEENRKPTAEELKADYTVSHTIFSYLPKL

>C_sp41_ZWL1

MSIKLEDLQSYNEAGLARGEGDDNPIPVLLLNKYRVRLMPLSSLPFVSNYSNISQLGLNSDEALVIDSPIESSEKQKKSSSLFAKRETVTVKEDDEKMETDSVEEDKENSVTEEHNGPLVTSFLTLDKLEKNGANDVEIVGLDCEVEFFDANPIPFEDGVSLQRFLRLESSKHFSAAVNKLPIWISTFSSKFPSLCWLAAGRTNKNVQVAGATRVLGYFKDDSEKLVKQLNEACGAAQVNRYRAVYDGIRRIATPTRDAPGEVIVDMRWTTKSSLVLLEQPDNAADCTIKIDLGWRDNRFFIDDAIFEQLFFVLNLADVLANPEKDVVFPPERDNYDNLVQEMKDLVEACSHEDNVFASNERSEEVTDKVWNIVRKCGDVKHATMLFKNFLQALTYGKIKSHVQEGNKSHLASLIRASKTCDFRMPILERLSTIEMMMEIGVESLRGRIINKFSDTLQFPSDELTFILKTCENDLSSGEVALNASVVSLLPITMALATVHQIFGLLNAKDHVILPDLARRVLTKFTSSMVEKAKRGETETDYTFETTLPLLRMNKDTFMDKRPRIWTCENTNTVGANVQARIMTALELESSLEHVNRLVNASRPVRPVEEENKKPTTEELNADYTVSHTILSYLPKL

>C_zanzibari_ZWL1

MSIKLEDLQSYNEAVSARGEEDEAPTPVLLLDKYRVRLIALSSLPFVSNFSNVSQLGFNSEDVLVIDSSIESSEKQKKANIFVKRETVELDDEEMETESIEEDKENTTNEDEGGPLVTSFLTLDKLERNNDVEIVGLDCEVEFFDTNPIPYEDGISLQRLRLESSKHFSAAVDKLPIWISTFGSNFPSICWLAAGRTNKNAQFAGATRVLGYFKDDSEKLIKQLNESCGAAQVNRYRAVYDGIRRIVTPTRPSPGEVIVDMRWNTKSSLVLLEQPDNAADCTIKIDLGWRDNRFFTDDAIFEQLFFVLNLADVLANPEKEVIFPVEYDKFDSLVQEMKDLVEACSHEDNVFASNEFRNEEVTDKVWNIVRKCGDVQHVTLLFKNFLQALTYGKIKSHVQEGNKSHLASLIRDSKTCDFRMPILERLSTISMMMEIGVESLRRRIINKFSDTLQFPTDELTFILKTCENDLSSIEGALNTSVVSLLPITMALATVHQVFGLLNVKDHVMLPDLARRVLTKFTSSMVEKAKKGETETDYTFETTLPLLRMNKDTFMYKRPRIWTCENTNTVGANVQARMMTSLELESSLEHVNRLVNASRPIRPIDEENRKPTAQELNADYTVSHTVFSYLPKM

>C_sinica_ZWL1

MSLKLEDLQSYNEAVLNREEEDEAPAPVLLLDKYRVRLIALSSLPFVSNYSNVSQLGLNSEDVLVIDSPIQSSEKQKKSSLFTKRATVPVKEDDEKMETESIEEDKENTAAENEGPLVTSFLTLEKLEKNDVEIVGLDFEVEFFDANPIPYEDGISLQRLRLESSKHFSPAVANLPIWISTIGSNFPSVCWLAAGRTNKNMQFVGATRVLGYFKDDSEKLVKQLNEACGAAQINRYRAVYDGIRRIATPSRDSPGEVIVDMRWNTKSSFALLEQPDNAADCTIKIDLGWRDNRFFIDDAIFEQLFFVLNLADVLVNPEKEVVFPTEYDKFDHLVQEMKDLIENCSQEDNVFASNEFRSEEVTDKVWNIVRKCGDVKHATMLFKNFLQALTYGKIKSHVQEGNKSHLASLIRASKTCDFRMPILERLSTIEMMMEIGVESLRRRIINKFSDTLQFPSDELTFILKTCENELSSGEGALNASAVSLLPITMALATVHQIFGLLNVKDHVILPDLARRVLTKFTSGMVEKAKRGETETDYTFETTLPLLRMNKDSFMDKRPRIWTCENTNTVGANVQARIMTAMELESSLEHVSRLVNASRPVRPIDEESRKPTAEELNADYTVSHTIFSYLPKL

>C_nigoni_ZWL1

MSIKLEDLHSFNEAVLSKGEEDEAPAPVLLLDKYRIRLVPITELPIVSNYSNTSQLGLNSEEVLVIDSPVESAEKQKTSSLLNRRECKKTIKSEKEDEPMDMETTEGDKENTVSEIGGGPLVTSFLTLDKLEKDGVNDVEIVGLDCEIQFFDANPIPFEDGISLQRFLRLEGSKHFSAAVDKLPIWISTIGHHFPSVCWLAAGRTNKNVQVSGATRILGYFNENSQKLVKQLNEACGAAQVNRYRAVYDGIRKVATPTREAPGEVIIDMRWNTKSSLVLLEQPDNAADCTIKIDLGWGDNRFFIDETIFEQLFFVLNLADVLANPEKEVVFRSESDKFDDLVQEMKQLVEACSHEDNVFASNEKSDQVTDKVWNIVRKCSDVKQATMLFKNFLQALTYGKIKSHVQEGNKSHLASLIRASKTCDFRMPILERLSTIEMMMEIGVESLRGRIINKFSDTLQFPSDELTFILKTCENDLSTLEGTLHSSVVSLLPITMAMATIHQIFGLLNAKDLVVLPDLARRVLTKYTTSMVEKAKRGETETDYMFETTLPLLRMNKEAFMHKRPRIWTCENTNTVGANVQTRAMTTLELEPSLEHVSRLVNASRPVRPIDEDNRKPTEEERNADYTVSHTIFSYLPKL

>C_briggsae_ZWL1

MSIKLEDLHSFNEAVLFKGEEDEAPAPVLLLDKYRVRLVPITELPLVSNYSNTSQLGLNSEEVLVIDSPIESAEKQKTSSLLNRRENKKTIKSEKEDESMDMETAEGDKENTVSETGGGPLVTSFLTLDKLEKNDVEIVGLDCEIQFFDANPIPFEDGISLQRLRLESSKNFSAAVDKLPIWISTIGHHFPSVCWLAAGRTNKNVQVSGATRILGYFNENSQKLVKQLNEACGAAQVNRYRAVYDVIRKIATPTREAPGEVIIDMRWNTKSSLVLLEQPDNAADCTIKIDLGWGDNRFFIDETIFEQLFFVLNLADVLANPEKEVVFRSESDKFDDLVQEMKQLVEACSHEDNVFASNEKFRSEQVTDKVWNIVRKCSDVKQATMLFKNFLQALTYGKIKSHVQEGNKSHLASLIRASKTCDFRMPILERLSTIEMMMEIGVESLRRRIINKFSDTLQFPSDELTFILKTCENDLSTLEGTLHSSVVSLLPITMAMATIHQIFGFLNVKDLVVLPDLARRVLTKYTTGMVEKAKRGETETDYVFETTLPLLRMNKEAFMHKRPRIWTCENTNTVGANVQTRAMTTLELEPSLEHVCRLVNASRPVRPIDEDNRKPTEEERNADYTVSHTIFSYLPKL

>C_remanei_ZWL1

MALKLEDLHNYNTLVATKGEEDEEPTPVLLLNKYRIRLLPLSSLPFVQNYSNISQLSLNSDDVLVIDSPIGATEKYSKKSIENSKKSMIKEEDESMEMEEDKENEGGEIGPLVTSFLTLEKLQKGVNDIEIVGLDCEVQFFDANPIPFEDGVSLQRLRLESSAHFSPAVDKLPIWISTISQNIPSVCWIASGRTQKNVQFSGVTRVIGHFNEINQKLIKQLNEAAGAAQVNRYRAVYDGIRKIATPTRESPGEVIIDMRWNTKSSLVLLEQPDNAADCIIKIDLGWRDNRFFIDESIFEQLFFVLNLADVLANPEKEVIFPVEYVKFGDLVKEMDEIVEACSHEDNVFASNEKFRNEEVTDKVWNIVRKCGDIKHATMLFKNFLQALTYGKIKSHVQEGNKSHLASLIRASKTCDFRMPILERLSTIEMMMEIGVESLRRIINKFSSTLQFPADELTFILKTCENELTTGEGALNSSVVSLLPITMALATVNQIFGLLNEKDYVLLPELARRVLTKFTSGMIEKAKRGETETDYTFETTLPLLRMSKEKFMDKRPRIWTCENMSTVGAHCQTRIMTSLELESSMEHVSRIVNAGRPIREIVETEETQRKPTIEELNADYTVSHTVFSYLPKM

>C_latens_ZWL1

MALKLEDLHNYNTLVAEKGEEDEEPTPVLLLNKYRIRLLPLSSLPFVQNYSNISQLSLNSDDVLVIDSPIGSSEKYSKKSLEISKKSAIKQEDESMEMEEDKENEGGEDGPLVTSFLTLEKLQKGNDVEIVGLDCEVQFFDANPIPFEDGVSLQRLRLESSAHFSPAVDKLPIWISTISSNIPSVCWIASGRTQKNIQFSGVTRVIGHFNESNQKLIKQLNEAAGAAQVNRYRAVYDGIRKIATPTREAPGEVIIDMRWNTKSSLVLLEQPDNAADCIIKIDLGWRDNRFFIDESIFEQLFFVLNLADVLANPEKEVVFPMEYVKFGDLVKEMDEIVEACSHEDNVFASNEKRNEEVTDKVWNIVRKCGDIKHATMLFKNFLQALTYGKIKSHVQEGNKSHLASLIRASKTCDFRMPILERLSTIEMMMEIGVESLRRRIVNKFSSTLQFPADELTFILKTCENELTTGEGSLNSSVVSLLPITMALATVNQIFGLLNEKDYVLLPELARRVLTKFTSGMIEKAKRGETETDYTFETTLPLLRMSKEKFMDKRPRIWTCENMSTVGAHCQTRIMTSLELESSMEHVSRIVNAGRPIREIETDAETQRKPTIEELNADYTVSHTVFSYLPKM

>C_sp51_ZWL1

MPIQLEDLQNYNESVKDWVEGENEPTPILLLNKYRVRLMSVSSLPFVSNYSNISQLGFSSDDVIVIDSAIETFQKEKRIHSLPEASKKQSCAAINEENVSMETEDFEHDKENTAVDEGPLVTSFLTLDKLQKNAVDDVEIVGLDCEIQFFDVNPVPFADGISIQRYLRLESSKHFSTDVDKLPVWISTMSSHFPSVCWLASGRTNKNVQVSGVTRILGYFNESSQKIVKQINEACGAAQVNRYRAVYDGIRKIETPNRNSPGEVIVDMRWNTKSSLVLLEQPDNAADCTIKIDLGWKDKRFFIDEAIFEQLFFVLNLADVLANPEREVVFPSEWAKFDDLVHEMNELVDSCSQEDNVFVSNEKSEEVTDKVWNIVRKCGDVKHATMLFKNFLQALTYGKIKSHVQEGNKSHLATLIRASKTCDFRMPILERLSTIEMMMEIGVESLRGRIINKFSSTLQFPSDELAFILKTCENDLSTGEEALNASVVSLLPITMALATVSQIFGLLNEKDHVILPDLARRVLTKFTSSMVEKAKRGETETEYLFETTLPLLRISKDKFMDKRPRIWTCENTNTVGANVQTRIMTTLELESPVEHVCRLVNESRPIRETKEENEKPSVDELNADYTVSHTVFSYLPKM

>C_sp44_ZWL1

MSLKLEELQSYNDAVKSRKEDEAEPAPLLLLNKYRVRLMPISSLPFISNFSNISQLGFNSDDIIVIDSPIGIQKEKKISFVDSAAKQNDMADSMETEEAHEDHDKENTVTHGGPLETSFLTLTKLQKNAVDDIEIVGLDCEVQFFDVNPIPFADGVSVQRYLRIESSNHFSPTVDRLPVWISTSTSHFPSVCWLASGRTNKNIQVSGATRILGYFDENSQKIVKQMNEACGAAQVNRYRAVYDGIRKIVTETRKYPGEVIVDMRWNTKSSLVLLEQPDNAADCTIKIDLGWKDKRFFIDDAIFEQLFFVLNLADVLANANTEKPVEFPLEWAKFDDLVQEMNDLVDACSQEDNVFVANEKSEEVTDKVWNIVRKCGDVKHATMLLKNFLQALTYGKIKSHVHEGNKSHLASLIRASKTCDFRMPILERLSTIEMMMEIGVESLRGRIINKFSSTLQFPSDELSFILKTCEKDLSTGEEALNASVVSLLPIAMALATVCQIFGLLNEKDHVILPDLARRVLTKFTGSMVEKAKKGELETNYTFETTLPLLRMSKDKFMDKRPRIWICENTNTVGANVQTRTMTALELESSVEHVNRLVNASRPIREMKEENEKPTAEELNADYTVSHTVFSYLPKM

>C_sp48_ZWL1

MSLKLNDLQSYNEQVESWVDGEEKPTPLLLLNKYRVRLMPVSSLPFVSNYSNISQLGFNSDDVLVVDSTVELFQKEKRSSVLNDATKHNIITIDDSMETEDFEQNKENSTVDGGPLVTSFLTLDKLQKNAVDDVEIVGLDCEVEFFDVNPIPFADGVSIQRYLRLESSKHFSPAVDKLPVWISTISSHFPSVCWLAAGRTTKNIQVSGVTRVLGYFNENSQKVVKQMNEACGAAQVNRYRAVYEGIRKIATPTRDAPGEVIVDMRWNTKSSLVLLEQPDNAADCTIRIDLGWRDKRFFIDEAIFEQLFFVLNLADVLANPEKEVVFPSESAKFDDLVQDMNELVDSCSQEDNVFVSNEKSEEVTDKVWNIVRKCGDVKHATMLFKNFLQALTYGKIKSHVQEGNKSHLASLIRASKTCDFRMPILERLSTIEMMMEIGVESLRGRIINKFSSTLQFPNDELSFILKTCENDLSTVEESLNASVVSLLPIAMALATVSQIFGLLNEKDHVILPDLARRVLTKFTSGMVEKAKKGETETEYTFETTLPLLRMSKENFMDKRPRIWTCENTNTVGANVQTRIMTALELESSVEHVSRIVNASRPVREMKEENEKPTAEELNADYTVSHTVFSYLPKM

>C_brenneri_ZWL1

MSLKLNDLQSYNEQVKSWVDGEEKPTPLLLLNKYRVRLMSVSSLPFVSNYSNISQLGFNSDDILVVDSTVELFQKEKRSSILDNAEKRSNIKIDDSMETEDFEQDKENNTVDGGPLVTSFLTLDKLQKNDVEIVGLDCEVQFFDVNPIPFADGISIQRYLRLESSKHFSPAVDKLPVWISTISSHFPSVCWLAAGRTTKNIQVSGVTRVLGYFNENSQKIVKQMNEACGAAQVNRYRAVYEGIRKIATPTRDAPGEVIVDMRWNTKSSLVLLEQPDNAADCTIKIDLGWRDKRFFIDEAIFEQLFFVLNLADVLANPEKEVVFPSESAKFDDLVQDMNELVDSCSQEDNVFVSNEKRSEEVTDKVWNIVRKCGDVKHATMLFKNFLQALTYGKIKSHVQEGNKSHLASLIRASKTCDFRMPILERLSTIEMMMEIGVESLRRRIINKFSSTLQFPTDELSFILKTCENDLSTVEDALNASVVSLLPIAMALATVSQIFGLLNEKDHVILPDLARRVLTKFTSGMVEKAKKGETETEYTFETTLPLLRMSKENFMDKRPRIWTCENTNTVGANVQTRIMTALELESSVEHVSRIVNASRPVREMKEENEKPTAEELDADYTVSHTVFSYLPKM

>C_wallacei_ZWL1

MAIKLEELQNYNEAVNAWEEGKEIPTPILLLNKYRLRLMSISSLPFVSNYSNVSQLGFNSEDVLVIDSPIDSPEKRKTSTLFRNSSKQSDAETKNNDENMETEETEEDDKENSVTHGGPLANDIEIEGLDCEFNFFDANPIPFADGISLQRFLRIESSNNFSPAVDKLPVWISTISPTFPSVCWLASGKTNKNIQVSGATRILGYFDENSQRTVKQIHEACGAAQVNRYRAVYDVIRKISTPNRSAPGEVIVDMRWNTKSSLVLLEQPDNAADCTIKIDLGWRDNRFFIDDAIFEQLFFVLNLADVLANPEKEVVFPSEWAKFEDLVQEMNELVDACSKEDNVFASNEKSEQVTDKVWNIVRKCADVKHATMLLKNFLQALTYGKIKSHVQEGNKSHLASLIRASKTGDFRMRILERLSTIEMMMEIGVESLRERIINKFSSTLQFPRDELIYILESCENDLSSTGEDALNTSAVSLLPITMALATVTQIFRLLNEKDEVILPDLARRVLTKFTSSIVEKAKKGETETDYIFETTLPLLRMSKDTFMYKRPRIWTCENTNTVGANVQTRMMSTLELESSVEHINRMVNASRPHRELKEENAKPTAEELKADYTVSHTVFSYLPKL

>C_doughertyi_ZWL1

MPLKLEELQSYNEAVDAWRDGDDFPTPVLLLEKYRLRVLPLSSLPFVSNYSNISQLGLNSEDVLVIDSPVESPEKQKRSSRSETDVKPSPMIKQESLSMETDDSGEDKENTVADGGPLVTSFLTLEKLQQNNDVNIVGLDCEVEFFDINPIPFEDGVSLQRLRLESVKYFAPTVEKLPVWISTMSSNFPSVCWLAAGRTNRNVQFAGVTRILGYFDETSQRTVKQMNEACGAAQVNRYRAVFDGIRRIATPTRSSPGEVIVDMRWNTKSSLVLLEQPDNAADCTIKIDLGWRDNRFFIDEAIFEQLFFVLNLADVLANPEKEVVFPSEWAKFDDLVQEMKLLVDICSKEDNVFASNEKFRSEEVTDKVWNIVRKCGDVKHATMLFKNFLQALTYGKIKSHVQEGNKSHLAALIRASKTGEFRMPILERLSTIEMMMEIGVESLRSRIINKFSSTLQFPSDEVAFILKSCEAELSNGDGALNSNVISLLPITMALATVNQIFGLLNEKDHVILPDLARRVLTKFTGSMIEKAKKGETETEYTFETTLPLLRMSKDKFMDKRPRIWTCENTNTVGAHVQTRMMTSLELESSVEHVNLMVNASRPIRDVKEGNEKPTAEELKADYTVSHTVFSYLPKM

>C_sp54_ZWL1

MSVKLEELQSYNEAVIAKGEDEKAPAPILLLDKYRLRLVTLSSIPFVSNYSNISQLGLNSDEVLVIDSSTELMGKEKKSSIFEKMPIKREEEHMETDDIEDDKENAATEDGPLVTSFLTLDKLQKDAVNDVEIVGLDCQVQFFDVNPIPFEDGVSLQRYLRLKSSEHFSATVDKLPVWISTISSNFPSVCWLSSGRTNKNVQISGVTRILGHFNENSHNRYRAVYDGIRKIATPTREAPGEVIIDMRWNTKSSLVLLEQPDNAADCTIKIDLGWRDKRFFIDEAIFEQLFFVLNLADVLANPEKEVVFPSEWPKFDDLVQEMNDLVEACSHEDNVFASNEKSEEVTDKVWNIVRKCGDVKHATMLFKNFLQALTYGKIKSHVQEGNKSHLASLIRASKTCDFRMPILERLSTIEMMMEIGVESLRGRIINKFSSTLQFPSDELTFILKTCENELSSGEGLLNSTVISLLPITMALATVNQIFGLLNEKDHVILPELARRVLTKYTSSMVEKSKRGETETDYTFETTLPLLRMSKDKFMDKRPRIWTCENANTVGANVQTRMMTALELESSLEHVNRIVNASRPVRDIKENEKPTVGELNADYTVSHTVFSYLPKI

>C_inopinata_ZWL1

MITLEQLQKYNEETISKKEDDEPAPVLLLNANRVRLMQLSSLPIVSNYSNISQLGLNSDNVIVIDLPIGSAEKTKTKASALENRKLPEQHEQMEMEEDKENTATEGGPLITSFLTLDKLQDIEIVGMDCEEQFFEGNPIPFEDGISLQRLRLNASKHFSPSIDKLPVWILTTSSHFPSVCWLSSGRTNKNVQFTGATRVLGHFDEKIHKLVKQMNEACGASHFNKYRAVYDEIRKVATPTRVSPGEVFIDVRWNTKSSLVLLDHPDNSADCTIKIDLGWKDKRFFTDEAIFEQLFFVLNLADVLVNPEKEVVFPSQYAIFDDLVQEMNVLIEASSHEDNVFASNEKFRNEEITDKVWNIVRKCGNLNHATMLLKNFLQALTYGKIKSHAQERNKSHLALLIRASKTCEFRMPILERLGTIAMMMEIGVESLKRRIINKFSNTLQFPSDELAFILKSCERDLANGEITLNSSTVSLLPITMAFATVNQIFALLNEKDHVILPDLARRVLTKYTSSMIEKAKRGETETEYSFETTLPLLRMCKERFSEKRPRIWTCENTCTVGANVQARTMTLLELESSLEHVTSLVNAHRPVREMKKEDEKASREDLEADYTVSHTVFSYLPKM

>C_elegans_ZWL1

MPLTIEQLKQYNEKMAAKGEEDDAPAPVLLldkyrvrvlpltsipfvmnhssvsqlslssedvlvvdfpskgtgsgagkaekkmifkkkiepmedsfedkenvneggplmtsfltleklrkgmVgdveiigyecetsffdanpiplsdgvslqrYirlncsehfspsistlpvwisttsskfpsicwlaagrtnrnvqfaaatrvlghfnnenservvkqlnqacgtsqlnkyravyeeirkiatenrpapgevtidvrwstkstlvllehpdnaadctikidlgwgdkrffvddeifeqlffvlnladvlanpdnevifpmappanfddlvqemdalveassrednvfvsneNfrggditdkvwnivrkcndvkqvtllfrnflqalaygkikshaqernkshlasliriskssefkipvlerlstidmmmeigveslrRrvidifssqllypsdelefilqtcendlpagngamnsaaisllpitmalatanriyellnekdhvilpdltrrilqkytasmieksrrgeteteytfettlpllrmykegfmskrpciwtcensntvganvqarvltslelqpslehinrlvndsrpvwtateekpamikdlnadytvvhtifsylpkm

>C_oiwi_ZWL1

MLLALEDLHNYNRGIERMSEDENEEEPEPLLLLNKYRLRLIPIDSLPIVSNHSNINHLGLNSEDALVIDLPVGPSKKETKLSVSESFSTKENQTVKMEGVEENKENASRNGGPLETSFLTLDKLQKGSGNDDEVEIVGFDSEVDFFATNPIPFSDGVSLQRYLRIASKEHFSPLVDKLPVWISTIGSDFSSVCWLSSGRTIKDGRFSAITRVIGHLNEHTEKLVKQLNEACGTAQSNRYRCIYDGIRRISNSTREAPGEVIIDMHWNKKSSLVLLEQPDNAADCTIKIDLGWRDNRFYIDEAVFDQLFFVLNLADVLANPEKEVIFPTELVKFDDLVKEMNELVEACSHEDKVFASNEKDDEVTDKVWNIVRKCGDVKHATMLLKNFLQALTYGKIKSHVQEGNKSHLASLIRASKTSDFRMPILERLSTIEMMMEIGVGYLRGRVINKFSSTLHFPSDELKFILTKCENDLSKDEGSLNSSVVSLLPITMALATVNQIFGLLNEKDHVVLPELTRRVLAKYTSSMVEKAKKGETETDYTFEVTLPLLRISKERLMDKRPRIWTCENTNTVGGNAQTRVMIALELESSLEHVNTLINASRPVREAKENDEKPNKEEMNADYTVSHTVFSYLPKM

>C_kamaaina_ZWL1

LNKYRLRLIPIDSLPIISNHSNINHLGLNSEDALVIDLPVGLSKKEKKLAVSESFSPKENQSMKMEGVEENKENTSGNGGPLETSFLTLEKLQKGEVEIVGFDSEVEFFATNPIPFSDAVSLQRYLRMASKEHFSPLVDKLPVWISTIASDFPSVCWLSSGRSSKDGRFSGITRVIGNLNEQTEKLVKQLNEACGTAQSNRYRCIYDGIRRISNSTREAPGEVIIDMHWNKKSSLVLLEQPDNAADCTIKIDLGWRDNRFYIDEAVFDQLFFVLNLADVLANPEKEVIFPTEFVKFDDLVKEMNELVEACSHEDKVFASNEKFRDDEVTDKVWNIVRKCGDVKHATMLLKNFLQALTYGKIKSHVQEGNKSHLASLIRASKTSDFRMPILERLSTIEMMMEIGVGYLRRRVINKFSSTLHFPSDELKFILNKCENDLSKDEGSLNPSVVSLLPITMALATVNQIFGLLNEKDHVVLPELTRRVLAKYTSSMVEKAKKGETETDYTFEVTLPLLRISKERLMDKRPRIWTCENTNTVGGNAQTRVMIALELESSLEHVNTLINASRPMREVKENDEKPNKEEMNADYTVSHTVFSYLPKL

>C_waitukubuli_ZWL1

MSVSVEQLQEYSRAVEEAAEEDAAPTPVLLLDKFRLRLFKITDLPIISNYSNIPQLGLSSDDVLLIDLPTGTEKKAKNVIKSTIPSSVESESEEVMEGIEEDEEDDKENTGGPLVTSFLTLEKLQKGLVDLEIAGFDFEAPKFDVNPVQMSDGISIQRLRLQSSDHFTAALSKLPVWIATSLPSVPSVCWLATGRTAKNLQFSGATRVLGYLNESTNKFVNQLNEACGAASVNRYRAVYDNIRRIATQTRENPGEVIVDMRWHTKNSLVLLEQPDNAAECTIKIDLGWRDKRFYIDESIFEQLFFVLNLADVLANPEKEVVFPNEFEKFDDLVELMNKLVETCSDEENVFVSKDFRSEEVTDKVWNIVRRCGDIKQATLLFKNFLQALTYGKIKSHVLEDNKSHLAALIRASKTCEFRMPILERLSTIEMMMEIGVQSLRRRIVDKFSNTLKFPSDELSLVLKTCENDLKSVECVMNVSVVSLLPITMALATVHQIFGLLNDKDHVILPDLARRVLMKYTSSMVEKSKKGETELNYSFETTLPLLRINKDRFMDKRPRIWTCENSKHVGANVHARMMTCLELETSLENVSSLVNSIRPVRETKEDEKPTVEELNAEYTVSHTVFTYFPKI

>C_panamensis_ZWL1

MLVGIEHLQEYNKAVESKTEEDEAPVPVLLLDKFRLRLIPITDIPIILNYSNIPQLGFSSDDVLLIDSPTGSEKASKITKTLKSVKVDAEEEEQMEVEEEEGDKENTATEGGPLQTSFLSLNKLQKRAEMEIDGFDFQAPKFDVNPIPLGDGISIQRSLRLESSNHFSAALSKLPVWIATTDSNLPSACWLATGRTGKNVQFSGATRILGYYNETTDKFVNQLNEAGGGALVNRYRAVYDRIRRIATKDRENPGEVMVDMRWNTKNSLVLLEQPVNAAECIIKIDLGWKDKRFYIDESIFNQLFFVLNLADVLANPDNEVVFPNESVKFDNLVKLMNELVELCSDEENVFVAKDFRSEEVTDKVWNIVRCCGDIKQATMLFKNFLQALTYGKIKSHVLEDNKSHLAALIRASKTCEFRMPILERLSTIEMMMEIGVQSLRRQIVDKFSKTLKFPSDELALVLKTCENDLKSVDCNLNASVVSLLPITMALATVNQIYGILNEKDHVILPDLARRVLMKYTSSMVEKCKRGETEVNYSFETTLPLLRINKEKFMDNRPRIWTCENSKHVGANVQTRMMTSLELETALDHVCSLVNSVRPTSEAKDNETRTAAELNADYTVSHTIFTFSPKV

>C_nouraguensis_ZWL1

MLIGIEQLKQYNAAVESKAEDDEAPAPVTLLDKFRLRLMPITDLPIISNYSNIAQLGFYSDDILLIDAPTGSAKKAKIAHMVLPTRQDNSEAMEVGDEEEDKENSAMEGGPLQTSFLTLEKLQRDVEIAGFNFEAPKFDVNPIALFDGISIQRLRLESTSIFSAAVSKLPIWIATTQSDFPSMCWLAAGRTNKNLQFSGATRVLGYYNETTDKFVNQLNEACGGALVNRYRAVYDGFRRIATQTRQHPGEVMMDMRWNTKNSLVLLEQPVNAAECTIKIDLGWKDERFYIDESIFEQLFFVLNLAEVLANPEEEVKFPNKWEKFDDLVKLMNELVLKCSDEENVFASKDTSEEVTDKVWNIVRRCRDIKEATLLFKNFLQALTYGKIKSHVLEDNKSHLAALIRASKTGEFRMPILERLSTIEMMMEIGVQRLRIVDKFSKTLKFPSDELTLMLKSCENDLKSVECSMNASVVSLLPITMALATVHQIFGILNEKDHVILPDLARRVLMKYTSSMIEKSKRGETELNYSFETTLPLLRINKDMFVDKRPRIWTCENSKHVGANVQARMMTSLELEMSLEHVSDLVNSIRPLRETKQDKIPTTEELNAEYTVSHTVFTYYSKV

>C_becei_ZWL1

MLVGIEQLQQYNTKVESKTEDDEAPAPVLLLEKFRLRLMPITDLPIISNYSNIAQLGFTTDNVLLIDAPARSAKKAKIAQVPQLTKRDDVEVMEVGDEEEEDKENTAMEGGPLETSFLTLEKLQKSSVDVEIAGFNFEAPKFDVNPVALCDGISIQRYLRLESSAVFTAAVSKLPVWIATTQSDFPSMCWLSAGRANKNLQFSGATRILGYYNEATDKFVNQLNEACGGALVNRYRAVYDGFRRVATQSRQHPGEVIMDMRWNTKNSLVLLEQPVNAAECTIKIDLGWKDERFYIDESIFEQLFFVLNLAEVLANPEEEVKFPKKWEKFDELVKLMNELVQKCSDEENVFASKDTSEEVTDKVWNIVRRCSDIKEATMLLKNFLQALTYGKIKSHVLENNKSHLAALIRASKTCEFRMPILERLSTIEMMMEIGVQRLRGRIVDMFSKTLKFPSDELTLMLKSCENDLKSVECSMNASVVSLLPITMALATVHQIFGILNEKDHVILPDLARRVLMKYTSSMIEKSKRGETELNYSFETTLPLLRINKDMFVDKRPRIWTCENSKHVGANVQARMMTSLELEMPLEHVSELVNSIRPVRETKEDTAPTTEELNAEYTVSHTVFTYYSKI

>C_yunquensis_ZWL1

MLVGLEHLQQYNTAVKSKTEEDEEPTPVLLLGKFRLRLMPIADLPIVSNYSNIAQLGFSSDDILLIDSPNGSEKKAKVQQVPFSIKQENGEAMEVGEDEDDKENAAMEDGPLETSFLTLEKLQRSSGNEKDIEIAGFDFEVPMFDVNPIPLCDGISIQRYLRLESSSLFTPAVSKLPVWISTTQSEFPSMCWLAAGRTNKNLQFSGATRILGYYNETTDKIVNQLNEACGGALVNRYRAVYDGICRIATQSRANPGEVIVDMRWNTKNSLVLLEQPVNAAECTIKIDLGWKDKRFYIDESMFEQLFFVLNLADVLANPEKEVVFPDKCEKFDDLVKLLNELVETCSDEDNVFVSKDSSEEVTDKVWNIVRCCGDIKQATLLFKNFLQALTYGKIKSHVLEDNKSHLAALIRASKTSEFRMPILERLSTIEMMMEIGVQSLRGRIVDKFSNTLKFPSDELTLILKSCENDLKANECAMNASVVSLLPITMALATVHQIFGILNEKDHVILPDLARRVLMKYTSNIIEKGKRGETELNYSFETTLPLLRIRKDEFMDRRPRIWTCENSKHVGANVHARMMTSLELETSLEHVSGLVNSIRPVIDVEDNKKLTAEELNAKYTVSHTVFTYFSKF

>C_macrosperma_ZWL1

MSVSVEQLHEYNRALETKAEDDEAPTPVLLLNKFRLRLVPITDLPIVSNYSNIPQLGFSSDDVILIDSPTGSEKKQKVSLEPQVVAETDDAMEMGENEEDKENATGGGPLETSFLTLAKLQRSSGKEVNVEIAGYDFEAPTFDVNPIPLGDGISIQRYLRLQSSDVFTAAVAKYPVWIATTEPSLPSMCWLAAGRTGKNLRFSGATRILGYYNESTDKLVNQLNEACGGALVNRYRAVYDGIRRIATPTRENPGEVMVDMRWNTKNSLVLLEQPVNAAECTIKIDLGWRDKRFYIDESIFEQLFFVLNLADVLANPEKEVVFPNEWEKFDDLVKLMNELVEMCSDEENVFVSKDTSEEVTDKVWNIVRRCGDIKQATLLFKNFLQALTYGKIKSHVLEDNKSHLAALIRASKTCEFRMPILERLSTIEMMMEIGVQSLRGRIVEKFSKTLKFPGDELTLMLKTCENDLKSGECAMNASVVSLLPITMALATVHQIFGILNEKDHVILPDLARRVLMKYTSSMIEKSKRGETELNYSFETTLPLLRINKDKFMDKRPRIWTCENSKHVGANVQARMMTSLELETSLDHVCSLVNSIRPIRETAEDESPSAAELNAEYTVSHTVFTYFPK

>C_sulstoni_ZWL1

MAICLEDLKSYQNAIRTKEENDEAPPPVLFQYRLRLIPISELPIVSNYSNFTQLGLNSDEVLVIDSSIESEKKETFSKVKKSAKIQAEQSDSMEVGGDEEEEADKENDTSATGPLVTSFLTLDKLQKVDMEIGDFEFEMPSFDANPIPPEDGISLLRLRLSSPNHFSDAVAKLPIWISTSSDSFPSTCWLSSGRTQKNVQFSGVTHVLGFFNEHSEKLVNQLNETCGGAQVNRYRAIYDGIRRIVTENQKTPGEVIVDMRWNTKNSLVLLEQPVNAAECTIKIDLGWKDKRFFVDETIFEQLFFVLNLADVLANPEKEVVFPSEWEKFDDLVKQMNELIEACSNEDNVFASNDSSEEVTDKVWNIVRRCGDVKQATLLFKNFMQALTYGKIKSHVLENNKSYLAALIRSSKTCDFRMPILERLSTIEMMMEIGVESLRSRIIHEFETILKFPADELSLILKTCETDLINGEGSMNASVISLLPIAMSLATVNQIFGLLNEKDHVILPDLARRVLTKYTNSMVEKSKRGETESHYTFETTLPLLRINKEKFMNKRPRVWTCENSKTVGANVQCRMMTVLELESSLEHVTRVVNAIRPLREENSLDINSDYTVSHTVFSYLPKM

>C_afra_ZWL1

MSICLENLKKYNDAVKAKGEDDEAPAPVLLMSQYRLRVIPITDLPIISNYSNIAQLGLNSDEVLVIDSPVEPEKKEKLLRVRNEERIETGSTESMEIEEEEEDKENTSATGPLVTSFLTLDKLQKHSERMKWKSPDLTLKRPVSMRIRSPRKTEFLFSDNHLSDSLAKLPIWISTSSASFPATCWLSSGRTQKSVQYSAVTRVLGFFDEHSEKLVNQLNEACGGAQVNRYRAIYDGIRRIATEQRETPGEVVVDMRWNTKNSLVLLEQPVNAAECTIKIDLGWKDKRFFIDESIFEQLFFVLNLADVLANPEKEVVFSCEYEKFDDLVKQMNELIEASSNEENVFTSNDTSEEVTDKVWNIVRRCGDVKQATLLFKNFMQALTYGKIKSHVLENNKSYLAALIRSSKTCDFRMPILERLSTIQMMMEIGVESLRSRIINKFESTLKFPADELSLILKTCETDLINGEGSMNASVISLLPIAMSLATVNQIFGLLNDKDHVILPDLARRVLTKYTGSMVEKSKRGETETNYTFETTLPLLRISKEKFMNKRPRIWSCENSKTVGANVQCRMMTVLELESSLEHVARAVNAIRPMRDGQDEETSHETNADYTVSHTVFSYLPKM

>C_sp49_ZWL1

MSITLEQLKAFNDAVDSKKEDDAAPTPVLLLNKYRVRLLPLSDLPIITHHSNLSQLALNSESVLVIDSPVNSEKPECRKLPTKTQNVCSDEKMDIEIIEKDENDKENVGGPLETTFLTLDKLQKGVAVVGDVEIEGFDIAAATFDVNPIGYSEAGVLHRLIRHNPTTHFSSPIHQYPIWISTFTDSLPHICWMSAGITQKNRQVTGITRVLGYLDENSEKLVNQLNASCGAGTVSRYRAVYDEIRKIRTAKLPCPGEVVLEMRWNSNGSVVLLDKPVNAAECTIRIDLGLQDSRFFADKSLYSELFFVLNLANVLAHPENEVLFPSCTEKFDVLVKEINSLVEKCSEDDKVLISGDTEEVTHHVWQIVRRCSDIKQATMLLKNFMQALTYKKIKSHVQESNKSHLASLIRSSKTGDFRMPILERLSTIEMMMEIGAESLRGRIIRNFSEKVAFPSEELKLILEACENDLKNSDVAVNAHAISLLPVAMGLATVSHIHSLLNEKDQVILPDVTRRVLQKYTKGMVEAMKRGENENEYIYETTLPVLRIKKEAFMAKRPRVWSCENMRTVGANVLSRMMTALELECPLSRVVDAVNSIRPHREEGDQNQEKALDLNSDYTVSHTVFSFMPKM

>C_sp25_ZWL1

MPITLEELNTYNEATEAKKEEDEPPVPVLLLALNSESVIVIDSPIRLEKNEARRVAPAPQKTVSDEAMDTEVTENDKENVGGPLETSFLTLDKLQKSGMIVGDIDIKGFDFAAPSFDANPIGYSEAGVLHTVLRHDPTTYFSAPIDQYPIWISTFTESFSHICWMAAGRTQKNRQFTSITRVLGFLDEHTEKLANQLNVACGGGTVSRYRAVYDEIRKIRTAKRPYPGEVVLEMRWNSNSSMVLLEKPVNAADCTIRVDLGWQDSRFFVDESLYTELFFVLSLANVLANPENEVQFQSKSEKFDELVKEINLLVAECADDNKVLVSNETEEVTHHVWQIVRRCGDIKQATLLLKNFMQALTYKKIKSHVQENNKSHLASLIRSSKTGDFRMPILERLSTIEMMMEIGVESLRGRIIKNYAKTVAFPSDELDLVFKNCESDLKNCGVTVNSHAISLLPVAMGLATVSQIHSLLNEKDHVILPDLTRRVLQKYTKEMVDATKRGEEDNEYVFETTLPVLRIKKEAFMTKRARVWSCEKVRTVGANVLSRMMTLLELECSLPRVVDAVNKIRPYEQDEDPNQEKISDSNSDYTVSHTAFSFMPVM

>C_imperialis_ZWL1

MPITLEQLKTYNEAVKTKGEEDEAPAPVLLLNKYRVRLLPLAELPIIENHSNLAQLALNSEEVLVIDSPSAPAALEKQPVKIVTKPVTSDEQMDVIEAGDDVEDKENVGGPLETTFLTLEKLQKGKKIVADVEIEGFDYALPTFDANPIAYTEADALHRILRLQPTAHFCSTIAHFPIWTSTFSSSFPHICWKSSNLTQKGQQVTGVTRVVGFYDEHSEKLVNQLNAACGGSAVSRYRAVYDQIRKISTAKRPFPGEIVLDMRWNTNGSVVLLEKPVNAAECTIRIDLGWQDRRFFADESLYTELFFVLNLANVLANPELDVHFPSSSDKFDALVKEMNALVENCSEDDKVLASSETEEVTNNVWHIVRRCGDIKQATLLLKNFMQALTYKKIKSHVQENNKSHLAALIRSSKTGDFRMPILERLSTVEMMMEIGVESLRGRIVKNFHESVRFASEELELVLKTCETELKNCNASINAHAVSLLPVAMGLATVSQIHALLNVKDHVILPDLTRRVLQKYTKSMVEAAKRGESESGYVFETTLPVLRIKKESFMTKRPRVWTCENVKMGGASVLSRMMTVLELESSLPRVTDAVNSIRPYREEMEESPENMKKQLDVNSDYTVSHTTFSYLPKIQ

>C_kamaaina_ZWL1_Exon1NotFound

LNKYRLRLIPIDSLPIISNHSNINHLGLNSEDALVIDLPVGLSKKEKKLAVSESFSPKENQSMKMEGVEENKENTSGNGGPLETSFLTLEKLQKGEVEIVGFDSEVEFFATNPIPFSDAVSLQRYLRMASKEHFSPLVDKLPVWISTIASDFPSVCWLSSGRSSKDGRFSGITRVIGNLNEQTEKLVKQLNEACGTAQSNRYRCIYDGIRRISNSTREAPGEVIIDMHWNKKSSLVLLEQPDNAADCTIKIDLGWRDNRFYIDEAVFDQLFFVLNLADVLANPEKEVIFPTEFVKFDDLVKEMNELVEACSHEDKVFASNEKFRDDEVTDKVWNIVRKCGDVKHATMLLKNFLQALTYGKIKSHVQEGNKSHLASLIRASKTSDFRMPILERLSTIEMMMEIGVGYLRRRVINKFSSTLHFPSDELKFILNKCENDLSKDEGSLNPSVVSLLPITMALATVNQIFGLLNEKDHVVLPELTRRVLAKYTSSMVEKAKKGETETDYTFEVTLPLLRISKERLMDKRPRIWTCENTNTVGGNAQTRVMIALELESSLEHVNTLINASRPMREVKENDEKPNKEEMNADYTVSHTVFSYLPKL

>C_tropicalis_zwl1_cDNAWithFrameshift

ATGTCTCTGAAAATTGAAGACTTGCAAACGTATAATTTAGCTGTCAATAAAGCGAAAGAAGGAGAAGAGAAACCAGCTCCTGTCCTATTGCTCAACAAGTATCGCATTCGGCTGATCCCAATCTCATCACTACCATTTGTGTCAAACTACTCGAATATTTCTCAGCTCGGGTTCAACTCAGATGATGTTATCGTCATTGACTCACCAAACGAATTAGCTGAAAAACAGAAGACATCTAACCTTTTCTGCAATACCTCGAAGAAAAAATCTACTTCTGTGGAGCAAGAAACTGAGCAGATGGAAACCGATGAACCTGAAGAAGACAAAGAGAATATTGATGCCCATGACGGGCCATTGGCAACATCTTTTCTTACTTTAGACAAATTACAAAAGGACATGGAGATTGCTGGGTTGGACTGTGAGGTCCATTTTTTCGATGTGAATCCACTCCTATTTACAGACGGTGTTTCTCTCCAGAGACTTCGAATTGAAAGCTCTAACTATTTTTCACCACCGGTCGACAAACTTCCTGTTTGGATTTCCACTATCAGTCCGAACTTTCCATTAGTTTGCTGGTTAGCTTGTGGAAAAACCAGCAAAAATGCACAAGTTTCGGGGACAACTCGAATTCTCGGATTTTTCGATGAAAGTAGTCAGAAAACTGTGAAACAAATACAGGAAGCATGTGGTGCTGCTCAAGTAAACCGGTACCGTTCTGTTTACGATGTTATTCGAAAAATTCCCACAACAACTCGCCCTGCACCCGGAGAAGTTATAGTAGACATGCGTTGGAACACAAAAAGTAGTCTCGTTCTTTTGGAACAACCAGATAATGCTGCCGATTGTACCATCAGAATTGACCTTGGTTGGAAGGACAACCGTTTTTTCATTGATGATGCTATCTTTGAACAGCTTTTTTTGTGCTTAATCTGGCCGAAGTTCTGGCTATTCCCGATAATGAAGTTACCTTCCCTTCGGAGTGGACAAAGTTCGAAGATCTTGTTCAAGAAATGAATGAACTCGTTGACGCTTGCTCCAAGGAAGACAACGTGTTTGCCTCAAATGAAAAGTTCAGAAGTGAACAAGTCACCGACAAGGTATGGAATATCGTTCGCAAGTGCGCGGATGTTAAACACGCGACAATGATTCTCAAAAACTTCTTACAAGCGTTGACTTATGGAAAGATCAAATCGCATGTACAGGAAGGAAACAAAAGCCACCTGGCCTCATTGATTCGTGCTTCGAAAACTGGTGATTTTAGAATGCCTATCCTCGAACGATTGAGTACTATAGAAATGATGATGGAAATCGGTGTAGAAAGTCTGAGACGTATCGTCAACAAATTTACGAGCACTCTCCAGTTTCCACGTGATGAACTTATTTACATCCTGGAAACTTGTCAAAACGATTTGTCGTCTACTGGTGAAGAAGCTTTGAATACAAGCGCTGTATCTCTTCTTCCAATTACCATGGCATTGGCAACTGTCACTCAAATTTTCCGACTTCTCAATGAAAAGGATCAAGTTATTCTACCAGATTTGGCAAGACGCGTTCTTACTAAATTCACAAGCGCCATGGTCGAAAAAGCAAAGAAAGGTGAAACTGAAACCGACTACATATTCGAGACAACTCTTCCTCTTCTTCGTATGAGTAAAGATACATTTATGTACAAAAGACCACGAATTTGGACTTGTGAAAATACAAATACTGTTGGAGCCAACGTACAAACTCGTATGATGACAACACTCGAGTTGGAGCCGTCGGTTGAACATATCAGTCGTATGGTTAATGCGAATCGACCACTTCGTGAGCTGAAAGAAGAGAATGTGAAGCCAACTGAAGAAGAACTCAGAGCGGACTACACAGTATCGCATACAGTTTTCTCCTATTTACCAAAGCCGTGA

**Supplementary Data S7: All SPDL-1 sequences used in this study**

>C_tribulationis_SPDL1

MPDDEEKLQLRADVERFKKAIRQKDEMIEEMEHELNNIGKTPQSDGRAEARERELAGTIRDLQFEMDGKDATIHGQTDIIASMREEIDKLEKKNRELINRPDCSEIDESNSFVESEMLRISEECEKFKELANTLGEENLDLKRAALELKEEYESACEHVKCLESHSKTKEEEIMRLEGEVFDLKNSTAGKFSNTGNSIFAEAMEAEKKLEEDLKVLYREKQSLMGMVKRLNLEKEDAEQRARSNMNRGLVVRNAINHIEVQELNRLNTRLRQLETERFQFWEKMFIKMKSVPKRELGSMFQGYFESFKCSITNMQSGYDELMKKNEQHITTIRGLQQEIETLRVKNEQLTFDVELLERKVRSAGNCELDNPQLRAPLKPMNNARPSFFVKPKKADPAPPLETSLSNMMMTPQKPSPQPMCRSTAKKEEASEWAERKLKAKAEKKSATPAPRYNFVKMSAPVSSTKFKPAILQMPSTPSAIPELEN

>C_sp41_SPDL1

MPDDEEKLQLRADVERFKRAIRQKDEMIEEMEHELNNLGKVPQSDGRAEAREKELAGQIRDLQFEMDGKDATIHNQTDLIASMRLEIDRLEKNLELINRPDCSDVDESNSFVESEMIRISEECEKFKELVNTLGEENLELKKSAIELKEEYENACDHVKSLESHSKTKEEEIMRLEGEVFDLKNSATGKFSSTGNSIFEEAMEAERKLEEDLKVLFKEKQSLMGMVKRLNIEKEEAEERARSNMNRGLVVRNAINHIDVQELNRLNTRVRQLETERFQFWEKMFIKIKSIPKKELGSMFLGYFESFKCSIANMKGGFEELMKKNEQQIITIRGLQQEIETLRSKNEQLTFDFEILERKVRSAENRELENPQLRAPLKPMNNARPSFFVKPKKADPAPSLEASMSNMMMTPQKPSPQPTCRSTAKKEETSEWAERKLKAKAEKKAATPAPRYNFVKLSAPISNTKFKPAILQMPSTPSAPSELEN

>C_zanzibari_SPDL1

MPDDEEKLQLRADVERLKRAVRQKDEMIEEMEHELNNYGKIPESDGRAEAREKELAGQIRDLQFEMDGKDAAIHDQTDIITSLRVEIDKLEKTNRELVNRTDCSEGDDSSSFVESEMLRISEECEKFKELANMLGEENLELKREAVELKENYESACDHVKSLESHSKTKEEEIMRLEGEVFDLKNSTTGKFASTGNSIFAEAMDAERKLEEDLKVLYREKQALMGMVKRLTIEKEEIEERARAQMNRGLVVRNAINHIDVQELNRLNTRIRKLETERFQFWEKMFIKIRSVSKKELGAMFLGYFESFKCSITNMKDGFDELMKKNEQHITTIRGQQQEIETLRMKNEQLTFDVECLERKVRSAGNCELENPQLREPLKPMNNARPSLFVKPKKTDSVPPLETSMASMMMTPQKPSPQPTCRSTAKKEETSEWAERKMKSKAEKKSATPAPRYNYVKMSVPVSNNRFKPATLTMPSTPSTHPELEN

>C_sinica_SPDL1

MPDDDEKLQLRADVERYKRAIQQKDEMIEEMEHELSSYGKAPESNGKAEAREKELAVKIRDLQYEMDGKDATIHDQTDLISSLRVEIDKLEQNRELINRHDITESDESNSFVESEMLRISDECEKFKELANMLGDENLELKKAAVELKEEYESACDHVKILESHSKTKEEEIMRLEGEIFDLKNSNTGKFSNSGNSIFAEAMEAEQKLEEDLKVLFREKQALMGMVKRLTIEKEEAEERARSHMNRGLVVRNAINHIDVQELNRLNTRVRKLETERFEFWQKMFIKTKSIPKKELGSMIVGYFESFKCSIASMTGGYEELMKKNEQHITTIRGLQQENETLRVKVEQLTFDVEILERKVRSAGNCELDNPQLREPLKPMNNARPSFFVKPKKADPAPPLETSMSNMMMTPQKPSPQLACRSTAKKEETSEFADRWMKAKSEKKAAAPSRYNFVKMKVPVPANKFKPPVLQMPSTPSADPELEN

>C_nigoni_SPDL1

MPDDEEKLQLRADVERFKRAIRQKDEMIEEMENELNSYGKQPVSNGKAEAREKELAGQMRDLQFEMDGKDATIHEQTELISSLKVEIDKLEKTNRELMNRSVCVDSDESNLLDESQMLRISEECEKLKELASALGEENLELKKDAVTLREDYESAVEHVKSLESHSRTKEEEITRLEGELFDLRNSCTGKFANTGNSIFAEAMDAERKLEEDLKTLYREKQSLMGMVKRLTMEKEEAEERARSHMNRGLVVRNAINHIDVQELNRLNKRVRQLETEKSHFWEKIFIKMRSIPKKELGALIVGHFGAFKCSIENMSEGYDELMKKNEQHVTTIRGLQQEIDTQRVKIEQLTFDVECLERKLRSSAADSEPEDPQLRAPLQPITNAARPSFFVKPKKTNPEPSLEMSMSNMMMTPQKPSAQITCRSTVKKEDDELSEWAERRLKAKAEKKSATPAAKYNFVKLTAPAPSNKFKPAVLQMPSIQSENTDEHEQ

>C_briggsae_SPDL1

MPDDEEKLQLRAdverfkrairqkdemieemenelnsygkqpvsngkaearekelagqmrdlqfemdgkdatiheqtelisslkveidklekTnrelinrsvcvdsdesnlfdesqilriseeceklkelasalgeenlelkkdavtlrenyesavehvksleshsrtkeeeitrlegelfdlrnsstgkfatagnsifaevmdaerqleedlktlyrekqslmsmvkrltmekeeaeerarshmnrglvvrnainhidvqelnrlnkrvrqletekshfwekifikmrsiskkelgalivghfgafkcsienmsegydelmkknehhvttirglqqevdthqvkieqlkfdveclerklrssaansepenpqlraplqpitnatrpsffvkpkkadpepsleismssmmmtpqkpsaqitfrstvkkeddelsewaerrmkakaekksatpaakynfvkltapapsnkfkpavlqmpsiqsenteehdnkcqifsiyefl

>C_remanei_SPDL1

MPDDDEKLQLRADVERYRKAIRQKDDMIEEMENELNHYGKPAISDGKAEAREKELNGRIRDLQFEMDEKDAAIHDQTDLISSLRSEIEKLEKTNRELINRSECNESDESNSFVESEMIRISEECEKFKEVASTLYEQNRELKKEAVELREEHDSAMGHVKNLESHIKTNEEEIARLEGEVFDLKNSNQGKHASTGNSIFAEVMEAEQKLEEDLKVLFREKQSLMSMVKRLTMEKDEAEERARSFMNRGLIVRNAINHIDVEEMRRLSARNRELETERTHFWERMFIKMKTVPKKEIGAIIVGYFESFKCSIASIKGGFDDLMKKNEQNLTIIRGQNQDLENQRVKIEQLQFDMECLERKLRSAVKAESEDSQLRAPLKPMNNSRPSFFVKPKKTDPVPPLETSMSNMMMTPQKPSPVITARSTAKKEDSSEWAERRLKAKAEKKSATPTPRYNYVTMSAPVPKFKAAVLQMPSTLPESQEN

>C_latens_SPDL1

MPDDEEKLQLRADVERYRKAIRQKDDMIEEMENELNHYGKPAISDGKAEAREKELNGRIRDLQFEMDEKDAAIHDQTDLISSLRSEIEKLEKTNRDLINRSDCNESDESNSFVESEMIRISEECEKFKEVASTLYEQNLELKKEAVELREEHDSAMGHVKNLESHIKTNEEEIARLEGEVFDLKNSNQGKHASTGNSIFAEVMEAEQKLEEDLKVLFREKQGLMSMVKRLTMEKDEAEERARSFMNRGLIVRNAINHIDVEEMRRLSARNRELETERTHFWERMFIKMKTVPKREIGPIIVGYFESFKCSIASIKGGFDELMKKNEQNLTIIRGQNQDLENQRVKIEQMQFDMECLERKLRSAAKAEIEDSQLRAPLKPMNNSRPSFFVKPKKTDSVPPLETSMSNMMMTPQKPSPAITARSTAKKEDSSEWAERRLKAKAEKKSATPTPRYNYVTMSAPVPKFKAAVLQMPSTLPESQEN

>C_sp51_SPDL1

MLDDEEKLQLKADVERFKKVIRQKDEMIEEMENELNSGKTTETDGKAEAREKGLNEQIRNMQFEIDGKDARIHEQTGLISSLRHEIDELEKSNREFINRSEFNESDESNSSVETEMLRISEECEKFKEVAATLYEQNSDLRKEVVELKEDHETAIEHVKSLEIHIKTKEEEIMRLEGELFDLKNSNHGKHASSGNSIFAEAMEAERKLEADLKVLFREKQSLLVTVKRLTVEKEDAEERARAFMNRGLVVRNAINHIDVEEMRRLRARVQHLESERIHLWERLFIKIKSVPKREIGSIMAGYFDSFKCSIASVTGGFEELMKKNEKYVTTIRGLQQENENQRVKIEQLQFDMECAERKMRSALNGDSENPQLRAPLKPLDNSRPSFFVKPKKSETVAQLETSMSSMLMTPQKPSVAMTASSTAKEEDASDWTERRLKAKAEKKSATPAVRYNFITLKAPAPSSKFKAAVLPMPSTPTEPKEN

>C_sp44_SPDL1_paralog1

MLDEEKLQLRADVERYKKAIRQKDEMIEEMENELNSHGKGATDEKAEAKEKELNGQIRDLQFEIDGKEATIHEQTDLISELRNEIDKLEKSNRELINRSDFIIESDESNSFGETEMLRLSEECEKFKEESTNLYEENLKLKKEVVELKEDHNSAMEHIRSLESHIKTKEEEIAMLEGEVFDLKNSNHGKHASSGNSIYLEAMEAERKLEEDLKVIFREKQSLMAMVKRLTLEKEDAEERARAFVSRGLVVRNAINHIDVAEMQRLRTRVNELETERIHLWERFFIKMKTVPKREIGSMMAGYFESFKCSIASVTGGFEDLMKRNEQYVTTIRGLQQETENQRVKIEQLQFDIECAERKMRSSSNADSEHPQLRAPLKAVDNSRPSFFVKPKKVEPVPQLETSMSNMLMTPQRSSEPIAETSTAKKEDMSDWAERRLKAKAEKKSATPAARYNYIKLTAPASSSKFKAAVLPMPSSSSDSKEN

>C_sp44_SPDL1_paralog2

MNSSSISEKPGRAKELTDAIITEIQSYEGKLKQTPFMKQVCAKLLALEMPETVGEQWEKKLANRTKNEIQWIRDTNLIPQMLKDEQISNDNKLKLGTIFGTAVIQSETPEDGHGPTDDSFHEQTESDESNSFGETEMLRLREECEKFKEESAKLYEENSKLKKEVVELKEDHNSAMEHIRSLESHIKTKGEEIAMLEGEVFDLKNSNHGKDASSGNSIYLEAMEAERKLEEDLKVIFREKQSLMAMVKRLTLEKEDAEERARAFVSRGVVVRTAINHIDVAEMQRLRTRVNELETERIHLWERLFIKMKTVPKREIGSIMAAYCESFKCSTASVTGGFEDLMKRNEQYVTTIRGLQQETENQRVKIEQLQFDIECAERKMRSSSKADSEHPQLPAPLKAVDNSRPSFFVKPKKVEPVPQLETSISNMLVTPQRSSEPIAETSTAKKEDMSDWAERRMKAEAASSSEFKADVSPMPSSSSGSKEH

>C_sp48_SPDL1

MSTKMLDDDEKLQLRADVERFKKAIRQKDEMIEEMENELNNHGKASESDVKAEAREKELNVKIRDLQFEMDGKDVTIHEQTDLISSLRNEIDKLEKSNRELANRSDTNESDESNSFVETEMLRISEECEKFKEVAATLYEQNLELKKEAIELKEEHDAAVEHVKSLESHIKTKEDEIARLEGEVFDLKNSNHGKHASSGNSIFAEAMEAERKLEEDLKTIFREKQALMAMVKRLTVEKEDAEERARAFMTRGLVVRNAINHIDVEEMRRLRARVQQLETERTHFWERLFIKMKTVPKREIGSIMAGYFESFKCSIASVTGGFEEIMKKNEQYVTTIRGLQQENENQRVRIEQLQFDIECAERKLRSAVNSDSEHPQLRAPLKPMDNSRPSFFVKPKKAEPVTQLDISMSNMLMTPQKPAAMITSNSTAKKEDASDWAERRLKAKAEKKSATPAARYNYITLKAPSAATKFKAAVLPMPSSDSDLKEN

>C_brenneri_SPDL1

MLDDDEKLQLRADVERLKKAIRQKDEMIEEMENELNNHGKASESDGKAEARERELNGKIRDLQFEIDGKDVTIHEQTDLISSLRNEIDKLEKSNRELANRSDTNESDESSSFVETEMLRISEECEKFKEVAGTLYEQNLELKKEAIELKEEHDAAVDHVKSLESHIKTKEDEIARLEGEVFDLKNSNHGKHASSGNSIFAEVQAMEAERKLEEDLKKIFREKQGLMAMVKRLTLEKEDAEERARAFMARGLVVRNAINHIDVEEMRRLRARVQQLETERTHFWERLFIKMKTVPKREIGSIMAGYFESFKCSIASVTGGFEEIMKKNEQYVTTIRGLQQEIENQRVKIEQLQFDIECAERKMRSVLSSDSENPQLRAPLKPMDNSRPSFFVKPKKAEPVTQLDISMSNMLMTPQKPAAMITSNSTAKKEDASDWAERRLKAKAEKKSATPAARYNYITLKAPSAATKFKAAVLPMPSSDSDLKEN

>C_wallacei_SPDL1

MPEDEEKLQLRADVERFKRAIRQKDEMIEEMENELNCRGKSPVSNSKAEAREKELNGQIRDLQFEMDGKETTIHEQTDLINSLRDEIDKLEKTNRELVNRSDCTESDESNSFVETEMLRISEECEKFKEVAASLYEQNVELKKEAVELREEHDAAVEHLKSLESHIKTKEEEITRLEGEVFDLKNSSQGKHATSGNSIFAEAMEAERKLEQDLKVLFREKQSLMAMVKRLTLEKEDAEDRARNFIDRGRAVRQAMSHIDVEEMRRLRTRVQQLESERSHLWERLFIRMKNVPKRELGSLVASYFESFRCSMASVTGGFEELMKKNEQYVTTIRGLQQEIENYRVKVEQLEFDNECSQRKLRLSANIEQENPQIRAPLKPMNNSRPSFFVKPKKIDPVSTLETSMSSMLMTPQKPSVPTTASSTAKKEASDWAERREKSKAEKKSATPAQHFNFVKLAAPVPKFKAAVLQMPSSPTELNEN

>C_tropicalis_SPDL1

MPDDEEKLQLRADVERFKKTIRQKDEMIEEMENELNNRGKSPVSNGKAEAKEKELSSQIRDLQFEMDGKDATIHEQTDLINTLRDEIDKLEKTNRELINRSYCAESDESSSFVETELLRISEECEKFKEVASSLFEQNVELKKEAVELREEHDAAVDHVRSLESHIKTKEEEITRLEGEVFDLRNSSQGKHASSGNSIFAEMEAERKLEEDLKLLFREKQSLMTMVKRLTLEKEDAEDRARNCIDRGRVVRQAMSHIDVEEMRRLRTRVHQLESERTHLWERLFIKMKSIPKRELGSLIASYFDSFKCSIASMKGGFEELLKKNEQYVTTIRGLQQELENQRGKVEQLEFDVECAERKLRSAANVELENPQLRAPLKPMNTARPSFFVKPKKIDSTTSLETSMSSMLMTPQKPYNPIAASSTAKTEASDWAERRQKAKAEKKSATPAQHFNYVKLAAPVPKFKAAVLQMPSSPTDSNPQ

>C_doughertyi_SPDL1_paralog1

MPDDEEKLQLQADVERYKKAIRQKDEMIEEMENELNSHGKLPVSDSKAEEREKALNGQIRDLRFEIDGKDATINEQTDIINNLRDEMEKLEKKNRELVNRSDCTEIDESNSFVESEMIRISEECEKFKEMASSLFEQNMELKKEAVELREEYDSAIELMKNLESHIKTKEEEIARLEGEVFDLKSSSQGKHASAGNSIFAEAMEAERKLEEDLKTLFREKQALMSMVKRLTLEKDEAEERARNFMNRGLIFRNSINHIDVEEMRRLRTRVQQLETERTNFWERLFIKMKTVPEREIGSIIAGYFESFKCSIANVTGGFEELMKKNEQYVTTIRGLQQEIENQRVKIEQMQFDIECSERKMRSALSLESETPSLRAPLKPMDNSRPSFFVKPKKADPVPVPNLEVSLSNILLTPQKPSAEVTTQSTTKKDDASEWTERRLKAKAEKKLATPALRYNYVQLKGPAPKFKPAVLPMPRTSPNSKEN

>C_doughertyi_SPDL1_paralog2

MRDDEEKLQLQADVERYKKAIRQKDEMIEEMENELSSYGKLPMSDSKAEEREKALNGQIRDLRFEIDGKDATIKEQTDLINDLRKESNSLEESNSLESEMIRISEECEKFKEMASSLFEQNMELNKEAVELREDYDSAIELMRNLESHIETKEEEIARLEGEVFDLKSSSQRKHASAGNSIFAEAMEAERKLEEDLKTIFREKQALLSMVKRLTLEKDEAEERARNFMNRGLIFRNSINHIDVEEMRRLRTRVQQLETERTNFWERLFIKMKTVPEREIGSIIAGYFESFKCSIANVTGGFEELMKKNEQYVTTIRGLQQEIENQRVKIEQMQFDIECSERKMRSALSSESETPSLRAPLKPMDNSRPSFFVKPKKADPVPTLTQKVAKEDDASEWTERRLKAKAEKKLATPALRYNYVQLKGPAPKLQPAVLPMPRASPNSKEN

>C_sp54_SPDL1

MPDDEEKLQLLADVERMKKTIRRKDDLIEEMENELNNFGKTSRSDGRAEAREKELNGQIRDLQTEMDGKDTTIREQNDLIITLRGDLVKLENSNRELINRSECVDSDESNSFVETEMLRISEECEKFKELASTLYEQNLDLKKEALELREEHDTVVEHVRSLQSHIKTQEEEIARLEGDIFDLKNSNQGKHAASGNSIFAEAMEAERKLEEDLKVLFREKQALMGMVKRLTMEKDEAEERARTFMNRGLIVRNAMNHIDVEEMRRLRARVYELETERTHFWERLFIKMRTIPKRDIGQLFAGYLDSFKCSIASVKGGFEELLKKNEQYVLAIRGMQQEIENQRVKIEQLEFDIECSERKMRSSANTDLDHPQLRAPLKPIDNSRPSFFIKPKKAYHVPQLEVSMSKMLMTPQKPSGETTAHSTVKKESTSEWAERRIKAKNEKKSSPAPKFKAAILQMPTALSESKEN

>C_inopinata_SPDL1

MVEDEEKLQLRADLERCRKMVRQKDEIIEEMEHDLNRRGKEYIFDKKLEAREKELNAQIRQLQVEVDGKDATIQEQRDLVEDLRRDICTLEKSRELLNRTSSPKSDESTSFEKSEKIRISEECEAFKEVASTLYEQNAELKKNAVELKEELESAMEHIRSLQSHIKTLEEESSRLEGELFDLKNSNNGKIASSGNSIFAEAMEAERKLEEDLKVLFREKQSLACRVKRLTMEKEEAEERARSFISRGLIIRNAINHIDVEEMRRLRTRAHELETERIHLWEKLFTKMKTTSKREMGAIIAGYFESFKCSIESVKGGFDDLMKKNENYVNMIRVQQKEILDLKEKIEQQKFDIECYERKMHSVANANLENAQLLEPLKAIDNARPSFFVKPKKVESINQLETSMSNTLITPQKEVLSEWAEKRMKAKAEKNSATPGPRYNYITLTGPTPKFKSAILQMPSTPSEPKND

>C_elegans_SPDL1

MPDDEEKLQLLADVERLKKILRQKDEMLEEMEDDLKNQGKPCSSKLSLEERAQELSEQLRDLHVEMDGKNATILDRDALIDSLRSEIDKLEKINKEFANGSVIPEHDDSNSFGESEMLRISEDCQKYKETATALYERNAELEKEAVNLKDEIESMMDHIRDLKNHMETRDEEIARLEGELFDERNSHEGKLAARGNSMFSEVIDAERKVEEDLKVLHGENRALKGMVKRLRMEVEEVEERLRSSTKRFNVTRMTTSDIDVKEMRRLRDRVCHLETERVHLWERMFIKMRSIPKREVGALITGYFKSFELSIASVKGGFDGLMKDNEKYVTIIRGLQQEVENLKADIVQLQFDNKCAHRKAAPVVNKDFEHPLLAAPLKTLNNGRPSFFIKPKNVEPMPQLGHSLSSIAVTPQKPAAKFTTRSSIKDDTSEWAERRMKAQAEKKLATPTPRYNYIKLSEPVPKFKPAVLQMPSTSETKEN

>C_oiwi_SPDL1

MPDDEEKLQLRADLERTKKMLHQKDDMIEELEQELNNKAQVSDWKSEAREKQLNGQIRDLQMEMDGKDLTIQEQNGLIGELRVAVDRLEKRNRELANRSILPESDDSCSFDESGLIKASEECEKLRQNVVTLQEHNMELQKELIEKKEEHEGAVEHIRNLESHMKTQYEEIARLEGEIFDMKNSAQGKHASCGNSIFAEAMEAERQLEEDLKKLFREKQALTSMVKRLTIEKEEAEERARAYMNRGLIVRNAINHIDVEEMRRLRARVHELETERTHLWERLFIKMKSVPKREIGAIIVGYFESFKCSIASITGGQSDLLKKNEQYVTTIRGLQQEVENQRVKIEQLQFDVECAERKMRSAVTSNSEGENPQLRAPLKSVDNSRPSFFVKPKKKELAPPPSLELSMANMEMTPQKPSLITNRSSTAKKEDTSEWAARQLKARADKKATPAPKYNYVSLKQPVPKFKTAVLQMPSSPTESKEN

>C_kamaaina_SPDL1

MPEDEEKLQLRADVDRLKKMVHQKDEMIEEMEHELNNKAPVSDWKAEAREKQLNEQIRDLQVEIDGKDVTIQEQQGLIGELRVEIDRLEKNRELANRSILPESDDSCSFDESGLIKASEECEKLRQNVVTLQEHNMELQKELIEKKEEHEGAVEHVRNLESHMKTQYEEIARLEGEVFDMKNSANGKHASCGNSIFAEMEAERQLEEDLKKLFREKQALTSMVKRLTLEKEEAEERARAYMNRGLIVRNAINHIDVEEMRRLRARVHELETERTHLWERLFIKMKSVPKREIGAIIVGYFESFKCSIASITGGQSELLKRNEQYVTTIRGLQQEVENQRVKIEQLQFDVECAERKMRSAVTSNSEGEIPQLRAPLKSIDNSRPSFFVKPKKKEPAPPPSLELSMANMEMTPQVTNRSSTAKKEETSEWAARHLKARADKKATPAPRYNFVELKQPVPKFKAAVLQMPSSPTERKEN

>C_waitukubuli_SPDL1

MPDDEEKLQLQADVERYKRMIRQKDEMIEDLEHELSNHGKTPVSDGKTEARERELNGQIRDLQLEIDGKDATIVQRDGVIDELREEIEKLEKSSRELANRSEYQDSDESNSFLEHEMVRISEELEKYKEATGNLFEQNSELKKNVLQLQEENENAMEHMKSLESHLKTREEQIASLESELFDLKNSSRGKHASSGNSIFAEVMEAERKLEGDLRNVFTQNQALSAAVRRLTLEKEDAEERARSAMNRGLVVRTAINHIDIEEMRRLRERVHELELERTHCWERLFIKMKTVPKRELGALVAGYFESFKCSIVSVKSGQAELLKKNEQYVTTIRGLQQENENQHVKIEQLQFDIESLTRKLKSSVNDSEPAQLRAPLKPIDNSRPSFFVRPKPVEVPVPTASMSNMHVTPQKPANLPNTCSTAESKNENTSYWADRQKVKAEKKAATPAARYNFVTLTEPKPKFKPATLMMPST

>C_panamensis_SPDL1

MPVDEEKLQLRADVERYKRMIRQKDEMIEEFEHELNSHGKAPVSDGRAEAREKELNGQIRELQAEIDEKDATIVGKDGVIGELREEISKLEKQSRELANRSECQDSDESSSFLEQEITRISEELHKYKEATETLFEQNSELKKEAVQLKEEHESAMEHVRNLESHLKTREEEVARLESELFDLKNSNQGKHASKGNSIFHEVMEAERKLEEDLRTVFAENQKLKALARRLNVEKEDAEERARSAMNRGFVVRTAINDVDVYEMNRLRERNRQLELERTSLWEKLFVKMRTIPKKELGFLVAGYFESFKLSIISVKSGHEELLKKNEQYVTMIRGLQQQNENDSVKIEQLKFDIEGLNRKLINATRDNDPARVPLKPVDNSRPCFAKKKPVETAAPVVSMSTMQITPQENTSYWEQRQKIKAARAAKTPVAPVSRYNFVQLIEPKPLFKPTTLVVPSTPKEE

>C_nouraguensis_SPDL1

MPDDEEKLQLRADVERYKKMIRQKDEMIEEMEHELNSRGKTPTSDGRAGAREKELGVQIRELQLEIDGKDATIEMKDNVIEELRGEIDKLEKSRDLANRSEYPESDESSSFLEHEMVRISDELEKYKEATANLFEQNTEFKKDNLQLKDEHENAMEHVKSLESHLKTREEEVVRLESELFDLKNSSNGKHASTGNSIFAEAMEAERKLEEDLRKLFSEKQALASMVRRLDMEKNEAEERARTAMNRIVVRTAINNVDIEEMRRLRERVRELELERTHLWERLFIRMKSIPKRELGSLITGYFESFKCSIVSMKSGQDEVMKKNEKYVTTIRGLQQENENLRVKVEQLEFDIDSLNRKLKSAATASSKFFSRILTNLYVHPDFESEPAQLRAPLKPFDNSRPSFFVKPKAAEKPTAMPNVSNMYVTPQSTNEALKENTSDWAERRQKTKAEKKQAATPATRYNFIQLTAPQPKFKPATLMMPSSDKED

>C_becei_SPDL1

MPEDEEKLQLRADVERYKKMIRQKDEMIEEMEHELSSRGKTPTSDGRAEAREKELGGQIRELQLEIDGRDATIETKDGVIGELRDEIERLEKQSRDLANRSEYPESDESSSFLEHEMVRISEELEKFKEATANLFEQNAELKKVELQLKEDHENAMEHVKSLESHLKTREEEIVRLESELFDLKNSSNGKHASTGNSIFAEVMEAERKLEGDLRTLFAEKQALTAMVRRLDMEKNEAEERARAAMNRIVVRTAINNVDVEEMRRLRERVRELELERTHLWERLFIKMKSVPKRELGSLIVGYFESFKCSIVSMKSGQDELMKKNEKYVSTIRGLQQENENLRGKIEQLEFDIDSLNRKLKSSANATSECSSCVLNIFFVHLDIETEPAQLRVPLKPIDNSRPSFFVRPKDVETPAAMPNMSSMYITPQQKPAKMPSASNTVEAPNEDTSEWAERRQKAKAEKKQAATPATRYNFIQLTAPKPKFKPATLMMPSTDKEN

>C_yunquensis_SPDL1

MPNDEETLQLRADVERYKKMIRQKDEMIEEMEHELNNHGKATVSDGKADVREKELGAQIRDLQLEIDGKDATIVMKDGVIEELRDEIKKLEKQSQELANRSDYPESDESGSFLENEMARISDELEKYKEATANLFEQNSVLKKEGLQLKEEYENAMDHVKSLESHLETRNGEIARLESELFDLKSSNQGKHASSGNSIFAEAMEAERKLEGDLRKVFHEKLALTNMVRRLNMEKDDAEERARNAMSRGRVVRTAINEIDIAEMRRLQERVRELELERTQLWEKLFIKMKSMPKRELPALITGYFESFRCSIVSMKSGQDAVMKNNEKYVTTIRGLQQQCEMDRAKIEQLEFDVESLTRKLKSAAKTSDDQSGAALSQDSLKPTVNSRPSLFAKPKPVESSASVPAQLRAPLEPINNSRPSFYVRPKPMESSAAKPVLGNIHSTPQKSVDSASTCSTVERPNESTSEWAERREKAKAERKQAATPAARYNFIKLSAPEPKFKPAKLMMPSSDQEN

>C_macrosperma_SPDL1

MPDDEEKLQLRADVDRYKRMIRQKDEMIEEMEHELSNHGKPATSDGRAEAREKELSGRIRDLQIEMDGKDTTIAIKDGLIDELREEIMKLEKNSRDLANRSEYQDSDESSSFLEHEMVRISEELEKYKEATANLFEQNSELKKEGLQLKEEHEAAVEHVKSLESHLRTREEEIVRLESELFDLKNSSQGKHASSGNSIFAEAMEAERKLEEDLRKVFREKQALTVMVRRLTLEKDDAEERARSAMSRGLVVRTAINHIDVEEMRRLRERVRELELERTHLWERLFIRMKTVPKRELGALISGYFESFKCSIVSIQSGQDEVMKKNEQYVTTIRGLQQENENQRVKIEQLQFDIECLNRKLKSAVIASNADSEPAQLRAPLKLIDNSRPSFFLKPKPVESTAAPIFEMSSMHVTPQKPANIPSACSTVESKKENTSEWAERREKVKAEKKQAATPATRYNFITLAAPQPKFKPPTLMMPSTPKEE

>C_sulstoni_SPDL1

MEDEEKLQLDVERLKRQIRQKDDMIEEMEQELTSGRHAHGPVSDGKAEAREKELNAQIRDLQFEMDGKTTTIGEKEELIEKLRDEIDRLEKNRELANRSVVSESDESSSFHEVEMVKLGEQLEECRHHIASLEEQNLQLRKEALQLKEEHEMVMDHVKNLESHIKTGEEEIARLEGEVFDLRHSNQGKHATTGNSIFAEAMEAEQKLEQDLKTCYAENQALKKKIRRVILEKEDAEELARSAMGRGVVVRTAINHIDVLEMNRLRAKVHELEKERTLFWLHLFEKMKAAKITRRELGGLIAGYFESFKCSIASVTGGQEEILKKNEQFVTTIRGLQQENENLRVKTEQLQFDIGCLNRKLRSAINLRQPLKPVDNSRPSFFHKRAPVVDPAVQSETSMSSSQVPEQLRQPLKPFDNSRPSFFQRRAPVVESAVPLDSSMSSMHMTPQKPAFPQTTCSSQKENTSEWAERRQKAKAERKQAATPAAYNYVTLLAPQPKFKLPTLKMPSSPTTPRVHEE

>C_afra_SPDL1

MEDEEKLQLRADVERLKRQIRQKDDMIEEMEHELTSGRHVPASDGRAEAREKELNTQIRDLQIEVDGKNEAIEKREELIEELRDEIDKLEKRNRDLANRSVCPEAEDSSSFHEMEILKMGEEVEECRRQMASLEEQNLELRKQALQLKEEHEMAVDHVRSLESHIKTGEEEIARLEGEVFDLKNSSQGKHASAGNSIFAEAMEAEQQLEQDLKKCFAENQALKKLLRRVTDEKEDAEERARSAMSRGLIVRTGVNHIDVMEMNRLRAKVQELETERVHFWHHLFQKMKAAKLTRRELGSLIVGYFESFKCSIASVKGGQEEILKKNEQYVTTIRGLQQEIENLNAKVETLQYDIECANRKLRRANNIGSKSGHSRSVLNSSSDSEADVPPQLRQPLKPVDNSRPSFFRRPAPAAEPSAPLETSLSFMNVTPQKPAFLQNASSTQKENTSEWTERRQKAKAEKKAATPATRYNYVTLSAPKPKFKQGTLQMPSTPASLVQEEAEHE

>C_sp49_SPDL1_paralog1

MVDDEEKLQLRADVERFKRMIRQKDEMIEEMEHELSRPKTPGDSLRTAARERDLATKIRDLEIEIDAKNVTITQKDGQIDELRDEVDKLEKSNRDLAYRPESPSQDSSSSFLENELARLSEELDTFRESTSSLMHQNLEYEKELLQLRSDLESATEHVKSLETHLKTREEELARLEGEMFELKASGHGKHASAGNSIFAEAMEHEKKLEEDLKFLFAQNQCLLKKVRRLQIEKEDAEQRAQSAMRRKYTVGTAINHQGFAEMDRLRQKVRELEVERTHLWERLFVKLRGVNKREFGPLIVGYHESFKLSITSVTGNQEKVLKENEELAKKVQGLETINADYQEQVEQLKYDLETAQRKAESAVDLQDSQYPLPKLTRPILTRKPKPEDDEEFTSGKSHPLLNAPLKPMSSAVRNSFFVKPNRDPIDKGIMNMSIENSLRMMPGTPRAQDQENVSDFTLHHQRAKAERKAAATPARPNYNFVQLLQPTRTTSKFKPAILQMPKPTDE

>C_sp49_SPDL1_paralog2

MVHDEEKLQLRADVERYKRVIRQKDEMLEEMEYELSRPKTPADSMLTAARERDLVTKIRDLEIEIDAKNVTITHKDGQIDELRDEVDKLEKSNRDLVYHSESPSQDSSSSFLENELARLSEELDTFRETTSSLINQNLEYEKELLQLRSDLESATEHVKNLENHLQTREEELARLEGEIVELKALGHGKFATAGNSIFAEAMEHEKKLEEDLKFLFAQNQRLLKKVRFLQIEKEDAEQRVQSAMRRKYTVTTAISYQGFAEMDRLRQKVRELEVERTHLWERLFVKMRCVSKREFGPVVVAHYESFKLSIASVTGNQENVLKENEELSKQVHRLENINAEYEEQVKQLKYDLETAERKAKSAVEEPKDLQDSQVPPPICTRPIFFSRKPKPEVAQNPEDEFTSGKSHPLLNAPLKPMSSAVRNSFFVKPSRDPIDRGIKNMSVENSLQMRASTLRVQDQEKLSDFTLHHQRAKAERKAAAPPAELKLNYIQLIQPTTTTSKLKPAILWMSKSTDK

>C_sp49_SPDL1_paralog3

MVDDKEKLQLRADVERFKRMIRQKDEMIEEMEHELSRPKTPGDSLRTAARERNLATKIRDLEIEIDAKNVTITQKDGQIDELRDEVDKLEKSNRDLAYRSESPSQDSSSSFLENELARLSEELDTFRETTSSLMHQNLEYEKELLQLRSDLESATEHVKSLETHLKTREEELARLEGEMFELKASGHGKHASAGNSIFAEAMEHEKKLEEDLKFLFAQNQCLLKKVRRLQIEKEDAEQRLKSATRRKYTVSTAINHQGFAEMDRLRQKVRELEVERTHLWERFFVKLKRVSKREFVPLIVGYHESFKLSIISVTGNQEKVLRENEELAKKVQGLETINADYEVQVAQLKYDLETAQRKAESAVDLQDSQYPLPKLTRPILTRKPKPEDDEEFTSGKSHPLLNAPLKPMSSAVRNSFFVKPNKDSIERRITNMSIQNSIRKAEMEAEAPPTTIKTNFVQLLQPTRTTSKFKPAILQMPKSTDE

>C_sp25_SPDL1

MVEDEEKLQLRADVERFRRMIRQKDEMIEEMEHELSRPKTPGDKLRLEARERDLASKIRDLELEIDAKNVTIHKKEGQIEELRDEVDKLEKSNRDLAYRPESPQQDTSSSFMENELARLSEELDKCRETNSALMQQNLELEKENIQLKSDHESAMEHVKSLEVHLKTREEELGRLESENFQMKAAGHGSHASAGNSIFAEAMEAERKLEEDLKSLFAQNQCLIKKVRRLQIDKDDAEQRVQSAMQRKYVVSSAINTQGYEEMVRLRQKVRELEVERTRLWEKLFVKLRRVNRKELGPLVLGYHESFKLSIASVTGNQENVLKENEELATKIQGLEKMNAEKEEKIEQLKYELATAQREKKVAAALESATEDFASEKDQNIPLKQSVLLNYSRKPKQPEIPENPDEEQFTSGRDHPLLNAPLKPMNSGVRPSFFVKPKKEPVERTMMNMSIDPQSSFQTIPQTPTSGNDRENISDFTFRYQKAKAERKAAATPAKLSFNYVQLIQPTTTSTSKFKPATLQMPKIDD

>C_imperialis_SPDL1

MVDDEEKLQLRADVERFKRMIRQKDEMIEEMEHELSRPKTPGDRVRMEAREREFAGKIRDLQIELDARNVTIQKKDAVIDELRDEVDKLEKNNRELAYRPDSPTHDSTSSFLENEMSRMSEELDKFRETTSSLMQQNMEYEKEVLQLRGDLESSMEHVRSLESHLKTREEELERLEAELFELKASGHGKAASAGNSIFAEAMEAERKLEEDLKSLFAQNQCLLKKVRRLHIEKEDAEERAQSAMSRKYTVRSAINTQGHAEMDRLRQKVRELETERTHLWERLFVKLRNVRRSELGALVVGYHESFKLSIASVKGNHEQVLRENEELAVKIQGLERENADYEEKIEQLKYDLATAQRKEKAAIALEADEDFSSGKELMDAPLKGPVGRPTFFVKPKQVEFSEIPEDEEFTSGASHPLLSAPLKVQNSAARPSFFVKPKSEHIERGMMNMSMTPKNASRLPPTTPSSLDQENMSDFALRHQRAKAERKAAATPAKPQYNFVQLVQPTAVSRFKPATLLPPKPAE

>C_japonica_SPDL1_IncompleteSequenceInfoInExon4

MVEDEEKLQLDIERLRRQIRQKDEMIEEMESELSREKTFIANDKAEIRERELNGQIRGLQIEIDGKDGTISQKDGVIDELRVEIEKLNRELANRSDYPESDESSSFLETEMMRISDELQKFKETTSSLLEKNLEYEKECLQLKSDYESAIDHVNSLESHLKTREEEISRLESEVFDLKCSAHGKIANAGNSIFAEAMDAEQKLEEDLKRVYHEKQYLMERLKRIQLEKEEAEERAQANLRRNCTVRTAVSHVDIEEMRRLRARVHELEVERTCLWERLFIKMRTVPKREMGGLFAGYLESFKPASLKTLFTDTKENTTSEYSERQQRIKAERKQAATPASRYNFVQLAVPSTVPKFKPPTLNMPST

**Supplementary Data S8: All NDC-80 sequences used in this study**

>C_tribulationis_NDC80_paralog1

MFGGRRTGGPGFNAGRLSTAVTPTKRFTDFGIGSTRKAGRLSMSQGPRPSFFTKGSTVPPRDVKSLQAANVQKIYNFLVEQDGSDAPSESSIRSPGGGKDFKMIFESMYQHLSKDYEFPLNARIEDEVMPIFKGLGYPFSLKTSFFQPMGGPHGWPHLLDALAWLVDVIRMNLAVSVDTQNILFGDFLDHQKVQEKALSYMWYSTLFRDYTNDRKAAEDKDGVFWKEAKANLRQYFENSNEYEEMITNLRNVLQQLRFDCDEIEAEKGQEQTYVEDIARMKDDIRKAVEYLESTQRVKENKEAEVTSIKQELDMKNAEMEKCLGMVYELKERIEHQKQVHGCSGKEVRQMNLENSKDKETVSELQAELDEISKETWRLKNDDSFKEQKAKFVQLVENITKLLAGLDVKLNLDPLSVPSDEKELKAGWETLNSVWVPEISRQMHQRKLELETDKARFVDKFAAAEERIQIENEMLCEAKKKEGRDERIQRIEREEWKVGRQQLEKRYDELENEREVLTKKMQMDGSLEKEIKEEKDKMAKMESEAEAKSQEVQAAIREKLEEMLVEIAEIGQEKYMFHVEATEVSRLISGMCSLDS

>C_tribulationis_NDC80_paralog2_pseudogene

RNVFLLPIFSPAPRLFDQITFPNDINIKRTFYHFMGNTLVTKTIDEAKRIDQRYGGRYLITTFEGAIIDQSGNLTGGGTPLTGRMNVTGASSSRFNNDIERKNHIFKTKTRMQQTEREINSIEAMLNAEMTKMQTDKASAAALENEYNQLNETLKNLRSHVEKLHDQSMACDHRLAQIAPIEEITASIAEITQELEVLRERQKRQVAQNQEAQHIVSSLSSKI

>C_sp41_NDC80

MFGGRRTGGPGFNAGRLSTAVTPTKRFTDYGIGSTRKSEAAGRLSMSQGHRPSLFQKGSAVPPRDVKSIQAANVQKIYNFLVEHDGSEAPAESIIRTPRGKKDFEAIFESMYQHLSKDYEFPTQGRIEDEVTQIFKGLGYPYPLKNSFFQPMGASHGWPHLLDALAWLVDVIRMNLAVSVDTQNILFGDFLEQQKVQEKALSYMWYSTLFRDYTNDRKAAEDKDGNFWKETKANLRQYFENSNEYEEMVTNLRNVLQQLRFDCDEIEAEKGQEQTYVEDIARMKDDIRKAMEYLDSTQRVKENKESEVTSVKQELEMKTAEMEKAIGMVNELKERIEQQKLVHGCSGKEVRQMNLENSKDKETVSELQAELDEISKETWRLKNDDSFKEQKAKFVQLVENITKLLAGLDVQLKLEPLSVPIDEKQLKAGWETLNSVWVPEISRQMHQRKLELETEKARFVDKFAAAEERIQIENEMLCEAKKKEGRDERIQRIEREEWKVARQQLEKRYDELENEREVLTKKMQMDGSLEKEIKEEKDRMAKLESEAEAKSQDVQMAIREKMEQMVVEIAEIGQEKTMFHGESTDVARVIGGMCSLDC

>C_zanzibari_NDC80

MFGDRRKTGGPSFNGGRLSTAVTPTKRFTDARLSMSQGHRPSLFQKGSAVPPRDVKTLKAANVSKIFNFLVESDGSEAPSESTIRSPPGKNDFIAIFESMYQHLSKDYEFPASARMEEEVSSIFKGLGYPFPLKNSYFQPMGGAHGWPHLLDALAWLVDVVKMNQAVSRDTQNILFGDFMDQGKVQEKALSYMWYSKVFRDYTNDRKAAEDKDGEFWTNSGAELRRYFENSNDNEEMMTNLQNVLQQLHFDCDEIEAEKGQEQTYVEDIARMKDDIRKAAEYLESTQRVKELKQAEFTAVKQDLDSRKAELEKAVGMVIELKERIEQQKRIHGCSGKEVRQMNLENSKDKEMLSELQAELDEISKETWRLKNDDSFKDQTKKFVQVLVNIRKMLANLNIQLNLDLLQVPKDEQELKVCWETLNGVWVPEISRQMHQRKLELETEKARFVDKFAAAEERIQIENEKLCEAKKKEGRDERVQRIEREEWKVARQQLEKRYDELENEKEVLTKKMHLDGSLEKEIKEEKDKMTKMEQVAEAKRQDLEMAIREKLEKMVVEISEIGQEKIMFHAESTDVARVINEKCLLDC

>C_sinica_NDC80

MFGNDRRKTGGNFNTGRLSTAVTPTKRFTDFGMSSVRARHSISQSRPSLFTKGSAVAPRDVKSIQAANVQKIRNFLVETDGPEAPDEATIRSPRGKNDFIAIFESMYQHLSKDYEFPVQARVEEEVTQIFKGLGYPYPLKNSFFQPMGASHGWPHLLDALAWLIDVIRMNQRVSSDTQGIMFGDTFDQQKVQETALGYMWYSSLFRDYTNDRKAAEIKDGEFWVETKQKLRNFFENTNEFEDMAVNLKNVLQQLCFDCDEIEAERGQEQTYVEDIARMKDDIRKAVEYLESTQRLKELKETDVTSVKQELEFKKSEIEKVHGMVNELKERIEQQKRIHGCSGKEVRQMNQENGKDKETVSELQSELDELSKETWRLKNDDSFKEQKAKFVHIVENIMKILNGLKVEWNLAPLPVPSDERQLKTCWETLNGVWVPEISRQMHQKKLELETERARFAHKFAAAERIQIESEMLREAKKKEGRDERIQRIEREEWKVTRQQQEKRYDELENEKEMLKKNMHLDGSLEKEIKEETEKMIKIETESEAKSQDLQATLRAKMEQILVEIA

>C_nigoni_NDC80

MFGGERRKTGGMNLNGRQSLAITPTKRFTDHGLSSVRKTDARPSMPLHGQSRPSLFQKGSTIPPRDVRSLQAANVQKIYNFFVETDGAEAPSERSIRAPGSRREFVVLFESMYQHLSKDYEYPEPARLEDEVTQIFKGLGYPYPLKNSYYQPMGASHGWPHLLDALSWLVDVIKMNTTVAANTQGILFGDILEQSKVQEKVLNYSWFASIYKDYTNDRKGTEDKDSQFWKDAKNKLRQHFENSNEYDDFASNAKNVLQQLIFDCDEIESERGQEQTYVEDIARMRDDIRKAAEYLQSVELVKEHKDAEMVKVKGELDSKVAEKEKLLRMVNELKDRIEQQKIIHGCSGKEVRQMNLENSKDKEMVAELQAELDEVSKEMWRMKNDDSFKEQKAKFLQIVENITKLLSGLNVQLNLDPMPVPADEKQLKTCWETLNSVWVTEISRQMHQRKLDLDTEKSRSLDRFAAAQERIQIENEMLCEAKKKEGRDERTRRAERDEWKAARQQQEKRYDELENEKEVLMKKLHLDGSLEQEIKEEKDKMAKIEKEAEEKTQYLRSAIRQKVEAVEMELAEIGQEKTMFHAECVAVEKLVQGTCGATH

>C_briggsae_NDC80

MFGGERRKTGGMNLNGRQSLAITPTKRFTDHGLSSVRKRTDARPSMSLHGQSRPSLFQKGSTIPPRDVRSLQAANVQKIYNFFVETDGAEAPSERSIRAPGSRREFIVLFESMYQHLSKDYEYPEPARLEDEVTQIFKGLGYPYPLKNSYYQPMGASHGWPHLLDALSWLVDVIKMNTTVAANTQGILFGDFLEQSKVQEKVLNYSWFASIYKDYTNDRKGTEDKDSQFWKDAKNKLRQHFENSNEYEDVASNAKNVLQQLLFDCDEIESERGQEQTYVEDIARMRDDIRKAAEYLDSVERVKEHKDAEMVKVKGELESKVLEKEKLLRMVNELKDRIEQQKMIHGCSGKEVRQMNLENSKDKEMVAELQAELDEVSKEMWRMKNDDSFKEQKAKFLQIIENITKLLSGLNVQLNLDPMPVPADEKQLKVCWETLNTVWVTEISRQMHQRKLDLDTEKSRSADKFAAAQERIQIENEMLCEAKKKEGRDERTRRAERDEWKAARQQQEKRYDELENEKEVLMKKLHLDGSLEQEIKEEKDKMAKIEKEAEEKTQYLRSAIRQKVEAVEMEIAEIGQEKTMFHAECVAVEKLV

>C_remanei_NDC80

MFGTERRKTGGVNLNGRSSIAITPTKRFTDFGATSVRRTDGRPSLSVGQLGQQSSRPSIFKTSVHAPRDVKSFQAANAHKIFNFLVESDGADAPAESVIRTPRGKNDFIVIFESMYQHLSKDYEFPPLNQARIEEEVSSIFKGLGYPFHLKNSYFQPMGASHGWPHLLDALAWLVDFIKINKSVSADTQHIIFGDFLEPAKVQEKALSYAWFSTTFRDYTNDRKSAESSDSEFWVETKNKLRKYFEDSNEYEDMTANAQSALQQLRFDCDEIESERGQEQTYVEDIARLKDDIRKAMDYFESVQHLKERKENETAVIKEELEAKVAENEKIQMAVNELKERIEQQKRVHGLNGKEVRKMNLENSKDKDTVSDLQSEQEMLSKQLWRLRDENPFKDQKMKVIQIAENVTKILSGLNMQFELESLRPPENEKELRACWEILIGSWLPEINRQLHQRKLDVETEKSRFHQKFAAAERIQIENEVLDEVKKKEAREERIHRDERDEWKEARQKQEKRYDELENEKEMLKRKMQMDGSLEKEIKAEKEKKEKIKKDLEEMHQGLEAVFRKKLEKIASETAEIGNEKTMFRIEAIEVEKLIDRKCSNQKV

>C_latens_NDC80

MFGTERRKTGGVNLNGRSSIAITPTKRFTDFGTTSVRKRTDGRPSLSVGQLGQQSSRPSIFKTSVHAPRDVKSFQAANAHKIFNFLVESDGADAPAESVIRTPRGKNDFIVIFESMYQHLSKDYEFPPLNQARIEEEVQSIFKGLGYPFQLKNSYFQPMGASHGWPHLLDALAWLVDFIKINKSVSADTQHIIFGDFLEPAKVQEKALSYAWFSTTFRDYTNDRKSAEYSDSEFWIETKNKLRKYFEDSNEYEDMTANAQSALQQLRFDCDEIESERGQEQTYVEDIARLKDDIRKAMDYFESVQHLKERKENEMAVIKDELEAKVAENEKIQMAVNELKERIEQQKRVHGLSGKEVRKMNLENSKDKDTVSDLQSEQEMLSKQLWRLRDENPFKDQKMKVIQIAENVTKILSGLNMQFELETLRPPENEKELRACWEILIGSWLPEINRQLHQRKLDVETEKSRFHQKFAAAEERIQIETEVLDEIKKKEAREERIRRDERDEWKEARQKQEKRYDELENEKEMLKRKMHMDGSLEKEIKAEKEKKEKIKKDLEEIHQGLEAVFQKKLEQIASETAEIGNEKTIFRIEAIEVEKLIDRKCSNQKV

>C_sp51_NDC80

MFGERRKTGGMSLGGRQSIAITPTKRFTDLGIASVRKTDARSSMSVNQPRMSLFQNKGSSLPPREVKSIQQANVQKIHQFLVESDGDDAPAEVVVRTPRGKKDFTMVFEIIYQHLSKDYEYPHPVRLEDEIPQIFNGLGYPYPLKNSYFQPLGAGHGWPHLLDALGWLVDVVKMSKAVVPNTQSIIFGDFLEQSKVQEKTLNYAWTTNTYRDYTSDRKVIENREHPFWEQAKSQLRNYFESSNEYEDMESNAKNALEQLMFDCEEIESERGQEQHLQEEIAKMKDDIRKAEDYLLQTENLKSHKEKDLGKIRETLVEKQSELEKIQQLVNELKERIEKQKINHGLSGKEVRKLNLENFKDKETVSEVQGELEKLSKEIWTLNNDDRFKEQKSKFVQLTDNIAKLTTGLNIQLSLEPLKAPTDERQLKTHWETLSNIWLPEINRQLHQRKLELETELSRFADSFAAAEERIQIESETLCEAKKREAREERIRRNERDEWKTSRQELEKRYDELENEKEVLEKQMQIGGSLDKEIDEERARMLKTEQDLRAKEADLMAAIRQKLDGLFAGIVELDNEKMMILKEFNDLEMVMNVKCANY

>C_sp44_NDC80

MFGNDRRKTGGMNLGGRASIAITPTKRFTDFGTGSVRKTDARLSVNQPRPSLFQKGSSLPPREVKSIQQANAQKIYQFLVENDGDDAPSEASIRTPSGKKEFGAIFEAIYQHLSKDYEFNHQARAEDEVTPIFKGLGYPYPLKNSYFQPMGSPHGWPHMLDALGWLVDFVKINKEVNKDTQNIIFGDFLEQNKVQEKTLNYAWNTNTYREYTLDRKVLDDQSHPFWEQTRIQLREYFESSSDYGDMATNIKNALDQLMFDCEEIESERGQEQQLQEEIAKMKDDIRKAEDYLLQTENIKSHKEKDLEKTLQCLVERQSELEKVQIAVNELKERIEKQKINHGLSGKEVRKLNMENQKDKETVSEIQGELDKLSKKTWTLKNDDCFKEQKSKFVQLTDQISKLMSGLSVQLNLQPLKAPSDEQELKTHWETLSNIWLPEINRQLHQRKLELDTELSRFGDKFSAAEERIQIESEMLCEAKKREAREERIRRNERDEWKTSRQEMEKRYDELENEKEVLKKQIQIGGNLDKEIEEEKTKMVKIEEALRTKEADLIAKLRQKLEEIVVGLAEIDQEKMRIHKEFNDLEIVMNKKCAHY

>C_sp48_NDC80

MFGNDRRKTGGMNLGGGGRASIAITPTKRFTDFGASSVRKVDPRLSVNQPRPSLFQKGSSLAPREVKSLQAANIQKIYQFLVESDGDDAPSEQTIRAPRGKTDFVMIFETIYQHLSKDYEFPQQTRLEDEVVLIFKGLGYPFPLKSSHFQPMGTGHGWPHLLDALGWLVDFVKISKGVSTARQNILFGDFLEQDKVQEKALSYAWNTNTYRDFTSDRKVLDNRDHPFWEQTKSQLRTFFESSNEYEDMANTGKNALEQLMFDCEEIESERGQEQHLQEEITKMKDDIRKAEDYLLQTENLKVRKEKDLMTMRETLAEKKAMLEKVLQAVNELKERIERQKIDHGLSGKEVRQLNLENMKDKETVSEVQGELDALSKLSWTFKNDDCFREQKAKFVKLTVDIAKLMAGLDIQLNLEPLKAPRDEPELKTHWETLSTVWLPEINRQMHQRKLELETELSRFADKFAAAEERIQIETEMLCEAKKREAREERVRRNERDEWKIARQELEKRYDALENEKEVLKRQMQIGGNLDKEIDEEREKMRKIEEALRAKEAELETEIRLKLDEMFTQIAKIDAEKMMIFKQFTDLEAIMNANCARY

>C_brenneri_NDC80

MFGNDRRKTGGMNLGGGRASIAITPTKRFTDFGGASSIRKVDPRLSVNQPRPSLFQKGSTLPPREVKSLQAANVQKIYQFMVEADGDDAPSEQTIRAPHGKTDFVMIFETIYQHLSKDYEFPQQTRLEDEVIQIFKGLGYPFPVKPSYFQPMGVGHGWPHLLDALGWLVDFVKINKAVNKDTQNIIFGDFLEQQKVQEKTLNYAWNTNTYREYTSDRKVLDNRDHPFWEQTRSQLRAYFESSNEYEDMASNVKNALEQLMFDCEEIESERGQEQHLQEEITKMKDDIRKAEDYLLQTENLKAHKEKDLVKVRENLAEKQAVLEKVLQAVGELKERIERQKIDHGLSGKEVRQLNLENMKDKETVSEVQGELDALSKLSWTFKNDDCFREQKAKFVKLSVDITKLMAGLNIQLNLEPLKAPRDEPELKTHWETLSTVWLPEINRQMHQRKLELETELSRFANKFAAAEERIQIESETLCEAKKREAREERVRRNERDEWKTARQELEKRYDALENEKEVLKRQMQIGGNLVKEIDVEREKMAKIEEALRAKEAELQTEIRLKLDEMFAQIAKIDAEKMMIFKQFTDLEAIMNANCAHY

>C_wallacei_NDC80_paralog1

MYGNERRKTGAPNFSGRASLAITPSKRFTDVDMTSIRRSEVRPSLMGGQSRPSIFGKGSTLPPRDVKSMLYVNVQKIYHFLVEHDQGDAPSEQVIKSPQGKKDFIAIFESIYQHMSKDYEFPDGKRLEEEIIPLFKALGYPYPLKDSYFKPIGAGHGWPHLLDALGWLVDLVRIMLSVKNDTQNIMFGDFMEANKVQEKTLSYSWHTNIFREFTNDRKVIESREHPFWEEQKTNLRNVFDSSNEYGDMTNNYKNALEQLKFDCDEIESERGQETNLQEEIARMSDDIRKANDYREQTEQVRTLRETDFQKIQEALVEKKEENIRMQNTVNELKERIEQQKINHGLSGKQVREMNLENSKDKEHVSEIQGELDKLSKEMWTLNNENCFREQKAKFTEIIDHIQKTINGLDISIQLPHLKPPTDEKQLKSGWETLTNNWIPEINRHLHQKKLELETEQSRFADRFAATQERIQIEIEMLQESKKRETREDRIRQNERSEWKSARQELEKRYDEQENRLEVLKKQLKMDGSLEKEIEEEKEKMRKIEEEMTAKQEELVRILNGRLDEIVAMTAKIENEKLGIHKEFEEIERVILANQ

>C_wallacei_NDC80_paralog2

MYGNERRKTNFDGRPSNAITPNKRHTEGISSLRRSEVRPSIGGHPRQSIFGRASTIPSRDVKALAHGNIDKIYNFFVEKGEEPPCKSIIQSPTGKDFAVIFETIYQHLSEGFVCSYQHPEEDVIPIFKALNYPYTLKDSYFKPVGTSHGWPHLLDALGWLIDLVNISLSVNNNAQNMLFGDILERNQVEEKTLSYAWYTSIYRDYTNDRKVIENRNIPFWEKQRDALRNAFNGFDGNMDVIGRMKNELEQLKFDCDEIESETGQENQLKEEISRIKDDIRKADEYFEQTEQLREIREKNFQKILEELQEKQKENERIQKKVNEVKERIEEQIINHGIDGKQVREMNLENSKDKELVSEMRGELDKLSKEMWTLKNEDLFKEQKGKFQEVIDHIYKTINGLEIQITLTPMKSPSDERELKIGWDTLTNIWIPEINRHLHQRKLELETEQSRFSDKFAASQERIQIELEMLEEAKKREAREERQRQNERDEWKVSRQNLEKHYDELENEMEVLKKKLKMDGSLEKEIEEERVKIMKIEEELSSKHNELINEITRKLDEIFAATAEIENEKLLIQKEVNELENAMKSVGLQ

>C_tropicalis_NDC80

MYGSERRKTGAMNLGGRTSMAITPSRRFTDVESSSVRRSEVRPSLMGSQARPSIFGKGSTIPPRDVKSMIGANVEKIWHFLVEHDQGDAPSDSVIRNPQGKKDFVAIFESIYQHMSKDYEFPNGGRLEEEITPLFKALGYPYPLKDSYFKPIGATHGWPHLLDALGWLVDLVRIMLSVKDDTQNIMFGDFPEATEVHERALSYSWHTSIYREYTNDRKVIESKGHPFWEKQRAELRNVFDSSNEYGDMTSNYKNALEQLKFECDEIESERGQEANLLDEIARMKDDIRKAAAYQEQTENVKMLREADLEKLHAVLREKMEENQKMQKAVNELKERIEKQRVDHGLNGKQVRQMNLENSKDKEMVSEIQAELDKVSKETWMLNNENGFRDQKLKFLEIHEHIIKTIHGLNLPVNLPPLKPPSDELELKSGWETLTTVWIPDITRHLHQKKLELENEQSRFADRFAAAEERLQIENEMLEEAKKREQREERTRQNERSEWKTARQRLEKESDELENQLEVLKKQMQIGGNLEMEIEEERRKMKAIEEGMERKNEELIACLQKKLDEIFAATVEIENEKLAMQKEFEELEAVMQTNCH

>C_doughertyi_NDC80

MFGGERRKTGGMSLTGRASLAITPTKRFTDFGISSSRTNPRPSLNIGPRQSLFGSKSASVAPRDVKSIQQGNVQKIYRFFVETDGDDAPPESIIRAPGGKADFITIFEQIYQHLSKDYEFPQGARLEDEIAQIFKGLGYPFPLKKSFFQPMGCAHGWPHLLDALGWLVDLVRISIRVSSDTQNILFGDFMEESKLTSKFLNYAWHTKVYKDYTNDRKVIDNPHSQFWLEAKSQLRDHFETANEYDDWMDTAKNNLEQLKFDCDEIESERGQEQHLLDEIAKMKDDIRKADDYLQQTANLKVHKETALQEVQQTLLEKQGELEKIQKVVNELKERIEQQKVKHGLSGKEVRQMNLENQKDKETVSEIQGVLDQLSKEAWSLKNDDSFKEQKAKFVQLAEHITKIMSGLDIEINLARLSVPSDERQLKANWETLTNVWMPEVNRQLHQKKLDLETEISRFTDKFAAAEERNKIESEMLCEAKKKDAREERLYLNERKEWQEARQQLQKRCDELENEKEVLKKEMKMDGSLEKEIEEEKTKMAKEVKELRAKQTELEAVLRQKLDQIVAEVTDIQNEKLMVYKEFNDLGMIFESKLPNY

>C_sp54_NDC80_paralog1

MFGADRRKTGGGMSLGPRQSVAITPTKRFTDYGPSSMRKPDGRASLSHNIGQPRPSLFGTKNSAVAPRDVKGLVTANVSKIYSFLIEFEGSEAPSEKDIKSPRGKNDFTFVFELIYQHLSKDYEYPQSARIEDEIPQIFKGLQYPVTLKNSYFQPMGSSHGYPHLLDALAWLVDLVRINSCVSADTQNIIFGDFMEQAIVQEKTLNYAWMSTTFRDYTNDRKAAENPETPFWDETKCRLRKYFEESNEFDDMAKTASNGLEQLRFECEEIESERGNEQSLLEEIAKIKDDIRKATEFFESTEQVKELKEKELARVKLELEAKMAEHERAQRMVNELKARIEMQKTNHGLSGKEVRQLNIENNRDKETVAEIQAELDSLSKEAWRLNKDDSFKEQKAKFVQLYQKTMKIIAGLDIQLNLDPIKAPADERQLKTCWEALNGIWLPEINRQLHQKKLELETKNSRFADNFAHTEQRIQMESEMLCEAKKKEAREERIRRNERDEWKEARQQMEKRYDELENEKEVLKKQMQMDGSLEKEIEAERVKMTKIEKELEEKKKYLETSIRQKLDQIVAETAEIENEKIMFHVECTELEKTVKAMCLN

>C_sp54_NDC80_paralog2

MNSTFILNHFSLLRHVKKMFSGGPRDVKSLVTANVKKIYSFLIEFEGSEAPSEKDIKSPRGMNDFISIFESIFHHLNKDYKYPQSARVEEEISQIFKGLQYPFPLKVSFFQPFGSSHGYPHLLDALAWLVDLVRMNSCTNDTNEKEMLKMQMTKIEKELEEKKKNLETSIRQKLDQIVAETAEIENAKMMLLEMKCTELEKTVKAMCLN

>C_inopinata_NDC80

MYGDRRRTGGISLNGRASLAITPTKRYTDFRTETRPSLGQSSGQHRLSIFHKTPSVAPRDVKSIFTANVSKVHNFLAEHEGLEAPLESTIRNPRGRNDFICCFELIYQHLSKDYEFRKDERIEDEISQIFKGLGYPFALKNSYFQPMGSSHGYPHLLDALAWLVDIIRVNAAVSADTQNIIFGYFMEQSKAQERTLHYAWMSSIFREYTNERKVAENSESPFWGEVKNKLRKFFEDSGEFEDMTNSAASALEMLRFECDEIESEKDNEQSILEEITRIKDDIRKASAYLETTQQVLQHKEIELAGVQSEVADKIAENEKVQKMVNDLKTRIENQKNHHGLTGKEVRELNLENNRDKEMVSEIQAELNNLSKETWRLKDEDHFKEQRSKFVQLFDSVTKMIAGLNIQMNLKQISSPNNERELKVCWETLDNVWLPEINRQLHQRKLDLETEQSRFSSKFVAAAERIQMKSEILSEVKKNEIREERLRQNERDEWKENRKQKEKRYDELENEKEVLKTQMQMDGSLEKEIEAERISMAKIEKELEEKCKYLQSVIKKKLDQIVTETTAIESQKTIFHLQCTEIEKAVN

>C_elegans_NDC80

MFGDRRKTGGLNLNGRASIAITPTKRFTDYTGSTSVRKTDARPSLSQPRVSLFNTKNSSVAPRDVKSLVSLNGSKIYNFLVEYESSDAPSEQLIMKPRGKNDFIACFELIYQHLSKDYEFPRHERIEEEVSQIFKGLGYPYPLKNSYYQPMGSSHGYPHLLDALSWLIDIIRINSAVSEDTQNILFGDFMEQGKAQEKTLNYAWMTSTFRDYTNDRKAAENPSSSYWDDTKHRLRKYFEQSNEFEDMTKTAASALEMLNYECDEIEADKGNEASLKEEISRIRDDIRKAKDYLEQNLHVKQHMEKELAMVKSEQEEKISENEKVQKMVDDLKNKIELQKQIHGLTGKEVRQMNLDNNKDKEVVLEIQSELDRLSKETWKLKDEDFFKEQKSKFIHLAEQIMKILSGLNIQMNLEPLRAPTNERDLKDYWETLNKIWVPEISRQLHQRKLELETEQSRFSNKAVTAEERIQIQSETLCEAKKNEAREERIRRNERDSWKDARKHIEQRYEQLLNEKEVLLKQMKLDGSLEKEIEDETARMSATGEEHIQKRSQLEAGIRQILDLMVVEIAEIENKKIGFHVQCAGIEKAVL

>C_oiwi_NDC80

MFGDRRKTGGLNLNGPRASLAITPTKRFTDLGVNSVRKSTTRMSLSQTSGQPRMSLFQSKSIAPRDVKALVSSNVTKIYNFLVEHEATEAPSEKDIRSPRGKNDFVLFFELIYGHLSKDYEFPQDTRLEDEIPLIFKGLGYPYPLKNSYFQPMGSSHGYPHLLDALSWLIDLAKISSSVRSDTKNIIFGDFMDQNKALEKILSYSWLSTVYKDFTNDRKAAETDQFWVETKGTLRKHFEHNNEYDDMIETAKSALEQLRFECDEIESDKGNEQSLQEEIARIRDDIRKASDYLEQTEMLKERTQEELNEINTQLQVKSAENEKVQGMVNELKARLEQQKLHQGLTGKEVRQMNLENNKDKETVAELQAQLEELSKQAWKLKNDESFKEQKTKFVHLVENALKVLSGIGINTNSDAFRVPTNEKELQASWETMNNVWLPEINRQLHQRKLELETEKLQFADKFASAEERIQMETEMLREAKKKEVREDRLRRTEREEWKESRQQLEKRLDELENEKEVLKKQLQIDGSLEKEIETEKVRMAKIEKEMEAQKERLEAAIRKKMDEIVEETAGIENSKIMFHKSCVEVEKTVKTALKVSKK

>C_kamaaina_NDC80

MFGDRRKTGGLNLNGPRASLAITPTKRFTDLGVNSMRKSTARVSLSQTSGQPRMSLFQSKSTAPRDVKALVSSNVTKIYNFLVEHEATEAPSEKDIRSPRGKNDFVVFFELIYGHLSKDYEFPQDTRLEDEVKKIFKGLGYPYPLKNSYFQPMGSSHGYPHLLDALSWLIDLAKISSSVRSDTKNIIFGDFMDQNKALEKILSYSWLSTVYKDFTNDRKAAETDQFWVETKETLRKHFEHNNEYDDMIETAKSALEQLRFECDEIESEKGNEQSLQEEIARIRDDIRKASDYLEQTEMLKERTQEELNEIKTQLDTKSVESEKMQGMVNELKARIEQQKLHQGLTGKEVRQMNLENNKDKETVAELQAQLEELSKQAWKLKNDESFKEQKTKFVHLVENALKVLSGIGINTNSDAFKVPANEKELQASWEAINNVWLPEINRQLHQRKLELETEKLQFADKFASAEERIQMETEMLREAKKKEMREDRLRRTEREEWKESRQQLEKRLDELENEKEVLKKQLQIDGSLEKEIETERVRMAKIEKEMEAQKQRLEAAIRKKMDEIVEETSGIENSKMMFHKSCVEVEKTVKALKVSNNQ

>C_waitukubuli_NDC80

MFGDRRKTGGPGFGPRLSTAITPTKRFTDVGPTSVRKADSRLSMGQGGGQPRASLFQRSSAVASRDVKSLMAANVNKIYNFFIESEGSEAPSEAQIRNPKGRNDFITFFELLYQHLSCDYEFPQGARVEDEVSALFKNLGYPTALKNSFFQPMGSSHGYPHLIDALAWLVDLIRVNMCVSNDTQNIIFGDFLEQSKVQERTLSYCWMSSTYKEYTNDRKAAETRDGPFWDSTKIKLRKYFEDMNETGDMAASAKTSLEQLRFECEEIEADKGTEQNLIEEIAKIKDDFRKASDYCERLEAVERQREVDLDQVKMEVDVKKKEMSEVQEEVNSLKQRIEEQKQNHGLTGKQVRQLNSDNSRDKAIVSEIQSELERLSKELWLRKNEENFKEKKTMFLQLAEKLAKIGAGINVDLGLTSLRSPQNEELKIECEKLVGVWLPEVTRHLNQKKLELEMEKSRFNDKFASAACANMEQETLCEAQKKAAREERIRRNEREEWTESRLAKEKRYDELENELDVLKKNMQMGDSLVKEIEMERAKKVKLCEEQEKQKKTIEASIRQKMDQIMTETAEIDNEKMMIHVECNELEKMV

>C_panamensis_NDC80

MFGDRRKTGGPGFGPRLSTAITPTKRFTDVGAAASVRKVDNRLSMGGQHRASLFQRSSAIAPRDVKSLMSANVSKIYNFFLEFEGPDAPSESQIKNPKGRNDFISFFELLYQHLSCDYEFPANVRVEDEVQISTIFKNLAYPTPLKNSFFQPMGSSHGYPHLIDALAWLVDLIRNKCVSDDTQNIIFGDFLEPAKVQEKTLSYAWMSSTYREYTNDRKAAETKDGPFWEVTKNKLRKYFEDMSETQDLAASVKNSLEQIRFECEEIEADKGTEQSLIEEIAKIKDDLRKATDYAEKLEAVERQRLMDLDKVKADLDVKKKENSEVQTEVDGLKQRIEEQKQNHGLTGKEVRQLNSDNARDKAVSNELQSEMEKVAKDLWRNNNEENFKEQKRTSFSRLVERIEKILAGINVTLRLEPLRTPQTERDLKAGLDELMSVWLPEITRQLNQKKLELNMEESRFTDKFVAVQARVQLEKETLDEAKKKEAREERMRRNEREEWMESRLATEKRYDEMENEQDVLKKQLQMDGSLEREIEMEKARKVKISEEQENKKNKIDARIRQKMDEVMKETAEIDNGNLLFHVACNELEKMVLKTQNLI

>C_nouraguensis_NDC80

MFGERRKTGGQGFGPRISTAITPTKRFTDVGAAASVRRTDSRLSMGQSGGQHRASLFQRNSAAAPRDVKAMLGPNVNKIYNFFIEFEGSDAPAEAQIKCPKGRNDFINFFELLYQHLSCDYEFPPNARVEDEVLWIQIFKCIGYPVSLKNSYFQPMGSSHGYPHLVDALAWLVDFIRNKCVSDDTQNIIFGDFLEPAKVQEKTLSYAWMSSIFREYTNDRKAAETRDGPFWTITKERLRDYFEKMNEAQGLAVSLKNALEQIRFECEEIEADKGQGQTMLEEINKMKDDMRKAVDYNEALQTAQRQREIDLEKAMTDLEVRMKEDSEVQADVNELKKRIEEQKEKHGLTGKEVRQLNLENTRDKAVISEIQAELDKLSKELWLKKNEENFKDKRASFVRLAENIGKIVAEVYIDLGLEPLRSPQNEKELKSELEKMNSVWLPEVTRQLRQKQLNLNKEKSFFRDKERVQIERDTLCEAEKKMAREERIRRTERDEWKEIRLANEKRYDELENEQDVLKKQMQKDGSLDKEIELEMARKLKIDEESSKKKSALEAVIRRKLDHIMAETSKIDNEKLMFHTDCTEFEKQILKTRNMK

>C_becei_NDC80

MFGERRKTGGPGFGPRISTAITPTKRFTDVGAAASVRKTDSRLSMGQPGGQHRASLFQRNSAAAPRDVKSMLGANVSKIYNFFIEFEGSDAPAEAQIRNPKGRNDFINFFELLYQHLSCDYEFPPNARVEDEVIHIFKCIGYPVSLKNSYFQPMGSSHGYPHLVDALAWLVDFIRNKCVSDDTQNIIFGDFLEPAKVQEKTLSYAWMSSIFREYTNDRKAAETRDGPFWTVTKEKLKDFFEKMNEAQGLAASLKNALEQIRFECEEIEADEEINKMKDDMRKAVDYNDALQTAQRQREIDLEKAMTDLEIRMKENSEVQADVNELKQQIEEQKEKHGLTGKEVRQLNLENTRDKAVISEIQAELDKLSKELWLKKNEENFKDKRAIFVQLAEKIGKIVAEVHIDLGLESLRSPQNERELKSEQEKMNSVWLPEVTRQLRQKQLNLNKEKSFFRDKFAAAEERVQIKKVTLCEDEKEMAREERLRRNEREEWKEIRLVNEKRYDELENEQDVLKKQMQRDGSLDKEIELEMARKLKIDEESSKKKSALEAKIRQKLDQILVETSKINNEKLMFHSDCQEFEKQILKTRNMK

>C_yunquensis_NDC80

MFGDRRKTGGPGFGQRNSTAITPTKRFTDVGAASSVRKTDNRLSMGQTGGQHRASLFQRNSAAAPRDVKSMMGANVSKIYNFFLEFEGPADAPSEGNIRSPKGRNDFINFFELLYQHLSKDYEFPANARVEDEIIHIFKFLGYPISLKNSYFQPMGSSHGYPHLVDALAWLVDFIRCSISVSGDTQTIYYGDFLDQAKVQERTLSYAWMTTIFREYTNDRKAAETRDGPFWTMTKEKLRTYFEKMNEDQDLTASVKNALDQIRFECEEIEADKGKGQEISEEIVKVKDDIVKAIDYGEALEAAQKQQESDLEKATAELRERVDENSEVQANVNELKQRIEDQKKKHGLTGKEVRQLNSENTRDKAMITEIQAELDKLSKELWLKKNEENFKDKRESFVQLVERIGKIVAEVSIDLGLEPLRSPRNERELKTELEKLNGIWLPEVTRQLRQKQLNLDKEKAFFRDKFAAAEERVQIERDTLCEAEKKMAREERVRRHERDEWKESRLVIEKRYDELENEQDVLKKQMQMDGSLDKEIELEKAKKVKIDEEIAKKKAALEAAMRQKLDKIMVETAKIDNEKILFHVECTEFEKQILKTRNLN

>C_macrosperma_NDC80

MFGDRRKTGGPGFGPRISTAITPTKRFTDVGAAASARKVDSRLSMGQPGGQHRASLFQRSSAVAPRDVKSMLAANVNKIYNFLVEYEGTDAPSEAQIRCPKGKNDFINFFELLYQHLSCDYEFPANGRLEEEISNTFRHLGYLPHLKNSYFQPMGSSHGYPHLVDALAWMVDIIRCNKCVSDDTQNIIFGDFLEPAKVQEKTLSYAWMSSTFREYTNDRKAAEAINGPFWNSTKNKLRKFFEDMNDTQDLTASAKNALEQLRFECEEIEADKGTEQSLLEEITRIKDDIRKAICHYEDTKRVLNQRESDLEKVKAELDTRIKENNEVQAEVNLLKNRIEEQKEKHGLTGKEVRQLNSNNTRDKAVVSEIQTELDKLSKELWLMKSEDTFKEKKTCFVHLTERIGKIVAGIEIDLGLEPLRSPQNERELKVEWEKLNNVWLPEITRQLHQKKLELEMEKSRFSDKFAAIEERVQMERETLCEAKKKESREERLRRNEREEWKESRLLKEKRYDELENELDVLKRQMQMDGSLDREIEIEKAKKVKNDEELKKKKAAIEASIRQKLDQIVAETAKIENEKTLFHVECIEFEKLILNTQNLV

>C_sulstoni_NDC80

MYGGDRRKTGGFGFGNPRTSAITPTKRFTDLGTTSSVRKDHGRLSMSQTSGHHRVSLFGGASATGPKDVRANFDANAKKIYNFLVEHEGSEAPSENVIRCPAGKSDFVTVFELMYQHLSKDYEFPQNVRVEEEVSNIFKALGYAPPLKNSYFQPMGSAMGYPHLVEALAWLVDLIRINASVSADTQPIIFGDVMDQEKVHEKTLSYAWMSTTFRDYTNDRKAAESGTGPFWEETKSKLRDYFEKSDEIEDVATNYKSALEQLRFECEEIEAERGNELTLLEEIAKIKDDVRKALEYDETTARVQKHKEDELKAVKGTLEQKMAENSKVQAEVAELKERIEVQKQMHGLTGKEVRQLNSDNNRDKETVTELQAELDEISKQMWRLKNEDTFKQQKTNFVRLVESVEKIVSGLDIEMGLEPLRAPSNESELKSRWDELNSVWIPEINRQLHQKKLEFETEQSRFADRFAAAERVQMERELLCEAKKKEAREDRVRRNERDEWKENRLQREKKYDELENEKDVLTKKMQMDGTLDKEMEEERTKAWKLEQELQKEADAVENAIRKNLDLIVNEVAQIENDKMLFHVECCEFEKLIVKTQNMF

>C_afra_NDC80

MYGGDRRKTGGFGFGNPRTSAITPTKRFTDLGTTSSVRRVSLFGGGSAAGPKDVKGNFEANVKKIHSFLIEHEGNEAPADTVIRSPAGRNDFLTVFELMYQHLSNNYEFQQGVRIEDEVSNLINASVGADTQTILFGDVMDQAKVQEMTLRYSWMSTTFRDYTIDRRAAESGTGPFWDETKNKLREYFEKSDEIEDLVTSYKTALEQSAFECQEIEADKGNEQHLLEEISKMKDDVRKAMEYAESTARVQKHKEEEMKTVKATLETKIAENSKVQAEVAELKERIEVQKQLHGLTGKEVRQLNSDNNRDRETVTELQAELDEVSRQMWRLKSEDTFKQQKANFIRLVESVEKIVSGLELQLGLDAILAPSDESELKVRWDELNNVWIPEVNRRLHQKKLEFETEQSIFVDKFAAAERVQMERELLCEAKKREAREDRVRRNERDEWKENRLQREKKYDELENEKYVLKKKMQMDGSLDKEVEEEQAKARKIEEELQKEADAIENSIRKNLDQIVSETAQIENDKMLFHVECCEFEKLI

>C_sp49_NDC80

MMFGGPRRKTGGPNFGATGRTSTAITPTKRTTDLGAAHSVRKNDNRMSFGQGPSNSRPSLFQRTGGPSKDFKAAVPGNAAKVYNFFVDTEGANAPSEKTIRTPAGKGDFVTMFELLYQHLSKDYEFPSNGVRLEDEFTNIMKALGYPHALKPSFFQPIGSSHGYPHLLDALAWLVEAVEINEKVRLDTQNIIMGDFMEPDQVQDKVLSYSWLSKTFFEYSNNRKVLEDKADPFWATTKCDLRNFFENNDESADMITSSKNMLEQLRFDCEEIEADKGNEQGLLDEIARIRDDIRKAQLYLDETMRVVAQSNHEYQTVVDACEAKLAELEKVKTEVAELKERIEEQKRLHGLSGKEVRQMNAENSKDKEMLSDVNAEIERATKMMWRLRDAADYKEPSTRFYRLISHMHQILDDADVHPQLEPIASPSSELDLKAGWDAINNLWLPEVNRQLQHKKLELETENARFLNKFEAIEERATMEQEMLNEATKKEDREERVRRNEREEWKAARFQKEKRCDQLENEKDILVKQMHLDGSLDRDLQEETEKAAKLSETLEQKQAAFEQKVREKIDEMMASTAEMEQEKLEFHKECVELEMVVKNS

>C_sp25_NDC80

MFGGPRRKTGGPGFSTTGRTSTAITPTKRNTDLGTAQSVRKNDGRMSFGANPSGSRPSLFQRTVAPSKDVKAAVPANVARIYNFFVETEGPNAPSEKMIRSPAGKGDLITMFELIYQHISKDYEFPSVGVRLEDEFTNIMKALGYPNALKQSFFQPIGSSHGYPHLLDALAWLVEAVEINEKIRLVTQNILVGDFMEPDQVQDKVLSYSWFSKTFLEYTNNRKVLEDRSDPFWANTKRELRDFFERNDESADMIATSKNMLEQLMYDCEEIEADKGNEQGLIDEIARIRDDVRKAQAYLEETSRVVAQSIHEFQSVSDAFEAKRTEFEAVKTEIAELKERIEEQKRNHGLCGKEVRRLNLENIKDKETVAELQAELEKTSKIIWNLRDAGSFKEQKARFDRLITHMKMILDDTDIHFQLESITSPRTELELKAGWDTVNNVWLPEVNRQLQRKLLELETENARFASKFGAKEEHVTMEQEMLNEATKKEDREERVRRNEREEWKAARFQKEKRFDQLENEKDILMKQLHLDGSLDRELEEETEKSIKLTEALEQKKAEFEQKVREKIDLMMVSTAEMEQERLDVHKEFVELETIINSS

>C_imperialis_NDC80_paralog1

MFGGPRRKTGGPNFSATGRTSTAITPTKRNTDLGAVQSVRKTDTRMSFGQGPSAPRASLFQRAGLPSRDVKACFASNVSRIYNFFVETEGSSAPSEKTIRSPNGKGDFISMFELIYQHLSKDYEFPTQNRLEEEFSNILKSLGYPYPLKNSFFQPIGSSHGYPHLVDALAWLVEVCEVNEKVRLATQNILLGDFMEPEQMKDKFISYSWFSKVFLEFTNQRKAAEDKTDPFWESTRRELREFFEQNDESVEMIASLKNMLEQLRFDCEEVEEDKGNEQSLLDEISRLRDDVRKAQSYLEETARVVAQSHHEFATVKEAVDAKMEEFERVKAEVAELKARIEQQKELHGLTGKEVRLMNVENNKDKETVAEIQNELDRTSKIIWRLRDAGSFKEQKSRFERLVEHMLKIVDDADIHLTLDPINSPSTELELRAGWEAVNTVWLPEVNRLLQRKKLELEGENARFSSKFGAVEEKAILEKELLNEATKQEAREERVRRNERDEWKESRLQKERRCDQLENEKDVLVKQLHLDGSLDVEIREKADEMEKLVVELAGKKKTLEGEVRKRIDEMMESTVQMEHEKLLFHRECVELEALVKSQ

>C_imperialis_NDC80_paralog2

MFGNARRKTGGPNFATGRPSTAITPTKRITDLGAAQSVRKNDSRMSFGPSAHRTSLFQRAGPHSKDMKACFAANVAKIYHFFVETEGSEAPPEKMIRSPTKNEFITIFELVYQHLSKDYEFHMQNRLEEEFTSILKALGYPYPLKNSFFQPIGSSHGYPNLVDALAWLVEVVEVNSAVSRVTQNILIGDFMEPELAEDKVLSYSWFSKTFLEFTNNRKALEDKSDPFWESTRLGLREYFERNDETAEMIASCKNTLEQMRFDCEEIEEDKGNEQSLLDEISRLRDDVRKAEAYLLETSRALEQATQEHESVKEAVETKGAEFEQVKMEVARVKELIEQQKEQHGLTGKEVRAMNVECQRDKETVSEIQMELDKVSKVIWRLRDAGSFKEQKSRFDRLIEHMHTILVDADIHLELEPICSPSTEIELRAGWEAINGVWLPEVNRQLQHKKLELEAENARFSSKFGVVEEKAILEQELLNEAIKQEARDERVRRNEREEWKASRLQKEQRCDELENEQDVLVKQLNLDGNLDAEIREAAQKAERLFLELQAKKEQFEGAVRAKIDEMMESTVAMEQEKLMLHKECSELESSVKSTL

>C_japonica_NDC80

MFGERRKTGGANFGFAGGRPSTAITPTKRFTDLGTAQSTRRVDNRLSMGQNGGRLSVFQRGSAVAPRDVKSAQASNVTKIYNFLLEHEQSGAPPEKIIRQPTGKNDFITMFEQLYQHLSKDYEFPQGARVEDEFTGIMKGLGYPFPLKNSFFQPMGSSHGYPHLLDALAWLIDVISLNQSVSSVTQNILLGDFMEAAEAQDKIISYSFYSSTFREYTYDRKAIESKDAPFWAETKERLRDYFEKNDEITDLVTSTKQVFEQLRFECEEIEADKGDEQGLLEEIARIKDDVRKAHEYLESTERVQKQTGVELGIVQESLEAKKAEFEAVQAEVNELKRRIEVQRQQHGLTGKEVRQLNVENNRDKEAVHEIQTELDNVSKTLWRMRDEDTFREQKANFVLVIENMEKILLDANVKIGLDALRPPQNERDLKVGWDALNNQWLPEVNRQLQHKKLELDDEKTQFSSRFAAIEERVQMQQELLREANKKEAREERVRRNERDEWKTDRLQREKRLDELENEKDVLKNQMQTGGSLDREIEEEKAKSAKLVEALEQKKASLAEGIRQKLDETMREITKIDLENATFHAECTDIEKLVLKTRNMNQ

**Supplementary Data S9: All HIM-10 sequences used in this study**

>C_tribulationis_NDC80_paralog1

MFGGRRTGGPGFNAGRLSTAVTPTKRFTDFGIGSTRKAGRLSMSQGPRPSFFTKGSTVPPRDVKSLQAANVQKIYNFLVEQDGSDAPSESSIRSPGGGKDFKMIFESMYQHLSKDYEFPLNARIEDEVMPIFKGLGYPFSLKTSFFQPMGGPHGWPHLLDALAWLVDVIRMNLAVSVDTQNILFGDFLDHQKVQEKALSYMWYSTLFRDYTNDRKAAEDKDGVFWKEAKANLRQYFENSNEYEEMITNLRNVLQQLRFDCDEIEAEKGQEQTYVEDIARMKDDIRKAVEYLESTQRVKENKEAEVTSIKQELDMKNAEMEKCLGMVYELKERIEHQKQVHGCSGKEVRQMNLENSKDKETVSELQAELDEISKETWRLKNDDSFKEQKAKFVQLVENITKLLAGLDVKLNLDPLSVPSDEKELKAGWETLNSVWVPEISRQMHQRKLELETDKARFVDKFAAAEERIQIENEMLCEAKKKEGRDERIQRIEREEWKVGRQQLEKRYDELENEREVLTKKMQMDGSLEKEIKEEKDKMAKMESEAEAKSQEVQAAIREKLEEMLVEIAEIGQEKYMFHVEATEVSRLISGMCSLDS

>C_tribulationis_NDC80_paralog2_pseudogene

RNVFLLPIFSPAPRLFDQITFPNDINIKRTFYHFMGNTLVTKTIDEAKRIDQRYGGRYLITTFEGAIIDQSGNLTGGGTPLTGRMNVTGASSSRFNNDIERKNHIFKTKTRMQQTEREINSIEAMLNAEMTKMQTDKASAAALENEYNQLNETLKNLRSHVEKLHDQSMACDHRLAQIAPIEEITASIAEITQELEVLRERQKRQVAQNQEAQHIVSSLSSKI

>C_sp41_NDC80

MFGGRRTGGPGFNAGRLSTAVTPTKRFTDYGIGSTRKSEAAGRLSMSQGHRPSLFQKGSAVPPRDVKSIQAANVQKIYNFLVEHDGSEAPAESIIRTPRGKKDFEAIFESMYQHLSKDYEFPTQGRIEDEVTQIFKGLGYPYPLKNSFFQPMGASHGWPHLLDALAWLVDVIRMNLAVSVDTQNILFGDFLEQQKVQEKALSYMWYSTLFRDYTNDRKAAEDKDGNFWKETKANLRQYFENSNEYEEMVTNLRNVLQQLRFDCDEIEAEKGQEQTYVEDIARMKDDIRKAMEYLDSTQRVKENKESEVTSVKQELEMKTAEMEKAIGMVNELKERIEQQKLVHGCSGKEVRQMNLENSKDKETVSELQAELDEISKETWRLKNDDSFKEQKAKFVQLVENITKLLAGLDVQLKLEPLSVPIDEKQLKAGWETLNSVWVPEISRQMHQRKLELETEKARFVDKFAAAEERIQIENEMLCEAKKKEGRDERIQRIEREEWKVARQQLEKRYDELENEREVLTKKMQMDGSLEKEIKEEKDRMAKLESEAEAKSQDVQMAIREKMEQMVVEIAEIGQEKTMFHGESTDVARVIGGMCSLDC

>C_zanzibari_NDC80

MFGDRRKTGGPSFNGGRLSTAVTPTKRFTDARLSMSQGHRPSLFQKGSAVPPRDVKTLKAANVSKIFNFLVESDGSEAPSESTIRSPPGKNDFIAIFESMYQHLSKDYEFPASARMEEEVSSIFKGLGYPFPLKNSYFQPMGGAHGWPHLLDALAWLVDVVKMNQAVSRDTQNILFGDFMDQGKVQEKALSYMWYSKVFRDYTNDRKAAEDKDGEFWTNSGAELRRYFENSNDNEEMMTNLQNVLQQLHFDCDEIEAEKGQEQTYVEDIARMKDDIRKAAEYLESTQRVKELKQAEFTAVKQDLDSRKAELEKAVGMVIELKERIEQQKRIHGCSGKEVRQMNLENSKDKEMLSELQAELDEISKETWRLKNDDSFKDQTKKFVQVLVNIRKMLANLNIQLNLDLLQVPKDEQELKVCWETLNGVWVPEISRQMHQRKLELETEKARFVDKFAAAEERIQIENEKLCEAKKKEGRDERVQRIEREEWKVARQQLEKRYDELENEKEVLTKKMHLDGSLEKEIKEEKDKMTKMEQVAEAKRQDLEMAIREKLEKMVVEISEIGQEKIMFHAESTDVARVINEKCLLDC

>C_sinica_NDC80

MFGNDRRKTGGNFNTGRLSTAVTPTKRFTDFGMSSVRARHSISQSRPSLFTKGSAVAPRDVKSIQAANVQKIRNFLVETDGPEAPDEATIRSPRGKNDFIAIFESMYQHLSKDYEFPVQARVEEEVTQIFKGLGYPYPLKNSFFQPMGASHGWPHLLDALAWLIDVIRMNQRVSSDTQGIMFGDTFDQQKVQETALGYMWYSSLFRDYTNDRKAAEIKDGEFWVETKQKLRNFFENTNEFEDMAVNLKNVLQQLCFDCDEIEAERGQEQTYVEDIARMKDDIRKAVEYLESTQRLKELKETDVTSVKQELEFKKSEIEKVHGMVNELKERIEQQKRIHGCSGKEVRQMNQENGKDKETVSELQSELDELSKETWRLKNDDSFKEQKAKFVHIVENIMKILNGLKVEWNLAPLPVPSDERQLKTCWETLNGVWVPEISRQMHQKKLELETERARFAHKFAAAERIQIESEMLREAKKKEGRDERIQRIEREEWKVTRQQQEKRYDELENEKEMLKKNMHLDGSLEKEIKEETEKMIKIETESEAKSQDLQATLRAKMEQILVEIA

>C_nigoni_NDC80

MFGGERRKTGGMNLNGRQSLAITPTKRFTDHGLSSVRKTDARPSMPLHGQSRPSLFQKGSTIPPRDVRSLQAANVQKIYNFFVETDGAEAPSERSIRAPGSRREFVVLFESMYQHLSKDYEYPEPARLEDEVTQIFKGLGYPYPLKNSYYQPMGASHGWPHLLDALSWLVDVIKMNTTVAANTQGILFGDILEQSKVQEKVLNYSWFASIYKDYTNDRKGTEDKDSQFWKDAKNKLRQHFENSNEYDDFASNAKNVLQQLIFDCDEIESERGQEQTYVEDIARMRDDIRKAAEYLQSVELVKEHKDAEMVKVKGELDSKVAEKEKLLRMVNELKDRIEQQKIIHGCSGKEVRQMNLENSKDKEMVAELQAELDEVSKEMWRMKNDDSFKEQKAKFLQIVENITKLLSGLNVQLNLDPMPVPADEKQLKTCWETLNSVWVTEISRQMHQRKLDLDTEKSRSLDRFAAAQERIQIENEMLCEAKKKEGRDERTRRAERDEWKAARQQQEKRYDELENEKEVLMKKLHLDGSLEQEIKEEKDKMAKIEKEAEEKTQYLRSAIRQKVEAVEMELAEIGQEKTMFHAECVAVEKLVQGTCGATH

>C_briggsae_NDC80

MFGGERRKTGGMNLNGRQSLAITPTKRFTDHGLSSVRKRTDARPSMSLHGQSRPSLFQKGSTIPPRDVRSLQAANVQKIYNFFVETDGAEAPSERSIRAPGSRREFIVLFESMYQHLSKDYEYPEPARLEDEVTQIFKGLGYPYPLKNSYYQPMGASHGWPHLLDALSWLVDVIKMNTTVAANTQGILFGDFLEQSKVQEKVLNYSWFASIYKDYTNDRKGTEDKDSQFWKDAKNKLRQHFENSNEYEDVASNAKNVLQQLLFDCDEIESERGQEQTYVEDIARMRDDIRKAAEYLDSVERVKEHKDAEMVKVKGELESKVLEKEKLLRMVNELKDRIEQQKMIHGCSGKEVRQMNLENSKDKEMVAELQAELDEVSKEMWRMKNDDSFKEQKAKFLQIIENITKLLSGLNVQLNLDPMPVPADEKQLKVCWETLNTVWVTEISRQMHQRKLDLDTEKSRSADKFAAAQERIQIENEMLCEAKKKEGRDERTRRAERDEWKAARQQQEKRYDELENEKEVLMKKLHLDGSLEQEIKEEKDKMAKIEKEAEEKTQYLRSAIRQKVEAVEMEIAEIGQEKTMFHAECVAVEKLV

>C_remanei_NDC80

MFGTERRKTGGVNLNGRSSIAITPTKRFTDFGATSVRRTDGRPSLSVGQLGQQSSRPSIFKTSVHAPRDVKSFQAANAHKIFNFLVESDGADAPAESVIRTPRGKNDFIVIFESMYQHLSKDYEFPPLNQARIEEEVSSIFKGLGYPFHLKNSYFQPMGASHGWPHLLDALAWLVDFIKINKSVSADTQHIIFGDFLEPAKVQEKALSYAWFSTTFRDYTNDRKSAESSDSEFWVETKNKLRKYFEDSNEYEDMTANAQSALQQLRFDCDEIESERGQEQTYVEDIARLKDDIRKAMDYFESVQHLKERKENETAVIKEELEAKVAENEKIQMAVNELKERIEQQKRVHGLNGKEVRKMNLENSKDKDTVSDLQSEQEMLSKQLWRLRDENPFKDQKMKVIQIAENVTKILSGLNMQFELESLRPPENEKELRACWEILIGSWLPEINRQLHQRKLDVETEKSRFHQKFAAAERIQIENEVLDEVKKKEAREERIHRDERDEWKEARQKQEKRYDELENEKEMLKRKMQMDGSLEKEIKAEKEKKEKIKKDLEEMHQGLEAVFRKKLEKIASETAEIGNEKTMFRIEAIEVEKLIDRKCSNQKV

>C_latens_NDC80

MFGTERRKTGGVNLNGRSSIAITPTKRFTDFGTTSVRKRTDGRPSLSVGQLGQQSSRPSIFKTSVHAPRDVKSFQAANAHKIFNFLVESDGADAPAESVIRTPRGKNDFIVIFESMYQHLSKDYEFPPLNQARIEEEVQSIFKGLGYPFQLKNSYFQPMGASHGWPHLLDALAWLVDFIKINKSVSADTQHIIFGDFLEPAKVQEKALSYAWFSTTFRDYTNDRKSAEYSDSEFWIETKNKLRKYFEDSNEYEDMTANAQSALQQLRFDCDEIESERGQEQTYVEDIARLKDDIRKAMDYFESVQHLKERKENEMAVIKDELEAKVAENEKIQMAVNELKERIEQQKRVHGLSGKEVRKMNLENSKDKDTVSDLQSEQEMLSKQLWRLRDENPFKDQKMKVIQIAENVTKILSGLNMQFELETLRPPENEKELRACWEILIGSWLPEINRQLHQRKLDVETEKSRFHQKFAAAEERIQIETEVLDEIKKKEAREERIRRDERDEWKEARQKQEKRYDELENEKEMLKRKMHMDGSLEKEIKAEKEKKEKIKKDLEEIHQGLEAVFQKKLEQIASETAEIGNEKTIFRIEAIEVEKLIDRKCSNQKV

>C_sp51_NDC80

MFGERRKTGGMSLGGRQSIAITPTKRFTDLGIASVRKTDARSSMSVNQPRMSLFQNKGSSLPPREVKSIQQANVQKIHQFLVESDGDDAPAEVVVRTPRGKKDFTMVFEIIYQHLSKDYEYPHPVRLEDEIPQIFNGLGYPYPLKNSYFQPLGAGHGWPHLLDALGWLVDVVKMSKAVVPNTQSIIFGDFLEQSKVQEKTLNYAWTTNTYRDYTSDRKVIENREHPFWEQAKSQLRNYFESSNEYEDMESNAKNALEQLMFDCEEIESERGQEQHLQEEIAKMKDDIRKAEDYLLQTENLKSHKEKDLGKIRETLVEKQSELEKIQQLVNELKERIEKQKINHGLSGKEVRKLNLENFKDKETVSEVQGELEKLSKEIWTLNNDDRFKEQKSKFVQLTDNIAKLTTGLNIQLSLEPLKAPTDERQLKTHWETLSNIWLPEINRQLHQRKLELETELSRFADSFAAAEERIQIESETLCEAKKREAREERIRRNERDEWKTSRQELEKRYDELENEKEVLEKQMQIGGSLDKEIDEERARMLKTEQDLRAKEADLMAAIRQKLDGLFAGIVELDNEKMMILKEFNDLEMVMNVKCANY

>C_sp44_NDC80

MFGNDRRKTGGMNLGGRASIAITPTKRFTDFGTGSVRKTDARLSVNQPRPSLFQKGSSLPPREVKSIQQANAQKIYQFLVENDGDDAPSEASIRTPSGKKEFGAIFEAIYQHLSKDYEFNHQARAEDEVTPIFKGLGYPYPLKNSYFQPMGSPHGWPHMLDALGWLVDFVKINKEVNKDTQNIIFGDFLEQNKVQEKTLNYAWNTNTYREYTLDRKVLDDQSHPFWEQTRIQLREYFESSSDYGDMATNIKNALDQLMFDCEEIESERGQEQQLQEEIAKMKDDIRKAEDYLLQTENIKSHKEKDLEKTLQCLVERQSELEKVQIAVNELKERIEKQKINHGLSGKEVRKLNMENQKDKETVSEIQGELDKLSKKTWTLKNDDCFKEQKSKFVQLTDQISKLMSGLSVQLNLQPLKAPSDEQELKTHWETLSNIWLPEINRQLHQRKLELDTELSRFGDKFSAAEERIQIESEMLCEAKKREAREERIRRNERDEWKTSRQEMEKRYDELENEKEVLKKQIQIGGNLDKEIEEEKTKMVKIEEALRTKEADLIAKLRQKLEEIVVGLAEIDQEKMRIHKEFNDLEIVMNKKCAHY

>C_sp48_NDC80

MFGNDRRKTGGMNLGGGGRASIAITPTKRFTDFGASSVRKVDPRLSVNQPRPSLFQKGSSLAPREVKSLQAANIQKIYQFLVESDGDDAPSEQTIRAPRGKTDFVMIFETIYQHLSKDYEFPQQTRLEDEVVLIFKGLGYPFPLKSSHFQPMGTGHGWPHLLDALGWLVDFVKISKGVSTARQNILFGDFLEQDKVQEKALSYAWNTNTYRDFTSDRKVLDNRDHPFWEQTKSQLRTFFESSNEYEDMANTGKNALEQLMFDCEEIESERGQEQHLQEEITKMKDDIRKAEDYLLQTENLKVRKEKDLMTMRETLAEKKAMLEKVLQAVNELKERIERQKIDHGLSGKEVRQLNLENMKDKETVSEVQGELDALSKLSWTFKNDDCFREQKAKFVKLTVDIAKLMAGLDIQLNLEPLKAPRDEPELKTHWETLSTVWLPEINRQMHQRKLELETELSRFADKFAAAEERIQIETEMLCEAKKREAREERVRRNERDEWKIARQELEKRYDALENEKEVLKRQMQIGGNLDKEIDEEREKMRKIEEALRAKEAELETEIRLKLDEMFTQIAKIDAEKMMIFKQFTDLEAIMNANCARY

>C_brenneri_NDC80

MFGNDRRKTGGMNLGGGRASIAITPTKRFTDFGGASSIRKVDPRLSVNQPRPSLFQKGSTLPPREVKSLQAANVQKIYQFMVEADGDDAPSEQTIRAPHGKTDFVMIFETIYQHLSKDYEFPQQTRLEDEVIQIFKGLGYPFPVKPSYFQPMGVGHGWPHLLDALGWLVDFVKINKAVNKDTQNIIFGDFLEQQKVQEKTLNYAWNTNTYREYTSDRKVLDNRDHPFWEQTRSQLRAYFESSNEYEDMASNVKNALEQLMFDCEEIESERGQEQHLQEEITKMKDDIRKAEDYLLQTENLKAHKEKDLVKVRENLAEKQAVLEKVLQAVGELKERIERQKIDHGLSGKEVRQLNLENMKDKETVSEVQGELDALSKLSWTFKNDDCFREQKAKFVKLSVDITKLMAGLNIQLNLEPLKAPRDEPELKTHWETLSTVWLPEINRQMHQRKLELETELSRFANKFAAAEERIQIESETLCEAKKREAREERVRRNERDEWKTARQELEKRYDALENEKEVLKRQMQIGGNLVKEIDVEREKMAKIEEALRAKEAELQTEIRLKLDEMFAQIAKIDAEKMMIFKQFTDLEAIMNANCAHY

>C_wallacei_NDC80_paralog1

MYGNERRKTGAPNFSGRASLAITPSKRFTDVDMTSIRRSEVRPSLMGGQSRPSIFGKGSTLPPRDVKSMLYVNVQKIYHFLVEHDQGDAPSEQVIKSPQGKKDFIAIFESIYQHMSKDYEFPDGKRLEEEIIPLFKALGYPYPLKDSYFKPIGAGHGWPHLLDALGWLVDLVRIMLSVKNDTQNIMFGDFMEANKVQEKTLSYSWHTNIFREFTNDRKVIESREHPFWEEQKTNLRNVFDSSNEYGDMTNNYKNALEQLKFDCDEIESERGQETNLQEEIARMSDDIRKANDYREQTEQVRTLRETDFQKIQEALVEKKEENIRMQNTVNELKERIEQQKINHGLSGKQVREMNLENSKDKEHVSEIQGELDKLSKEMWTLNNENCFREQKAKFTEIIDHIQKTINGLDISIQLPHLKPPTDEKQLKSGWETLTNNWIPEINRHLHQKKLELETEQSRFADRFAATQERIQIEIEMLQESKKRETREDRIRQNERSEWKSARQELEKRYDEQENRLEVLKKQLKMDGSLEKEIEEEKEKMRKIEEEMTAKQEELVRILNGRLDEIVAMTAKIENEKLGIHKEFEEIERVILANQ

>C_wallacei_NDC80_paralog2

MYGNERRKTNFDGRPSNAITPNKRHTEGISSLRRSEVRPSIGGHPRQSIFGRASTIPSRDVKALAHGNIDKIYNFFVEKGEEPPCKSIIQSPTGKDFAVIFETIYQHLSEGFVCSYQHPEEDVIPIFKALNYPYTLKDSYFKPVGTSHGWPHLLDALGWLIDLVNISLSVNNNAQNMLFGDILERNQVEEKTLSYAWYTSIYRDYTNDRKVIENRNIPFWEKQRDALRNAFNGFDGNMDVIGRMKNELEQLKFDCDEIESETGQENQLKEEISRIKDDIRKADEYFEQTEQLREIREKNFQKILEELQEKQKENERIQKKVNEVKERIEEQIINHGIDGKQVREMNLENSKDKELVSEMRGELDKLSKEMWTLKNEDLFKEQKGKFQEVIDHIYKTINGLEIQITLTPMKSPSDERELKIGWDTLTNIWIPEINRHLHQRKLELETEQSRFSDKFAASQERIQIELEMLEEAKKREAREERQRQNERDEWKVSRQNLEKHYDELENEMEVLKKKLKMDGSLEKEIEEERVKIMKIEEELSSKHNELINEITRKLDEIFAATAEIENEKLLIQKEVNELENAMKSVGLQ

>C_tropicalis_NDC80

MYGSERRKTGAMNLGGRTSMAITPSRRFTDVESSSVRRSEVRPSLMGSQARPSIFGKGSTIPPRDVKSMIGANVEKIWHFLVEHDQGDAPSDSVIRNPQGKKDFVAIFESIYQHMSKDYEFPNGGRLEEEITPLFKALGYPYPLKDSYFKPIGATHGWPHLLDALGWLVDLVRIMLSVKDDTQNIMFGDFPEATEVHERALSYSWHTSIYREYTNDRKVIESKGHPFWEKQRAELRNVFDSSNEYGDMTSNYKNALEQLKFECDEIESERGQEANLLDEIARMKDDIRKAAAYQEQTENVKMLREADLEKLHAVLREKMEENQKMQKAVNELKERIEKQRVDHGLNGKQVRQMNLENSKDKEMVSEIQAELDKVSKETWMLNNENGFRDQKLKFLEIHEHIIKTIHGLNLPVNLPPLKPPSDELELKSGWETLTTVWIPDITRHLHQKKLELENEQSRFADRFAAAEERLQIENEMLEEAKKREQREERTRQNERSEWKTARQRLEKESDELENQLEVLKKQMQIGGNLEMEIEEERRKMKAIEEGMERKNEELIACLQKKLDEIFAATVEIENEKLAMQKEFEELEAVMQTNCH

>C_doughertyi_NDC80

MFGGERRKTGGMSLTGRASLAITPTKRFTDFGISSSRTNPRPSLNIGPRQSLFGSKSASVAPRDVKSIQQGNVQKIYRFFVETDGDDAPPESIIRAPGGKADFITIFEQIYQHLSKDYEFPQGARLEDEIAQIFKGLGYPFPLKKSFFQPMGCAHGWPHLLDALGWLVDLVRISIRVSSDTQNILFGDFMEESKLTSKFLNYAWHTKVYKDYTNDRKVIDNPHSQFWLEAKSQLRDHFETANEYDDWMDTAKNNLEQLKFDCDEIESERGQEQHLLDEIAKMKDDIRKADDYLQQTANLKVHKETALQEVQQTLLEKQGELEKIQKVVNELKERIEQQKVKHGLSGKEVRQMNLENQKDKETVSEIQGVLDQLSKEAWSLKNDDSFKEQKAKFVQLAEHITKIMSGLDIEINLARLSVPSDERQLKANWETLTNVWMPEVNRQLHQKKLDLETEISRFTDKFAAAEERNKIESEMLCEAKKKDAREERLYLNERKEWQEARQQLQKRCDELENEKEVLKKEMKMDGSLEKEIEEEKTKMAKEVKELRAKQTELEAVLRQKLDQIVAEVTDIQNEKLMVYKEFNDLGMIFESKLPNY

>C_sp54_NDC80_paralog1

MFGADRRKTGGGMSLGPRQSVAITPTKRFTDYGPSSMRKPDGRASLSHNIGQPRPSLFGTKNSAVAPRDVKGLVTANVSKIYSFLIEFEGSEAPSEKDIKSPRGKNDFTFVFELIYQHLSKDYEYPQSARIEDEIPQIFKGLQYPVTLKNSYFQPMGSSHGYPHLLDALAWLVDLVRINSCVSADTQNIIFGDFMEQAIVQEKTLNYAWMSTTFRDYTNDRKAAENPETPFWDETKCRLRKYFEESNEFDDMAKTASNGLEQLRFECEEIESERGNEQSLLEEIAKIKDDIRKATEFFESTEQVKELKEKELARVKLELEAKMAEHERAQRMVNELKARIEMQKTNHGLSGKEVRQLNIENNRDKETVAEIQAELDSLSKEAWRLNKDDSFKEQKAKFVQLYQKTMKIIAGLDIQLNLDPIKAPADERQLKTCWEALNGIWLPEINRQLHQKKLELETKNSRFADNFAHTEQRIQMESEMLCEAKKKEAREERIRRNERDEWKEARQQMEKRYDELENEKEVLKKQMQMDGSLEKEIEAERVKMTKIEKELEEKKKYLETSIRQKLDQIVAETAEIENEKIMFHVECTELEKTVKAMCLN

>C_sp54_NDC80_paralog2

MNSTFILNHFSLLRHVKKMFSGGPRDVKSLVTANVKKIYSFLIEFEGSEAPSEKDIKSPRGMNDFISIFESIFHHLNKDYKYPQSARVEEEISQIFKGLQYPFPLKVSFFQPFGSSHGYPHLLDALAWLVDLVRMNSCTNDTNEKEMLKMQMTKIEKELEEKKKNLETSIRQKLDQIVAETAEIENAKMMLLEMKCTELEKTVKAMCLN

>C_inopinata_NDC80

MYGDRRRTGGISLNGRASLAITPTKRYTDFRTETRPSLGQSSGQHRLSIFHKTPSVAPRDVKSIFTANVSKVHNFLAEHEGLEAPLESTIRNPRGRNDFICCFELIYQHLSKDYEFRKDERIEDEISQIFKGLGYPFALKNSYFQPMGSSHGYPHLLDALAWLVDIIRVNAAVSADTQNIIFGYFMEQSKAQERTLHYAWMSSIFREYTNERKVAENSESPFWGEVKNKLRKFFEDSGEFEDMTNSAASALEMLRFECDEIESEKDNEQSILEEITRIKDDIRKASAYLETTQQVLQHKEIELAGVQSEVADKIAENEKVQKMVNDLKTRIENQKNHHGLTGKEVRELNLENNRDKEMVSEIQAELNNLSKETWRLKDEDHFKEQRSKFVQLFDSVTKMIAGLNIQMNLKQISSPNNERELKVCWETLDNVWLPEINRQLHQRKLDLETEQSRFSSKFVAAAERIQMKSEILSEVKKNEIREERLRQNERDEWKENRKQKEKRYDELENEKEVLKTQMQMDGSLEKEIEAERISMAKIEKELEEKCKYLQSVIKKKLDQIVTETTAIESQKTIFHLQCTEIEKAVN

>C_elegans_NDC80

MFGDRRKTGGLNLNGRASIAITPTKRFTDYTGSTSVRKTDARPSLSQPRVSLFNTKNSSVAPRDVKSLVSLNGSKIYNFLVEYESSDAPSEQLIMKPRGKNDFIACFELIYQHLSKDYEFPRHERIEEEVSQIFKGLGYPYPLKNSYYQPMGSSHGYPHLLDALSWLIDIIRINSAVSEDTQNILFGDFMEQGKAQEKTLNYAWMTSTFRDYTNDRKAAENPSSSYWDDTKHRLRKYFEQSNEFEDMTKTAASALEMLNYECDEIEADKGNEASLKEEISRIRDDIRKAKDYLEQNLHVKQHMEKELAMVKSEQEEKISENEKVQKMVDDLKNKIELQKQIHGLTGKEVRQMNLDNNKDKEVVLEIQSELDRLSKETWKLKDEDFFKEQKSKFIHLAEQIMKILSGLNIQMNLEPLRAPTNERDLKDYWETLNKIWVPEISRQLHQRKLELETEQSRFSNKAVTAEERIQIQSETLCEAKKNEAREERIRRNERDSWKDARKHIEQRYEQLLNEKEVLLKQMKLDGSLEKEIEDETARMSATGEEHIQKRSQLEAGIRQILDLMVVEIAEIENKKIGFHVQCAGIEKAVL

>C_oiwi_NDC80

MFGDRRKTGGLNLNGPRASLAITPTKRFTDLGVNSVRKSTTRMSLSQTSGQPRMSLFQSKSIAPRDVKALVSSNVTKIYNFLVEHEATEAPSEKDIRSPRGKNDFVLFFELIYGHLSKDYEFPQDTRLEDEIPLIFKGLGYPYPLKNSYFQPMGSSHGYPHLLDALSWLIDLAKISSSVRSDTKNIIFGDFMDQNKALEKILSYSWLSTVYKDFTNDRKAAETDQFWVETKGTLRKHFEHNNEYDDMIETAKSALEQLRFECDEIESDKGNEQSLQEEIARIRDDIRKASDYLEQTEMLKERTQEELNEINTQLQVKSAENEKVQGMVNELKARLEQQKLHQGLTGKEVRQMNLENNKDKETVAELQAQLEELSKQAWKLKNDESFKEQKTKFVHLVENALKVLSGIGINTNSDAFRVPTNEKELQASWETMNNVWLPEINRQLHQRKLELETEKLQFADKFASAEERIQMETEMLREAKKKEVREDRLRRTEREEWKESRQQLEKRLDELENEKEVLKKQLQIDGSLEKEIETEKVRMAKIEKEMEAQKERLEAAIRKKMDEIVEETAGIENSKIMFHKSCVEVEKTVKTALKVSKK

>C_kamaaina_NDC80

MFGDRRKTGGLNLNGPRASLAITPTKRFTDLGVNSMRKSTARVSLSQTSGQPRMSLFQSKSTAPRDVKALVSSNVTKIYNFLVEHEATEAPSEKDIRSPRGKNDFVVFFELIYGHLSKDYEFPQDTRLEDEVKKIFKGLGYPYPLKNSYFQPMGSSHGYPHLLDALSWLIDLAKISSSVRSDTKNIIFGDFMDQNKALEKILSYSWLSTVYKDFTNDRKAAETDQFWVETKETLRKHFEHNNEYDDMIETAKSALEQLRFECDEIESEKGNEQSLQEEIARIRDDIRKASDYLEQTEMLKERTQEELNEIKTQLDTKSVESEKMQGMVNELKARIEQQKLHQGLTGKEVRQMNLENNKDKETVAELQAQLEELSKQAWKLKNDESFKEQKTKFVHLVENALKVLSGIGINTNSDAFKVPANEKELQASWEAINNVWLPEINRQLHQRKLELETEKLQFADKFASAEERIQMETEMLREAKKKEMREDRLRRTEREEWKESRQQLEKRLDELENEKEVLKKQLQIDGSLEKEIETERVRMAKIEKEMEAQKQRLEAAIRKKMDEIVEETSGIENSKMMFHKSCVEVEKTVKALKVSNNQ

>C_waitukubuli_NDC80

MFGDRRKTGGPGFGPRLSTAITPTKRFTDVGPTSVRKADSRLSMGQGGGQPRASLFQRSSAVASRDVKSLMAANVNKIYNFFIESEGSEAPSEAQIRNPKGRNDFITFFELLYQHLSCDYEFPQGARVEDEVSALFKNLGYPTALKNSFFQPMGSSHGYPHLIDALAWLVDLIRVNMCVSNDTQNIIFGDFLEQSKVQERTLSYCWMSSTYKEYTNDRKAAETRDGPFWDSTKIKLRKYFEDMNETGDMAASAKTSLEQLRFECEEIEADKGTEQNLIEEIAKIKDDFRKASDYCERLEAVERQREVDLDQVKMEVDVKKKEMSEVQEEVNSLKQRIEEQKQNHGLTGKQVRQLNSDNSRDKAIVSEIQSELERLSKELWLRKNEENFKEKKTMFLQLAEKLAKIGAGINVDLGLTSLRSPQNEELKIECEKLVGVWLPEVTRHLNQKKLELEMEKSRFNDKFASAACANMEQETLCEAQKKAAREERIRRNEREEWTESRLAKEKRYDELENELDVLKKNMQMGDSLVKEIEMERAKKVKLCEEQEKQKKTIEASIRQKMDQIMTETAEIDNEKMMIHVECNELEKMV

>C_panamensis_NDC80

MFGDRRKTGGPGFGPRLSTAITPTKRFTDVGAAASVRKVDNRLSMGGQHRASLFQRSSAIAPRDVKSLMSANVSKIYNFFLEFEGPDAPSESQIKNPKGRNDFISFFELLYQHLSCDYEFPANVRVEDEVQISTIFKNLAYPTPLKNSFFQPMGSSHGYPHLIDALAWLVDLIRNKCVSDDTQNIIFGDFLEPAKVQEKTLSYAWMSSTYREYTNDRKAAETKDGPFWEVTKNKLRKYFEDMSETQDLAASVKNSLEQIRFECEEIEADKGTEQSLIEEIAKIKDDLRKATDYAEKLEAVERQRLMDLDKVKADLDVKKKENSEVQTEVDGLKQRIEEQKQNHGLTGKEVRQLNSDNARDKAVSNELQSEMEKVAKDLWRNNNEENFKEQKRTSFSRLVERIEKILAGINVTLRLEPLRTPQTERDLKAGLDELMSVWLPEITRQLNQKKLELNMEESRFTDKFVAVQARVQLEKETLDEAKKKEAREERMRRNEREEWMESRLATEKRYDEMENEQDVLKKQLQMDGSLEREIEMEKARKVKISEEQENKKNKIDARIRQKMDEVMKETAEIDNGNLLFHVACNELEKMVLKTQNLI

>C_nouraguensis_NDC80

MFGERRKTGGQGFGPRISTAITPTKRFTDVGAAASVRRTDSRLSMGQSGGQHRASLFQRNSAAAPRDVKAMLGPNVNKIYNFFIEFEGSDAPAEAQIKCPKGRNDFINFFELLYQHLSCDYEFPPNARVEDEVLWIQIFKCIGYPVSLKNSYFQPMGSSHGYPHLVDALAWLVDFIRNKCVSDDTQNIIFGDFLEPAKVQEKTLSYAWMSSIFREYTNDRKAAETRDGPFWTITKERLRDYFEKMNEAQGLAVSLKNALEQIRFECEEIEADKGQGQTMLEEINKMKDDMRKAVDYNEALQTAQRQREIDLEKAMTDLEVRMKEDSEVQADVNELKKRIEEQKEKHGLTGKEVRQLNLENTRDKAVISEIQAELDKLSKELWLKKNEENFKDKRASFVRLAENIGKIVAEVYIDLGLEPLRSPQNEKELKSELEKMNSVWLPEVTRQLRQKQLNLNKEKSFFRDKERVQIERDTLCEAEKKMAREERIRRTERDEWKEIRLANEKRYDELENEQDVLKKQMQKDGSLDKEIELEMARKLKIDEESSKKKSALEAVIRRKLDHIMAETSKIDNEKLMFHTDCTEFEKQILKTRNMK

>C_becei_NDC80

MFGERRKTGGPGFGPRISTAITPTKRFTDVGAAASVRKTDSRLSMGQPGGQHRASLFQRNSAAAPRDVKSMLGANVSKIYNFFIEFEGSDAPAEAQIRNPKGRNDFINFFELLYQHLSCDYEFPPNARVEDEVIHIFKCIGYPVSLKNSYFQPMGSSHGYPHLVDALAWLVDFIRNKCVSDDTQNIIFGDFLEPAKVQEKTLSYAWMSSIFREYTNDRKAAETRDGPFWTVTKEKLKDFFEKMNEAQGLAASLKNALEQIRFECEEIEADEEINKMKDDMRKAVDYNDALQTAQRQREIDLEKAMTDLEIRMKENSEVQADVNELKQQIEEQKEKHGLTGKEVRQLNLENTRDKAVISEIQAELDKLSKELWLKKNEENFKDKRAIFVQLAEKIGKIVAEVHIDLGLESLRSPQNERELKSEQEKMNSVWLPEVTRQLRQKQLNLNKEKSFFRDKFAAAEERVQIKKVTLCEDEKEMAREERLRRNEREEWKEIRLVNEKRYDELENEQDVLKKQMQRDGSLDKEIELEMARKLKIDEESSKKKSALEAKIRQKLDQILVETSKINNEKLMFHSDCQEFEKQILKTRNMK

>C_yunquensis_NDC80

MFGDRRKTGGPGFGQRNSTAITPTKRFTDVGAASSVRKTDNRLSMGQTGGQHRASLFQRNSAAAPRDVKSMMGANVSKIYNFFLEFEGPADAPSEGNIRSPKGRNDFINFFELLYQHLSKDYEFPANARVEDEIIHIFKFLGYPISLKNSYFQPMGSSHGYPHLVDALAWLVDFIRCSISVSGDTQTIYYGDFLDQAKVQERTLSYAWMTTIFREYTNDRKAAETRDGPFWTMTKEKLRTYFEKMNEDQDLTASVKNALDQIRFECEEIEADKGKGQEISEEIVKVKDDIVKAIDYGEALEAAQKQQESDLEKATAELRERVDENSEVQANVNELKQRIEDQKKKHGLTGKEVRQLNSENTRDKAMITEIQAELDKLSKELWLKKNEENFKDKRESFVQLVERIGKIVAEVSIDLGLEPLRSPRNERELKTELEKLNGIWLPEVTRQLRQKQLNLDKEKAFFRDKFAAAEERVQIERDTLCEAEKKMAREERVRRHERDEWKESRLVIEKRYDELENEQDVLKKQMQMDGSLDKEIELEKAKKVKIDEEIAKKKAALEAAMRQKLDKIMVETAKIDNEKILFHVECTEFEKQILKTRNLN

>C_macrosperma_NDC80

MFGDRRKTGGPGFGPRISTAITPTKRFTDVGAAASARKVDSRLSMGQPGGQHRASLFQRSSAVAPRDVKSMLAANVNKIYNFLVEYEGTDAPSEAQIRCPKGKNDFINFFELLYQHLSCDYEFPANGRLEEEISNTFRHLGYLPHLKNSYFQPMGSSHGYPHLVDALAWMVDIIRCNKCVSDDTQNIIFGDFLEPAKVQEKTLSYAWMSSTFREYTNDRKAAEAINGPFWNSTKNKLRKFFEDMNDTQDLTASAKNALEQLRFECEEIEADKGTEQSLLEEITRIKDDIRKAICHYEDTKRVLNQRESDLEKVKAELDTRIKENNEVQAEVNLLKNRIEEQKEKHGLTGKEVRQLNSNNTRDKAVVSEIQTELDKLSKELWLMKSEDTFKEKKTCFVHLTERIGKIVAGIEIDLGLEPLRSPQNERELKVEWEKLNNVWLPEITRQLHQKKLELEMEKSRFSDKFAAIEERVQMERETLCEAKKKESREERLRRNEREEWKESRLLKEKRYDELENELDVLKRQMQMDGSLDREIEIEKAKKVKNDEELKKKKAAIEASIRQKLDQIVAETAKIENEKTLFHVECIEFEKLILNTQNLV

>C_sulstoni_NDC80

MYGGDRRKTGGFGFGNPRTSAITPTKRFTDLGTTSSVRKDHGRLSMSQTSGHHRVSLFGGASATGPKDVRANFDANAKKIYNFLVEHEGSEAPSENVIRCPAGKSDFVTVFELMYQHLSKDYEFPQNVRVEEEVSNIFKALGYAPPLKNSYFQPMGSAMGYPHLVEALAWLVDLIRINASVSADTQPIIFGDVMDQEKVHEKTLSYAWMSTTFRDYTNDRKAAESGTGPFWEETKSKLRDYFEKSDEIEDVATNYKSALEQLRFECEEIEAERGNELTLLEEIAKIKDDVRKALEYDETTARVQKHKEDELKAVKGTLEQKMAENSKVQAEVAELKERIEVQKQMHGLTGKEVRQLNSDNNRDKETVTELQAELDEISKQMWRLKNEDTFKQQKTNFVRLVESVEKIVSGLDIEMGLEPLRAPSNESELKSRWDELNSVWIPEINRQLHQKKLEFETEQSRFADRFAAAERVQMERELLCEAKKKEAREDRVRRNERDEWKENRLQREKKYDELENEKDVLTKKMQMDGTLDKEMEEERTKAWKLEQELQKEADAVENAIRKNLDLIVNEVAQIENDKMLFHVECCEFEKLIVKTQNMF

>C_afra_NDC80

MYGGDRRKTGGFGFGNPRTSAITPTKRFTDLGTTSSVRRVSLFGGGSAAGPKDVKGNFEANVKKIHSFLIEHEGNEAPADTVIRSPAGRNDFLTVFELMYQHLSNNYEFQQGVRIEDEVSNLINASVGADTQTILFGDVMDQAKVQEMTLRYSWMSTTFRDYTIDRRAAESGTGPFWDETKNKLREYFEKSDEIEDLVTSYKTALEQSAFECQEIEADKGNEQHLLEEISKMKDDVRKAMEYAESTARVQKHKEEEMKTVKATLETKIAENSKVQAEVAELKERIEVQKQLHGLTGKEVRQLNSDNNRDRETVTELQAELDEVSRQMWRLKSEDTFKQQKANFIRLVESVEKIVSGLELQLGLDAILAPSDESELKVRWDELNNVWIPEVNRRLHQKKLEFETEQSIFVDKFAAAERVQMERELLCEAKKREAREDRVRRNERDEWKENRLQREKKYDELENEKYVLKKKMQMDGSLDKEVEEEQAKARKIEEELQKEADAIENSIRKNLDQIVSETAQIENDKMLFHVECCEFEKLI

>C_sp49_NDC80

MMFGGPRRKTGGPNFGATGRTSTAITPTKRTTDLGAAHSVRKNDNRMSFGQGPSNSRPSLFQRTGGPSKDFKAAVPGNAAKVYNFFVDTEGANAPSEKTIRTPAGKGDFVTMFELLYQHLSKDYEFPSNGVRLEDEFTNIMKALGYPHALKPSFFQPIGSSHGYPHLLDALAWLVEAVEINEKVRLDTQNIIMGDFMEPDQVQDKVLSYSWLSKTFFEYSNNRKVLEDKADPFWATTKCDLRNFFENNDESADMITSSKNMLEQLRFDCEEIEADKGNEQGLLDEIARIRDDIRKAQLYLDETMRVVAQSNHEYQTVVDACEAKLAELEKVKTEVAELKERIEEQKRLHGLSGKEVRQMNAENSKDKEMLSDVNAEIERATKMMWRLRDAADYKEPSTRFYRLISHMHQILDDADVHPQLEPIASPSSELDLKAGWDAINNLWLPEVNRQLQHKKLELETENARFLNKFEAIEERATMEQEMLNEATKKEDREERVRRNEREEWKAARFQKEKRCDQLENEKDILVKQMHLDGSLDRDLQEETEKAAKLSETLEQKQAAFEQKVREKIDEMMASTAEMEQEKLEFHKECVELEMVVKNS

>C_sp25_NDC80

MFGGPRRKTGGPGFSTTGRTSTAITPTKRNTDLGTAQSVRKNDGRMSFGANPSGSRPSLFQRTVAPSKDVKAAVPANVARIYNFFVETEGPNAPSEKMIRSPAGKGDLITMFELIYQHISKDYEFPSVGVRLEDEFTNIMKALGYPNALKQSFFQPIGSSHGYPHLLDALAWLVEAVEINEKIRLVTQNILVGDFMEPDQVQDKVLSYSWFSKTFLEYTNNRKVLEDRSDPFWANTKRELRDFFERNDESADMIATSKNMLEQLMYDCEEIEADKGNEQGLIDEIARIRDDVRKAQAYLEETSRVVAQSIHEFQSVSDAFEAKRTEFEAVKTEIAELKERIEEQKRNHGLCGKEVRRLNLENIKDKETVAELQAELEKTSKIIWNLRDAGSFKEQKARFDRLITHMKMILDDTDIHFQLESITSPRTELELKAGWDTVNNVWLPEVNRQLQRKLLELETENARFASKFGAKEEHVTMEQEMLNEATKKEDREERVRRNEREEWKAARFQKEKRFDQLENEKDILMKQLHLDGSLDRELEEETEKSIKLTEALEQKKAEFEQKVREKIDLMMVSTAEMEQERLDVHKEFVELETIINSS

>C_imperialis_NDC80_paralog1

MFGGPRRKTGGPNFSATGRTSTAITPTKRNTDLGAVQSVRKTDTRMSFGQGPSAPRASLFQRAGLPSRDVKACFASNVSRIYNFFVETEGSSAPSEKTIRSPNGKGDFISMFELIYQHLSKDYEFPTQNRLEEEFSNILKSLGYPYPLKNSFFQPIGSSHGYPHLVDALAWLVEVCEVNEKVRLATQNILLGDFMEPEQMKDKFISYSWFSKVFLEFTNQRKAAEDKTDPFWESTRRELREFFEQNDESVEMIASLKNMLEQLRFDCEEVEEDKGNEQSLLDEISRLRDDVRKAQSYLEETARVVAQSHHEFATVKEAVDAKMEEFERVKAEVAELKARIEQQKELHGLTGKEVRLMNVENNKDKETVAEIQNELDRTSKIIWRLRDAGSFKEQKSRFERLVEHMLKIVDDADIHLTLDPINSPSTELELRAGWEAVNTVWLPEVNRLLQRKKLELEGENARFSSKFGAVEEKAILEKELLNEATKQEAREERVRRNERDEWKESRLQKERRCDQLENEKDVLVKQLHLDGSLDVEIREKADEMEKLVVELAGKKKTLEGEVRKRIDEMMESTVQMEHEKLLFHRECVELEALVKSQ

>C_imperialis_NDC80_paralog2

MFGNARRKTGGPNFATGRPSTAITPTKRITDLGAAQSVRKNDSRMSFGPSAHRTSLFQRAGPHSKDMKACFAANVAKIYHFFVETEGSEAPPEKMIRSPTKNEFITIFELVYQHLSKDYEFHMQNRLEEEFTSILKALGYPYPLKNSFFQPIGSSHGYPNLVDALAWLVEVVEVNSAVSRVTQNILIGDFMEPELAEDKVLSYSWFSKTFLEFTNNRKALEDKSDPFWESTRLGLREYFERNDETAEMIASCKNTLEQMRFDCEEIEEDKGNEQSLLDEISRLRDDVRKAEAYLLETSRALEQATQEHESVKEAVETKGAEFEQVKMEVARVKELIEQQKEQHGLTGKEVRAMNVECQRDKETVSEIQMELDKVSKVIWRLRDAGSFKEQKSRFDRLIEHMHTILVDADIHLELEPICSPSTEIELRAGWEAINGVWLPEVNRQLQHKKLELEAENARFSSKFGVVEEKAILEQELLNEAIKQEARDERVRRNEREEWKASRLQKEQRCDELENEQDVLVKQLNLDGNLDAEIREAAQKAERLFLELQAKKEQFEGAVRAKIDEMMESTVAMEQEKLMLHKECSELESSVKSTL

>C_japonica_NDC80

MFGERRKTGGANFGFAGGRPSTAITPTKRFTDLGTAQSTRRVDNRLSMGQNGGRLSVFQRGSAVAPRDVKSAQASNVTKIYNFLLEHEQSGAPPEKIIRQPTGKNDFITMFEQLYQHLSKDYEFPQGARVEDEFTGIMKGLGYPFPLKNSFFQPMGSSHGYPHLLDALAWLIDVISLNQSVSSVTQNILLGDFMEAAEAQDKIISYSFYSSTFREYTYDRKAIESKDAPFWAETKERLRDYFEKNDEITDLVTSTKQVFEQLRFECEEIEADKGDEQGLLEEIARIKDDVRKAHEYLESTERVQKQTGVELGIVQESLEAKKAEFEAVQAEVNELKRRIEVQRQQHGLTGKEVRQLNVENNRDKEAVHEIQTELDNVSKTLWRMRDEDTFREQKANFVLVIENMEKILLDANVKIGLDALRPPQNERDLKVGWDALNNQWLPEVNRQLQHKKLELDDEKTQFSSRFAAIEERVQMQQELLREANKKEAREERVRRNERDEWKTDRLQREKRLDELENEKDVLKNQMQTGGSLDREIEEEKAKSAKLVEALEQKKASLAEGIRQKLDETMREITKIDLENATFHAECTDIEKLVLKTRNMNQ

>C_tribulationis_HIM10

MNNAKQVILVNHDPKSIITKLNSKLHLGLTPENVTHPAETALTIFMSFVRYVLNVSDHSLTTLPLSASGEHDPELHKRSVQLTLVYQSMKAFITDNSGQKLDMRMCDLVTPAKEPLRFRKLLAFLVEFIKLHEISAPIFNEISDEFSEQKHAMESLQEDIATAERKKNELLSKQSLRKRRENELMDDHSKIKNELNGIVNQYNDNLAMTNDMEKQKAELIQQIEDIEREILTAKKTVEHLTEEVLESPEELKKEMKERKRRIEEFREALAASRQTLKSKVEAREICANSEKNLPVIQQRIQSWKEVREDILDLMDEIQEKLRKLDEMQEQLAFTTDKKNNSGKRMIEQAEMHEQLRKEHLQRSENLNKNIEEIRQIASLGKNQPDVSRDIEKKRQELLAAKNAHSELMSRIMNSTKDAFVKFRKIDAHFKATQNVALEKHCAMDRAKNRLSNSYKSRLPSDYTFSASSINESDAENLDPQSPVFENFSVFKN

>C_sp41_HIM10

MNNQKQVIVMNYDARSIITKLNPKLHLGLSPDNITNPAETSLTIFMSFVRYVLNVSDHSLTTLPLSASGEHDPELHKRSVQLTLVYQCMKAFIKDNSGQKMDMRMCDLVTPGKEPARFKKLLCFLVDFIKLHEMAAPIFNEISDEFSEQKHAMEALQEDIAIAERKKNELLSKQNLRKRRENELMDDHSKIKNELNGVVNQYNDNLVITNEMEKQKVELIQQIEDIEREIMTAKKTVEHLTEEVLESPEELKREMKERKKQIEEFRDSLAASRQTLKTKLEAREICANSEKNLPVIQQRIQSWKEVREDILDLMDEIEEKLRKLNEMQEQLAFTTDKKNSSGKRMIERAEMHEQLKREHLQRSEDLNKNIEEILGPHSERISQMCREILRSRDKNSSPPKTPTANECLAS*TRRRTRSPNFEKSTLASRRPNAWQSRKGVQSIVPRIASATRSNPAFPATTRSVLPVSTTRNRRILTHSRRRSRTFLYLTI

>C_zanzibari_HIM10

MNNGKQVILFNHDAKTIVTKLNAKLHLGLSTDNIINPTAETALTVFMSFVRYILNVADQSLTTLPLSASSEHDPEFHKRSVQLTIVYQSMKAFITDNSGQKLDMRMCDLVTPAKDATRFKRLLSFLVEFIKLHEMASPIFNEISEDFSEQKNMVDRLQEDIADAERRKNELLSRQSLRKRRENELMDDHSKIKNELNGVVNQYTENLAITEDIEKQKVELFQQIEDIEREIMTAKKTADHLTEEVLESPEELKNEMRERKKQIEEFRESLAASRNTLRLKLEARDICANSEKNLPVIEQRIEAWKEVREDILDLMDEIEEKFRKLNEMQEQLAFTADKKETSGKRMIEQAEMHEQLKKEHLQRSEKLNKNIEEIVQIASLGKNQPDVSRDIEKKQRQELLAAKNAHSERMSRIMNSTKDAFAKFRKIDAHFKETQRVAMEKQCAMDRAKNRLANSFKSRLPSDYTFSASTINEQDSENFDPESPVFENFSVFKN

>C_sinica_HIM10

MNNGKQVILVPNDVRTIVAVLNKKLRLGLSQDSINIPTETAFLVYMNFVRVVLDVNDIHLATLPMSANTDHDPELHRKSVPLIIVFQCMKAFISDNSGQKLDMRMCDFVIPGRDIPRFKKLMSFLVEFIKLHEMAAPVFNEISGEFSDQKQEIEALQREINDTEKKKTELLSRQSLRKRRENELMDDHSKIKNELNGIVCQYSDNLAATEEMEKQKAELIQQIEDINREILTTKKTAEHLSEEVLESPEELKREMTERKKQIEELRECLAASRQTLKAKLDARDICANSEKNVPVMQQRLETWNTVREDILTLMDVIEAELRKLAEMEEQLAFTTDKKNNSGKRMIEQAEMHEQLKREHLQRNEKLSKNIEEIQIAALGKNQPEVSRDIEKKRQELLAAKNAHSERMSQIVRSTKEACGKFQKIDAHFKDLHRVAQEKRCAMDRAKNRLCNSFKSRLPSDYTFSASSINEEASENCDPQSPVFESFSVFKN

>C_nigoni_HIM10

MSNAKSVILIQYDTRLIAKVLQQKLGLGLTADSILNPDVSSFPSIAEVAQSIFSNFIRYILNVSEQSLLTLPLSADSGHDPELHKKTIPLVIIYQCMKAFIVDNSDRKLELTMCDLVTPEKQPNRFRRLTSFLVDFIKLHEMSAPIFNEISDEFSEQKQEMERMQDEIIQAEKRKNDLISKQSLRKRRENELMNEHSKIKSELAGVVSQYTDVLERTEEIEKQEKELIQQIEEIEREIMTAKKTVEHLTEEVLASPEELKQEMANRKKQIEELKESLAASRQTLKSKLEARNICANAEKNVPLINQRIQAWSEVREDILDMMDEVEEKLRKLNEIQEQLAFAADKKTSSEKRMIEQTEMHEQLRREHLQRSAALGTSQPDVSRDIENKQRQELLAVKNAHSEKMALINNAQKDALAKHRRIDDHFKETLRVAVEKRNAMERIKNRVSNAYTGRLPSDYTFSASSINESENCDPQSPVFENFSVFNN

>C_briggsae_HIM10

MSNAKSVILIQYDTRLIAKVLQQKLGLGLTADSILNPDAEVAQSIFSNFIRYILNVSEQSLLTLPLSADSGHDPELHKKTIPLVIIYQCRLTSFLVDFIKLHEMSAPMFNEISDEFSEQKQEMERMQDEIIQAEKRKNDLISKQSLRKRRENELMNEHSKCKSELAGVVNQYTDVVERTEDIEKQKTELIQQIEEIEREIMTAKKTVEHLTEEVLASPEELKQEMANRKKQIEELKESLVVSRQTLKSKLEARNICANAEKNVPLINQRIQAWSEVREDILDMMDEVEEKLRKLNEIQEQLAFAADKTTSSEKRMIEQTEMHEQLRKEHLQRSEKLYKNIDEIARQIAALGTSQPDVSRDIENKRQELLAAKNAHSEKMALINNAQKDALAKHRKIDDCFKETQRVAVEKRNAMERIKNRVSNSYTGRLPSDYTFSASSINESENCDPQSPLSNLGPRSLKHVQSAASSSSSVVLLHKSYTESSAPPSTNRKLPYKTLRLASTQQVTEFIQNGELPTFKKQGSRSTNNPWRRHGALQKLCETGGFAPNSGSKKERKNRGSLLAVKQNVKVPPAPTSSQMTLNKVKEQGQEIKLALKAPKNVAVDVAAQNLAEQVKAFVLRYGSELVEDERTAQSK

>C_remanei_HIM10

MSNAKSVVLVMFDPRKISTTLNQKLQVGVTPDNILNPTAEIVQQIYLNFVRVVINISENSLHTLPLNADSDFDQELHKKSIPLAIVYQSMKAFIKDNSGGKLDLTMCDLVTPGKNPQRFRKLSSFLADFIKLDEIAAPIFNEISEEFSDQKVEMEALQEEIVAAEKRKDELVARQSQRRRRENELMDDHNKKKSELAGIINQYTEIGVKTEELEKQKNELIRQIEETEKESITAKKTVELLNEEVLASPEELRQEMTERKKQIEDLKESIITAKQALQDKLEARDICANADKNVPVIEQKIQAWAEEREDILDLMDEVDENLRKLSEMEEQLTFTTDKKSNHGKRMIEQAEMHEQLRREHLQRSEELNKNIEEITGQIAALGKNQPSVSRDIEEKRQELLALKNAYSEQLAKYRNSSRDSFNKFRKINALFNEVQRVSLEKKNAMDRAKNRLQNMLIGRLPSDYTFSTSSINDSENCDPISPIESDFSVFKN

>C_latens_HIM10

MSNAKSVVLVMFDPRKISTTLNQKLQVGVTPDNILNPTAEIVQQIYLNFVRVVINISENSLHTLPLNADSDFDQELHKKSIPLAIVYQSMKAFIKDNSGGKLDLTMCDLVTPGKNPQRFRKLSSFLADFIKLDEIAAPIFNEISEEFSGQKLEMEALQEEIVAAEKRKDELVARQSQRRRRENELMDDHNKKKSELAGIINQYTEIGVKTEELEKQRNELIRQIEEIEKESITAKKTVELLHEEVLASPEELRQEMAERKKQIVDLKESIITAKRQALQDKLEARDICANAEKNVPVIEQKIQAWAEEREDILDLMDEVDENLRKLSEMEEQLTFTTDKKSNHGKRMIEQAEMHEQLRREHMQRSEELNKNIDEITGQIAALGKNQPRVSRDIEEKRQELLALKNAYSEQLAKYRNSSRDSFNKFRKINAHFNEVQRVALEKKNAMDRAKNRLQNMLIGRLPSEYTFSTSSINDSENCDPISPIDSDFSVFKN

>C_sp51_HIM10

MAHQKQVMLCTYDARIIAKSLSQKLQLGLTADNIINPTAENAQQIFSQFARIILNVSEHSLTTLPLSADVDHDHELHRKSVPLTIVYQSMKAFIDDNSGGKLELTMCDLTTPAKNEIRFRKLTSFLHDFIKLHEVASPVFNEICDEFSDRKLDMERIQEEVNIAEKKKEELLAKQASRKRRENELMNDHNKLKTELNNVVNQYMKNSEITSDIDKQTEEAFRQIESVERETVTGKKTVEHLTEEVLTSPEELKHEMIQRKKHIEELKECLKSSKQSLQVQLEARDICINSEKSVPVVVEKIRVWSEVRDDILDLIDSVEENLRKLNEKQEHLAFTADKRTKIAERVIEQAQMHDQLRKEHLQRTEELQANIEKIAALGKNQPDVSKEIAQKQREELLSVKNAFSETIAKINNSCRDAISKFKKIDVQFRETQRTSAEKRNAVQRAKDRLRTACIGRLPSDYTFSTSSINDSENCDPLSPIRLSNFNVFK

>C_sp44_HIM10_paralog1

MLHQKQVVLIGHDHRIIARSLSQRLQLGLTPDNILNPTAEVSQQIFTHFARLILNVSEHSLTTLPLSADIDHDHEMHRKSIPLIIVYQSMKAFIDDNSGGKLELTMCDLVTPGKNPQKFKRLTSFLHDFIKFHEVATPIFNEISDEFSDRKLEMDQLQEELRDAEKKKDELLGKQASRKRRENELMKDHNTIKTELSSVVDKYMKNSELTNNIDKQSEEAIRQIEEVERETLTGKKTVEHLTEEVLNSPEELKQEMEQRRAHIEELKECLKASQNLQAKLEAREICQNSEKNVPVVLEKIGVWIEVRDEILDLIDSVEKNHRNLNEKNEQLSFTSNKKTNTNERMIEQAEMHQQLRKEHLQRMKELQANIEDIQRQIANLGINQPDVSKENAEKREALISVKNAHSETVSKIFSSCQEAVSKYEKIVARFKETQNKSMEKKVAFDRAKDRLRAACVGRLPSDYTFSTSTLNDTENYDPLSPIAPSDVNVFK

>C_sp44_HIM10_paralog2

MQHQKQVVLKDYDVRTLARSLGQRLQLGLTPEDFINPTAECSQQIFTNFARLILNISEHSLSTLPLSASEIDINHEMHRKSIPLIIVFQSMKAFVHDNSGGKLDLSMCDLVTPGRNPQRFKKLTSLLYDFIKLHEAAAPIFDEIAEEFSDRKIEMDQLQEELRAAEKKKDELLGKQASRKRRENELMNHHNKFKTELSSVVDQYTKNAELSNNNDKQSEEAFRQIEEVEREIITGKKTIEHLTEEILDSPEELKQEMEQRRAHIEELKECLKASRQNLQDKLEARDICINSEKSGPVVHEKLGAWKAVREEILALIELIEQNQRDLNDEYEKLTFIANKKTSVNERMVEQAEMYEQLRKEHLQRMHNLQANIEDIQRQIASLGINQPEVSKENAAKREELISAKNQHSETIAKILSSCKEATAKYEKIRADYIQTQRKAIEQRVAAARAKDRLRAACVGPLPSDYTFSTSTLTETENTEPLSPIAPSDFNVFK

>C_sp48_HIM10

MQHQKQVVLVNYNPRDIAKSLSQKLQLGLTGESITNPTGEVSQQIFSQFARIILNVSENSLQQLPLTAGCDHDHELHRKSIPLIIVYQCMKAFIEDNSGGKLSFSMCDLVNPQRDATKFKRLTSFLHDFIRLHEFAAPIFNEICDEFSDRKQEMELIQEELRAAEKRKDDLVAKQASRKRRENELMNDHNKLKTELNNVVNQYMKNTEMSNEIDKQAEEALRQVEEVERETITGKKTIEHLTEEVLSSPEELKQEMAQRKKHIEELKECLKVSRQALQTKQEARDICTNAEKNVPVVAEKIEVWAEVRDDILDLIDSVEENVRKLNEMQEDLALTANKKTKANELMVEQSQMHEQLRNEHLQRTQKLQANIEEITKIAGLGKNQPEVSRENSKKQQELIVVKNAHSQTVARIVNSIQDSVSKFQKLELQFRETQKSALEKRNAVQRANDRLRAACVGRLPSDYTFSTSSLCDSENHDPLSPIAPADFNVFK

>C_brenneri_HIM10

MQHQKQVILISYDQRTIARSLSLKLQLGLTGESITSPTAETSQQIFTQFARIILNVPEHSLTTLPMSAGADHDNDLHRKSIPLIIVYQSMKAFIEDNSGGKLSLSMCDLVNPSKDPQKFKRLTSFLHDFIRLHEYASPIFNEICEEFSDQKQEMELIKEELAAAEKRKNDLVAKQASRKRRENELMKNHNELKTELNNVVNQYMKNSELSNEIDKQTEEACRQVEEVERETITGKKTIEYLTEEVLSSPEELKQEMAQRKKHIEELKECLKISSRQALQLKQEARDICINAEKNVPVVTEKIEVWAEVRDDILDLMDSVEENVRKLNEMQEDLALTANKKAKANEQMVEQSQMHEQLRKEHLQRTQELQANIEEITRKIAGLGKNQPEVSRENSKKQQELIAVKNAHSETVAKIINSIQDSVSKFQKLEQQFRETQKSALEKRNAVQRANDRLRAACVGRLPSDYTFSTSSLCDSENHDPLSPIAPADFNVFK

>C_wallacei_HIM10

MSNQPQVVLTLYDARLLAKSLSQKLQLGLTAENFLHPTAEVAQAVLTQFARIILNVPEHSLSTLPLSSNCDFDPELQRKSIPVVLVYLSKAFIKDNSGGKLELTMCDLTMPSKGNNNRFRKLASFLHDFIKLHEFASPIFNEICEEFSDRKLEMEEVQEELIAAEKKKKDLLAKQASRKRRENELMNDHNKLKTELNNIVNQYMKNTELTSDIDKQSEEAIRQIEEIERETLTGKKTVEHLNEEVLSSPEELKQEMDKRKKHIEELKECLKVSRQNLQTKLEARDICATAEKTLPVVVEKLEAWSEVRDDILDLMDAVDGNLRKLNEMNEQLTFTANKKITVGERLVEQSQMQEQLRKEHLQRTEELEANIEEISRIAALGKNQPDVSRDIANKRQELIAVKNAHSETIAKLTNSSHDAVSKFRRIDAQFRETQRVSLEKRNAVQRAKDRFRNACVGRLPSDYTFSTSSINDSENCDPQTPMESDFNVFN

>C_tropicalis_HIM10

MSNQMQVVLTMFDAKIVAKALSQKLQLGLTGENITNPTEVAQNVLSQFARIILNVPEHSLSTLPLSSNCDFDHELQRKGIPVILVYLSMKAFIRDNSGGKLELTMCDLTMPAKTPNRFRKLASFLHDFIRLHEFATPFFSEICEEFSDRKLEMEEVQEELMAAEKKKNDLLAKQASRKRHENELMNDHNKLKTELNNIVQQYTKNTEITKEVDKQTEETMRQIEEVERETLTGKKTVEHLTEEVLTSPEELKQEMDTRKKHIEELKECLKVSKRQSLQSRLQARDICTTAEKNLPVAVEKLQVWSDVRDDILDLIDAVDLNFRKLNELTDDLTINTDKKRNLGERLNEQSQMQEQLRREHMQRTEELQANIEEIKKISSLGSNQPDVSRDILKKKEELIAIKNAHSETVAKLTNSSVDAMSKFSRIDAQFRETQRVSLEKRNAVHIAKSRVRNACIGRLTSDYTFSTSSMIDSENCDPLSPVESDFSVFN

>C_doughertyi_HIM10

MSNQRNAVLTMFDSKNVSKLLNQKLQLGLTPDNITTPTAEIAHQVFSQFARMILNVSEHSLSTLPLSVDSSDHDQEMHRKSIPIVIVYQSMKAFIKDNAGIDLTMCDLTTPAKVPNRFRRIASFLYDFIRLHEFASPIFNEISEEFADQKLEMASIQEELVVAEKRKNDLLSKQALRKRRENELMNDHNKLKTELNNVVNQYMKNSESSNDIDKQTEETTRQIESVEMETITGKKTIEHLNEEVLSSPDELKQEMFERKKHIEELKECLKVSRQNLQAKREARDICIAAGKNVPIVIEKTEVWSEVRDEIVDWMDMVDENRRKLSEMQEQLAFTTDKKAKAEQRIEEQKQVHEQLRQEHLQRSQKLQADIEEITRITALGKNQPEVSRDIAKKREELIAVKNAYSETVAEITNSCQSAISKFHKIDSQFKDTQRKAMEKQNSVHRAKDRFRNSFVGQLPSEYTFSTSSINDSENFDPQSPMESDFNVFN

>C_sp54_HIM10_paralog1

MANARPVVLIMYDARLIAKQLSQKLQVALTAESILTPTAEIAQQIYYNFVRLFLNVSEHSLTTLPLSANSDHDQELHRKSISLVIVYQSMKAFIKDNSGEKLDLTMCDLVTPGKIPQRFRKLTSFLVDFMKLHEMASPIFNEISEEFSDRKLEMEAIQEELLAAERRKNDLLSRQSLRKRREHELINDHNKVKGELNNIVSQFQKNVDDSTELDKQRNEAKQQINAFEKEIDTGKKTVEHLNEEVLASPEELRQEMAERKKQIEELKDCLKSSRENLQTKLEARDICINSEKNLPVINEKIKMWAEVRENIIDLIDIVNENLRKLNEMEEQLVFTTDKKKNTGERMVEQAEMHQQLRKEHLQRIQELQNNIEDITRQITAMGKNQPDVSRGIEKKQRQELLATKNAHSATVANASNACQDALAKFRKVDSLFRDTQRIALEKKTAGDRAMGRLRNSFIGRLPSDYTFSTSSINDSENCDPCSPVDSEFSVFK

>C_sp54_HIM10_paralog2

MANARPVVLIMYDARLIAKQLSQKLQVALTAENILTPTAEIAQQIYYNFVRLFLNVSDHSLTTLPLSADSDHDQELHRKSISLVIVYQSMKAFIKDNSDKKLDLTMCDLVTPGKIPQRFRKLTSFLVDFMKLHQMASPIFNEISEEFSDRKLEMEAIQEELLAAKKRKNDLVSRQSLRKRREHELMNDHNKVKEELSNIVSQYMENKDYSTALDEQKDEAEQQIEAFKKEIIAGKKTVELLNEEILDSPEELRQEMAERKKQIEELKDCLKSSRENLQTMLEARDICINSEKNVPVINEKIKMWTEVRENIIDLIDIVNENRRKLKEIKEQFVFTADKKENTGERMVEQAEMHQQLRKEHLQRIQELQNNIEDITRQITAMGKNQPDISRDIEKKRQELLATKNAHSVTVAKTSNTCQDALAKFRKVDSLFRDTQRIALEKKTAGDRAMGRVRNSFIGPLRNDYTFSTSSINDSDENCDPCSPVDSEFSVFK

>C_inopinata_HIM10

MSTSKTVVLVMYDPRMIAKYLNQKLQVGLTPDDILAPTAEISQLVFTNFVRHVLGVSEQSLNTLPLAVDFGPDQEMQRNSIPIIIVYQCMKAFIIDNSDKKLDLTMCDLVAPAKIPSRFRKLTSFLVDFLKFNDLATPVFNEISDEFSDRKLEMEALQEELVAVEKRKNELISRQNLRKRRKHELINEHNKLKEELNKMVSEYTENQSNTVLLNKKKEDAKQQIEYLEKEVLTGKKTIDHLTEEVLESPEELKQEMSERKKQIEELNDCLKCSRNLKSKLEDLEICVNAEKNVPVVVDKITTWSVLREEILDLIDVENENLRKLKEMEEQLSFTSNKTEAANKRILEQAELHEQLRKQHLERSDEWQKKIEEITRQISAMKFNQPDVSREIEKKRSDLLAAKNAHSEAISRMTNSCKETMSKFRKIEAVFKDTQRTSAEKKTAGDRAFDRLRNACVGCLPSDYTFSTSSISHSENCDPIETEFTVFK

>C_elegans_HIM10

MSNVVLIVYDPRMISKYLGQKLHMGLVADDIIKPTAEIAQQIFANFVRLVLNVSESSLTTLPLSANCDYDPELHKKSIPIIILFQCMKAFIKDNSGNKLDLTMCDLVTPAKHEHRFRKLTSFLVDFLKLHELATPAFNEISEEFSDRKFEMEKIREELLEAEKKKNDLLAKQSIRKRHEHELINEQSNAKAELKNVVNEYTETRQINEELDKQKEEAILHIQALEKEMLTGKKTIEHLNEEVLTSPEQLKQEMEERKRHIEELRDCLESSKKGLQAKLEAREICINSEKNVPVIIEKIHQWTEVREVIIDLIDVESENLRKLKEMEEQLDFMMKEMETAQKRLVEQSETHEQLRIEHTQKSEERQRRIEEITEQIANLKTSQPDVSQEIAKKKQELLALKNAHSETISQITNSCQDAVAKFAKLNAMFKETQKVAFEKNTAAAREMERLKSSLTGRLLSDYTFGSSTIDAGENTENCDPQPNDSSFSVFK

>C_oiwi_HIM10

MTSARPAVLITYDARLVAKFLSQKLQVGLTAENILTPTAEIAQQVMVNFVRLIIGVGEHSLSTLPLSADCDHDPELFRNSIPLVIVYQFMKAFMMDYSGEKLDLTMCDLVTPAKFPQRFRKLTSFLMDFIKLHERAAPLFDEISEEFRDRKIEMESLQEDLINEEKRKNDLISRQNLRRRREHELINDHNKVKGELNSIVNQYTENADISADVDKKKKDAKAQIEDFEREIITGKKQLEYLTEEVLDSPEELRKEMEQRKHQIAELRECLESSRNLQAKMEALDICSNSDKNVPVVNERIRIWSEKREAILDLIDSVEEDHRKLESLEEKLIFKQDEKNNVANQLLKCADSHEELRKEHLKRMEEFNGKIADITQQIATLGKNQPEASRDIEKKKHELISLKNMHSQTVAQLSNTCKDTLAKFRKVDALFKETQRVGLEKKTAGDRAMERLKSACVGRLPSDYTFSTSSINDSENFEPHSPFDPESSVFK

>C_kamaaina_HIM10

MTSARPAVLITYDARLVAKFLSQKLQVGLTAENILTPTAEIAQQVMVNFVRLIIGVGEHSLSTLPLSADCDHDPELFRNSIPLVIVYQFMKAFMMDYSGEKLDLTMCDLVTPAKFPQRFRKLTSFLMDFIKLHERAAPLFDEISEEFRDRKIEMESLQEDLINEEKRKNDLISRQNLRRRREHELINDHNKVKGELNSIVNQYTKNAEISADLDKKKKDAEAQIADFEREIITGKKTFEFLTEEVLDSPEELKKEMEQRKHQIAELRECLESSRTNLQVKMEALDICSNSEKNVPVVNERIRIWSEKREEILDLIDAVEEDHRKLESLEEKLIFKQDEKNNAAKQLINCADSHEELRKEHLKRMEEFNGNISDITRQIATLGKNQPEASRDIEKKKHELISMKNIHSQTVAKLSNTCKDTLAKFHKVDALFKEIQRVGLEKKTAGDRAMERLKSACVGRLPSDYTFSTSSINDSENCEPHSPFDPESSVFK

>C_waitukubuli_HIM10

MATQKPVALVPMDKVSICKILNQKLQIGLTQDTLITPTVEIAQLMYMNFVRSLLNVSESCLSTLPLSATCDHDPELHRRSIPIIIAYQCMKAFIKDHFGDKLDFQMCDLVYPQKTPGRFKRIAGFLADYIRFHEKGLPVFNEVSEEFSYQKQEVELLQEELLEEEKRKNALLAQQNQRKRREHELINEHNKANAELNGKIAQYEASTANAEVLEKEKLEAMDAVDRMENEIISCRKMVDHLKEEVLSSPEELKREMALRKKQIEELKECLNGAKWSLAERLEAIEICTSFEKYKPTVDEKMRQFAAMKEEIIQLFDAVHENQRVLSDLEDEKKFTEEKRKNVIEVLGENAHNHAELREKHLQRIEELNRKIEEIMQQIADLGKNQPDVSRDIGKKQRQKLLSVKNATSELVAMFEQEIRETLTKFQKVLAMFQSVSRGADEKRVAFDRAKSRVVNSCNGQLRTDYTFSTSSIADDENTAPDAIVDGDFEVFK

>C_panamensis_HIM10

MAMQKPVVLMPMEKAPLIKLLNQKLQLGITLETLTTPTADLAQKMYRTFVRQILNVSESCLSTLPLSADCDHDPELHRNSIPIIIVYQCMKAFIKDHSGDKLDLTMCDLIQPHRVPGRFKKMATFLADYMRFHEIGNPVFNEISEEFSYQKQEVEMLHIELQNEESRKNGLLANQNLRKRREHELINEHNKVKTEFGNMVAQYEASHAAAEALNKEKEEALEETDRMEMEIISGKKMVDHLKEEVLSSPEELKVEMANRKKLIEELKDNLKATKKSYTERMEAIEICASFEKNKLMIEEKFRQFAMVKDEIMELLDAENENQRKLDDMEVEHRYMVEQRKNMHELLEEKALNHAQLRKEHLQRNEELNKKIEEITKKIAALGKNQPDVSRDIEKKRQELLAVKNCNSEIVAKAEHETREKLAKFQKVQTAFLKVHRNAEEKKTACERRISRVVNASVGRFSDHTFSTSSINEDSENTAPDSLQFNVFK

>C_nouraguensis_HIM10_paralog1

MAQKPVVLVALDKAVICKILKPKLHLGQLTPEDINNPSSEIAQQIFSNFVRYTLNVSESCMSTLPLSATSDHDPELHRRSIPIIIVFQCLKAFIKDHSGDKLDLTMCDFVNPQRINGRFKKITSFLADYIRFHENAQPIFNEVSEEFSYQRQEEQQLNEELQEEEKRKEMLISQQNARKRKDNELCNEHIKLKEELMNFMVACDEKKAFVEALFTEKEATEEKTESIENEILSGKKMVDHLKEEILSSPEELKREMAARKKQIEELKECLAGSKLALAERMEAIEICSSAEKNVPAIQEKINQFAMMKEEILELLDAVNEDNRKLSDLEDELKFTEEKKIKTRELMVDSAQLHEQVRKEHLQRNEQLNEKIKEITQISQMGTNQPDVSRDIEKKQELRAAKNATSKAVAKVVEETRETMAKYEKVLAMFQKVQRDAVEKQVAADRATSRLVNACVGPLISDYTFSTSSCCAEDDENTAPGVGTNFNVFPK

>C_nouraguensis_HIM10_paralog2_pseudogene

DKEAEMELRKKVLEERANKSVDDKETRALIRDMMENEAELKNARNEHEKLRLKLQQMEKKLIVGGENLLEKVEEQAKLLEISNREMESSKQSEERLRSQLEEKTAWKVEIEERYSSLQEESAAKTRKTKRVTNELREVRMELKDVEEEHQRQLEAMLEDARQLRK

>C_becei_HIM10

MAQKPVVLVTLDKAVICKILKPKLQLGHLTPEDINNPTEIAQQIFSNFVRYVLNVSESCMSTLPLSASCDYDPELHRRSIPIIIVFQCLKAFIKDHSGDKLDLTMCDFVNPHRINGRFKKITSFLADYIRFHENAQPVFNEISEEFSYQRQEEEQLNEELQEEEKRKEMLISQQSARKRKENELYNEQIKLKEEFSAIVGKCEEKKVFSETLFAEKVAAEEKTEGIENEILSGKKMVDHLKEEILSSPEELKQEMTARKKQIEELKECLAGSKLALAERMEAIEICTNVEKNVPAIQEKINQFVIMKEEIIELMDAVNEDNRKLNDLEDELKFTEEKKIKTRELMVDSAQLHEQLRKEHLQRNEQLNEKIKEITQISAMGKNQPDVSRDIEKKQELRAAKNATSEAVAKVVEETRETMAKYEKVLAMFQKVQRDAVEKQVAADRATSRLVNACIGPLISDYTFSTSSCCAEDDENTVPGGGTDFNVFPK

>C_yunquensis_HIM10

MAAQKPVVLVTLDKAVICKILKPKLHLGSLTPEDIMSPSAEIAQLVFSNFVRHTLNVSETCMTTLPLSATCDHDPELHRRSVPIIIVLQYMNAFIRDHSGGKLDMTMCDLIHPERIPGRFKKLTSFLADYIRFHENAQPVFNEISEEFSYQKLEEEQLRKELEEEEKRKASLTSQQNLRKRRENELRNELAKVKSEMFDIVATCESKKATFATLLSEKNAAVEETARIESDILSGRKMVDHLKEEVLSSPEELKLEMAARKKQIEELKECLKGSKQALVERIEAIEICASAEKNEPVILEKINQFAAMKEDIIELLDAVNEDNRKLSDLEDELKFTVEKKNNVHELMGEKAQLHAQLRNEHLQRTEQLNEKINEITKQIAAMGTNQPDVSREIEKKSQELRDAKNANSEAVAAVIQETRETMAKYNRVLAKFQQLQKDAAEKQVAAERATSRLVNACVGKLLTDYTFSTSSICTEEDENAAPDGQSFNVFQK

>C_macrosperma_HIM10

MASQKPVVLVPMDKVPICKILNQALQLGLTPDNITNPTAEIAQQVYINFVRLILNVSESCLSTLPLSANCNYDPELHRRSIPIIIVYQCKAFIKDHSGDKLDLTMCDLVQPQRVQGRFKKLATFLADYIKFHENGQPVFNEISEEFSYRKLEVEQLQLAIEDEERRKNDLLSQQNLRKRREHELINEHNKVKSEFSGVVGQYEANKITAGELLKQNEEAVEQIEQVENEILRSRKMAEHLKEELLSSPEELRLEMAARKKQIEELNECLKGSKVALAERMEAIDICINVEKNTPAVNEKLKMWAAVKEEIIMLFDAVNEDHRKLSDLENEQRFTAEKKKNVHELIGEQAQRHAQLQKEHLQRNVELNNKIDEITKQIAALGKNQPDVSRDIEKKRQELLAVKNANSESVAKVAQECREMFTKYQNVLARFQEVRRDANEKQVAVDRTTSRVVNACDGRLPTDYTFSTSSCCTEDGENITPDTQLRTDFNVFK

>C_sulstoni_HIM10

MASVRPVVMIKHDFKSISRNLNAKLHLNTRPEDISNPTVAELAQNVYMNFVRLILNVPDHSLSILPINSNVDYDPDQHRNSVRLIIVYQCMKAFVTDHSGRAFDLSMCDLVLPGKIPGRFQKLMSFLVDFMKLYQLAKPIFSEISEEFSYRKQEVEELKQALYEEEKRKKEMLAQQSLRRRREHELIDEHAKVTHELNGIVQQYTASTTRATELEKQKEEAIQSIERLESETLSGKKMVEHLNEEFLASPEELKREMAERKKQIEELTECRNSSKAKLAALEICRHIEKNFPASAEKIKIFRNVRAEILKLFDAVNENLRQLTDLEQELKFTTEKTKKSHEMMEEQAEMHKQLRNEHLQRSRELDSRIEEISREIAAMVKNQPDLSRDIENKRQELLVFKNEHSQTVSRILRHCEDLLVKYRKVHAMFEETQRTAQEKKTAGERAKGRVRTACFGRLPTDYTFNTSSLNEEKDENCRPDENFTVFK

>C_afra_HIM10

MASVRQIVLIKYDVKQISKVLNAKFQLGTKPDDIIKPSAELAQNIYMNFARLVLCIPDHSLTTLPISAYTDFDQDQHRNSVRLSLVYQCKAFIVDMSLGALSLSMCDLVVPDRTPGRFQKLMSFLVDFMKFHQVAEPTFSEISEEFSHRKKEVEELKQLLYEEERRKSELIAQQSLRKRREHELIDEHTRVNNELSGIIQQYTANTTTAGELDKQKEEALLTIERLEMETISGKKMVEHLNEEFLTSPDELRREMAERKRQIEELTECRNSAKEICRNIEKNFPASSEKIKTFQNVRSEIVKLFDAVNGNLRQLEDMEQELNFTTEKTRKSHEMMEEQAEMHKQLRKEHLQRSDELDARIEEITREIAAMGKNQPDLSREIEKKRQELLAFKNAHSQTVARILRHCEDLLAKYRKVHAMFEETRRNAEEKRIAGERAKGRVLNACSGRLPTDYTFNTSSVNEDLDENCRPDENFTVFN

>C_sp49_HIM10

MASEKEKRTVVLTMYDPNTIAKVLSSKLKLGLTPDDIINPTVRPTAFLVFQCFVRHVLDVSDASMSSLPLSAQSDEMDHDSHKRTIQLAIVYQCLKAFIADNSGNKIILSMCDIVQPAAVQNRFKRITSFLVDFIRLHDTALPIWDEIREEFSDRKHEVQSLQSDLITEEKRKNALLTQQSQRKRREHDLINDHNKVKQELTTTIDQYQANLAELEELKKKEKETNEETERLGTEILSGRKMVEHLSEELLTDPEELKKEMKTRKKEIEEMRGRLLNAKNILQKREEEIKICAEADRNEILFNEKLDVWEQIKADIVSLREEISDNVRTLSELEEKLKLTCEKRKLVAERMKEQAETQNQLRQQNSDRNEELQNKIDEVTEEIAALGRNQPDVSRIIEKKRQELLAVKNAMSSKEAECANSCKETLLKLRRMEAMFEELHRVSLEKRTAADRARYRVKTACVGNSMADYTFTSESIDENNPPNP

>C_sp25_HIM10

MTSEKEKRSVVLTIYDPNTIAKILNAKLKLGLTPDDIINPTHATAFQVFQSFVRHVLGVSDAAMNSLPLAAQSDDFDHDSHRKTIQLGIVYQCLKAFIADNSGQKIILSMCDIVQPAAIQHRFKKVTSFLADFIKLREVALPIWDEIREEFSDRNHEVQSLQGDLIAAEKRKNALLSQQSQRKRREHELINEHNKVNQELTNIVEQYTANMKEVEDRKKKKEETLNEIERLVSEILSGRKMVEHLGEEVLTSPEDLKNEMAGRKKQIEELKEHLIQARQSVQEKEEAIKICAEAERNEGVFNDKLDAWEKVKSDIVSIREEINENLRAFSELEEKLKLLSEKRKKVAERMQEQAETQNQLRQQNSERNDELQNNIDAVTEQIAALGKSQPDVSRIIEKKRQDLLAVKNAMSANESECANSCKETLFKLRKMEAMFDELHRVSLEKRTAADRARCRIKTACVGNRMADYTFSTESIDENNPPNSSFNIFK

>C_imperialis_HIM10

VHLYMYDARLIAKVLSSKLKLGLTPDDITNPTSEVAIQVFTNFVRFVLDVSETSLTSLPLTAQVDDLDYESHRKTIPLVIVYQCLKAFVSDNSGKKLILTMCDFVNPAAIQNRFKKVTSFLVDFIKLHGHALPIWDDIRDEFSDRKHEVQSLQNDLVTEEKRKNALLSQQSQRKRREHELINEHNKVNTELTNVVGQYTANMEEVEERKKKKEEAYDKIERLTNEIISGRKMVEHLGEEVLSSPEELKNEMAMRKKQIEELREHLAQARKALQEKDEAVKICTEAERNEVVFNEKLVMWDQVREDIVTIREEINENLRALSEYEEKLKLTIEKRKIVGERMREQSEQQEEMREQHSERNKQLQLKIDAVTEQIAALGKNQPDVSRDIEKKRQELLAVKNMMSEKEAECANSCRETLLKLRKMETMYEELFRVALEKRTAADRAACRVKNACAGVSMADYTFNTESIDENNPPPHTSFKVFN

>C_japonica_HIM10

MASGRPVLLTILDMRTILRVLNGKLHLGLTQENILTPTAEVAQQVFYNFVRYVLSVPESSLTTLPLTADVDVDNEMNRKSIPLVIVYQCMKAFIKDNTGGKLDLTMCDFVTPAKIQNRFKKLTSFLADFIRLHDMAMPLWNEISDEFGYRKHELESLQSEVMAVEKRKDDLLAQQSLRKRREHELINEHNKVKSELNKIIGQYNSNKSAAEERSKQKEEAIELIEKVENDVISGKKMVEHLSGEVLSSPEELKAEMEARRKQIEELRDCLRHSRKTLQNKEEALKICAEAEKNVPVLIDKINSWSELQEEIAELIDVINDNMRKLAELEENLQLTIEKKKKVGERMDEQAKLQTQLRRQHFQRNEDLQHKIEEITAEISALGKNQPDVSRDIERKRQELLLVKNALSEDIAELTNWCHESMSKFRKVQELFGETHRIALEKQTAGKRAKHRVRNAIFGPLPTEYTFDYTKTLSMDENDVGGNGSSVDFKVFK
